# Cyclin-G-associated kinase GAK/dAux regulates autophagy initiation via ULK1/Atg1 in glia

**DOI:** 10.1101/2021.08.16.456579

**Authors:** Shiping Zhang, Shuanglong Yi, Linfang Wang, Shuhua Li, Honglei Wang, Li Song, Jiayao Ou, Min Zhang, Ruiqi Wang, Mengxiao Wang, Yuchen Zheng, Kai Yang, Tong Liu, Margaret S. Ho

## Abstract

Autophagy is a major means for the elimination of protein inclusions in neurons in neurodegenerative diseases such as Parkinson’s disease (PD). Yet, the mechanism of autophagy in the other brain cell type, glia, is less well characterized and remains largely unknown. Here we present evidence that the PD risk factor Cyclin G-associated kinase (GAK)/dAuxilin (dAux) is a new component in glial autophagy. GAK/dAux directly interacts with ULK1/Atg1 via its uncoating domain. Lack of GAK/dAux increases the autophagosome number and size in adult fly glia and mouse microglia, and generally upregulates levels of components in the initiation and PI3K class III complexes including ULK1/Atg1, demonstrating that GAK/dAux regulates the onset of glial autophagy. Consistently, lack of GAK/dAux enhances Atg1 and Atg9 trafficking to autophagosomes, promoting autophagy initiation. On the other hand, lack of GAK/dAux impairs the autophagic flux and blocks substrate degradation, suggesting that GAK/dAux might play additional roles in glial autophagy. Our findings identify a new autophagy factor in glia; considering the pivotal role of glia under pathological conditions, targeting glial autophagy is potentially a therapeutic strategy for PD.

## Introduction

As the primary catabolic mechanism, macroautophagy (henceforth referred to as autophagy) was initially characterized as a bulk and non-selective degradation pathway induced by nutrient deprivation (1-3). Constitutive autophagy at the basal level, however, contributes to intracellular homeostasis in non-starved cells by degrading cellular materials like damaged organelles or misfolded proteins (4, 5). Interestingly, suppression of basal autophagy in the mouse nervous system results in the accumulation of ubiquitinated proteins and neurodegeneration (6, 7), whereas enhancing basal autophagy in the *Drosophila* nervous system promotes longevity and oxidant resistance (8). Furthermore, disease-associated mutations in core autophagy genes have been identified, supporting the link between autophagy and PD (9, 10). These findings mainly focus on a role for autophagy in neurons, as perturbation of such process leads to catastrophic consequences including pathological protein aggregation, deleterious lesions, and cell death (11). Whether autophagy in glia, the other major brain cell type, is executed via similar interplay using the same components as in neurons or other tissues, and whether it drives or contributes to disease pathology, remain largely unclear. Studies from yeast and mammals provided fundamental insights into *de novo* autophagosome biogenesis mediated by the evolutionarily conserved autophagy-related (ATG) genes (4, 5, 12). Upon autophagy initiation, stress induces the recruitment of ATG1/UNC-51 like autophagy activating kinase 1 (ULK1) complex to the phagophore assembly site or the pre-autophagosomal structure (PAS). The ATG1/ULK1 complex facilitates the assembly of phosphatidylinositol 3-kinase (PI3KC3) class III complex for PI3P production and promote the nucleation of the isolated membranes (also termed phagophores) (13). Whereas phagophore expansion is achieved by two ubiquitin-like conjugation steps: the formation of ATG12-ATG5 conjugate and ATG8 lipidation, additional lipids and proteins contributing to membrane expansion are delivered by the ATG9-containing vesicles derived from the Golgi apparatus, recycling endosomes or the plasma membrane (14-18). The expanded phagophores are sealed into a double-layered autophagosome, which eventually fuses with lysosome to form autolysosome, the milieu for degrading autophagic cargoes by acidic hydrolases.

Here we present evidence that the PD risk factor Cyclin G-associated kinase (GAK)/dAuxilin (dAux) is a new component in glial autophagy. GAK/dAux interacts with ULK1/Atg1 via the J domain with uncoating activity. GAK/dAux regulates the protein levels of ULK1/Atg1 and other components in the initiation and PI3K complexes controlling autophagy initiation. *In-vivo*, Lack of GAK/dAux increases the autophagosome number and size, and regulates the trafficking of ULK1/Atg1 and Atg9 to the autophagosomes. On the other hand, lack of GAK/dAux impairs autophagic flux and substrate degradation. Taken together, our findings identify a key factor regulating glial autophagy via controlling the activity of the initiation regulator ULK1/Atg1.

## Results

### Lack of dAux selectively increases the number of autophagosomes in adult fly glia

Identified in a behavior screen, the *Drosophila* homolog of the PD risk factor GAK, dAux, has been implicated in a broad spectrum of parkinsonian symptoms (19). GAK/dAux is a serine/threonine kinase that regulates clathrin-uncoating, cell cycle progression, and hepatitis C entry (20-23). Whereas another closely-related homolog for *GAK* (*DNAJC26*), *AUXILIN* (*DNAJC6*), is dominantly expressed in neurons, there is only one *Drosophila* homolog *dauxilin* (*daux*) that exhibits a higher homology with GAK (GAK 41.66%, AUXILIN 34.52%). GAK and dAux are both ubiquitiously expressed and shares a conserved kinase domain. To investigate dAux function at the cellular level, an RNAi line targeting *Drosophila daux* (*daux-*RNAi, V16182) was expressed to reduce *daux* expression in neurons or glia by the driver *GMR57C10-Gal4* or *repo-Gal4*, respectively. qRT-PCR, western blot (WB), and immunostaining analyses indicated that *daux* expression was efficiently reduced by the RNAi (Figures S1A-S1D). Under these conditions, the morphology, distribution, and number of the autophagic organelles labeled by the marker mCherry.Atg8a were analyzed in different adult brain regions, including the region “Glia” enriched of glial processes, glial nuclei, and neuronal nuclei, and the region “Antenna Lobe, AL” enriched of axon neuropils in close proximity to glial processes (Figures 1A and 1B). Interestingly, downregulating *daux* expression by *daux-*RNAi pan-neuronally did not cause severe change in the number or size of the Atg8a-positive puncta (Figures S1E and S1F). Instead, downregulating *daux* expression in glia caused a significant increase in the number of Atg8a-positive puncta (Figures 1C and 1D). These results, confirmed by one additional *daux*-RNAi (*daux*-RNAi^#2^, BL39017), an endogenous protein trap *pmCh*.*Atg8a*, anti-Atg8a antibodies, and Mosaic Analysis with a Repressible Cell Marker (MARCM) clones of *daux* mutant carrying EMS point mutations *I670K, L8H*, or *G257E* (24) (Figures S1G, S1H, and 1E-1J,), indicated an increase in glial autophagic structures in the absence of dAux. Results from using a different control *UAS-luciferase-*RNAi (*luc*-RNAi) were also consistent (Figures S1I and S1J). Ultrastructural analysis by Transmission Electron Microscopy (TEM) also showed an increase in the number of autophagosomes, which exhibits typical double-membrane structure with a characteristic cleft in-between and contains non-degraded cytoplasm, upon *daux*-RNAi expression in glia (pink asterisks, Figures 1K and 1L). Moreover, whereas the Rab7-positive late endosomes appeared in clusters with puncta number unaffected (Figures S1K and S1L), the formation of the Rab5-positive early endosomes remained largely intact upon glial dAux depletion (Figure S1M). Notably, expression of the fly dAux or human GAK in glia rescued the *daux-*RNAi-induced increase in the autophagosome number (Figures 1O and 1P). Taken together, these results indicate that dAux regulates glial autophagosome number and the function of GAK/dAux is conserved across species.

**Figure 1.**
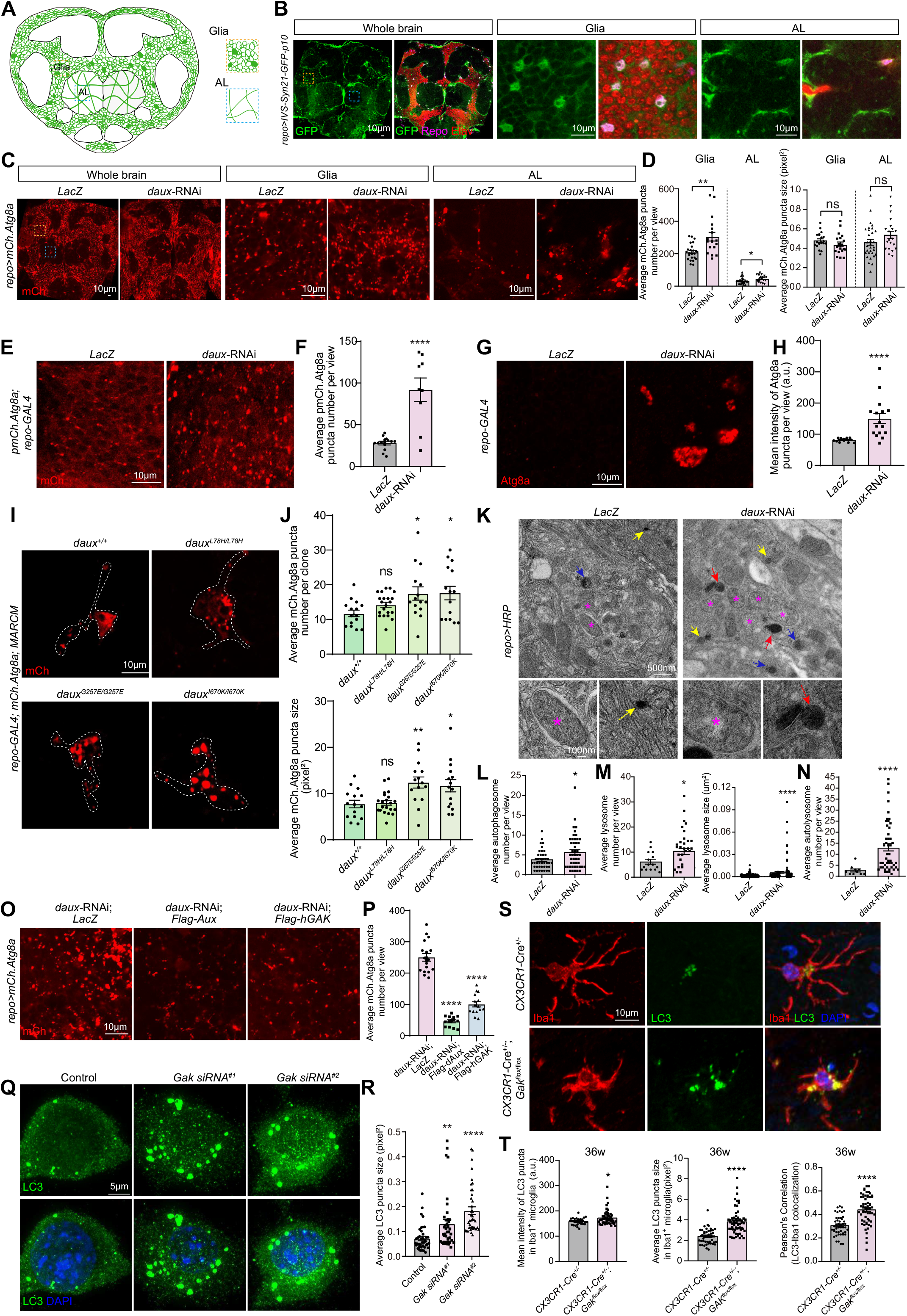
Lack of Gak/dAux increases the number and size of autophagosome in glia. (**A** and **B**) Illustration (A) and representative images (B) of an adult fly brain showing the expression pattern of glial cells. *repo-GAL4* drives the expression of *UAS-IVS-Syn21-GFP-p10* (*repo>IVS-Syn21-GFP-p10*) that labels both membrane and nucleus of glia. Dashed squares in yellow and blue indicate the relative positions of “Glia” and the antenna lobe (“AL”) assessed throughout the study. The regions “Glia” are enriched with GFP-positive glial membrane (green), GFP- and Repo-positive glial nuclei (green and magenta), as well as Elav-positive neuronal nuclei (red). The regions “AL” are enriched with axon neuropils and glial processes (green). (**C** and **D**) Representative images (C) and quantifications (D) of autophagosomes in the Whole brain, “Glia”, and “AL”. Glial autophagosomes (red) were labeled with *repo-GAL4* driven *UAS-mCherry*.*Atg8a* (*UAS-mCherry*.*Atg8a; repo-GAL4*). Note that downregulating glial *daux* induced an increase in the autophagosome number in both “Glia” and “AL”. (**E**-**H**) Representative images (E and G) and quantifications (F and H) of the “Glia” region in adult fly brains carrying an endogenous mCherry protein trap at the promoter of *Atg8a* (*pmCh*.*Atg8a*, E) or stained with the anti-Atg8a antibodies (G). Note that similar to *UAS-mCh*.*Atg8a*, the autophagosome number and fluorescent intensities in “Glia” region increased upon expressing the *daux*-RNAi in glia. (**I** and **J**) Glial MARCM clones of *daux* mutants (*daux*^*L78H*^, *daux*^*G257E*^, and *daux*^*I670K*^) expressing the *UAS-mCh*.*Atg8a* by *repo-GAL4*, and adult fly brain “Glia” region were shown. Dashed lines encircle the clonal area. Note that the number and size of glial autophagosomes per clone increased in *daux*^*G257E*^ and *daux*^*I670K*^ mutants. (**K-N**) Representative TEM images (K) and quantifications of autophagosome (L), lysosome (M), and autolysosome (N) in “Glia” region of control and *repo>daux-*RNAi adult fly brains. Note an increase in the number of autophagosome (pink asterisks), lysosome (yellow arrows) with the presence of large lysosome-like structure (red arrows), and autolysosome (blue arrows) when *daux-*RNAi was expressed in glia. (**O** and **P**) Co-expression of fly dAux or human GAK (hGAK) rescued the *daux*-RNAi-induced increase in glial autophagosome number in “Glia” region. (**Q** and **R**) Representative images (Q) and quantifications (R) of LC3-positive autophagosomes in IMG cells treated with either *Gak* siRNA. Note that LC3-positive autophagosomes were enlarged upon abolishing *Gak* expression. (**S** and **T**) Representative images (S) and quantifications (T) of LC3-positive autophagosome in mouse SN microglia deficient of *Gak* at 36 weeks. In addition to the increase in the intensity and size of LC3 in Iba1^+^ microglia, the colocalization of Iba1 with LC3 also increased upon microglial Gak depletion. Scale bars and the sample number n are indicated in the Figures. For each experiment, more than three biologically independent replicates were done. Whereas results were consistent, representative results from one experiment was shown. A serial confocal Z-stack sections were taken with 0.4 μm each, and representative single layer images acquired at the similar plane of brains across all genotypes are shown, except mouse brain slices (S) shown as maximal projection. For TEM, more than 40 images from at least 5 independent adult fly brains of each genotype were analyzed. Colocalization is analyzed using the Pearson’s Correlation and present as the Pearson’s R value. Quantification figures from the same images but with different parameters are shown under the same label unless noted otherwise. Data are shown as mean ± SEM. P-values of significance (indicated with asterisks, ns no significance, * p<0.05, ** p<0.01, and *** p<0.001) are calculated by two-tailed unpaired t-test, Mann-Whitney test, ordinary one-way ANOVA followed by Tukey’s multiple comparisons test, or Kruskal-Wallis tests followed by Dunn’s multiple comparisons test. All genotypes analyzed are listed in Table S1.

### Lack of Gak increases the number and size of autophagosomes in mouse microglia

In addition, the mouse immortalized microglial cell line (IMG) and the primary mouse microglia were examined for the effect of lacking Gak on autophagic structures. The IMG cells were stained positive for ionized calcium-binding adapter molecule 1 (Iba1) or transmembrane protein 119 (TMEM119), confirming their microglial identity (Figures S2A and S2B). Consistent with the previously reported expression in microglia and astrocytes (25, 26), Gak was also detected in the IMG cells (Figures S2C-S2G) (27). Treatment with either of the two siRNAs targeting *Gak* reduced its expression in IMG as shown by qRT-PCR, WB, and immunostaining analyses (Figures S2C-S2G). *Gak* mRNA and protein expression were also reduced down to 50% and 60%, respectively in *Gak* conditional knockout (cKO) mouse microglia (Figures S2H-S2J). We failed to collect *Gak* cKO primary microglia for WB analysis of Gak protein levels due to inefficient growth.

In IMG cells, treatment of either siRNA increased autophagosome size labeled by the microtubule-associated protein 1A/1B-light chain 3 (LC3) (Figures 1Q and 1R). Both the intensity and size of LC3 in microglia, and the LC3-Iba1 colocalization increased in *Gak* cKO microglia, suggesting increased autophagosome number in microglia (Figures 1S and 1T). Interestingly, the size of autophagosome consistently increases in the IMG, *Gak* cKO microglia, and fly *daux* mutant MARCM clones, but not *daux-*RNAi brains, possibly owing to the incomplete knock down by RNAi (Figures 1D, 1J, 1R, and 1T). Taken together, these results suggest that Gak regulates autophagosome biogenesis in mouse microglia.

### Lack of GAK/dAux promotes autophagy initiation

Based on the increase in the autophagosome number, it is conceivable that lack of glial dAux induces or blocks autophagy. Interestingly, Atg8a-II protein levels in adult fly brains downregulating glial dAux expression increased, which further increased in the presence of Bafilomycin A1 (BafA1), an inhibitor blocking lysosome acidification and autophagosome-lysosome fusion (28, 29) (Figures 2A and 2B). These results indicate that lack of glial dAux still triggers increase in the autophagosome number even when blocking the fusion, reinforcing a role for dAux in autophagy initiation. Using the anti-ULK1 antibodies that also recognize the *Drosophila* Atg1 proteins as validated by *atg1*-RNAi (Figures S3A and S3B), an increase in the number of ULK1-positive puncta was detected upon glial dAux depletion (Figures 2C and 2D). In addition, components of the initiation complex including Atg13, Atg6, and Atg14 were all affected as the number of mCherry.Atg13-, GFP.Atg6-, and GFP.Atg14-positive puncta increased upon glial dAux depletion (Figures 2C-2F). Notably, omegasomes labeled by two independent reporters: GFP.ZFYVE1 (12, 30) and EGFP.DFCP1 (31), exhibited significant increase in number upon glial *daux*-RNAi expression, with the size of EGFP.DFCP1-positive omegasomes became significantly larger as fused by multiple smaller rings (Figures 2G and 2H). Taken together, these results demonstrate the increased formation of autophagosome precursors in the absence of glial dAux.

**Figure 2.**
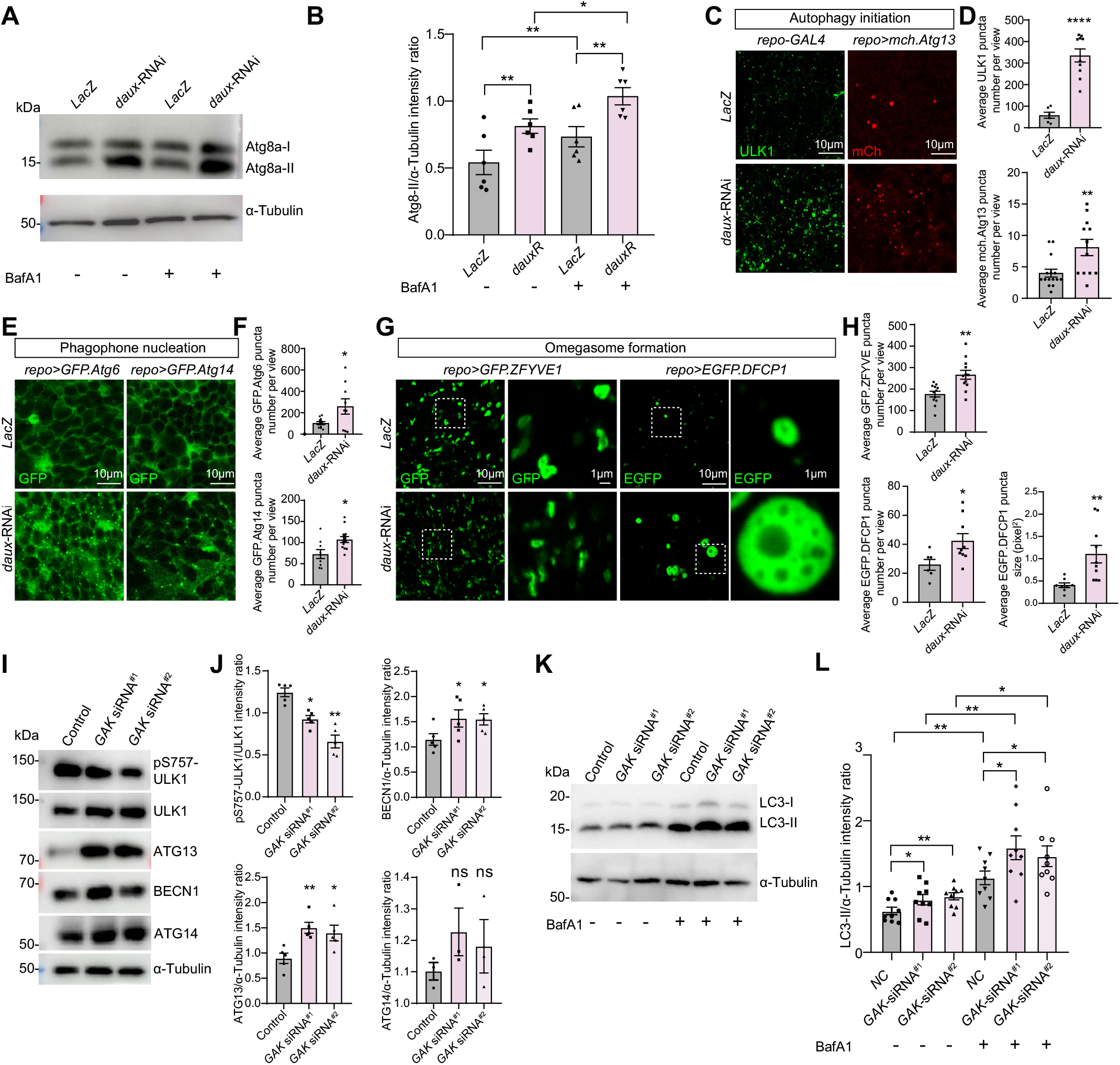
Lack of GAK/dAux promotes autophagy initiation. (**A** and **B**) Representative WB images (A) and quantifications (B) of Atg8a-II protein levels in the control and *repo>daux-*RNAi adult fly brains. Note that lack of glial dAux increased Atg8a-II protein levels, and BafA1 treatment further increased the Atg8a-II levels. (**C**-**H**) Representative images (C, E, G) and quantifications (D, F, H) of ULK1-, mCh.Atg13-, GFP.Atg6-, GFP.Atg14-, GFP.ZFYVE1-, and EGFP.DFCP1-positive puncta in “Glia” region of control and *repo>daux-*RNAi adult fly brains. *Drosophila* Atg1 was detected using the mammal anti-ULK1 antibodies. Note that lack of glial dAux increased the puncta number of all above, indicating a general upregulation in autophagy initiation, phagophore nucleation, and omegasome formation. (**I**-**L**) Representative WB images (I, K) and quantifications (J, L) of different autophagy protein levels in *GAK* siRNA-treated HeLa cells. Results from two independent siRNAs, *GAK* siRNA^#1^ and *GAK* siRNA^#2^, were consistent. Note that *GAK* knockdown significantly increased the protein levels of pS757-ULK1, ATG13, BECN1, and LC3-II. LC3-II protein level enhanced by *GAK* knockdown was further enhanced in the presence of BafA1. Scale bars and the sample number n are indicated in the Figures. For each experiment, more than three biologically independent replicates were done. Whereas results were consistent, representative results from one experiment was shown.A serial confocal Z-stack sections were taken with 0.4 μm each, and representative single layer images acquired at the similar plane of brains across all genotypes are shown. Colocalization is analyzed using the Pearson’s Correlation and present as the Pearson’s R value. Data are shown as mean ± SEM. P-values of significance (indicated with asterisks, ns no significance, * p<0.05, ** p<0.01, and *** p<0.001) are calculated by two-tailed unpaired t-test, Mann-Whitney test, or ordinary one-way ANOVA followed by Tukey’s multiple comparisons test.

To better understand the mechanism of dAux-mediated autophagy initiation, we turned to a different system, HeLa cells, for further characterization. A knockdown system was established using two independent siRNAs targeting *GAK* (siRNA^#1^ and siRNA^#2^). Treatment of either siRNA reduced *GAK* expression in HeLa cells as shown by WB and immunostaining analyses (Figures S3C-S3F). Interestingly, ULK1, ATG13, and BECLIN1 protein levels significantly increased while ATG14 protein levels exhibited an increasing trend upon *GAK* siRNA treatment, suggesting a general upregulation in autophagy initiation (Figures 2I and 2J). In addition, the mTORC1-mediated phosphorylation on the ULK1 Serine 757 (pS757-ULK1) decreased, suggesting that the ULK1 activity is enhanced as mTORC1 inhibition is relieved (Figures 2I and 2J). In addition, a *GAK* KO cell line was created by silencing the *GAK* expression in HeLa cells using CRISPR-Cas9 (Figures S3G-S3J). Consistent to the observations in flies, lack of GAK causes an increase in LC3-II protein levels in both *GAK* siRNA-treated and KO cells (Figures 2K and 2L, S3K and S3L). LC3-II protein levels also further increased in the presence of BafA1 in *GAK* siRNA-treated cells, reinforcing a direct role of GAK in autophagy initiation (Figures 2K and 2L). Taken together, these results indicate that autophagy initiation is induced in the absence of GAK/dAux; GAK/dAux regulates the onset of glial autophagy.

### GAK/dAux interacts with ULK1/Atg1 and regulates its protein levels

As shown in flies and HeLa cells, lack of GAK/dAux enhances ULK1/Atg1 activity and induces autophagy initiation. We next tested if ULK1/Atg1 acts downstream of GAK/dAux in regulating autophagy initiation. Co-immunoprecipitation (Co-IP) analysis using fly S2 cells expressing Flag-dAux and Myc-Atg1 revealed a direct interaction between dAux and Atg1 when pulling down Myc-Atg1 with the anti-Myc beads. Reverse pull down by Flag-tagged dAux also revealed similar interaction (Figure 3A). To determine which dAux domain is responsible for Atg1 interaction, different dAux deletion variants were made and expressed in flies for testing their interaction with Atg1. All variants expressed properly except the one lacking PTEN domain (dAux^ΔPTEN^, Figures 3B and 3C). Co-IP analysis showed that Atg1 interacts with most of the dAux variants except the one lacking J domain (dAux^ΔJ^, Figure 3D). In addition, overexpression of dAux^ΔKinase^ or dAux^ΔJ^ enhanced ULK1 levels (Figures 3E and 3F). Importantly, the early-onset PD SNP mutation in the J domain of AUXILIN (DNAJC6) (32, 33) is conserved in dAux, and overexpressing the dAux carrying this mutation dAux^DNAJC6m^ enhanced ULK1 levels (Figures 3E and 3F). These results implicate that dAux kinase and uncoating function are important for regulating the ULK1/Atg1 protein level.

**Figure 3.**
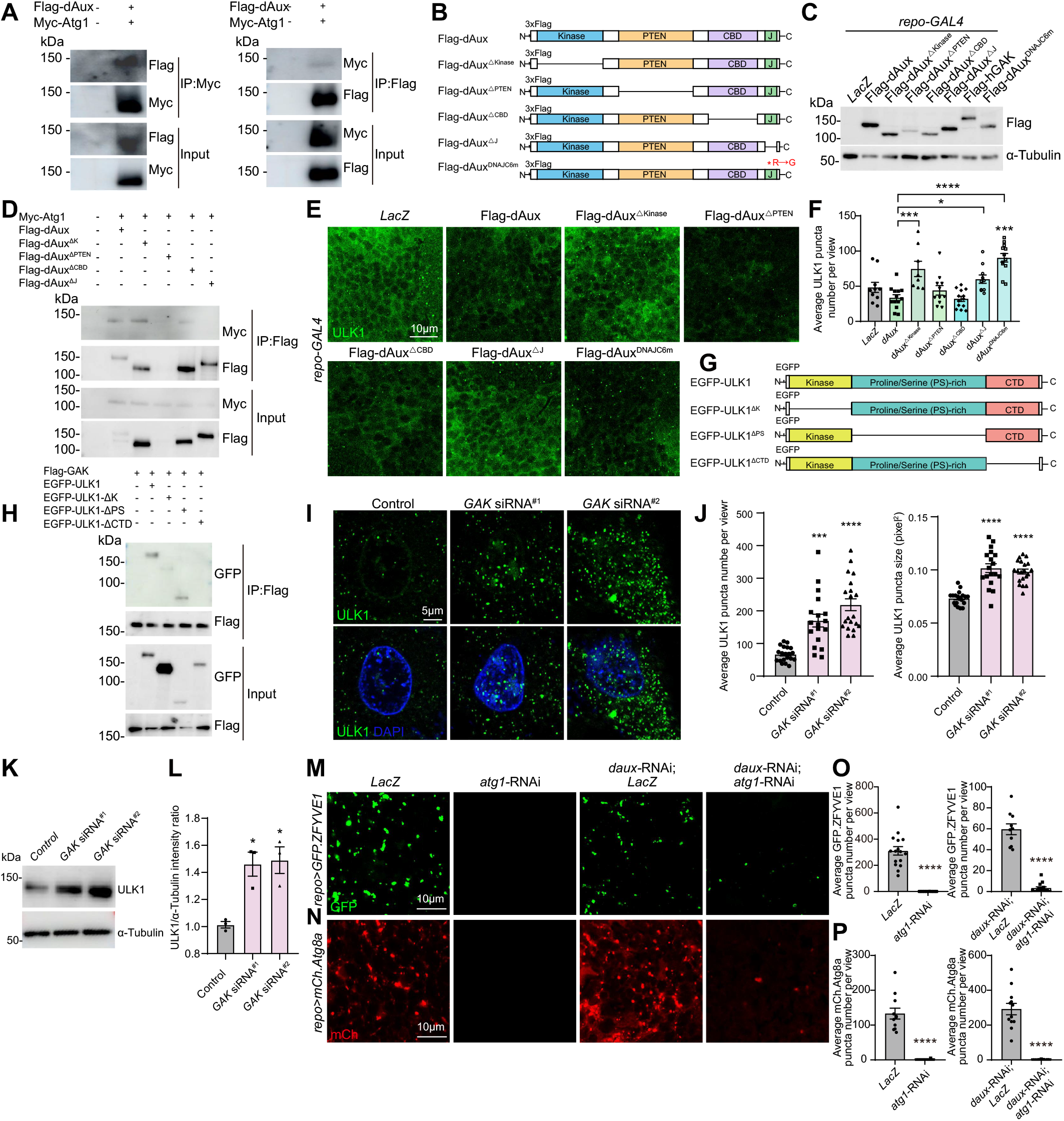
GAK/dAux interacts with ULK1/Atg1 and regulates its protein levels. (**A**) Co-IP analysis using fly S2 cells expressing Flag-dAux, Myc-Atg1, or both revealed a direct interaction between dAux and Atg1. dAux or Atg1 was detected in the pull down of Myc-Atg1 or Flag-dAux, respectively. (**B** and **C**) dAux variants lacking different domains or carrying an early-onset PD SNP mutation (DNAJC6m) were constructed and expressed in fly, adult heads were used for WB to check the expression level. Note that dAux^ΔPTEN^ expressed poorly. (**D**) Different dAux variants were expressed in S2 cells, and analyzed for their interaction with Atg1. Note that J domain was required for dAux-Atg1 interaction. (**E** and **F**) Representative images (E) and quantifications (F) of the ULK1 puncta number in the “Glia” region of adult fly brains expressing different dAux variants. Note that the ULK1 puncta number increased when expressing dAux^ΔKinase^, dAux^ΔJ^, or dAux^DNAJC6m^. (**G** and **H**) N-terminal EGFP-fused ULK1 variants lacking different domains were constructed, expressed in HEK293FT cells, and analyzed for their interaction with GAK. Note that ULK1 lacking the C-terminal domain CTD failed to interact with GAK. (**I**-**L**) The representative immunostaining (I) and WB (K) images and quantifications (J and L) of the ULK1 protein level in *GAK* siRNA-treated HeLa cells. Note that *GAK* knockdown caused an increase in the number and size of the ULK1 puncta, also the ULK1 protein level. (**M**-**P**) Epistatic genetic analysis revealed that dAux acts upstream of Atg1 in regulating autophagy initiation. Co-expression of *atg1*-RNAi in glia rescueed and further suppressed the *daux*-RNAi-induced increase in the number of GFP.ZFYVE1- and mCh.Atg8a-positive puncta in “Glia” region. Scale bars and the sample number n are indicated in the Figures. For each experiment, more than three biologically independent replicates were done. Whereas results were consistent, representative results from one experiment was shown. A serial confocal Z-stack sections were taken with 0.4 μm each, and representative single layer images acquired at the similar plane are shown. Data are shown as mean ± SEM. P-values of significance (indicated with asterisks, ns no significance, * p<0.05, ** p<0.01, and *** p<0.001) are calculated by Mann-Whitney test, ordinary one-way ANOVA followed by Tukey’s multiple comparisons test, or Kruskal-Wallis tests followed by Dunn’s multiple comparisons test.

On the other hand, GAK interacted with ULK1 in HEK293FT cells, but failed to interact with ULK1 lacking the C-terminal domain (ULK1^ΔCTD^), suggesting that CTD is required for GAK-ULK1 interaction (Figures 3G and 3H). Consistent to the enhanced ULK1 activity, the ULK1 protein level was also elevated in *GAK* siRNA-treated HeLa cells (Figures 3I-3L). To further test the relationship between dAux and Atg1, epistatic genetic analysis between dAux and Atg1 was conducted. Silencing *atg1* expression in glia caused a dramatic decrease in the number of omegasomes and autophagosomes, reinforcing the master regulatory role of Atg1 in autophagy initiation (Figures 3M-3P).

Co-expression of *atg1*-RNAi and *daux*-RNAi rescued and further suppressed the *daux*-RNAi-induced increase in the number of omegasome and autophagosome, suggesting that dAux acts upstream of Atg1 in regulating autophagy initiation (Figures 3M-3P). Taken together, these results demonstrate that GAK/dAux interacts with ULK1/Atg1 and regulates its protein level.

### dAux regulates Atg1 independently of its kinase function

Given that both dAux kinase and J domains are implicated in the Atg1 activation, phosphoproteomic analysis was subsequently conducted using adult fly head samples collected from control and *repo>daux*-RNAi flies (Figure S4A). These results revealed significant differences on the phosphorylation profiles of autophagy-related proteins (Figure S4B). Nonetheless, no significant changes were detected for Atg1 phosphorylation when downregulating dAux expression in glia. Only minor alternations on the two Atg1 residues, S282 (RNAi/LacZ ratio 1.131) and T292 (RNAi/LacZ ratio 1.053), were identified (Table S2). In addition, our Co-IP analysis revealed that dAux^ΔKinase^ interacts with Atg1, indicating that the kinase domain is not essential for the interaction (Figure 3D). Despite that dAux^ΔKinase^ expression enhances ULK1 levels, its impact on autophagy remains similar to dAux when analyzing the protein levels of Atg8a-II, Ubi, and P62 (Figures S4C-S4F), suggesting that lacking kinase domain does not hinder dAux function in autophagy. Hence, it is unlikely that dAux regulates Atg1 in a kinase-dependent manner.

### Lack of GAK/dAux promotes ULK1/Atg1 trafficking to autophagosomes

One of the major steps in initiating autophagy is the ULK1/Atg1 transport to the autophagy initiation sites for forming the autophagosomes. We next analyzed the colocalization of ULK1/Atg1 with LC3/Atg8a as an indicator for the ULK1/Atg1 trafficking efficiency. Since an increase in the ULK1/Atg1 protein levels was also detected, we carefully chose the Manders’ Correlation method when analyzing the colocalization given that the change in protein level is also considered in this statistical approach (34-36). Consistent to the observations in flies and mice, the number of LC3-positive puncta labeled by the transfected RFP-LC3 increased in the *GAK* siRNA-treated or KO cells (Figures 4A and 4B, S5A and S5B). In addition, the ULK1/Atg1 trafficking to the autophagosomes was enhanced as shown by the increased colocalization of ULK1/Atg1 and LC3/Atg8a in flies expressing *daux*-RNAi, *GAK* siRNA-treated, and KO HeLa cells (Figures 4A-4D, S5A and S5B). In contrast, the colocalization of ULK1 with TGN38-marked trans-Golgi network (TGN) decreased, further suggesting that ULK1 trafficking to the autophagosomes is enhanced when lacking GAK (Figures 4E and 4F, S5C and S5D). In support of these findings, ULK1 trafficking to the omegasomes also increased as the colocalization of ULK1 with transfected GFP-DFCP1 increased in the *GAK* siRNA-treated (Figures S5E and S5F) and KO cells (Figures S5G and S5H). Taken together, these results indicate that GAK/dAux regulates autophagy initiation via ULK1/Atg1 trafficking.

**Figure 4.**
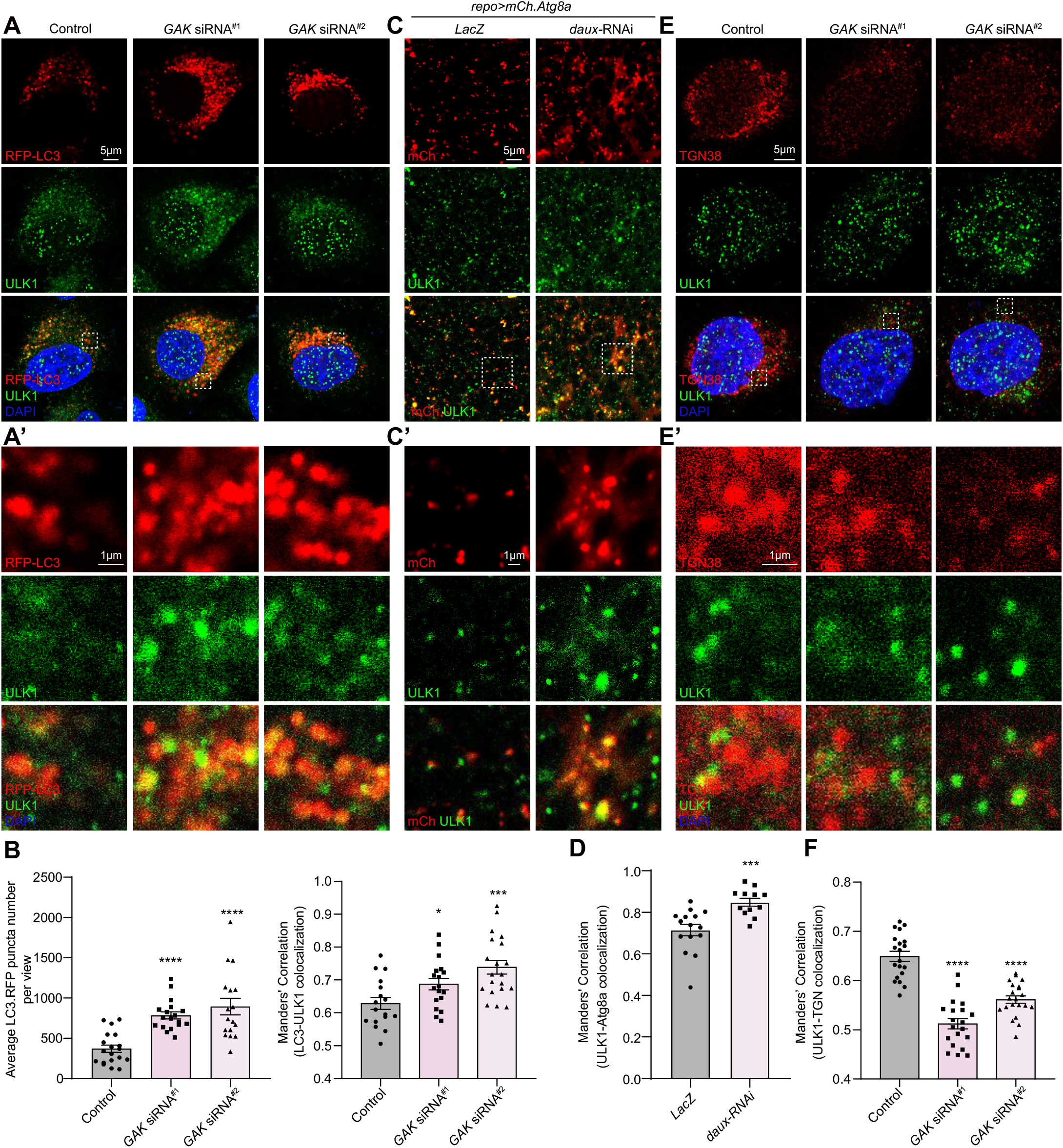
GAK/dAux regulates ULK1/Atg1 trafficking from TGN to autophagosomes. (**A** and **B**) HeLa cells transfected with RFP-LC3 in the presence or absence of the *GAK* siRNA were collected for immunostaining analysis with the anti-ULK1 antibodies. Note that both the LC3-positive puncta number and the ULK1 (green)-LC3 (red) colocalization increased upon GAK depletion. (**C** and **D**) ULK1-Atg8a colocalization was analyzed in “Glia” region of adult fly brains expressing mCh.Atg8a (*repo>mCh*.*Atg8a*) to label glial autophagosomes. Note that ULK1 (green)-Atg8a (red) colocalization increased in the absence of glial dAux. (**E** and **F**) ULK1 (green)-TGN38 (red) colocalization is analyzed in *GAK* siRNA-treated HeLa cells. Note that ULK1-TGN38 colocalization decreased, suggesting enhanced ULK1 trafficking to the autophagosomes upon GAK depletion. Areas enclosed by the white dashed squares in the representative images (A, C, and E) are enlarged in A’, C’, and E’. HeLa cell nuclei were stained with DAPI (blue). Scale bars and the sample number n are indicated in the Figures. For each experiment, more than three biologically independent replicates were done. Whereas results were consistent, representative results from one experiment was shown. A serial confocal Z-stack sections were taken with 0.4 μm each, and representative single layer images acquired at the similar plane are shown. Colocalization is analyzed using the Manders’ Correlation, taking into account the change in the protein level. Data are shown as mean ± SEM. P-values of significance (indicated with asterisks, ns no significance, * p<0.05, ** p<0.01, and *** p<0.001) are calculated by two-tailed unpaired t-test or ordinary one-way ANOVA followed by Tukey’s multiple comparisons test.

### Glial dAux regulates Atg9 trafficking to autophagosomes via Atg1

In addition to promote Atg1 trafficking, lack of dAux increased the glial Atg9-Atg8a colocalization analyzed by the *mCherry*.*Atg8a* and *EGFP*.*Atg9* reporters, indicating that dAux also regulates Atg9 trafficking to autophagosomes (Figures 5A, 5A’, and 5C). Next, live-cell imaging analysis was conducted, and the dynamic of Atg9 trafficking was analyzed by Imaris (Figures 5E-H, Supplementary Video 1 and 2). Interestingly, the speed of both Atg8a- and Atg9-positive puncta decreased upon glial dAux depletion, suggesting that these puncta are less mobile (Figures 5E and 5H). Notably, the duration of the Atg9-Atg8a contact was significantly longer, and the percentage of Atg9-positive puncta being trafficked to the vicinity of Atg8a-positive puncta (with a radius of 0.35 μm) was also significantly greater (Figures 5F and 5G). These results demonstrate that glial dAux regulates the dynamic of Atg9 trafficking to and its fall off from autophagosomes.

**Figure 5.**
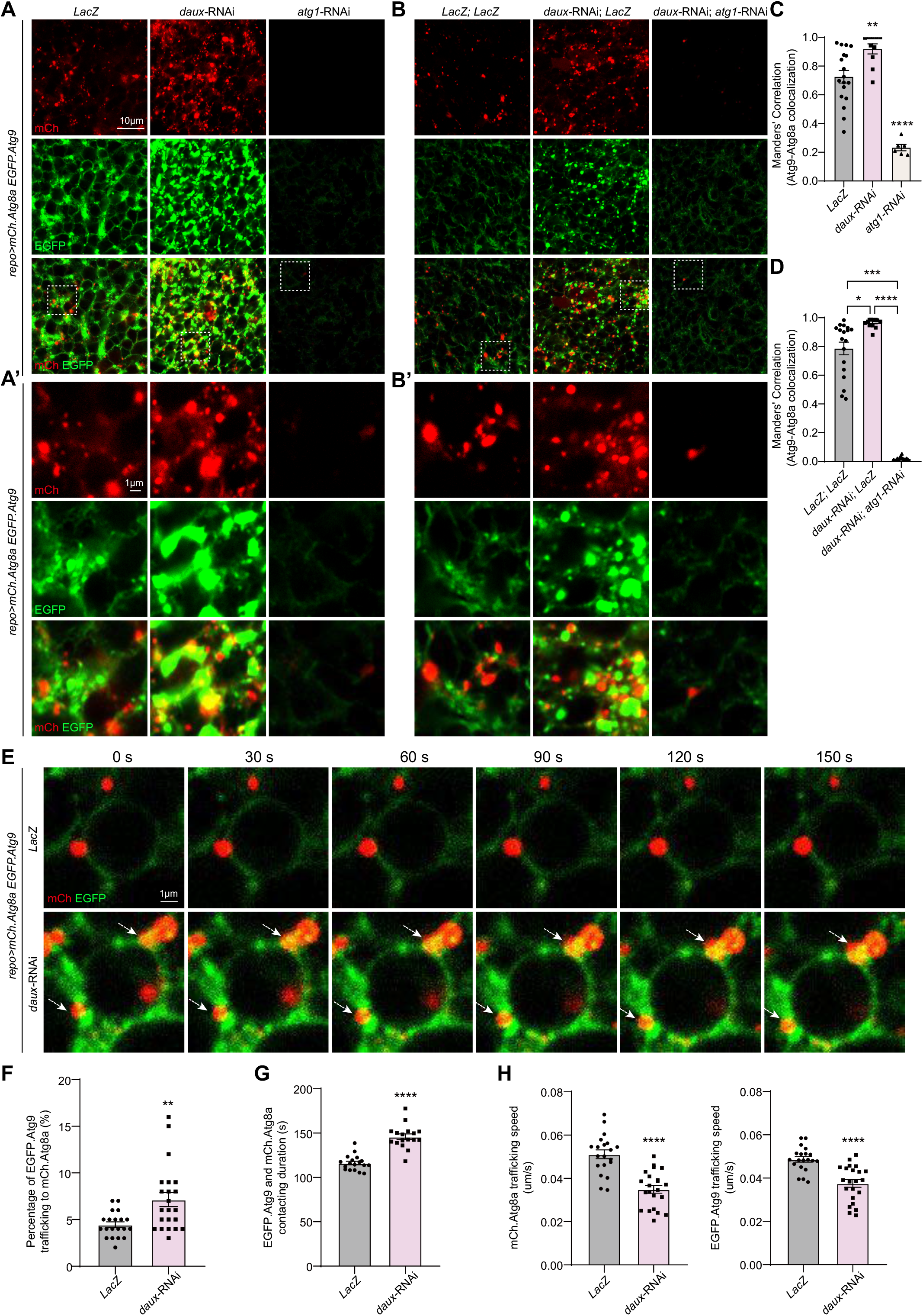
dAux-mediated Atg9 trafficking depend on Atg1. (**A**-**D**) Atg9 trafficking is investigated by analyzing the Atg9 (green)-Atg8a (red) colocalization in the “Glia” region of adult fly brains expressing the *UAS-EGFP*.*Atg9* and *UAS-mCh*.*Atg8a* reporters (*UAS-mCh*.*Atg8a; repo-GAL4, UAS-EGFP*.*Atg9*). Note that *daux*-RNAi or *atg1*-RNAi expression in glia caused a significant increase or decrease in the Atg9-Atg8a colocalization, respectively. Co-expression of *daux*-RNAi and *atg1*-RNAi rescued and further suppressed the increased Atg9-Atg8a colocalization. Areas enclosed by the white dashed squares in the representative images (A and B) are enlarged in A’ and B’. (**E**-**H**) Live-cell imaging (E) and quantifications (F-H) of Atg9 trafficking to Atg8a in “Glia” region of adult fly brains. Note the percentage of Atg9-positic puncta being trafficked to the vicinity of Atg8a-positive puncta (with a radius of 0.35 um) was increased (F), the duration of the Atg9-Atg8a contacting was longer (G), and the speed of both Atg8a- and Atg9-positive puncta were decreased upon glial dAux depletion (H). Scale bars and the sample number n are indicated in the Figures. For each experiment, more than three biologically independent replicates were done. Whereas results were consistent, representative results from one experiment was shown. A serial confocal Z-stack sections were taken with 0.4 μm each, and representative single layer images acquired at the similar plane are shown. Colocalization is analyzed using the Manders’ Correlation, taking into account the change in the protein level. Data are shown as mean ± SEM. P-values of significance (indicated with asterisks, ns no significance, * p<0.05, ** p<0.01, and *** p<0.001) are calculated by two-tailed unpaired t-test, Mann-Whitney test, ordinary one-way ANOVA followed by Tukey’s multiple comparisons test, or Kruskal-Wallis tests followed by Dunn’s multiple comparisons test.

Consistent to the notion that Atg1 regulates Atg9 phosphorylation and transport (37-39), lacking Atg1 abolished Atg9 trafficking to the autophagosomes as the Atg9-Atg8a colocalization decreased significantly (Figures 5A, 5A’, and 5C). These results suggest that dAux and Atg1 regulates Atg9 trafficking in opposite means. Interestingly, co-expression of *atg1*-RNAi and *daux*-RNAi rescued and further suppressed the dAux-mediated Atg9 trafficking (Figures 5B, 5B’, and 5D), suggesting that dAux regulates Atg9 trafficking via Atg1. Given that Atg9 is a downstream target of Atg1, it is likely sthat dAux-mediated Atg1 activation promotes Atg9 trafficking to the autophagosomes, hence initiating autophagy.

### Lack of GAK/dAux impairs autophagic flux and substrate degradation

To further demonstrate a role for dAux in glial autophagy, the autophagic flux was analyzed using a dual reporter *UAS-GFP-mCherry-Atg8a* in *Drosophila* adult brains. Whereas mCherry-positive puncta were GFP-negative and autophagy proceeded normally in the control animals, autophagic flux was impaired in the absence of glial dAux as both the GFP- and mCh-positive puncta colocalization and the intensity of GFP puncta increased significantly upon glial dAux depletion. In the presence of BafA1, the GFP- and mCh-positive puncta colocalization further increased upon depleting glial dAux and a similar trend of GFP puncta intensity was detected (Figures 6A and 6B). As these non-quenched GFP signals could represent either the non-fused autophgasomes or the fused but dysfunctional autolysosomes, we next checked whether the fusion of autophagosomes and lysosomes is affected by glial dAux depletion. Interestingly, analysis of Lamp1-Atg8a colocalization with Stimulated Emission Depletion (STED) imaging on adult fly brains expressing *daux*-RNAi revealed an increased number of enlarged and well-colocalized autolysosomes, indicating that lysosomes are able to fuse with autophagosomes (Figures 6C and 6D). Ultrastructural TEM analysis also showed an increase in the number of autolysosomes, which have only one limiting membrane and contain electron dense cytoplasmic materials and/or organelles, upon *daux*-RNAi expression in glia (blue arrows, Figures 1K and 1N). Reducing the expression of fusion factors such as *syx17* or *vamp7* caused a block in fusion (40), and co-expression of either RNAi reverted the *daux*-RNAi-induced increase in Lamp1-Atg8a colocalization (Figures S6A and S6B). These results suggest that some successful fusion occurs so that increasing number of enlarged autolysosomes is detected.

**Figure 6.**
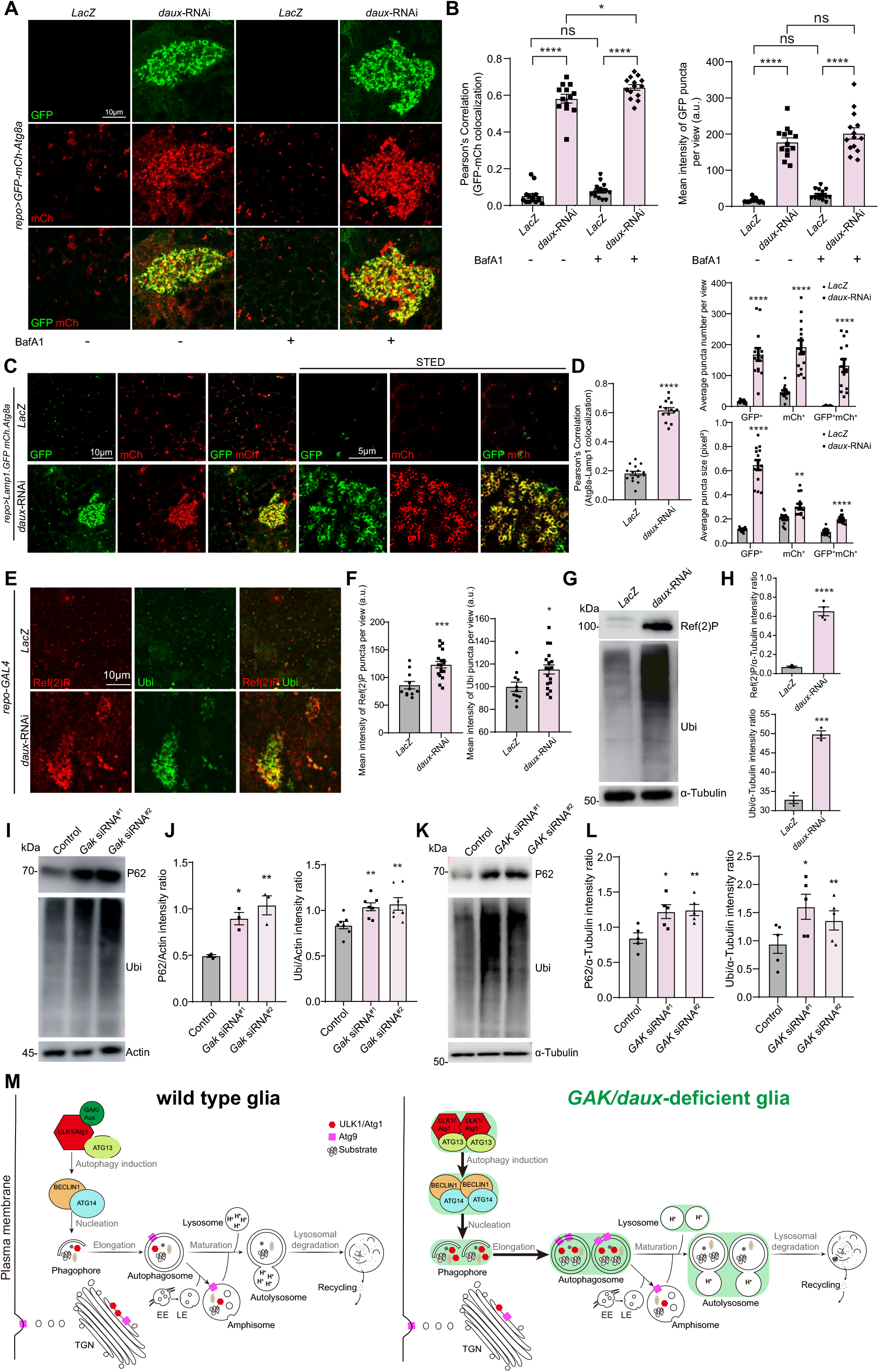
Lack of GAK/dAux impairs autophagic flux and substrate degradation. (**A** and **B**) Autophagic flux was impaired as examined by expressing the flux reporter *UASp-GFP-mCh-Atg8a*. Both the GFP- and mCh-positive puncta colocalization and the intensity of GFP puncta in “Glia” region increased upon depleting glial dAux. In the presence of BafA1, the GFP- and mCh-positive puncta colocalization further increased upon depleting glial dAux, a similar trend of GFP puncta intensity was detected but not statistically significant. Note that GFP- and mCh-positive puncta were enlarged and potentially dysfunctional due to the non-quenched GFP signals. (**C** and **D**) Representative confocal/STED images (C) and quantifications (D) of Lamp1- and Atg8a-positive puncta in “Glia” region of adult fly brains. Note that dAux depletion in glia increased the number and size of glial autolysosomes and the colocalization of autophagosomes and lysosomes. (**E**-**H**) Autophagic substrates such as Ref(2)P (the *Drosophila* P62 homolog) and Ubiquitin (Ubi) accumulated in *repo>daux-*RNAi adult fly brains as revealed by immunostaining (“Glia” region) and WB analyses. (**I** and **J**) P62 and Ubi protein levels increased upon *Gak* depletion in IMG cells. (**K** and **L**) P62 and Ubi protein levels increased upon *GAK* depletion in HeLa cells. (**M**) A model illustrating the regulating roles of GAK/dAux in glial autophagy. In *GAK/daux-*deficient glia, lack of GAK/dAux activates ULK1/Atg1, upregulates protein levels of components in the initiation and PI3P class III complexes, and enhances Atg1 trafficking to the autophagosomes (thicker black arrows). dAux also regulates Atg9 trafficking to the autophagosomes via Atg1. These actions promote autophagy initiation and start the glial clearance program for substrates degradation. On the other hand, despite the benefits of inducing glial autophagy, lack of GAK/dAux also disrupts autophagic flux and substrate degradation. GAK/dAux regulatory steps are highlighted in green. Scale bars and the sample number n are indicated in the Figures. For each experiment, more than three biologically independent replicates were done. Whereas results were consistent, representative results from one experiment was shown. A serial confocal Z-stack sections were taken with 0.4 μm each, and representative single layer images acquired at the similar plane of brains across all genotypes are shown. Data are shown as mean ± SEM. P-values of significance (indicated with asterisks, ns no significance, * p<0.05, ** p<0.01, and *** p<0.001) are calculated by two-tailed t-test or ordinary one-way ANOVA followed by Tukey’s multiple comparisons test.

As a consequence, inhibiting dAux expression in glia caused the accumulation of autophagic substrates such as Ref(2)P (the *Drosophila* ortholog of P62) and Ubiquitin (Ubi) (Figures 6E-6H). P62 and Ubi were also accumulated in *Gak* siRNA-treated IMG cells (Figures 6I and 6J), *GAK* siRNA-treated and *GAK* KO HeLa cells (Figures 6K, 6L, S3K, and S3L). Furthermore, increased Ref(2) puncta observed upon *daux*-RNAi expression in glia were well localized with autophagosomes and lysosomes (Figure S6C), suggesting that theses fused autolysosomes are able to recognize and encircle protein substrates, but failed to degrade them. Taken together, these findings suggest that GAK/dAux regulates glial autophagy via the initiation step but with an ultimate effect on impaired flux and substrate degradation (Figure 6M).

## Discussion

Our findings pinpoint a role for GAK/dAux in autophagy initiation. Despite that the ultimate consequence for lacking GAK/dAux blocks autophagic substrate degradation and a potential compensatory mechanism leading to the increased autophagosome biogenesis might exist, our evidence indicates that GAK/dAux functions directly on autophagy initiation by modulating ULK1/Atg1 activity and protein level. First, the dAux-mediated increase in the Atg8a-II/I protein levels is further enhanced by the presence of BafA1, indicating that autophagy is promoted even when substrate degradation is blocked. Lack of GAK/dAux also causes an increase in phagophore nucleation and omegasome formation, suggesting that these earlier steps in autophagy initiation are all affected. Furthermore, the levels of components in the initiation and PI3K class III complexes including ULK1/Atg1, ATG13, BECLIN1, and ATG14 all increase in the absence of GAK, indicating a general upregulation in autophagy initiation. It has been shown that mTOR senses the nutrient-rich condition and blocks autophagic induction by phosphorylating ULK1 at S757 to prevent its activation (41). In the absence of GAK, S757 phosphorylation is significantly decreased, indicating enhanced ULK1 activity. Thus, GAK/dAux suppresses autophagy initiation; it is conceivable that GAK/dAux acts as a glial sensor to control the onset of autophagy for sensing pathological signals like protein inclusions.

ULK1/Atg1 is a master regulator initiating autophagy. In complex with ATG13 and FIP200, ULK1/Atg1 initiates autophagy by phosphorylating a vast array of downstream components including the factors in the initiation and PI3P class III complexes (42-44). The ULK1/Atg1 activity undergoes intricate controls by post-translational modification including phosphorylation, acetylation, or ubiquitination (45, 46). These modifications can either activate or inhibit ULK1/Atg1 activity, thereby promoting or inhibiting autophagy, respectively. Despite that a direct interaction between GAK/dAux and ULK1/Atg1 is detected, our phosphoproteomic analysis reveals no significant alternation on Atg1 phosphorylation upon glial dAux depletion. In addition, dAux lacking the kinase activity remains associated with Atg1. Thus, it is unlikely that GAK/dAux regulates ULK1/Atg1 via phosphorylation.

Previous studies reported that inhibiting ULK1 kinase activity, either by ULK1 inhibitors or expressing a kinase-dead ULK1, caused accumulation of stalled early autophagosomal structures (47, 48). While we observe some phenotypes similar to stalled early autophagosomes, but with crucial differences. First, instead of lacking LC3-II flux, we observed increased LC3-II flux with/without BafA1 when downregulating GAK/dAux both in HeLa cells and fly. Immunostaining of dual autophagy flux reporter in adult fly brains with BafA1 got similar results. Second, the accumulated autophagosomes were able to fuse with lysosomes to form autolysosomes when downregulating glial dAux. Third, we observed substantial co-localization of autophagosomes, lysosomes, and Ref(2)P, suggesting that theses fused autolysosomes are able to recognize and encircle protein substrates, but failed to degrade them. Last but most importantly, we observed decreased S757 phosphorylation in the absence of GAK, indicating enhanced ULK1 activity.

As we demonstrated that dAux J domain is required for its interaction with Atg1, it is conceivable that dAux regulates Atg1 via its uncoating function. Expression of dAux lacking the J domain or dAux carrying a conserved early-onset PD SNP mutation in the J domain causes an increase in the ULK1 protein level. These results implicate the relevance of dAux clathrin-uncoating activity with ULK1/Atg1. In addition, it is also possible that GAK/dAux contributes to the recycling of ULK1/Atg1 from autophagosomes to TGN, thus leading to similar enhancement on autophagy initiation. To this end, our live-cell imaging analysis supports the notion that dAux plays dual roles in the context of Atg9 trafficking to autophagosomes. As increased amount of Atg9-positive puncta is detected at the vicinity of Atg8a-positive puncta, the contact duration time between Atg9 and Atg8a is also significantly enhanced in the absence of dAux. Thus, dAux might be needed for both the anterograde trafficking to and the recycling from autophagosomes. On the other hand, our results support a competition model for how GAK initiates autophagy. GAK interacts with the C-terminus of ULK1, the domain also required for its interaction with Atg13 (49). It is reasonable to spectulate that GAK competes with Atg13 for ULK1 binding. In the absence of GAK, increased amounts of ULK1 proteins are relieved from the inhibition for interaction with Atg13 and together trafficked to the omegasomes/autophagosomes for autophagy initiation.

Despite an effect on autophagy initiation, autophagic flux and substrate degradation were impaired in the absence of glial dAux. Recently, two independent studies reported the involvement of GAK in autophagy and lysosomal homeostasis. Munson et. al. reported that GAK inhibition alters lysosomal morphology in U2OS cells (50). After treatment of GAK kinase inhibitor, increased lysosome numbers and large autolysosome structures were observed, possibly mediated by the increased TFEB nuclear localization (50). Miyazaki et. al. found that *GAK* KO or GAK kinase inhibitor treatment could induce autophagosome accumulation and enlarged autolysosomes in A549 cells, which were caused by impaired autophagosome-lysosome fusion and autophagic lysosome reformation (ALR) (51). Given that we demonstrated that the fusion can still occur, it is possible that lysosome function is affected so that flux and substrate degradation are impaired upon glial dAux depletion. Consistent to this notion, the number and size of glial Lamp1-positive lysosomes were increased in adult fly brains expressing the *daux*-RNAi (Figures S6D and S6E). Ultrastructural TEM analysis also showed an increase in the number and size of lysosome, which exhibits smaller and electron dense structures with single-membrane, upon *daux*-RNAi expression in glia (yellow and red arrows, Figures 1K and 1M). Thus, it is feasible that GAK/dAux regulates autophagy at multiple steps via different mechanisms.

In the present study, we identify a new autophagy factor in glia and elucidate its regulatory mechanism on autophagy initiation. Our findings provide new insights on glial autophagy and advance the understanding of PD pathology from a glial perspective. Future work will be required to investigate how GAK/dAux plays different roles in glia and neurons, and how it contributes to neurodegeneration in a tissue-specific manner.

## Materials and Methods

All animal experiments were conducted according to proper regulation and guideline under ShanghaiTech. Details of fly and mouse genotypes are included in SI Appendix and Table S1. Methods including cell lines and transfections, molecular cloning, immunohistochemistry, *in-vivo* time lapse analysis, TEM, phosphoproteomic analysis, biochemistry, confocal microscopy, and statistical analysis are described in the SI Appendix.

## Supporting information

Vedio1

Vedio2

## Acknowledgments

We thank Bloomington *Drosophila* Stock Center, Vienna *Drosophila* RNAi Center, Tsinghua Fly Center, the Core Facility of *Drosophila* Resource and Technology, Shanghai Institute of Biochemistry and Cell Biology, Chinese Academy of Sciences, Developmental Studies Hybridoma Bank, Chao Tong, Henry C. Chang, Lei Xue, Henry Sun, Tor Erik Rusten, Gabor Juhasz, Patrik Verstreken, Aike Guo, Yijun Liu, and Gerald M Rubin for fly stocks and antibodies; Chih-Hao Lee for IMG cell line; Yanfeng Liu for HeLa and HEK293FT cell lines; Kangmin He for HeLa *GAK* KO cell line; Jiawei Zhou for *CX3CR1-Cre* mice. We also thank the Molecular Imaging Core Facility (MICF), the Molecular and Cell Biology Core Facility (MCBCF), and the Multi-Omics Core Facility (MOCF) at the School of Life Science and Technology, ShanghaiTech University for providing technical support; Yu Kong and Lijun Pan in Electron Microscopy Facilities of Center for Excellence in Brain Science and Technology, Chinese Academy of Science for assistance with TEM sample preparation; Wei-Chung Lee for help on TEM; Ho lab members for discussion and comments. This work was supported by grants from ShanghaiTech and National Natural Science Foundation of China (31871039 and 32170962).

## Author contributions

S.Z, S.Y, L.W, and M.S.H conceived and designed the study. S.Z, S.Y, L.W, S.L, H.W, L.S, J.O, M.Z, R.W, M.W, Y.Z, K.Y, T.L, and M.S.H performed the experiments. S.Z, S.Y, L.W, S.L, and M.S.H analyzed the data. M.S.H wrote the paper with the input of S.Z, S.Y, and L.W. All authors read and approved the manuscript.

