## Supplementary Materials for "Cyclin-G-associated kinase GAK/dAux regulates autophagy initiation via ULK1/Atg1 in glia"

**This PDF file includes:**

Materials and Methods
Supplementary Figures S1-S6
Supplementary Video Legends
Key resources table
Table S1
Table S2

#### Materials and Methods

##### *Drosophila* genetics

Flies were raised under standard conditions at 25℃ with 70% humidity. All fly crosses were carried out at 25°C with standard laboratory conditions unless noted otherwise. All strains were obtained from Bloomington *Drosophila* Stock Center (BDSC), the Vienna *Drosophila* RNAi Center (VDRC), Tsinghua Fly Center, or as gifts from colleagues. MARCM clonal analyses were carried out using flies with the following genotypes: *yw hs-flp/repo-GAL4; UAS-mCherry.Atg8a/+; FRT*$^{82B}$*tub-GAL80*$^{ts}$*/FRT*$^{82B}$*daux*. Flies were heat-shocked at 37℃ for 30 minutes at 4-5 days after cross, then recovered at 25℃. Brains of adult flies at 3- or 10-day-old were dissected. Detailed fly genotypes for all experiments are listed in Table S1.

##### Mouse genetics

The B6;129S6-*Gak*$^{tm2Legr}$/Mmjax mice (Cat. #36793-JAX, the Jackson laboratory, Bar Harbor, ME, USA) possess loxP sites flanking exon 1 of *cyclin G associated kinase* (*GAK*) gene. *CX3CR1-Cre* transgenic mice were kindly provided by Jiawei Zhou (Institute of Neuroscience, Chinese Academy of Sciences). *CX3CR1-Cre*$^{+/-}$ littermates were used as controls. Mice were housed in the animal facility with maximum of five mice per cage under a 12 hours light/dark cycle at 22 ± 2°C. Food and water were available *ad libitum*. The use and care of animals was approved and directed by the Animal Care and Use Committee of National Center for Protein Science Shanghai. Detailed genotypes for all experiments are listed in Table S1.

##### Cell lines and transfections

IMG cell line was a gift from Chih-Hao Lee (1). Cells were cultured in Dulbecco's modified Eagle medium (Cat. #11965092, Gibco, New York, NY, USA) with high glucose (4.5

g/L), 10% fetal bovine serum (Cat. #10099141, Gibco) and 100 units/mL penicillin-streptomycin (Cat. #15140-122, Gibco) at 37°C with 5% CO2 humidified atmosphere. IMG cells were transfected using Lipo 2000 (Cat. #11668019, Life Technologies, Carlsbad, CA, USA) according to the manufacturer's protocol. In brief, cells were seeded on a 12-well plate overnight. 40 pmol siRNA (GenePharma) and/or 2 µl Lipo 2000 was diluted with 100 µl Opti-MEM (Cat. #31985070, Gibco), mixed, and incubated at room temperature for 20 minutes, then added dropwise to cells cultured at 37°C with 5% CO2. After 6 hours, cells were changed into growth medium and harvested after 72 hours.

HeLa (CRM-CCL-2) and HEK293FT (PTA-5007) cell lines were gifts from Yanfeng Liu. Cells were cultured similarly as IMG cells, and transfected using Lipo Plus (Cat. #Q03003, Sagecreation) according to the manufacturer's protocol. Cells were seed on a 6- or 12-well plate overnight and 20 µmol siRNA (GenePharma) were used at a concentration of 1:400 with 1µg or more high purity plasmid. After 24 hours, cells were changed into growth medium and harvested after more than 24 hours.

*Drosophila* S2 cells were cultured in Schneider's *Drosophila* medium (Cat. #21720024, Gibco) at 28°C and transfected with using the Effectene Transfection Reagent (Cat. #301425, Qiagen) according to the manufacturer's protocol. Cells were transfected 6 hours after seeding and $CuSO_4$ was add at a final concentration of 1 mM 24 hours after transfection, and cells were harvested at another 24 hours.

**Molecular biology**

Plasmid cloning: Plasmids were constructed by polymerase chain reaction (PCR) using DNA templates from *Drosophila* or HeLa cell. These plasmids express dAux; dAux truncated variants including $dAux^{\Delta Kinase}$ (124-948 bp), $dAux^{\Delta PTEN}$ (1717-2121 bp), $dAux^{\Delta CBD}$ (2542-3174 bp), $dAux^{\Delta J}$ (3217-3441 bp), and $dAux^{DNAJC6m}$ (3355 bp, Arginine 927 to Glycine); Atg1;

mCherry.Atg13; EGFP.Atg9; human EGFP.DFCP1; human GAK; human ULK1; ULK1 truncated variants including ULK1$^{\Delta K}$ (46-843 bp), ULK1$^{\Delta PS}$ (835-2496 bp) and ULK1$^{\Delta CTD}$ (2497-3150 bp); human DFCP1; and human LC3. The corresponding DNA sequences were subcloned into *pUAST-attB* vector containing a 3xFlag or a 6xMyc epitope tag, or mammalian cell line vector *pEGFP-C1* (Clontech, Cat. #6084-1), *pERFP-LC3* (modulate form *pEGFP-C1),* and *pRK5-Flag* by Genscript (Nanjing, China). Fly microinjection was conducted by the *Drosophila* Core Facility, Institute of Biochemistry and Cell Biology, Chinese Academy of Sciences.

Primers used for amplification:

*daux*-F: CGCGGAATTCGGCGAGTTCTTTAAGTCGCTCAACCTCAAC

*daux*-R: CGCGCTCGAGTTACGCATTAAACATATTTTGCTGCGTGGC

*GAK*-F: CGGCCGCGAATTCATCGATAGATCTGATATCAACAAGTTTG

*GAK*-R: GAAAGCTGGGTCGAATTCGCCCTTTCAGAAGAGGGGCCGGGAGCCCTG

*EGFP-DFCP1*-F: GGACTCAGATCTCGAGGGAGTGCCCAGACTTCCC

*EGFP-DFCP1*-R: GTCGACTGCAGAATTCTTAAAGGTCACCGGGCTTTTTATTGC

*RFP-LC3*-F: CTCAAGCTTCGAATTCCCCGTCGGAGAAGACCTTCAAG

*RFP-LC3*-R: GATCCCGGGCCCGCGGTACCTTACACTGACAATTTCATCCCGAACGT

*Flag-GAK*-F: GGGTCGACCGCGGCCGCTTTATCGCTGCTGCAGTCGGCGC

*Flag-GAK*-R: AATAGGGCCCTCTAGATCAGAAGAGGGGCCGGGA

Anti-dAux antibodies: anti-dAux antibodies were generated by ABclonal Biotechnology co., Ltd (Harrogate, UK). *daux* cDNA fragment encoding 694-1157 a.a. was subcloned into the pET-28a-SUMO vector, and purified proteins were injected into rats using standard protocols to generate the antibodies.

GAK KO cell line: *GAK* expression in HeLa cells was silenced using the CRISPR-Cas9 approach. The sgRNA sequence targeting the *GAK* gene (5'-GTCGGCGCTCGACTTCTTGG-3') was cloned into the vector pSpCas9(BB)-2A-GFP (PX458) (Cat. #48138, Addgene). HeLa cells were then transfected with the constructed plasmid using Lipofectamine™ 3000 Transfection Reagent (Cat. # L3000015, Invitrogen) according to the manufacturer's instructions. 48 hours after transfection, the cells were subjected to single cell sorting (FACSAria II, BD Biosciences) to plate GFP-positive single cells into 96-well plates. The monoclonal cell populations were screened for loss of GAK protein expression by western blot. Multiple clones were selected and the clone A1 was used as the *GAK* KO and the clone D2 was used as a wild type control throughout the study.

qRT-PCR: Total RNAs were extracted from adult fly heads or IMG cells using TransZol Up (Cat. #ET111-01, TransGen, Beijing, China). cDNAs were reversely transcribed using HiScript III RT SuperMix (Cat. #R323-01, Vazyme, Nanjing, China). qRT-PCR reactions were performed using ChamQ Universal SYBR qPCR Master Mix (Cat. #Q711-02, Vazyme) and ABI 7500 RT-PCR system. The mRNA levels were normalized to *rp49* (flies) or *Actin* (IMG cells), respectively. The expression levels were analyzed by ΔΔCT method. Results are representative of at least three biological replicates.

Primers used are listed below:

Flies:

*rp49*-F: CCACCAGTCGGATCGATATGC

*rp49*-R: CTCTTGAGAACGCAGGCGACC

*daux*-F: CAGCGGCTACCACCAATCCTTC

*daux*-R: TGCGTCGGCGTCCTGATAGA

IMG cells:

*Actin*-F: CTAAGGCCAACCGTGAAAAG

*Actin*-R: ACCAGAGGCATACAGGGACA

*Gak*-F: GGTCATCCAGTCTGTGGCTAAC

*Gak*-R: TTGATTGCAGACTCCACACC

**Immunohistochemistry**

Adult fly brains: Brains were dissected and fixed in 4% formaldehyde for 40 minutes, then washed with PBT (PBS + 0.1% TX-100) for 3 times and dissected further to remove additional debris in PBS solution. Clean and fixed brains were blocked in PBT solution with 5% Normal Donkey Serum (NDS) and subsequently stained with primary antibodies at 4°C overnight and then secondary antibodies at room temperature for 2 hours. Primary antibodies used: mouse anti-Atg8a (1:1000, gift from Chao Tong), rabbit anti-Ref(2)P (1:500, Cat. #ab178440, Abcam, Cambridge, UK), mouse anti-mono- and polyubiquitinylated conjugates antibody (FK2, 1:500, Cat. #BML-PW8810, Enzo Life Sciences, Farmingdale, NY, USA), rabbit anti-ULK1 (1:100, Cat. #8045S, Cell Signaling Technology, CST, Danvers, MA, USA), mouse anti-Repo (1:100, Cat. #8D12, Developmental Studies Hybridoma Bank, DSHB, Iowa City, IA, USA), and rat anti-Elav (1:500, Cat. #7E8A10, DSHB).

IMG cells: Cells were seeded on glass coverslips in 12-well plates the day before fixation with cold methanol (Cat. #100141190, Sinopharm, Beijing, China) for 15 minutes, followed by washing with PBT (PBS + 0.1% TX-100) for 15 minutes, then blocked in PBS with 5% NDS for 1 hour. Cells were stained with primary antibodies at 4°C overnight and then secondary antibodies at room temperature for 2 hours. Primary antibodies used: rabbit anti-Iba1 (1:500, Cat. #019-19741, Wako), rabbit anti-TMEM119 (1:500, Cat. #ab185333, Abcam), rabbit anti-LC3B (1:200, Cat. #2775, Cell Signaling Technology, CST, Danvers, MA, USA), and rabbit anti-GAK (1:50, Cat. #12147-1-AP, Proteintech, Chicago, IL, USA). Nuclei were labeled by DAPI (Cat. #C1005, Beyotime, Shanghai, China) at room temperature for 5-10 minutes.

HeLa cells: After 48 hours transfection, cells were cultured on coverslips, and washed with PBS three times (3 minutes for each time) before fixation with 4% formaldehyde for 15 minutes, then washed with PBS three times, permeabilized with saponin (Cat. #P0095, Beyotime, Shanghai, China), blocked in PBS with 1% BSA for 1 hour, and washed with PBT (PBS + 0.1% TX-100) for three times. Cells were stained with primary antibodies at 4°C overnight and then secondary antibodies at room temperature for 2 hours. Primary antibodies used: rabbit anti-ULK1 (1:100, Cat. #8045S, CST), rat-anti TGN38 (1:100, Cat. # 610898, BD Transduction Laboratories) and rabbit anti-GAK (1:50, Cat. #12147-1-AP, Proteintech). Nuclei were labeled by DAPI (Cat. #C1005, Beyotime) at room temperature for 5-10 minutes.

Mouse brain slices: To prepare brain slices, mice were anesthetized with isoflurane and perfused with 0.9% saline, followed by 4% paraformaldehyde in PBS. The brains were removed and post-fixed overnight in 4% paraformaldehyde in PBS at 4°C. After dehydration in 20% sucrose, a series of coronal sections (10 μm thick) across the Substantia Nigra were cut by a Thermo CryoStar NX50 cryostat (Thermofisher Scientific, USA). The sections were permeabilized with 0.3% Triton X-100 in PBS containing 5% normal goat serum for 1 h at 37°C. After blocking with 5% normal goat serum in PBS, the samples were incubated with primary antibodies at 4°C overnight. After three washes with PBS, samples were incubated for 1 hour with secondary antibodies conjugated with Alexa-fluorescein. Primary antibodies used: rabbit anti-Iba1 (1:500, Cat. #019-19741, Wako), rabbit anti-GAK (1:50, Cat. # 12147-1-AP, Proteintech), and mouse anti-LC3 (1:200, Cat. ##83506, CST).

For all samples, secondary antibodies used were from Jackson ImmunoResearch (West Grove, PA, USA): donkey anti-mouse Cy3 (1:1000, Cat. #715-166-151), donkey anti-rabbit Cy3 (1:1000, Cat. #711-166-152), donkey anti-rat Cy3 (1:1000, Cat. #712-165-153), donkey anti-mouse Cy5 (1:500, Cat. #715-175-150), donkey anti-rabbit Cy5 (1:500, Cat. #711-175-152), and donkey anti-rat Cy5 (1:500, Cat. #712-175-150).

***In-vivo* time lapse analysis**

Live adult fly brains were dissected and kept in saline solution for tracing the puncta in a single focal plane using Nikon C2 microscope with a 40X water objective. The trafficking percentage and speed was analyzed by Imaris software (Bitplane, Zurich, Switzerland) using Autoregressive Motion mode. The contacting duration were calculated manually.

**Transmission electron microscopy**

Adult fly brains were fixed, embedded, stained, and dehydrated according to previous protocols (2). Ultrathin sections (70 nm) of each brain were cut with a Diatomediamond knife on a Leica EM UC7 ultra-microtome (Leica Microsystems, Germany) and collected on copper grids. Images were acquired with Talos L120C transmission electron microscope (Thermo Fisher Scientific, Waltham, MA, UAS) operating at an acceleration voltage of 120 kV.

**Phosphoproteomic analysis**

Samples for phosphoproteomic analysis were prepared and analyzed as described previously (PTM Bio, Hangzhou, China) (3). 300 fly adult brains per sample were sonicated three times on ice using a high intensity ultrasonic processor (Scientz) in lysis buffer (8M urea, 1% Protease Inhibitor Cocktail). For digestion, the protein solution was reduced with 5 mM dithiothreitol for 30 minutes at 56°C and alkylated with 11 mM iodoacetamide for 15 mins at room temperature in darkness. The protein sample was then diluted by adding 100 mM TEAB (tetraethylammonium bromide, TEAB) to urea concentration less than 2 M. Finally, trypsin was added at 1:50 trypsin-to-protein mass ratio for the first digestion overnight and 1:100 for a second 4-hour digestion. After trypsin digestion, peptide was desalted by Strata X C18 SPE column (Phenomenex) and vacuum-dried. Peptide was reconstituted in 0.5 M TEAB and processed according to the manufacturer's protocol for TMT kit/iTRAQ kit. Briefly, one unit of TMT/iTRAQ reagent were thawed and reconstituted in acetonitrile. The peptide mixtures were then incubated for 2 hours at room temperature and pooled, desalted and dried by vacuum centrifugation. The tryptic peptides were dissolved in 0.1% formic acid (solvent A), directly loaded onto a home-made reversed-phase analytical column (15 cm length, 75 μm i.d.). The gradient was comprised of an increase from 6% to 23% solvent B (0.1% formic acid in 98% acetonitrile) over 26 minutes, 23% to 35% in 8 minutes and climbing to 80% in 3 minutes then holding at 80% for the last 3 minutes, all at a constant flow rate of 400 nL/min on an EASY-nLC 1000 UPLC system. The peptides were subjected to NSI source followed by tandem mass spectrometry (MS/MS) in Q ExactiveTM Plus (Thermo) coupled online to the UPLC. The electrospray voltage applied was 2.0 kV. The m/z scan range was 350 to 1800 for full scan, and intact peptides were detected in the Orbitrap at a resolution of 70,000. Peptides were then selected for MS/MS using NCE setting as 28 and the fragments were detected in the Orbitrap at a resolution of 17,500. A data-dependent protocol that alternates between one MS scan followed by 20 MS/MS scans with 15.0s dynamic exclusion was conducted. Automatic gain control (AGC) was set at 5E4. Fixed first mass was set as 100 m/z.

**Biochemistry**

Western blot analysis: Fly adult heads were homogenized using a motorized pestle (Cat. #116005500, MP Biomedicals, Irvine, CA, USA) in lysis buffer (0.4% NP-40, 0.2 mM EDTA, 150 mM NaCl, 20% glycerol, 100 mM Tris-HCl pH7.5, 2% Tween 20, 0.5 mM phosphodiesterase inhibitors, and 1 mM PMSF). For Atg8a detection, fly heads were boiled with 1XSDS-Dye in 100°C for 10 minutes. Proteins were separated on SDS-PAGE gels and transferred to PVDF membranes (Cat. #IPFL00010, Millipore, Billerica, MA, USA). The membranes were blocked by 5% fat-free milk diluted in PBS with 0.1% Tween-20 for 40 minutes. Samples were incubated with the primary antibodies at 4°C overnight and HRP-conjugated secondary antibodies at room temperature for 2 hours. Primary antibodies used: rat anti-dAux (1:1000, this study), rabbit anti-GABARAP+GABARAPL1+GABARAPL2 antibody (1:1000, Cat. #ab109364, Abcam), rabbit anti-Ref(2)P (1:1000, Cat. #ab178440, Abcam), mouse anti-Mono- and polyubiquitinylated conjugates antibody (FK2, 1:500, Cat. #BML-PW8810, Enzo Life Sciences), mouse anti-α-Tubulin (1:5000, Cat. #T9026, Sigma), rabbit anti-Myc (1:2000, Cat. #0912-2, Hua An Biotechnology), and mouse anti-Flag (1:1000, Cat. #F3165, Sigma, St. Louis, MO, USA). Bands were visualized by Clarity Western ECL Substrate (Cat. #WBKLS0500, Millipore).

For IMG cells, cells were treated and lysed in 2XSDS loading buffer at 95℃ for 10 minutes. Primary antibodies used: rabbit anti-GAK (1:50, Cat. #12147-1-AP, Proteintech), mouse anti-P62 (1:1000, Cat. #ab56416, Abcam), and mouse anti-Ubiquitin (1:1000, Cat. #3936S, CST).

For HeLa cells, cells were lysed in lysis buffer (0.4% NP-40, 0.2mM EDTA, 150mM NaCl, 20% glycerol, 100mM Tris-HCl pH7.5, 2% Tween 20, 0.5mM phosphodiesterase inhibitors, and 1mM PMSF) in 4℃ for 30 minutes. The cell lysates were clarified by centrifugation at 13,000 rpm, at 4°C for 10 minutes, then boiled in 100℃ for 5 minutes, samples were collected and stored in -80°C. Primary antibodies used: rabbit-anti-ULK1 (1:1000, Cat. #8045S, CST), rabbit-anti-p-ULK1 S757 (1:1000, Cat. #14202T, CST), rabbit-anti-ATG13 (1:1000, Cat. #13273S, CST), rabbit-anti-BECLIN1 (1:1000, Cat. #3495T, CST), rabbit-anti-ATG14 (1:1000, Cat. #96752S, CST), rabbit-anti-LC3 (1:1000, Cat. #2775, CST), mouse-anti-P62 (1:1000, Cat. #ab56416, Abcam), mouse-anti-Ubi (1:1000, Cat. #3936S, CST), rabbit-Myc (1:2000, Cat. #0912-2, Hua An Biotechnology), and mouse-anti-Flag (1:1000, Cat. #F3165, Sigma, St. Louis, MO, USA).

For all examples, secondary antibodies used are from Jackson ImmunoResearch: goat anti-mouse-HRP (1:5000, Cat. #115-035-003), goat anti-rabbit-HRP (1:5000, Cat. #111-036-003) and goat anti-rat-HRP (1:5000, Cat. #112-035-003). Results are representative of at least three biological replicates.

Co-immunoprecipitation: HEK293FT or S2 cells were transfected and lysed in lysis buffer (0.4% NP-40, 0.2 mM EDTA, 150 mM NaCl, 20% glycerol, 100 mM Tris-HCl pH7.5, 2% Tween 20, 0.5 mM phosphodiesterase inhibitors, and 1 mM PMSF) in 4 ℃ for 30 minutes. The cell lysates were clarified by centrifugation at 13,000 rpm, at 4 °C for 10 minutes, the soluble supernatant was incubated with prewashed anti-FLAG(R) M2 beads (Cat. #A2220; Sigma) at 4°C overnight. After pull-downs, gels were extensively washed in lysis buffer for 25 minutes three times and then subjected to Western blot analysis. The following antibodies were used for blotting: rabbit-Myc (1:2000, Cat. #0912-2, Hua An Biotechnology), mouse-anti-Flag (1:1000, Cat. #F3165, Sigma) and rabbit-anti-GFP (1:2000, SB-AB0047, ShareBio).

BafilomycinA1 Treatment: BafilomycinA1 (BafA1, Cat. #BML-CM110-0100, ENZO) was diluted in DMSO (Cat. #D8371-50, solarbio) in a concentration of 1 mM, then added to 0.5 ml of liquid yeast for a final concentration of 4 μM. Yeast containing BafA1 was added to the top of the fly food in standard vials kept overnight to allow the evaporation of residual DMSO. Flies of different genotypes (25 females and 25 males) were then put in each vial and kept at 25°C for 5 days. After treatment, flies were collected for western blot analysis. HeLa cells were treated with 100 nM BafA1 for 3 hours after transfection.

**Confocal microscopy and statistical analysis**

Images of adult fly brains, mouse brain slices, IMG cells and HeLa cells were acquired by scanning a serial Z-stack of sections, each of 0.4 μm thickness, using Nikon A1, C2, or Spinning Disk confocal microscope (Tokyo, Japan) with the 40X or 60X oil objective. Super-resolution images were taken by STED SP8 (Zeiss, Germany) for a single layer section on the desired regions using 100X oil objective. The whole fly brain was positioned so that they can be scanned anteriorly to posteriorly (top to bottom). Representative single layer images at the similar plane are shown, except mouse brain slice images shown as maximal projection. For number and size analysis, original and unmodified images were imported into ImageJ (National Institutes of Health), and the intensity threshold for the relevant channel was set so that maximum number of dots were selected without miscounting two adjacent dots into one. For colocalization measurements, primary images were imported and analyzed automatically using the colocalization plugin with the intensity correlation tool, showing as Pearson's Correlation (R value, no threshold) or Manders' Correlation (M1 or M2). All data were imported into GraphPad Prism 8 and shown in a scatter plot with bar form. When calculating the statistical significance, Shapiro-Wilk normality test was used to check normal distribution, for normally distributed datasets, two-tailed t-test or ordinary one-way ANOVA followed by Tukey's multiple comparisons test was used, for datasets that did not have a normal distribution, Mann-Whitney test or Kruskal-Wallis tests followed by Dunn's multiple comparisons test were used. P value less than 0.05 is considered significant. ns: no significance, $p \geq 0.05$; *: $p<0.05$; **: $p<0.01$; ***: $p<0.001$; ****: $p<0.0001$.

### Figure S1

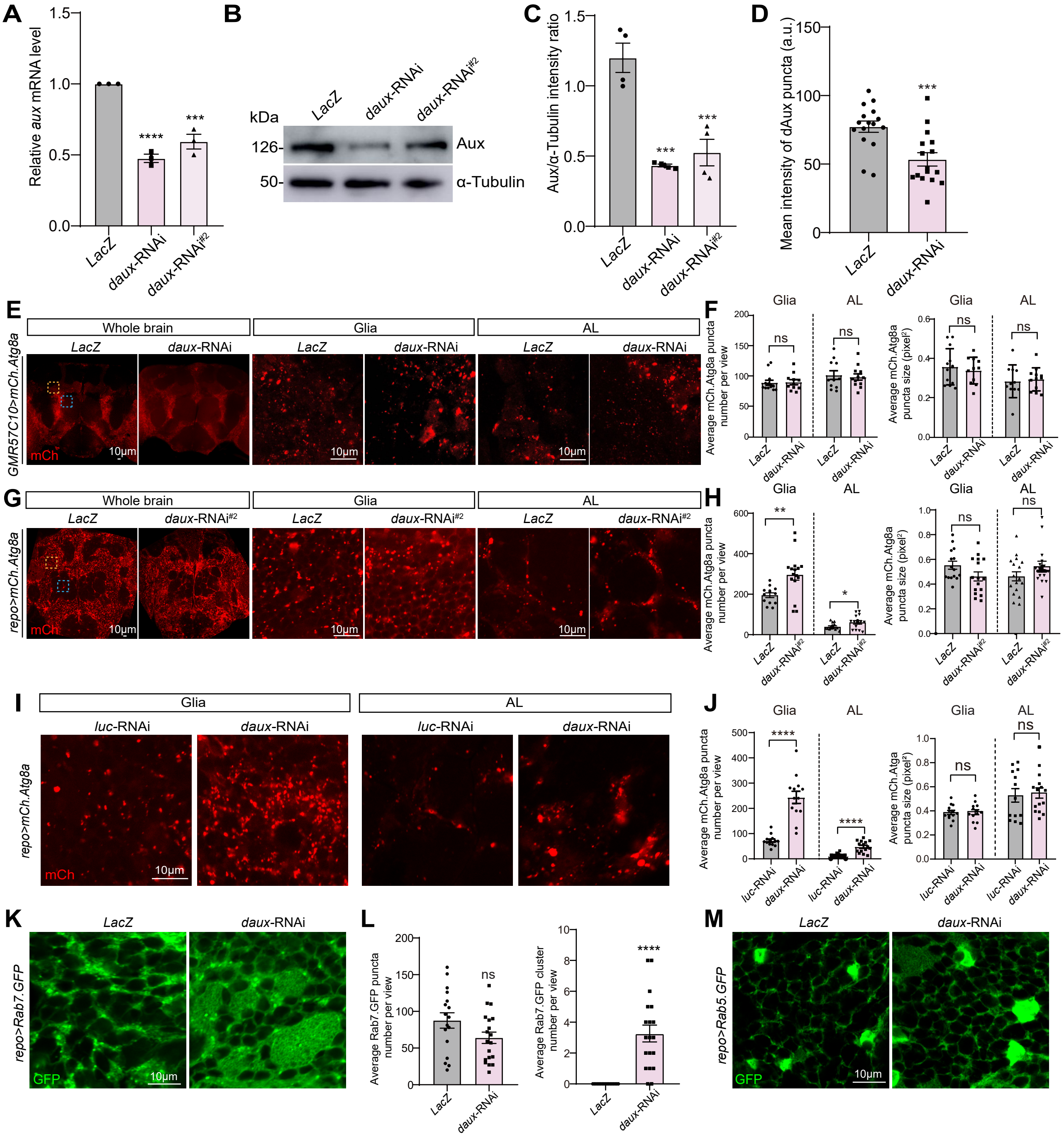
A
Relative aux mRNA level
LacZ
daux-RNAi
daux-RNAi#2
B
kDa
126-
50-
Aux
α-Tubulin
C
Aux/α-Tubulin intensity ratio
D
Mean intensity of dAux puncta (a.u.)
E
Whole brain
Glia
AL
GMR57C10>mCh.Atg8a
mCh
10μm
F
Average mCh.Atg8a puncta number per view
Average mCh.Atg8a puncta size (pixel²)
ns
G
repo>mCh.Atg8a
H
I
luc-RNAi
J
Average mCh.Atga puncta size (pixel²)
K
repo>Rab7.GFP
GFP
L
Average Rab7.GFP puncta number per view
Average Rab7.GFP cluster number per view
M
repo>Rab5.GFP

**Figure S1 (related to Figure 1). Lack of dAux selectively increases the number of autophagosomes in adult fly glia**

(**A**-**D**) *daux*-RNAi efficiency was analyzed by qRT-PCR (A), WB (B and C), and immunostaining with the anti-dAux antibodies (D). RNA and protein samples were collected from the control and *repo>daux*-RNAi adult fly heads. Two independent RNAi lines: V16182 (*daux*-RNAi) and BL39017 (*daux*-RNAi$^{\#2}$), were used. Note that both *daux* mRNA and protein levels decreased when expressing either *daux*-RNAi in glia. *rp49* and α-Tubulin served as controls for qRT-PCR and WB, respectively. (**E** and **F**) Representative images (E) and quantifications (F) of neuronal autophagic structures in Whole brain, "Glia" and "AL" region when *daux*-RNAi was expressed under the control of a pan-neuronal driver *GMR57C10-Gal4*. Note that autophagic structures labeled by *mCh.Atg8a* (red) did not exhibit significant alternation in the overall morphology and distribution upon neuronal dAux depletion. **(G** and **H)** Representative images (G) and quantifications (H) of glial autophagic structures in the Whole brain, "Glia", and "AL" region upon expressing *daux*-RNAi$^{\#2}$ (BL39017) in glia. Note that similar to the other RNAi, expressing *daux*-RNAi$^{\#2}$ in glia increased the number of autophagic structures in both "Glia" and "AL" region. **(I** and **J)** A different control *UAS-luciferase*-RNAi (*luc*-RNAi) was used for comparison with *daux*-RNAi. The number and size of autophagic structure in "Glia" and "AL" region were assessed in flies expressing the control *luc*-RNAi or *daux*-RNAi. Note that results are consistent to the main Figures using *UAS-LacZ* as a control throughout the study. (**K**-**M**) Representative images (K and M) and quantifications (L) of Rab7.GFP and Rab5.GFP in "Glia" region of fly brains expressing *daux*-RNAi in glia. Glial late endosomes and autophagosomes labeled by Rab7.GFP, early endosomes labeled by Rab5.GFP remained largely unaffected. Scale bars and the sample number n are indicated in the Figures. For each experiment, more than three biologically independent replicates were done. Whereas results were consistent, representative results from one experiment was shown.

A serial confocal Z-stack sections were taken with 0.4 μm each, and representative single layer images acquired at similar focal planes are shown. All quantification figures from the same images but with different parameters quantified are listed under the same label unless noted otherwise. Data are shown as mean ± SEM. P-values of significance (indicated with asterisks, ns no significance, * $p<0.05$, ** $p<0.01$, and *** $p<0.001$) are calculated by two-tailed unpaired t-test, Mann-Whitney test, or ordinary one-way ANOVA followed by Tukey's multiple comparisons test. All genotypes analyzed are listed in Table S1.

Figure S2

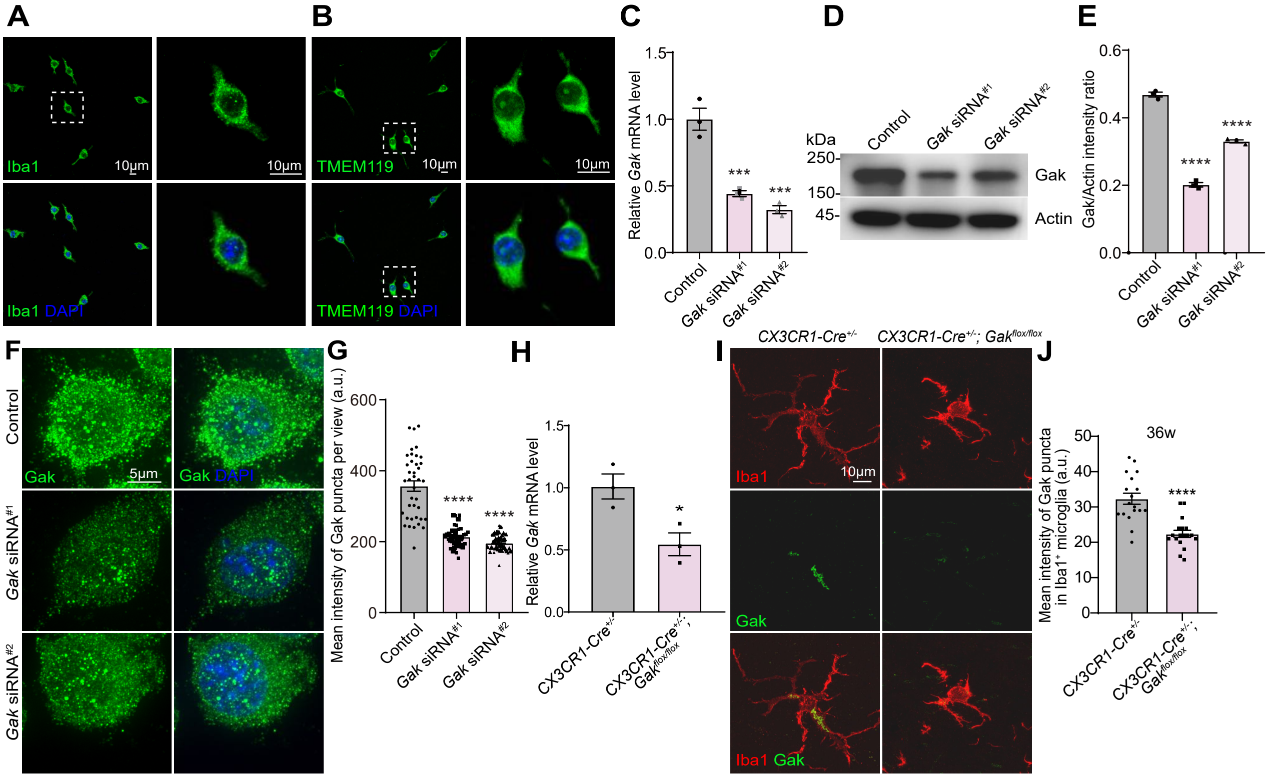
A
Iba1
10μm
10μm
Iba1 DAPI
B
TMEM119
10μm
10μm
TMEM119 DAPI
C
Relative Gak mRNA level
1.5
1.0
0.5
0.0
***
***
Control
Gak siRNA#1
Gak siRNA#2
D
kDa
250-
150-
45-
Control
Gak siRNA#1
Gak siRNA#2
Gak
Actin
E
Gak/Actin intensity ratio
0.6
0.4
0.2
0.0
****
****
Control
Gak siRNA#1
Gak siRNA#2
F
Control
Gak siRNA#1
Gak siRNA#2
Gak
5μm
Gak DAPI
G
Mean intensity of Gak puncta per view (a.u.)
600
400
200
0
****
****
Control
Gak siRNA#1
Gak siRNA#2
H
Relative Gak mRNA level
1.5
1.0
0.5
0.0
*
CX3CR1-Cre+/-
CX3CR1-Cre+/-; Gakflox/flox
I
CX3CR1-Cre+/-
CX3CR1-Cre+/-; Gakflox/flox
Iba1
10μm
Gak
Iba1 Gak
J
36w
Mean intensity of Gak puncta in Iba1+ microglia (a.u.)
50
40
30
20
10
0
****
CX3CR1-Cre+/-
CX3CR1-Cre+/-; Gakflox/flox

**Figure S2 (related to Figure 1). Lack of Gak increases the number and size of autophagosomes in mouse microglia**

(**A** and **B**) Immortalized microglial (IMG) cells were verified as stained positive for microglial specific markers Iba1 or TMEM119 (green). DAPI is a nucleus marker (blue). (**C**-**G**) *Gak* expression was efficiently silenced when transfecting either *Gak* siRNA to IMG cells. Note a reduction in *Gak* mRNA and protein levels as detected by qRT-PCR (C), western blot (WB, D and E), and immunostaining with the anti-Gak antibodies (F and G) upon treatment with the *Gak* siRNA. (**H**) *Gak* mRNA expression level was reduced in the primary microglia of *Gak* cKO mice ($CX3CR1\text{-}Cre^{+/-}$; $GAK^{flox/flox}$) as revealed by qRT-PCR analysis. (**I** and **J**) Gak protein expression was reduced in *Gak* cKO microglia ($CX3CR1\text{-}Cre^{+/-}$; $GAK^{flox/flox}$) as revealed by anti-Gak immunostaining on mouse substantia nigra (SN) microglia. Scale bars and the sample number n are indicated in the Figures. For each experiment, more than three biologically independent replicates were done. Whereas results were consistent, representative results from one experiment was shown. A serial confocal Z-stack sections were taken with 0.4 μm each, and representative single layer images acquired at the similar plane are shown, except mouse brain slice images (I) shown as maximal projection. Data are shown as mean ± SEM. P-values of significance (indicated with asterisks, ns no significance, * $p<0.05$, ** $p<0.01$, and *** $p<0.001$) are calculated by two-tailed unpaired t-test or ordinary one-way ANOVA followed by Tukey's multiple comparisons.

Figure S3

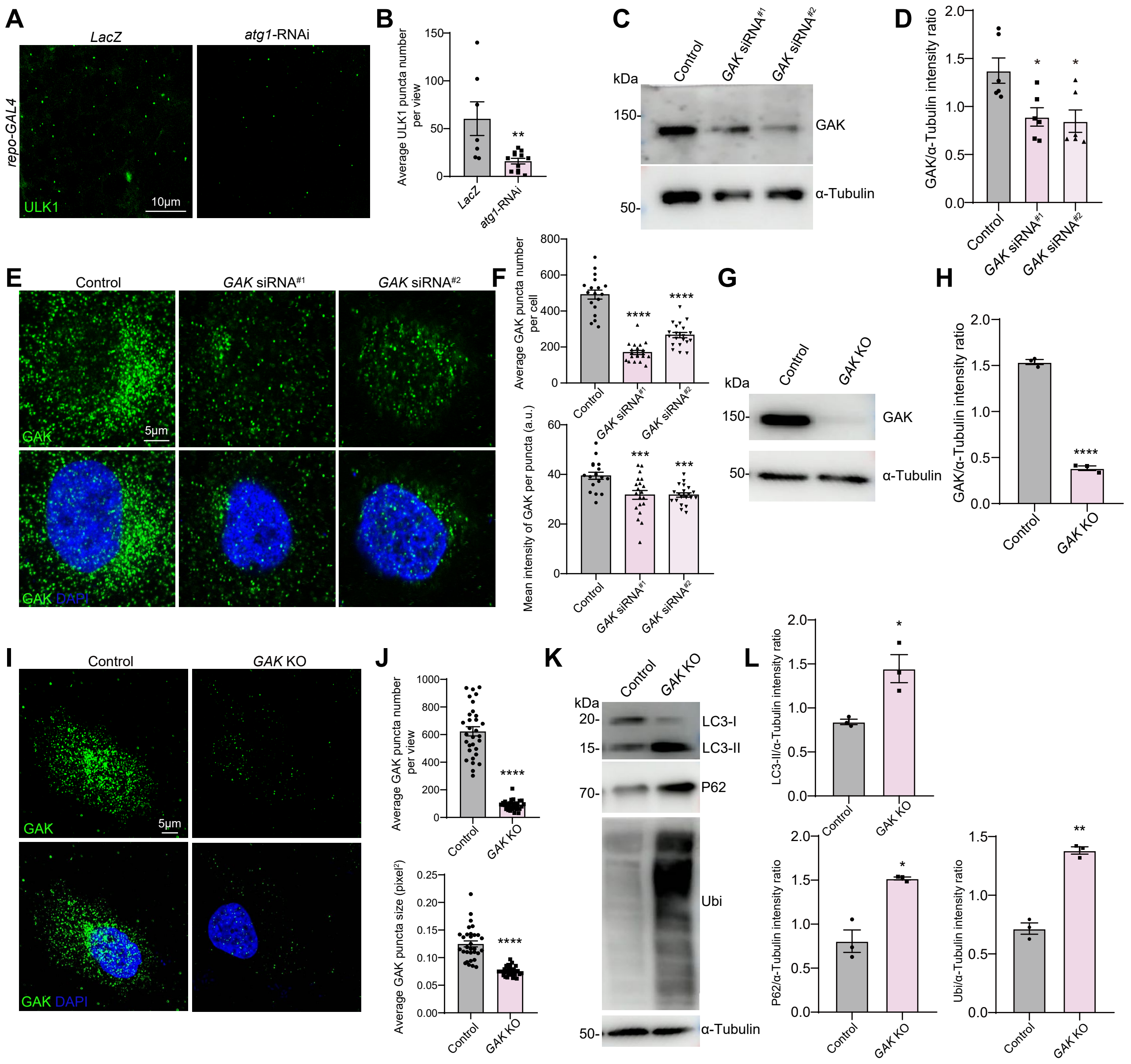

A
LacZ
atg1-RNAi
repo-GAL4
ULK1
10μm
B
Average ULK1 puncta number per view
LacZ
atg1-RNAi
C
Control
GAK siRNA#1
GAK siRNA#2
kDa
150-
50-
GAK
α-Tubulin
D
GAK/α-Tubulin intensity ratio
E
Control
GAK siRNA#1
GAK siRNA#2
GAK
5μm
GAK DAPI
F
Average GAK puncta number per cell
Mean intensity of GAK per puncta (a.u.)
G
Control
GAK KO
H
GAK/α-Tubulin intensity ratio
I
Control
GAK KO
GAK
5μm
GAK DAPI
J
Average GAK puncta number per view
Average GAK puncta size (pixel²)
K
kDa
20-
15-
70-
50-
LC3-I
LC3-II
P62
Ubi
α-Tubulin
L
LC3-II/α-Tubulin intensity ratio
P62/α-Tubulin intensity ratio
Ubi/α-Tubulin intensity ratio

**Figure S3 (related to Figures 2). *GAK* expression is efficiently silenced in *GAK* siRNA-treated or KO cells**

(**A** and **B**) Representative images (A) and quantification (B) of the ULK1-positive puncta number in the “Glia” region of adult fly brains. Note that the ULK1 puncta number decreased upon expressing *atg1*-RNAi in glia, validating the *atg1*-RNAi efficiency as well as fly Atg1 detection by the anti-ULK1 antibodies. (**C**-**F**) *GAK* siRNA efficiency was analyzed by WB (C and D) and immunostaining with the anti-GAK antibodies (E and F). (**G**-**J**) GAK protein levels were detect by WB (G and H) and immunostaining (I and J) in HeLa cells silencing *GAK* expression with the CRISPR-Cas9 (*GAK* KO). The clone A1 is used for the *GAK* KO cells throughout the study. Control is a wild-type clone selected after CRISPR-Cas9 (D2). (**K** and **L**) Representative WB images (K) and quantifications (L) of LC3-II, P62, and Ubi proteins level in *GAK* KO cells. Scale bars and the sample number n are indicated in the Figures. For each experiment, more than three biologically independent replicates were done. Whereas results were consistent, representative results from one experiment was shown. A serial confocal Z-stack sections were taken with 0.4 μm each, and representative single layer images acquired at similar focal planes are shown. Data are shown as mean ± SEM. P-values of significance (indicated with asterisks, ns no significance, * $p<0.05$, ** $p<0.01$, and *** $p<0.001$) are calculated by two-tailed unpaired t-test, Mann-Whitney test, ordinary one-way ANOVA followed by Tukey’s multiple comparisons test, or Kruskal-Wallis tests followed by Dunn’s multiple comparisons test.

Figure S4

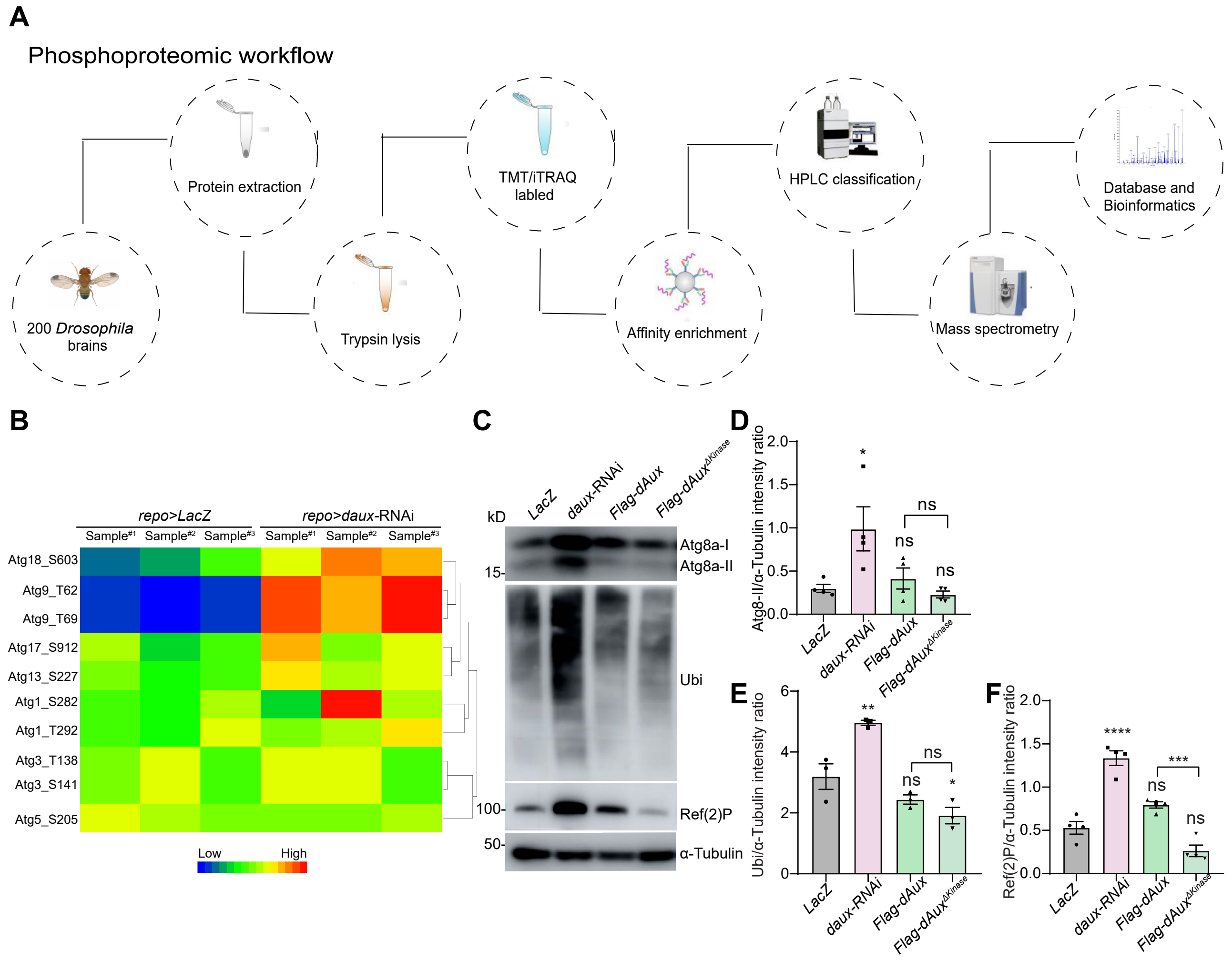
A
Phosphoproteomic workflow
200 Drosophila brains
Protein extraction
Trypsin lysis
TMT/iTRAQ labled
Affinity enrichment
HPLC classification
Mass spectrometry
Database and Bioinformatics
B
repo>LacZ
repo>daux-RNAi
Atg18_S603
Atg9_T62
Atg9_T69
Atg17_S912
Atg13_S227
Atg1_S282
Atg1_T292
Atg3_T138
Atg3_S141
Atg5_S205
Low
High
C
LacZ
daux-RNAi
Flag-dAux
Flag-dAuxΔKinase
kD
Atg8a-I
Atg8a-II
Ubi
Ref(2)P
α-Tubulin
D
Atg8-II/α-Tubulin intensity ratio
E
Ubi/α-Tubulin intensity ratio
F
Ref(2)P/α-Tubulin intensity ratio
ns

**Figure S4 (related to Figure 3). Phosphoproteomic analysis on glial dAux substrates**

(**A**) Workflow chart of phosphoproteomic analysis. (**B**) Phosphoproteomic analysis of the control and *repo>daux-RNAi* adult fly brains. Note that phosphorylations of different Atg proteins were altered upon glial dAux depletion. (**C**-**F**) dAux lacking the kinase activity impacted minimally on autophagy. Note that expression of Atg8a-II protein level remained similar and not significantly different from the LacZ control upon glial dAux or dAux$^{\Delta Kinase}$ expression (C and D). Ubi and P62 protein levels were slightly affected by glial dAux$^{\Delta Kinase}$ (E and F). Sample number n are indicated in the Figures. For each experiment, more than three biologically independent replicates were done. Whereas results were consistent, representative results from one experiment was shown. Data are shown as mean ± SEM. P-values of significance (indicated with asterisks, ns no significance, * $p<0.05$, ** $p<0.01$, and *** $p<0.001$) are calculated by ordinary one-way ANOVA followed by Tukey's multiple comparisons test or Kruskal-Wallis tests followed by Dunn's multiple comparisons test.

### Figure S5

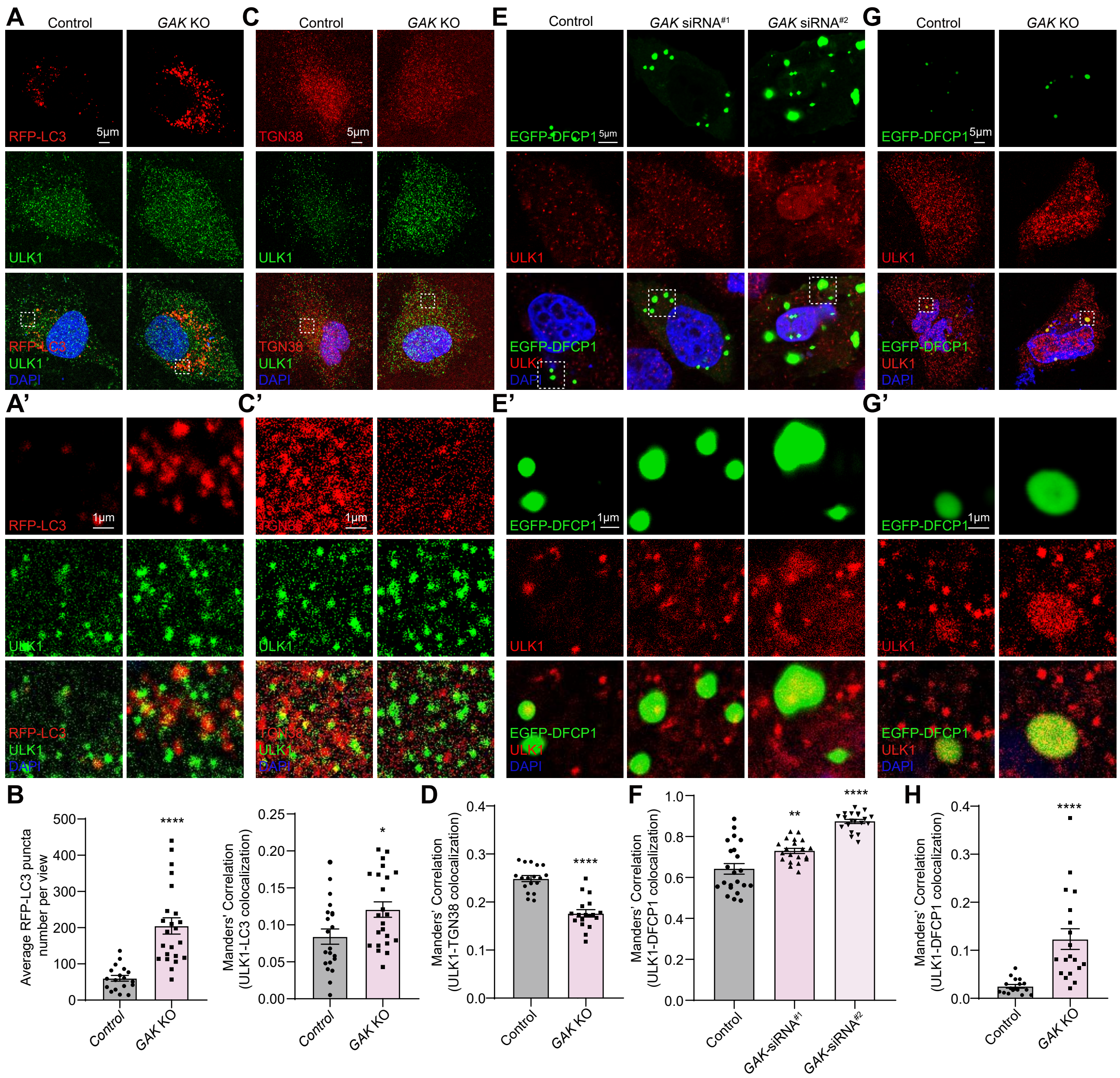

A
Control
GAK KO
C
E
GAK siRNA#1
GAK siRNA#2
G
RFP-LC3
ULK1
DAPI
TGN38
EGFP-DFCP1
5μm
1μm
A'
C'
E'
G'
B
Average RFP-LC3 puncta number per view
Manders' Correlation (ULK1-LC3 colocalization)
D
Manders' Correlation (ULK1-TGN38 colocalization)
F
Manders' Correlation (ULK1-DFCP1 colocalization)
GAK-siRNA#1
GAK-siRNA#2
H
****
**
*

**Figure S5 (related to Figure 4). GAK regulates ULK1 trafficking to omegasomes and autophagosomes**

(**A**-**H**) *GAK* siRNA-treated or KO cells were transfected with RFP-LC3 or EGFP-DFCP1 for subsequent immunostaining analysis with the anti-ULK1 or anti-TGN38 antibodies. Areas enclosed by the white dashed squares in A, C, E, and G are enlarged in A’, C’, E’, and G’. Note that the LC3 puncta number (A and B) and the colocalization of ULK1 (green)-LC3 (red) (A and B) or ULK1 (red)-DFCP1 (green) (E-H) were increased in the absence of *GAK*. ULK1-DFCP1 colocalization were analyzed in both *GAK* siRNA-treated (E and F) or KO (G and H) cells. The ULK1 (green)-TGN (red) colocalization was decreased (C and D). Scale bars and the sample number n are indicated in the Figures. For each experiment, more than three biologically independent replicates were done. Whereas results were consistent, representative results from one experiment was shown. A serial confocal Z-stack sections were taken with 0.4 μm each, and representative single layer images acquired at similar focal planes are shown. Colocalization is analyzed using the Manders’ Correlation, taking into account the change in the protein level. Data are shown as mean ± SEM. P-values of significance (indicated with asterisks, ns no significance, * $p<0.05$, ** $p<0.01$, and *** $p<0.001$) are calculated by two-tailed unpaired t-test, Mann-Whitney test, or ordinary one-way ANOVA followed by Tukey’s multiple comparisons test.

### Figure S6

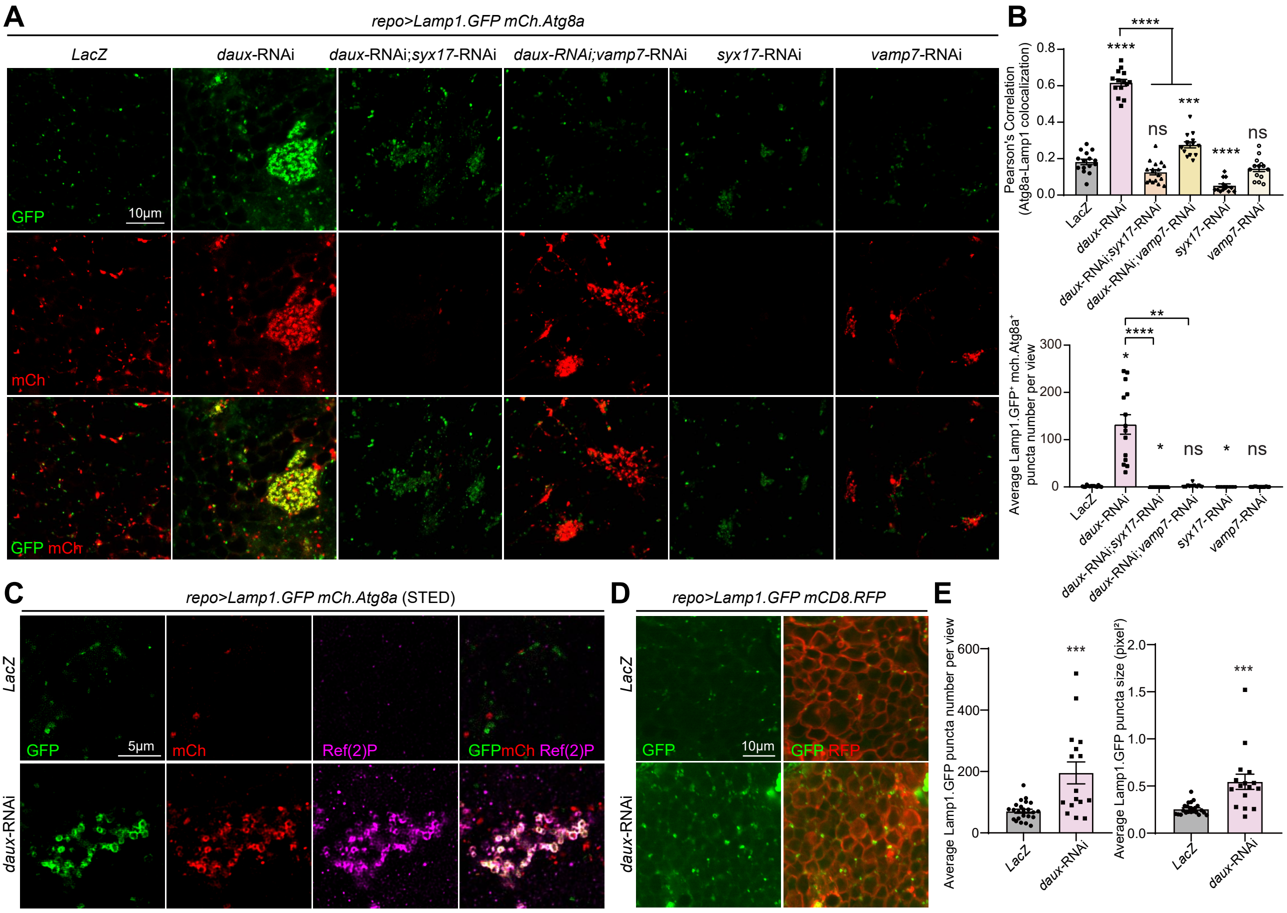

A
repo>Lamp1.GFP mCh.Atg8a
LacZ
daux-RNAi
daux-RNAi;syx17-RNAi
daux-RNAi;vamp7-RNAi
syx17-RNAi
vamp7-RNAi
GFP
10μm
mCh
GFP mCh
B
Pearson's Correlation (Atg8a-Lamp1 colocalization)
****
****
ns
***
****
ns
Average Lamp1.GFP+ mch.Atg8a+ puncta number per view
**
****
*
*
ns
*
ns
C
repo>Lamp1.GFP mCh.Atg8a (STED)
LacZ
daux-RNAi
GFP
5μm
mCh
Ref(2)P
GFPmCh Ref(2)P
D
repo>Lamp1.GFP mCD8.RFP
GFP
10μm
GFP RFP
E
Average Lamp1.GFP puncta number per view
***
Average Lamp1.GFP puncta size (pixel²)
***

**Figure S6 (related to Figure 6). Lack of GAK/dAux impairs substrate degradation**

(**A** and **B**) Representative images (A) and quantifications (B) of Lamp1.GFP- and mCh.Atg8a-positive puncta in the "Glia" region of adult fly brain with the indicated genotypes. Note that dAux depletion in glia increased the number of autolyososmes as revealed by the increase in colocalization and number of dual positive puncta. Co-expression of *syx17*-RNAi or *vamp7*-RNAi in the presence of *daux*-RNAi blocked fusion, hence the colocalization of Atg8a and Lamp1. (**C**) Representative STED images of Lamp1.GFP-, mCh.Atg8a-, and anti-Ref(2)P-positive puncta in the "Glia" region of adult fly brains. Note that substantial co-localization of Lamp1.GFP-, mCh.Atg8a-, and anti-Ref(2)P were observed, indicating that theses fused autolysosomes were able to recognize and encircle protein substrates. (**D** and **E**) Representative images (D) and quantifications (E) of lysosomes in the "Glia" region of adult fly brain. Glial lysosomes (green) are labeled with *repo-GAL4* driven *UAS-Lamp1.GFP* (*UAS-Lamp1.GFP; repo-GAL4*, *UAS-mCD8.RFP*). Note that downregulating glial dAux induces an increase in the lysosome number and size. Scale bars and the sample number n are indicated in the Figures. For each experiment, more than three biologically independent replicates were done. Whereas results were consistent, representative results from one experiment was shown. A serial confocal Z-stack sections were taken with 0.4 µm each, and representative single layer images acquired at similar focal planes are shown. Colocalization is analyzed using the Pearson's Correlation and present as the Pearson's R value. Data are shown as mean ± SEM. P-values of significance (indicated with asterisks, ns no significance, * $p<0.05$, ** $p<0.01$, and *** $p<0.001$) are calculated by two-tailed unpaired t-test, Mann-Whitney test, ordinary one-way ANOVA followed by Tukey's multiple comparisons test, or Kruskal-Wallis tests followed by Dunn's multiple comparisons test.

#### Supplementary Video Legend

**Videos. Live-cell imaging of Atg9 trafficking to Atg8a**

Live-cell imaging of Atg9 trafficking to Atg8a in the “Glia” region of control (Video 1) and *aux*-RNAi (Video 2) adult fly brains. Note the percentage of Atg9-positic puncta being trafficked to the vicinity of Atg8a-positive puncta was increased, the duration of the Atg9-Atg8a contacting was longer, and the speed of both Atg8a- and Atg9-positive puncta were decreased upon glial dAux depletion. Adult fly brains were dissected and kept in saline solution, and a single focal plane were recorded. More than three biologically independent replicates were done. Whereas results were consistent, representative results from one experiment was shown.

**Key Resources Table**

| REAGENT or RESOURCE | SOURCE | IDENTIFIER |
|---|---|---|
| Antibodies | | |
| mouse anti-Atg8a | Chao Tong | Liu, W. *et al. Autophagy* (2018) |
| rabbit anti-Ref(2)P | Abcam | Cat. #ab178440 |
| mouse anti-Mono- and polyubiquitinylated conjugates | Enzo Life Sciences | Cat. #BML-PW8810 |
| rat monoclonal anti-Elav | DSHB | Cat. #7E8A10 |
| mouse monoclonal anti-Repo | DSHB | Cat. #8D12 |
| rat anti-Aux | This study | ABclonal |
| rabbit anti-Iba1 | Wako | Cat. #019-19741 |
| rabbit anti-TMEM119 | Abcam | Cat. #ab185333 |
| rabbit anti-LC3B | Cell Signaling Technology | Cat. #2775 |
| mouse anti-LC3 | Cell Signaling Technology | Cat. #83506 |
| rabbit anti-GAK | Proteintech | Cat. #12147-1-AP |
| rabbit anti-GABARAP +GABARAPL1+GABARAPL2 | Abcam | Cat. #ab109364 |
| mouse anti-α-Tubulin | Sigma | Cat. #T9026 |
| rabbit anti-Myc | Hua An Biotechnology | Cat. #0912-2 |
| mouse anti-Flag | Sigma | Cat. #F3165 |
| mouse anti-P62 | Abcam | Cat. #ab56416 |
| mouse anti-Ubiquitin | Cell Signaling Technology | Cat. #3936S |
| rabbit anti-ULK1 | Cell Signaling Technology | Cat. #8045S |
| rabbit anti-p-ULK1(Ser757) | Cell Signaling Technology | Cat. #14202T |

| | | |
|---|---|---|
| rabbit anti-ATG13 | Cell Signaling Technology | Cat. #13273S |
| rabbit anti-ATG14 | Cell Signaling Technology | Cat. #96752S |
| rabbit anti-BECLIN1 | Cell Signaling Technology | Cat. #3495T |
| Rat-anti-TGN38 | BD Transduction Laboratories | Cat. # 610898 |
| goat anti-mouse-HRP | Jackson ImmunoResearch | Cat. #115-035-003 |
| goat anti-rabbit-HRP | Jackson ImmunoResearch | Cat. #111-036-003 |
| goat anti-rat-HRP | Jackson ImmunoResearch | Cat. #112-035-003 |
| donkey anti-mouse Cy3 | Jackson ImmunoResearch | Cat. #715-166-151 |
| donkey anti-rabbit Cy3 | Jackson ImmunoResearch | Cat. #711-166-152 |
| donkey anti-rat Cy3 | Jackson ImmunoResearch | Cat. #712-165-153 |
| donkey anti-mouse Cy5 | Jackson ImmunoResearch | Cat. #715-175-150 |
| donkey anti-rabbit Cy5 | Jackson ImmunoResearch | Cat. #711-175-152 |
| donkey anti-rat Cy5 | Jackson ImmunoResearch | Cat. #712-175-150 |
| Chemicals, peptides, and recombinant proteins | | |
| penicillin-streptomycin | Gibco | Cat. #15140-122 |
| Bafilomycin A1 | ENZO | Cat. #BML-CM110-0100 |
| papain | Roche | Cat. #5401119001 |
| liberase TM | Roche | Cat. #5401119001 |
| Experimental models: Cell lines | | |
| Immortalized microglial cell line (IMG) | Chih-Hao Lee | McCarthy, R. C. *et al. J Neuroinflammation* (2016) |
| HeLa (CRM-CCL-2) cell line | Yanfeng Liu | |
| HEK293FT (PTA-5007) cell line | Yanfeng Liu | |

| Experimental models: Organisms/strains | | |
|---|---|---|
| mice B6;129S6-*Gak*$^{tm2Legr}$/Mmjax | The Jackson laboratory | Cat. #36793-JAX |
| *w*$^{1118}$ | Bloomington Stock Center | RRID:BDSC_5905 |
| *UAS-LacZ* | Bloomington Stock Center | RRID:BDSC_1777 |
| *UAS-luciferase-RNAi* | Bloomington Stock Center | RRID:BDSC_31603 |
| *UAS-daux*-RNAi$^{\#2}$ | Bloomington Stock Center | RRID:BDSC_39017 |
| *UAS-atg1*-RNAi | Bloomington Stock Center | RRID:BDSC_80434 |
| *UAS-syx17*-RNAi | Bloomington Stock Center | RRID:BDSC_ 25896 |
| *UAS-vamp7*-RNAi | Bloomington Stock Center | RRID:BDSC_43543 |
| *UAS-Rab7.GFP* | Bloomington Stock Center | RRID:BDSC_42705 |
| *UAS-Rab5.GFP* | Bloomington Stock Center | RRID:BDSC_43336 |
| *UAS-mCherry.Atg8a* | Bloomington Stock Center | RRID:BDSC_37750 |
| *UAS-HRP* | Aike Guo | RRID:BDSC_8763 |
| *UAS-daux*-RNAi | Vienna Drosophila Resource Center | VDRC16182 |
| *UAS-atg1*-RNAi | Tsinghua Fly Center | THU2357 |
| *UAS-Lamp1.GFP* | Helmut Krämer | Pulipparacharuvil, S. *et al.* *J Cell Sci* (2005) |
| *daux*$^{L78H}$ | Henry C. Chang | Zhou, X. *et al.* *Development* (2011). |
| *daux*$^{I670K}$ | Henry C. Chang | Zhou, X. *et al.* *Development* (2011). |
| *daux*$^{G257E}$ | Henry C. Chang | Zhou, X. *et al.* *Development* (2011). |
| *pmCherry-Atg8* | Tor Erik Rusten | Katheder, N. S. *et al.* *Nature* (2017) |

| | | |
|---|---|---|
| *UASp-GFP-mCherry-Atg8a* | Henry Sun | Lee, Y. M. & Sun, Y. H. *et al. PLoS genetics* (2015) |
| *UAS-GFP.Atg6* | Chao Tong | Liu, W. *et al. Autophagy* (2018) |
| *UAS-GFP.Atg14* | Chao Tong | Liu, W. *et al. Autophagy* (2018) |
| *UAS-GFP.ZFYVE1* | Chao Tong | Liu, W. *et al. Autophagy* (2018) |
| *10xUAS-IVS-Syn21-GFP-p10* | Gerald M Rubin | Pfeiffer, B. D. *et al. Proc Natl Acad Sci U S A* (2012) |
| Oligonucleotides | | |
| *daux*-F: CGCGGAATTCGGCGAGTTCTTTAAGTCGCTCAACCTCAAC | This study | N/A |
| *daux-R: CGCGCTCGAGTTACGCATTAAACATATTTTGCTGCGTGGC* | This study | N/A |
| *GAK*-F: CGGCCGCGAATTCATCGATAGATCTGATATCAACAAGTTTG | This study | N/A |
| *GAK*-R: GAAAGCTGGGTCGAATTCGCCCTTTCAGAAGAGGGGCCGGGAGCCCTG | This study | N/A |
| *EGFP-DFCP1-F: GGACTCAGATCTCGAGGGAGTGCCCAGACTTCCC* | This study | N/A |
| *EGFP-DFCP1-R: GTCGACTGCAGAATTCTTAAAGGTCACCGGGCTTTTTATTGC* | This study | N/A |
| *RFP-LC3-F: CTCAAGCTTCGAATTCCCCGTCGGAGAAGACCTTCAAG* | This study | N/A |
| *RFP-LC3-R: GATCCCGGGCCCGCGGTACCTTACACTGACAATTTCATCCCGAACGT* | This study | N/A |

| | | |
|---|---|---|
| *Flag-GAK-F: GGGTCGACCGCGGCCGCTTTATC GCTGCTGCAGTCGGCGC* | This study | N/A |
| *Flag-GAK-R: AATAGGGCCCTCTAGATCAGAAGA GGGGCCGGGA* | This study | N/A |
| qRT-PCR: *rp49*-F: CCACCAGTCGGATCGATATGC | This study | N/A |
| qRT-PCR: *rp49*-R: CTCTTGAGAACGCAGGCGACC | This study | N/A |
| qRT-PCR: *daux*-F: CAGCGGCTACCACCAATCCTTC | This study | N/A |
| qRT-PCR: *daux*-R: TGCGTCGGCGTCCTGATAGA | This study | N/A |
| qRT-PCR: *Actin*-F: CTAAGGCCAACCGTGAAAAG | This study | N/A |
| qRT-PCR: *Actin*-R: ACCAGAGGCATACAGGGACA | This study | N/A |
| qRT-PCR: *Gak*-F: GGTCATCCAGTCTGTGGCTAAC | This study | N/A |
| qRT-PCR: *Gak*-R: TTGATTGCAGACTCCACACC | This study | N/A |
| SiRNA sequence | | |
| IMG *Gak* siRNA[#1]: CCUGGAUGCUUGUGAUAUUTT (5’-3’) | This study | N/A |
| IMG *Gak* siRNA[#2]: GGACCAAACAGCAAGACUUTT (5’-3’) | This study | N/A |
| HeLa *GAK* siRNA[#1]: CACCAGAAAUCAUAGACUUTT (5’-3’) | This study | N/A |
| HeLa *GAK* siRNA[#2]: CCCGAACAUUGUCCAGUUUTT (5’-3’) | This study | N/A |
| HeLa *GAK* KO sgRNA: GTCGGCGCTCGACTTCTTGG (5’-3’) | | |
| Software and algorithms | | |
| Fiji | http://fiji.sc | RRID:SCR_002285 |
| Imaris | https://imaris.oxinst.cn | RRID:SCR_007370 |

| | | |
|---|---|---|
| NIS-Elements Basic Research | Nikon | RRID:SCR_002776 |
| GraphPad Prism8 | GraphPad Prism | RRID:SCR_002798 |
| Others | | |
| DMSO | solarbio | Cat. #D8371-50 |
| Schneider’s medium | Invitrogen | Cat. #21720024 |
| DAPI | Beyotime | Cat. #C1005 |
| DMEM | Gibco | Cat. #A1896701 |
| Dulbecco’s modified Eagle medium | Gibco | Cat. #11965092 |
| Fetal bovine serum | Gibco | Cat. #10099141 |
| Lipo 2000 | Life Technologies | Cat. #11668019 |
| Opti-MEM | Gibco | Cat. #31985070 |
| PVDF membranes | PVDF membranes | Cat. #IPFL00010 |
| ECL Substrate | Millipore | Cat. #WBKLS0500 |

**Table S1**

Detailed fly or mouse genotypes in each experiment categorized by Figures.

| Figure | Genotype |
|---|---|
| Fig. 1 | |
| (B) | repo-GAL4, 10xUAS-IVS-Syn21-GFP-p10/+ |
| (C and D) | UAS-mCherry.Atg8a/+; repo-GAL4/UAS-LacZ |
| | UAS-mCherry.Atg8a/UAS-daux-RNAi; repo-GAL4/+ |
| (E and F) | pmCherry.Atg8a/+; repo-GAL4/UAS-LacZ |
| | pmCherry.Atg8a/UAS-daux-RNAi; repo-GAL4/+ |
| (G and H) | repo-GAL4/UAS-LacZ |
| | UAS-daux-RNAi/+; repo-GAL4/+ |
| (I and J) | yw hs-flp/repo-GAL4; UAS-mCherry.Atg8a/+; $FRT^{82B}$ tub-$Gal80^{ts}$/ $FRT^{82B}$ $Sb^{1}$ |
| | yw hs-flp/repo-GAL4; UAS-mCherry.Atg8a/+; $FRT^{82B}$ tub-$Gal80^{ts}$/ $FRT^{82B}$ $daux^{L78H}$ |
| | yw hs-flp/repo-GAL4; UAS-mCherry.Atg8a/+; $FRT^{82B}$ tub-$Gal80^{ts}$/ $FRT^{82B}$ $daux^{G257E}$ |
| | yw hs-flp/repo-GAL4; UAS-mCherry.Atg8a/+; $FRT^{82B}$ tub-$Gal80^{ts}$/ $FRT^{82B}$ $daux^{I670K}$ |
| (K-N) | UAS-HRP/+; repo-GAL4/UAS-LacZ |
| | UAS-HRP/UAS-daux-RNAi; repo-GAL4/+ |
| (O and P) | UAS-mCherry.Atg8a/UAS-daux-RNAi; repo-GAL4/UAS-LacZ |
| | UAS-mCherry.Atg8a/UAS-daux-RNAi; repo-GAL4/UAS-3xFlag-dAux |
| | UAS-mCherry.Atg8a/UAS-daux-RNAi; repo-GAL4/UAS-3xFlag-hGAK |
| (S and T) | CX3CR1-$Cre^{+/-}$ |
| | CX3CR1-$Cre^{+/-}$; $Gak^{flox/flox}$ |
| Fig. 2 | |
| (A and B) | repo-GAL4/UAS-LacZ |
| | UAS-daux-RNAi/+; repo-GAL4/+ |
| (C and D) | repo-GAL4/UAS-LacZ |
| | UAS-daux-RNAi/+; repo-GAL4/+ |
| | UAS-mCherry.Atg13/+; repo-GAL4/UAS-LacZ |
| | UAS-mCherry.Atg13/UAS-daux-RNAi; repo-GAL4/+ |
| (E and F) | UAS-GFP.Atg6/+; repo-GAL4, UAS-mCD8.RFP/UAS-LacZ |
| | UAS-GFP.Atg6/UAS-daux-RNAi; repo-GAL4, UAS-mCD8.RFP/+ |
| | UAS-GFP.Atg14/+; repo-GAL4, UAS-mCD8.RFP/UAS-LacZ |
| | UAS-GFP.Atg14/UAS-daux-RNAi; repo-GAL4, UAS-mCD8.RFP/+ |
| (G and H) | UAS-GFP.ZFYVE1/+; repo-GAL4, UAS-mCD8.RFP/UAS-LacZ |
| | UAS-GFP.ZFYVE1/UAS-daux-RNAi; repo-GAL4, UAS-mCD8.RFP/+ |
| | UAS-EGFP.DFCP1/+; repo-GAL4/UAS-LacZ |
| | UAS-EGFP.DFCP1/UAS-daux-RNAi; repo-GAL4/+ |

| | |
|---|---|
| Fig. 3 | |
| (C) | *repo-GAL4/UAS-LacZ* |
| | *repo-GAL4/UAS-3xFlag-dAux* |
| | *repo-GAL4/UAS-3xFlag-dAux*$^{\Delta Kinase}$ |
| | *repo-GAL4/UAS-3xFlag-dAux*$^{\Delta PTEN}$ |
| | *repo-GAL4/UAS-3xFlag-dAux*$^{\Delta CBD}$ |
| | *repo-GAL4/UAS-3xFlag-dAux*$^{\Delta J}$ |
| | *repo-GAL4/UAS-3xFlag-hGAK* |
| | *repo-GAL4/UAS-3xFlag-dAux*$^{\Delta DNAJC6m}$ |
| (E and F) | *repo-GAL4/UAS-LacZ* |
| | *repo-GAL4/UAS-3xFlag-dAux* |
| | *repo-GAL4/UAS-3xFlag-dAux*$^{\Delta Kinase}$ |
| | *repo-GAL4/UAS-3xFlag-dAux*$^{\Delta PTEN}$ |
| | *repo-GAL4/UAS-3xFlag-dAux*$^{\Delta CBD}$ |
| | *repo-GAL4/UAS-3xFlag-dAux*$^{\Delta J}$ |
| | *repo-GAL4/UAS-3xFlag-dAux*$^{\Delta DNAJC6m}$ |
| (M and O) | *UAS-GFP.ZFYVE1/+; repo-GAL4, UAS-mCD8.RFP/UAS-LacZ* |
| | *UAS-GFP.ZFYVE1/+; repo-GAL4, UAS-mCD8.RFP/UAS-atg1*-RNAi |
| | *UAS-GFP.ZFYVE1/UAS-daux*-RNAi*; repo-GAL4, UAS-mCD8.RFP/ UAS-LacZ* |
| | *UAS-GFP.ZFYVE1/UAS-daux*-RNAi*; repo-GAL4, UAS-mCD8.RFP/ UAS-atg1*-RNAi |
| (N and P) | *UAS-mCherry.Atg8a/+; repo-GAL4/UAS-LacZ* |
| | *UAS-mCherry.Atg8a/+; repo-GAL4/UAS-atg1*-RNAi |
| | *UAS-mCherry.Atg8a/UAS-daux*-RNAi*; repo-GAL4/UAS-LacZ* |
| | *UAS-mCherry.Atg8a/UAS-daux*-RNAi*; repo-GAL4/UAS-atg1*-RNAi |
| Fig. 4 | |
| (C and D) | *UAS-mCherry.Atg8a/+; repo-GAL4/UAS-LacZ* |
| | *UAS-mCherry.Atg8a/UAS-daux*-RNAi*; repo-GAL4/+* |
| Fig. 5 | |
| (A and C) | *UAS-mCherry.Atg8a/+; repo-GAL4, UAS-EGFP.Atg9/UAS-LacZ* |
| | *UAS-mCherry.Atg8a/UAS-daux*-RNAi*; repo-GAL4, UAS-EGFP.Atg9/+* |
| | *UAS-mCherry.Atg8a/+; repo-GAL4, UAS-EGFP.Atg9/UAS-atg1-*RNAi |
| (B and D) | *UAS-mCherry.Atg8a/UAS-LacZ; repo-GAL4, UAS-EGFP.Atg9/UAS-LacZ* |
| | *UAS-mCherry.Atg8a/UAS-daux*-RNAi*; repo-GAL4, UAS-EGFP.Atg9/ UAS-LacZ* |
| | *UAS-mCherry.Atg8a/UAS-daux*-RNAi*; repo-GAL4, UAS-EGFP.Atg9/ UAS-atg1*-RNAi |
| (E-H) | *UAS-mCherry.Atg8a/+; repo-GAL4, UAS-EGFP.Atg9/UAS-LacZ* |

| | |
|---|---|
| | *UAS-mCherry.Atg8a/UAS-daux*-RNAi*; repo-GAL4, UAS-EGFP.Atg9/+* |
| Fig. 6 | |
| (A and B) | *UASp-GFP-mCherry-Atg8a/+; repo-GAL4/UAS-LacZ* |
| | *UASp-GFP-mCherry-Atg8a/UAS-daux*-RNAi*; repo-GAL4/+* |
| (C and D) | *UAS-Lamp1.GFP, UAS-mCherry.Atg8a/+; repo-GAL4/UAS-LacZ* |
| | *UAS-Lamp1.GFP, UAS-mCherry.Atg8a/UAS-daux*-RNAi*; repo-GAL4/+* |
| (E-H) | *repo-GAL4/UAS-LacZ* |
| | *UAS-daux*-RNAi*/+; repo-GAL4/+* |
| Fig. S1 | |
| (A-C) | *repo-GAL4/UAS-LacZ* |
| | *UAS-daux*-RNAi*/+; repo-GAL4/+* |
| | *UAS-daux*-RNAi$^{\#2}$*/+; repo-GAL4/+* |
| (D) | *repo-GAL4/UAS-LacZ* |
| | *UAS-daux*-RNAi*/+; repo-GAL4/+* |
| (E and F) | *UAS-mCherry.Atg8a/+; GMR57C10-GAL4/UAS-LacZ* |
| | *UAS-mCherry.Atg8a/UAS-daux*-RNAi*; GMR57C10-GAL4/+* |
| (G and H) | *UAS-mCherry.Atg8a/+; repo-GAL4/UAS-LacZ* |
| | *UAS-mCherry.Atg8a/UAS-daux*-RNAi$^{\#2}$*; repo-GAL4/+* |
| (I and J) | *UAS-mCherry.Atg8a/+; repo-GAL4/UAS-luc-RNAi* |
| | *UAS-mCherry.Atg8a/UAS-daux*-RNAi*; repo-GAL4/+* |
| (K and L) | *UAS-Rab7.GFP/+; repo-GAL4, UAS-mCD8.RFP/UAS-LacZ* |
| | *UAS-Rab7.GFP/UAS-daux*-RNAi*; repo-GAL4, UAS-mCD8.RFP/+* |
| (M) | *repo-GAL4, UAS-Rab5.GFP/UAS-LacZ* |
| | *UAS-daux*-RNAi*/+; repo-GAL4, UAS-Rab5.GFP/+* |
| Fig. S2 | |
| (H-J) | *CX3CR1-Cre*$^{+/-}$ |
| | *CX3CR1-Cre*$^{+/-}$*; Gak*$^{flox/flox}$ |
| Fig. S3 | |
| (A and B) | *repo-GAL4/UAS-LacZ* |
| | *repo-GAL4/UAS-atg1*-RNAi |
| Fig. S4 | |
| (A and B) | *repo-GAL4/UAS-LacZ* |
| | *UAS-daux*-RNAi*/+; repo-GAL4/+* |
| (C-F) | *repo-GAL4/UAS-LacZ* |
| | *UAS-daux*-RNAi*/+; repo-GAL4/+* |
| | *repo-GAL4/UAS-3xFlag-dAux* |
| | *repo-GAL4/UAS-3xFlag-dAux*$^{\Delta Kinase}$ |
| Fig. S6 | |
| (A and B) | *UAS-Lamp1.GFP, UAS-mCherry.Atg8a/+; repo-GAL4/UAS-LacZ* |
| | *UAS-Lamp1.GFP, UAS-mCherry.Atg8a/UAS-daux*-RNAi*; repo-GAL4/+* |

| | |
|---|---|
| | *UAS-Lamp1.GFP, UAS-mCherry.Atg8a/UAS-daux*-RNAi*; repo-GAL4/ UAS-syx17*-RNAi |
| | *UAS-Lamp1.GFP, UAS-mCherry.Atg8a/UAS-daux*-RNAi*; repo-GAL4/ UAS-vamp7*-RNAi |
| | *UAS-Lamp1.GFP, UAS-mCherry.Atg8a/+; repo-GAL4/UAS-syx17-*RNAi |
| | *UAS-Lamp1.GFP, UAS-mCherry.Atg8a/+; repo-GAL4/UAS-vamp7-*RNAi |
| (C) | *UAS-Lamp1.GFP, UAS-mCherry.Atg8a/+; repo-GAL4/UAS-LacZ* |
| | *UAS-Lamp1.GFP, UAS-mCherry.Atg8a/UAS-daux*-RNAi*; repo-GAL4/+* |
| (D and E) | *UAS-Lamp1.GFP/+; repo-GAL4, UAS-mCD8.RFP/UAS-LacZ* |
| | *UAS-Lamp1.GFP/UAS-daux*-RNAi*; repo-GAL4, UAS-mCD8.RFP/+* |

#### Table S2

Phosphoproteomic analysis table.

| Protein accession | Position | Amino acid | *daux*-RNAi /*LacZ* Ratio | *daux*-RNAi/ *LacZ* P value | Regulated Type | Protein description | Gene name |
|---|---|---|---|---|---|---|---|
| Q8SYH8 | 159 | S | 1.394 | 0.000208735 | Up | RE57644p OS=Drosophila melanogaster OX=7227 GN=CG17159 PE=2 SV=1 | CG17159 |
| Q8SYH8 | 162 | S | 1.375 | 0.000935167 | Up | RE57644p OS=Drosophila melanogaster OX=7227 GN=CG17159 PE=2 SV=1 | CG17159 |
| M9NDC4 | 136 | S | 1.327 | 0.01608733 | Up | Uncharacterized protein, isoform G OS=Drosophila melanogaster OX=7227 GN=Dmel\CG4896 PE=4 SV=1 | Dmel\CG4896 |
| A0A0B4KEU9 | 489 | S | 1.533 | 0.010906411 | Up | Missing-in-metastasis, isoform K OS=Drosophila melanogaster OX=7227 GN=mim PE=4 SV=1 | mim |
| E5DK16 | 74 | S | 2.388 | 0.003577414 | Up | Lipin isoform J OS=Drosophila melanogaster OX=7227 GN=Lpin PE=1 SV=1 | Lpin |
| Q7KMM8 | 862 | S | 1.343 | 0.019456285 | Up | BcDNA.GH03482 OS=Drosophila melanogaster OX=7227 GN=Kank PE=2 SV=1 | Kank |
| Q9VH10 | 1288 | S | 1.565 | 0.000800557 | Up | Uncharacterized protein, isoform B OS=Drosophila melanogaster OX=7227 GN=CG34108 PE=1 SV=3 | CG34108 |
| Q9VH10 | 1382 | S | 1.316 | 0.012321285 | Up | Uncharacterized protein, isoform B OS=Drosophila melanogaster OX=7227 GN=CG34108 PE=1 SV=3 | CG34108 |
| Q9VH10 | 911 | S | 1.415 | 0.001008757 | Up | Uncharacterized protein, isoform B OS=Drosophila melanogaster OX=7227 GN=CG34108 PE=1 SV=3 | CG34108 |
| Q9VH10 | 551 | S | 1.521 | 0.00297187 | Up | Uncharacterized protein, isoform B OS=Drosophila melanogaster OX=7227 GN=CG34108 PE=1 SV=3 | CG34108 |
| Q9VH10 | 2562 | S | 1.441 | 0.007780387 | Up | Uncharacterized protein, isoform B OS=Drosophila melanogaster OX=7227 GN=CG34108 PE=1 SV=3 | CG34108 |
| A0A0B4JD03 | 536 | S | 1.4 | 0.016718244 | Up | Suppressor of variegation 2-10, isoform J OS=Drosophila melanogaster OX=7227 GN=Su(var)2-10 PE=1 SV=1 | Su(var)2-10 |
| C8VV76 | 483 | S | 1.755 | 0.01158269 | Up | LP09925p OS=Drosophila melanogaster OX=7227 GN=CG18659-RA PE=2 SV=1 | CG18659-RA |
| C8VV76 | 487 | S | 1.755 | 0.01158269 | Up | LP09925p OS=Drosophila melanogaster OX=7227 GN=CG18659-RA PE=2 SV=1 | CG18659-RA |
| A0A0B4JD23 | 254 | S | 1.393 | 0.015395676 | Up | Uncharacterized protein, isoform J OS=Drosophila melanogaster OX=7227 GN=Dmel\CG2246 PE=1 SV=1 | Dmel\CG2246 |
| A0A0B4JD39 | 407 | S | 1.323 | 0.028591521 | Up | Tramtrack, isoform H OS=Drosophila melanogaster OX=7227 GN=ttk PE=4 SV=1 | ttk |
| A0A0B4JD78 | 135 | S | 1.363 | 0.000441883 | Up | Protein kinase C OS=Drosophila melanogaster OX=7227 GN=aPKC PE=1 SV=1 | aPKC |
| A0A0B4JD78 | 141 | S | 1.363 | 0.000441883 | Up | Protein kinase C OS=Drosophila melanogaster OX=7227 GN=aPKC PE=1 SV=1 | aPKC |
| Q9VF03 | 372 | S | 1.37 | 0.002610043 | Up | Brahma associated protein 155 kDa OS=Drosophila melanogaster OX=7227 GN=mor PE=1 SV=3 | mor |
| Q9VF03 | 335 | S | 1.339 | 0.032788717 | Up | Brahma associated protein 155 kDa OS=Drosophila melanogaster OX=7227 GN=mor PE=1 SV=3 | mor |
| A0A0B4KH90 | 11 | S | 3.531 | 0.000566016 | Up | Uncharacterized protein, isoform R OS=Drosophila melanogaster OX=7227 GN=CG14890 PE=4 SV=1 | CG14890 |
| A0A0B4JDC3 | 603 | S | 1.375 | 0.011415672 | Up | Uncharacterized protein, isoform B OS=Drosophila melanogaster OX=7227 GN=Dmel\CG6621 PE=4 SV=1 | Dmel\CG6621 |
| A0A0B4JDC8 | 959 | S | 1.445 | 0.000319047 | Up | Uncharacterized protein, isoform C OS=Drosophila melanogaster OX=7227 GN=CG6934 PE=4 SV=1 | CG6934 |
| E2QCZ8 | 441 | S | 1.324 | 0.005909136 | Up | Smallish, isoform F OS=Drosophila melanogaster OX=7227 GN=smash PE=1 SV=1 | smash |
| A0A0B4K623 | 520 | S | 0.509 | 0.000711556 | Down | Pugilist, isoform E OS=Drosophila melanogaster OX=7227 GN=pug PE=1 SV=1 | pug |
| A0A0B4K6Q6 | 648 | S | 1.381 | 0.013458858 | Up | Prospero, isoform H OS=Drosophila melanogaster OX=7227 GN=pros PE=4 SV=1 | pros |
| A0A0B4K6Q6 | 651 | S | 1.381 | 0.013458858 | Up | Prospero, isoform H OS=Drosophila melanogaster OX=7227 GN=pros PE=4 SV=1 | pros |
| A0A0B4K6Q6 | 654 | S | 1.381 | 0.013458858 | Up | Prospero, isoform H OS=Drosophila melanogaster OX=7227 GN=pros PE=4 SV=1 | pros |
| A0A0B4K6V2 | 18 | S | 1.672 | 0.000747983 | Up | Like-AP180, isoform G OS=Drosophila melanogaster OX=7227 GN=lap PE=1 SV=1 | lap |
| A0A0B4K6V2 | 305 | S | 2.211 | 0.003464117 | Up | Like-AP180, isoform G OS=Drosophila melanogaster OX=7227 GN=lap PE=1 SV=1 | lap |
| A0A0B4K657 | 2399 | S | 0.736 | 0.025570717 | Down | Bitesize, isoform I OS=Drosophila melanogaster OX=7227 GN=btsz PE=4 SV=1 | btsz |
| A0A0B4K657 | 793 | S | 0.765 | 0.008726264 | Down | Bitesize, isoform I OS=Drosophila melanogaster OX=7227 GN=btsz PE=4 SV=1 | btsz |
| Q59DW8 | 9 | S | 1.417 | 0.000602694 | Up | Gilgamesh, isoform E OS=Drosophila melanogaster OX=7227 GN=gish PE=1 SV=1 | gish |
| B7Z0T3 | 651 | S | 1.445 | 0.013692435 | Up | Mustard, isoform S OS=Drosophila melanogaster OX=7227 GN=mtd PE=1 SV=1 | mtd |
| B7Z0T3 | 424 | S | 1.346 | 0.007612693 | Up | Mustard, isoform S OS=Drosophila melanogaster OX=7227 GN=mtd PE=1 SV=1 | mtd |
| A0A0B4K699 | 529 | S | 2.282 | 0.004818737 | Up | Mustard, isoform Y OS=Drosophila melanogaster OX=7227 GN=mtd PE=1 SV=1 | mtd |
| B7Z0T3 | 777 | S | 2.766 | 0.001198369 | Up | Mustard, isoform S OS=Drosophila melanogaster OX=7227 GN=mtd PE=1 SV=1 | mtd |
| B7Z0T3 | 120 | S | 1.663 | 0.010677327 | Up | Mustard, isoform S OS=Drosophila melanogaster OX=7227 GN=mtd PE=1 SV=1 | mtd |
| B7Z0T3 | 122 | S | 1.994 | 0.003106987 | Up | Mustard, isoform S OS=Drosophila melanogaster OX=7227 GN=mtd PE=1 SV=1 | mtd |
| Q9VDG5 | 421 | S | 1.989 | 0.011125492 | Up | AT07459p OS=Drosophila melanogaster OX=7227 GN=Calx PE=1 SV=1 | Calx |
| A0A0B4K6D2 | 127 | S | 2.038 | 0.00010164 | Up | p23, isoform B OS=Drosophila melanogaster OX=7227 GN=p23 PE=1 SV=1 | p23 |
| Q9VC62 | 629 | S | 1.372 | 0.00054636 | Up | Extended synaptotagmin-like protein 2, isoform A OS=Drosophila melanogaster OX=7227 GN=Esyt2 PE=1 SV=1 | Esyt2 |
| Q9VC62 | 604 | S | 2.717 | 0.000958868 | Up | Extended synaptotagmin-like protein 2, isoform A OS=Drosophila melanogaster OX=7227 GN=Esyt2 PE=1 SV=1 | Esyt2 |
| A0A0B4KI37 | 1802 | S | 4.832 | 2.98969E-05 | Up | Scribble, isoform T OS=Drosophila melanogaster OX=7227 GN=scrib PE=1 SV=1 | scrib |
| A0A0B4KI37 | 1780 | S | 1.653 | 0.001462339 | Up | Scribble, isoform T OS=Drosophila melanogaster OX=7227 GN=scrib PE=1 SV=1 | scrib |
| A0A0B4KI37 | 1672 | S | 1.717 | 0.001475352 | Up | Scribble, isoform T OS=Drosophila melanogaster OX=7227 GN=scrib PE=1 SV=1 | scrib |
| A0A0B4KI37 | 1887 | S | 1.376 | 0.015884145 | Up | Scribble, isoform T OS=Drosophila melanogaster OX=7227 GN=scrib PE=1 SV=1 | scrib |
| A0A0B4K6M4 | 2370 | S | 1.754 | 0.003545052 | Up | Scribble, isoform O OS=Drosophila melanogaster OX=7227 GN=scrib PE=1 SV=1 | scrib |
| A0A0B4KEF4 | 1875 | S | 1.343 | 9.80401E-05 | Up | Down syndrome cell adhesion molecule 1, isoform CD OS=Drosophila melanogaster OX=7227 GN=Dscam1 PE=1 SV=1 | Dscam1 |
| A0A0B4K6W1 | 578 | S | 1.323 | 0.030417014 | Up | Suppressor of hairy wing, isoform C OS=Drosophila melanogaster OX=7227 GN=su(Hw) PE=4 SV=1 | su(Hw) |
| A0A0B4LJ24 | 610 | S | 1.462 | 0.006126777 | Up | Rho GTPase activating protein at 100F, isoform I OS=Drosophila melanogaster OX=7227 GN=RhoGAP100F PE=4 SV=1 | RhoGAP100F |
| A0A0B4K730 | 336 | S | 2.181 | 0.000148552 | Up | Ced-6, isoform F OS=Drosophila melanogaster OX=7227 GN=ced-6 PE=1 SV=1 | ced-6 |
| A0A0B4K730 | 357 | S | 2.606 | 0.002147572 | Up | Ced-6, isoform F OS=Drosophila melanogaster OX=7227 GN=ced-6 PE=1 SV=1 | ced-6 |

| | | | | | | | |
|---|---|---|---|---|---|---|---|
| A0A0B4LF82 | 909 | S | 1.425 | 0.02774767 | Up | Sin3A, isoform H OS=Drosophila melanogaster OX=7227 GN=Sin3A PE=1 SV=1 | Sin3A |
| A1Z9J3 | 8652 | S | 1.575 | 0.033893692 | Up | Short stop, isoform H OS=Drosophila melanogaster OX=7227 GN=shot PE=1 SV=1 | shot |
| A1Z9J3 | 5642 | S | 1.387 | 0.004212478 | Up | Short stop, isoform H OS=Drosophila melanogaster OX=7227 GN=shot PE=1 SV=1 | shot |
| E1JH65 | 4925 | S | 1.512 | 0.014517569 | Up | Short stop, isoform P OS=Drosophila melanogaster OX=7227 GN=shot PE=1 SV=1 | shot |
| A0A0B4K849 | 703 | S | 1.964 | 0.003719119 | Up | Hu li tai shao, isoform O OS=Drosophila melanogaster OX=7227 GN=hts PE=1 SV=1 | hts |
| Q7JRI6 | 137 | S | 3.179 | 0.000129326 | Up | RE18604p OS=Drosophila melanogaster OX=7227 GN=BcDNA:RE18604 PE=1 SV=1 | BcDNA:RE18604 |
| Q7JRI6 | 140 | S | 1.858 | 0.000371 | Up | RE18604p OS=Drosophila melanogaster OX=7227 GN=BcDNA:RE18604 PE=1 SV=1 | BcDNA:RE18604 |
| Q7JRI6 | 120 | S | 2.077 | 0.000249389 | Up | RE18604p OS=Drosophila melanogaster OX=7227 GN=BcDNA:RE18604 PE=1 SV=1 | BcDNA:RE18604 |
| A0A0B4K7K7 | 67 | S | 2.302 | 3.32582E-05 | Up | Wunen, isoform C OS=Drosophila melanogaster OX=7227 GN=wun PE=4 SV=1 | wun |
| A0A0B4K7K7 | 69 | S | 2.382 | 9.85516E-05 | Up | Wunen, isoform C OS=Drosophila melanogaster OX=7227 GN=wun PE=4 SV=1 | wun |
| A0A0B4K7K9 | 1173 | S | 0.756 | 0.008983762 | Down | Bruchpilot, isoform J OS=Drosophila melanogaster OX=7227 GN=brp PE=1 SV=1 | brp |
| A0A0B4K7K9 | 889 | S | 0.736 | 0.007517903 | Down | Bruchpilot, isoform J OS=Drosophila melanogaster OX=7227 GN=brp PE=1 SV=1 | brp |
| B7YZD8 | 1066 | S | 1.319 | 0.024827537 | Up | CAP, isoform Q OS=Drosophila melanogaster OX=7227 GN=CAP PE=4 SV=2 | CAP |
| B7YZD8 | 1733 | S | 1.388 | 0.002741192 | Up | CAP, isoform Q OS=Drosophila melanogaster OX=7227 GN=CAP PE=4 SV=2 | CAP |
| B7YZD8 | 1637 | S | 1.346 | 0.049562346 | Up | CAP, isoform Q OS=Drosophila melanogaster OX=7227 GN=CAP PE=4 SV=2 | CAP |
| Q9VU14 | 282 | S | 1.131 | 0.4748546 | Up | Autophagy-related 1, isoform A OS=Drosophila melanogaster OX=7227 GN=Atg1 PE=2 SV=1 | Atg1 |
| Q9VU14 | 292 | T | 1.053 | 0.4847272 | Up | Autophagy-related 1, isoform A OS=Drosophila melanogaster OX=7227 GN=Atg1 PE=2 SV=1 | Atg1 |
| A1Z8W9 | 588 | S | 1.343 | 0.020480154 | Up | Gartenzwerg, isoform A OS=Drosophila melanogaster OX=7227 GN=garz PE=1 SV=1 | garz |
| Q9VCH4 | 199 | S | 2.28 | 0.003183118 | Up | Myoblast city OS=Drosophila melanogaster OX=7227 GN=mbc PE=1 SV=2 | mbc |
| Q9VCH4 | 747 | S | 2.317 | 1.8949E-06 | Up | Myoblast city OS=Drosophila melanogaster OX=7227 GN=mbc PE=1 SV=2 | mbc |
| Q9VCH4 | 1871 | S | 1.869 | 0.000674945 | Up | Myoblast city OS=Drosophila melanogaster OX=7227 GN=mbc PE=1 SV=2 | mbc |
| Q9VCH4 | 1873 | S | 1.819 | 0.000715221 | Up | Myoblast city OS=Drosophila melanogaster OX=7227 GN=mbc PE=1 SV=2 | mbc |
| A1ZAN7 | 1008 | S | 1.842 | 0.000489072 | Up | Rho guanine nucleotide exchange factor 2, isoform E OS=Drosophila melanogaster OX=7227 GN=RhoGEF2 PE=1 SV=1 | RhoGEF2 |
| A1ZAN7 | 1012 | S | 2.526 | 0.001014044 | Up | Rho guanine nucleotide exchange factor 2, isoform E OS=Drosophila melanogaster OX=7227 GN=RhoGEF2 PE=1 SV=1 | RhoGEF2 |
| A1ZAN7 | 1976 | S | 1.639 | 0.010942829 | Up | Rho guanine nucleotide exchange factor 2, isoform E OS=Drosophila melanogaster OX=7227 GN=RhoGEF2 PE=1 SV=1 | RhoGEF2 |
| A1ZAN7 | 464 | S | 1.878 | 0.000329824 | Up | Rho guanine nucleotide exchange factor 2, isoform E OS=Drosophila melanogaster OX=7227 GN=RhoGEF2 PE=1 SV=1 | RhoGEF2 |
| A0A0B4KEE5 | 1696 | S | 0.731 | 0.023050404 | Down | Uncharacterized protein, isoform Q OS=Drosophila melanogaster OX=7227 GN=CG2088 PE=1 SV=1 | CG2088 |
| A0A0B4KEE5 | 1349 | S | 0.638 | 0.021947074 | Down | Uncharacterized protein, isoform Q OS=Drosophila melanogaster OX=7227 GN=CG2088 PE=1 SV=1 | CG2088 |
| A0A0B4KEE5 | 1354 | S | 0.719 | 0.001602077 | Down | Uncharacterized protein, isoform Q OS=Drosophila melanogaster OX=7227 GN=CG2088 PE=1 SV=1 | CG2088 |
| A0A0B4K7V4 | 673 | S | 1.447 | 0.028323888 | Up | Transporter OS=Drosophila melanogaster OX=7227 GN=Dmel\CG43066 PE=3 SV=1 | Dmel\CG43066 |
| A0A0B4K7V4 | 676 | S | 1.603 | 0.000510092 | Up | Transporter OS=Drosophila melanogaster OX=7227 GN=Dmel\CG43066 PE=3 SV=1 | Dmel\CG43066 |
| A0A0B4K7V4 | 632 | S | 1.366 | 0.003373463 | Up | Transporter OS=Drosophila melanogaster OX=7227 GN=Dmel\CG43066 PE=3 SV=1 | Dmel\CG43066 |
| A0A0B4K7Y7 | 1510 | S | 1.602 | 0.041748416 | Up | La related protein, isoform G OS=Drosophila melanogaster OX=7227 GN=larp PE=1 SV=1 | larp |
| A0A0B4K7Y7 | 562 | S | 1.306 | 0.008452874 | Up | La related protein, isoform G OS=Drosophila melanogaster OX=7227 GN=larp PE=1 SV=1 | larp |
| A0A0B4LF25 | 837 | S | 1.579 | 0.013270656 | Up | Prosap, isoform E OS=Drosophila melanogaster OX=7227 GN=Prosap PE=4 SV=1 | Prosap |
| A0A0B4LF25 | 849 | S | 1.579 | 0.013270656 | Up | Prosap, isoform E OS=Drosophila melanogaster OX=7227 GN=Prosap PE=4 SV=1 | Prosap |
| A0A0B4LF25 | 863 | S | 1.41 | 0.010782909 | Up | Prosap, isoform E OS=Drosophila melanogaster OX=7227 GN=Prosap PE=4 SV=1 | Prosap |
| A0A0B4LF25 | 873 | S | 1.41 | 0.010782909 | Up | Prosap, isoform E OS=Drosophila melanogaster OX=7227 GN=Prosap PE=4 SV=1 | Prosap |
| A0A0B4LF25 | 879 | S | 1.349 | 0.027384194 | Up | Prosap, isoform E OS=Drosophila melanogaster OX=7227 GN=Prosap PE=4 SV=1 | Prosap |
| A1Z9J3 | 8746 | S | 1.719 | 0.030328452 | Up | Short stop, isoform H OS=Drosophila melanogaster OX=7227 GN=shot PE=1 SV=1 | shot |
| A0A0B4K883 | 75 | S | 1.301 | 0.04236437 | Up | Painting of fourth, isoform C OS=Drosophila melanogaster OX=7227 GN=Pof PE=4 SV=1 | Pof |
| A1Z6Q0 | 81 | S | 1.464 | 0.016583288 | Up | Klaroid, isoform D OS=Drosophila melanogaster OX=7227 GN=koi PE=1 SV=1 | koi |
| A1Z6Q0 | 235 | S | 1.319 | 0.033929678 | Up | Klaroid, isoform D OS=Drosophila melanogaster OX=7227 GN=koi PE=1 SV=1 | koi |
| P92177 | 46 | S | 1.79 | 0.001049647 | Up | 14-3-3 protein epsilon OS=Drosophila melanogaster OX=7227 GN=14-3-3epsilon PE=1 SV=2 | 14-3-3epsilon |
| E1JH23 | 10 | S | 2.324 | 0.029849358 | Up | Oxysterol-binding protein OS=Drosophila melanogaster OX=7227 GN=Dmel\CG1513 PE=1 SV=1 | Dmel\CG1513 |
| A0A0B4KEJ0 | 184 | S | 1.343 | 0.003424623 | Up | Adh transcription factor 1, isoform E OS=Drosophila melanogaster OX=7227 GN=Adf1 PE=1 SV=1 | Adf1 |
| E1JH51 | 929 | S | 1.722 | 0.015883183 | Up | Suppressor 2 of zeste h29 OS=Drosophila melanogaster OX=7227 GN=Su(z)2 PE=4 SV=1 | Su(z)2 |
| A0A0B4KES0 | 1680 | S | 1.441 | 0.009036632 | Up | Slit, isoform F OS=Drosophila melanogaster OX=7227 GN=sli PE=4 SV=1 | sli |
| A0A0B4KFU3 | 365 | S | 1.403 | 0.014292191 | Up | Patronin, isoform M OS=Drosophila melanogaster OX=7227 GN=Patronin PE=1 SV=1 | Patronin |
| A0A0B4KFU3 | 380 | S | 1.403 | 0.014292191 | Up | Patronin, isoform M OS=Drosophila melanogaster OX=7227 GN=Patronin PE=1 SV=1 | Patronin |
| A0A0B4KG58 | 402 | S | 2.495 | 0.030086594 | Up | Patronin, isoform L OS=Drosophila melanogaster OX=7227 GN=Patronin PE=1 SV=1 | Patronin |
| A0A0B4KEY9 | 778 | S | 1.494 | 0.031652638 | Up | Activated Cdc42 kinase-like, isoform C OS=Drosophila melanogaster OX=7227 GN=Ack-like PE=3 SV=1 | Ack-like |
| A0A0B4KEY9 | 691 | S | 1.481 | 0.003196343 | Up | Activated Cdc42 kinase-like, isoform C OS=Drosophila melanogaster OX=7227 GN=Ack-like PE=3 SV=1 | Ack-like |
| A0A0B4KFI2 | 442 | S | 1.427 | 0.001444069 | Up | Painless, isoform C OS=Drosophila melanogaster OX=7227 GN=pain PE=4 SV=1 | pain |
| A0A0B4KF90 | 294 | S | 3.129 | 0.011259044 | Up | Like-AP180, isoform J OS=Drosophila melanogaster OX=7227 GN=lap PE=1 SV=1 | lap |
| A0A0B4KF92 | 189 | S | 0.652 | 0.00087315 | Down | Holocarboxylase synthetase, isoform B OS=Drosophila melanogaster OX=7227 GN=Hcs PE=1 SV=1 | Hcs |
| Q0E9C6 | 13 | S | 1.479 | 0.002075564 | Up | FI05595p OS=Drosophila melanogaster OX=7227 GN=Dmel\CG30015 PE=1 SV=1 | Dmel\CG30015 |
| Q0E9C6 | 1146 | S | 1.517 | 0.000351443 | Up | FI05595p OS=Drosophila melanogaster OX=7227 GN=Dmel\CG30015 PE=1 SV=1 | Dmel\CG30015 |

| | | | | | | | |
|---|---|---|---|---|---|---|---|
| Q0E9C6 | 330 | S | 1.317 | 0.000586521 | Up | FI05595p OS=Drosophila melanogaster OX=7227 GN=Dmel\CG30015 PE=1 SV=1 | Dmel\CG30015 |
| Q0E9C6 | 1051 | S | 1.436 | 0.000382023 | Up | FI05595p OS=Drosophila melanogaster OX=7227 GN=Dmel\CG30015 PE=1 SV=1 | Dmel\CG30015 |
| Q0E9C6 | 1042 | S | 1.563 | 0.016507344 | Up | FI05595p OS=Drosophila melanogaster OX=7227 GN=Dmel\CG30015 PE=1 SV=1 | Dmel\CG30015 |
| Q7K3G2 | 71 | S | 1.345 | 0.006134633 | Up | LD29875p OS=Drosophila melanogaster OX=7227 GN=Dmel\CG5742 PE=2 SV=1 | Dmel\CG5742 |
| A0A0B4KFC5 | 324 | S | 1.707 | 0.013870867 | Up | Lingerer, isoform K OS=Drosophila melanogaster OX=7227 GN=lig PE=1 SV=1 | lig |
| A1Z7C4 | 308 | S | 1.746 | 0.000112749 | Up | LDL receptor protein 1, isoform G OS=Drosophila melanogaster OX=7227 GN=LRP1 PE=1 SV=2 | LRP1 |
| A8JQV2 | 149 | S | 1.419 | 0.013408183 | Up | Uncharacterized protein, isoform E OS=Drosophila melanogaster OX=7227 GN=Dmel\CG17816 PE=1 SV=1 | Dmel\CG17816 |
| A8JQV2 | 112 | S | 1.319 | 0.005327561 | Up | Uncharacterized protein, isoform E OS=Drosophila melanogaster OX=7227 GN=Dmel\CG17816 PE=1 SV=1 | Dmel\CG17816 |
| A8JQV2 | 414 | S | 1.451 | 0.013721784 | Up | Uncharacterized protein, isoform E OS=Drosophila melanogaster OX=7227 GN=Dmel\CG17816 PE=1 SV=1 | Dmel\CG17816 |
| Q9W205 | 880 | S | 1.315 | 0.000221034 | Up | Jitterbug, isoform F OS=Drosophila melanogaster OX=7227 GN=jbug PE=1 SV=5 | jbug |
| Q9W205 | 996 | S | 1.343 | 0.031948239 | Up | Jitterbug, isoform F OS=Drosophila melanogaster OX=7227 GN=jbug PE=1 SV=5 | jbug |
| Q9W205 | 999 | S | 1.496 | 0.008844939 | Up | Jitterbug, isoform F OS=Drosophila melanogaster OX=7227 GN=jbug PE=1 SV=5 | jbug |
| A0A0B4KFH8 | 2666 | S | 1.431 | 0.03318578 | Up | Sodium channel protein OS=Drosophila melanogaster OX=7227 GN=NaCP60E PE=3 SV=1 | NaCP60E |
| Q9VH03 | 598 | S | 1.674 | 0.012325353 | Up | Nuak1 ortholog, isoform E OS=Drosophila melanogaster OX=7227 GN=Nuak1 PE=4 SV=2 | Nuak1 |
| A0A0B4KG24 | 1374 | S | 1.805 | 0.014954 | Up | Kinesin heavy chain 73, isoform D OS=Drosophila melanogaster OX=7227 GN=Khc-73 PE=1 SV=1 | Khc-73 |
| A0A0B4KFU3 | 1357 | S | 1.759 | 0.01667201 | Up | Patronin, isoform M OS=Drosophila melanogaster OX=7227 GN=Patronin PE=1 SV=1 | Patronin |
| Q8INH2 | 292 | S | 2.538 | 0.002391066 | Up | GH23935p OS=Drosophila melanogaster OX=7227 GN=Dmel\CG9813 PE=1 SV=1 | Dmel\CG9813 |
| A0A0B4LH87 | 1075 | S | 1.46 | 0.003979375 | Up | Phosphodiesterase OS=Drosophila melanogaster OX=7227 GN=Pde6 PE=3 SV=1 | Pde6 |
| A0A0B4LH87 | 1078 | S | 1.695 | 0.001832662 | Up | Phosphodiesterase OS=Drosophila melanogaster OX=7227 GN=Pde6 PE=3 SV=1 | Pde6 |
| A0A0B4LH87 | 187 | S | 1.886 | 0.003007487 | Up | Phosphodiesterase OS=Drosophila melanogaster OX=7227 GN=Pde6 PE=3 SV=1 | Pde6 |
| A0A0B4LH87 | 1050 | S | 1.364 | 0.013136857 | Up | Phosphodiesterase OS=Drosophila melanogaster OX=7227 GN=Pde6 PE=3 SV=1 | Pde6 |
| A0A0B4KGP0 | 451 | S | 1.889 | 0.002060541 | Up | Stumps, isoform F OS=Drosophila melanogaster OX=7227 GN=stumps PE=4 SV=1 | stumps |
| A0A0B4KGP0 | 452 | S | 1.639 | 0.002067272 | Up | Stumps, isoform F OS=Drosophila melanogaster OX=7227 GN=stumps PE=4 SV=1 | stumps |
| A0A0B4KGP0 | 454 | S | 1.889 | 0.002060541 | Up | Stumps, isoform F OS=Drosophila melanogaster OX=7227 GN=stumps PE=4 SV=1 | stumps |
| A0A0B4KGP0 | 543 | S | 1.612 | 0.015444914 | Up | Stumps, isoform F OS=Drosophila melanogaster OX=7227 GN=stumps PE=4 SV=1 | stumps |
| Q7K0G4 | 261 | S | 2.033 | 0.001690527 | Up | SD10629p OS=Drosophila melanogaster OX=7227 GN=Spred PE=2 SV=1 | Spred |
| Q8IPP4 | 1218 | S | 1.614 | 0.015390026 | Up | Canoe, isoform C OS=Drosophila melanogaster OX=7227 GN=cno PE=1 SV=1 | cno |
| A0A0B4KG76 | 271 | S | 1.305 | 0.010924578 | Up | Pollux, isoform F OS=Drosophila melanogaster OX=7227 GN=plx PE=4 SV=1 | plx |
| A0A0B4KG81 | 1757 | S | 1.337 | 0.001803516 | Up | Uncharacterized protein, isoform B OS=Drosophila melanogaster OX=7227 GN=Dmel\CG15099 PE=4 SV=1 | Dmel\CG15099 |
| A0A0B4KG81 | 815 | S | 1.328 | 0.003506486 | Up | Uncharacterized protein, isoform B OS=Drosophila melanogaster OX=7227 GN=Dmel\CG15099 PE=4 SV=1 | Dmel\CG15099 |
| Q9VCL1 | 186 | S | 1.357 | 0.026918407 | Up | Beaten path IV, isoform A OS=Drosophila melanogaster OX=7227 GN=beat-IV PE=1 SV=2 | beat-IV |
| A0A0B4KH86 | 108 | S | 1.97 | 0.000543331 | Up | Inwardly rectifying potassium channel 1, isoform F OS=Drosophila melanogaster OX=7227 GN=Irk1 PE=1 SV=1 | Irk1 |
| A0A0B4KH86 | 515 | S | 1.334 | 0.009276766 | Up | Inwardly rectifying potassium channel 1, isoform F OS=Drosophila melanogaster OX=7227 GN=Irk1 PE=1 SV=1 | Irk1 |
| Q8MQS2 | 342 | S | 1.72 | 0.004602476 | Up | GH11945p OS=Drosophila melanogaster OX=7227 GN=jus PE=2 SV=1 | jus |
| Q8MQS2 | 347 | S | 2.155 | 0.001382365 | Up | GH11945p OS=Drosophila melanogaster OX=7227 GN=jus PE=2 SV=1 | jus |
| B7Z0L6 | 672 | S | 1.441 | 0.005531342 | Up | Serine/threonine protein phosphatase 2A regulatory subunit OS=Drosophila melanogaster OX=7227 GN=wrd PE=1 SV=1 | wrd |
| A0A0B4KHE1 | 204 | S | 1.867 | 1.54409E-05 | Up | Uncharacterized protein, isoform B OS=Drosophila melanogaster OX=7227 GN=Dmel\CG15514 PE=4 SV=1 | Dmel\CG15514 |
| A0A0B4KHE9 | 124 | S | 1.355 | 0.002344909 | Up | 26S proteasome non-ATPase regulatory subunit 1 OS=Drosophila melanogaster OX=7227 GN=Rpn2 PE=3 SV=1 | Rpn2 |
| A0A0B4KHJ5 | 6 | S | 2.341 | 0.009782134 | Up | Glycogen [starch] synthase OS=Drosophila melanogaster OX=7227 GN=GlyS PE=1 SV=1 | GlyS |
| Q9V9Y0 | 16 | S | 1.563 | 0.018614611 | Up | GH11014p OS=Drosophila melanogaster OX=7227 GN=Dmel\CG1607 PE=1 SV=1 | Dmel\CG1607 |
| A0A0B4KHR3 | 390 | S | 1.413 | 0.00497033 | Up | Uncharacterized protein, isoform E OS=Drosophila melanogaster OX=7227 GN=Dmel\CG13604 PE=4 SV=1 | Dmel\CG13604 |
| Q9VAY4 | 365 | S | 1.655 | 0.001212057 | Up | Biorientation defective 1, isoform A OS=Drosophila melanogaster OX=7227 GN=BOD1 PE=1 SV=2 | BOD1 |
| A0A0B4KI34 | 598 | S | 1.535 | 0.002280587 | Up | CIN85 and CD2AP related, isoform J OS=Drosophila melanogaster OX=7227 GN=cindr PE=1 SV=1 | cindr |
| A0A0B4KI34 | 542 | S | 1.424 | 0.005413478 | Up | CIN85 and CD2AP related, isoform J OS=Drosophila melanogaster OX=7227 GN=cindr PE=1 SV=1 | cindr |
| A0A0B4KI38 | 12 | S | 1.504 | 0.040882419 | Up | Methylthioribose-1-phosphate isomerase OS=Drosophila melanogaster OX=7227 GN=Dmel\CG11334 PE=3 SV=1 | Dmel\CG11334 |
| A0A0B4KI69 | 612 | S | 1.329 | 0.003091603 | Up | G protein-coupled receptor kinase OS=Drosophila melanogaster OX=7227 GN=Gprk2 PE=3 SV=1 | Gprk2 |
| Q8IMF5 | 354 | S | 1.452 | 0.003277669 | Up | Microtubule-associated protein 205, isoform B OS=Drosophila melanogaster OX=7227 GN=Map205 PE=1 SV=1 | Map205 |
| Q8IMF5 | 516 | S | 1.645 | 7.73347E-05 | Up | Microtubule-associated protein 205, isoform B OS=Drosophila melanogaster OX=7227 GN=Map205 PE=1 SV=1 | Map205 |
| A0A0B4LEV8 | 334 | S | 1.503 | 0.000858738 | Up | Coilin, isoform E OS=Drosophila melanogaster OX=7227 GN=coil PE=1 SV=1 | coil |
| A0A0B4LEW1 | 846 | S | 1.857 | 0.001590455 | Up | Uncharacterized protein, isoform E OS=Drosophila melanogaster OX=7227 GN=BcDNA:GH05582 PE=4 SV=1 | BcDNA:GH05582 |
| A0A0B4LEW1 | 1025 | S | 1.352 | 0.014434163 | Up | Uncharacterized protein, isoform E OS=Drosophila melanogaster OX=7227 GN=BcDNA:GH05582 PE=4 SV=1 | BcDNA:GH05582 |
| A0A0B4LEW1 | 1027 | S | 1.352 | 0.014434163 | Up | Uncharacterized protein, isoform E OS=Drosophila melanogaster OX=7227 GN=BcDNA:GH05582 PE=4 SV=1 | BcDNA:GH05582 |
| A0A0B4LEW1 | 1029 | S | 1.352 | 0.014434163 | Up | Uncharacterized protein, isoform E OS=Drosophila melanogaster OX=7227 GN=BcDNA:GH05582 PE=4 SV=1 | BcDNA:GH05582 |
| A0A0B4LEW1 | 630 | S | 1.301 | 0.008166312 | Up | Uncharacterized protein, isoform E OS=Drosophila melanogaster OX=7227 GN=BcDNA:GH05582 PE=4 SV=1 | BcDNA:GH05582 |
| A0A0B4LEW1 | 841 | S | 1.893 | 0.00146471 | Up | Uncharacterized protein, isoform E OS=Drosophila melanogaster OX=7227 GN=BcDNA:GH05582 PE=4 SV=1 | BcDNA:GH05582 |
| A0A0B4LEW1 | 614 | S | 1.469 | 0.0050095 | Up | Uncharacterized protein, isoform E OS=Drosophila melanogaster OX=7227 GN=BcDNA:GH05582 PE=4 SV=1 | BcDNA:GH05582 |
| A0A0B4LEW1 | 1140 | S | 1.621 | 0.030181414 | Up | Uncharacterized protein, isoform E OS=Drosophila melanogaster OX=7227 GN=BcDNA:GH05582 PE=4 SV=1 | BcDNA:GH05582 |
| A1Z7J7 | 492 | S | 1.303 | 0.006458651 | Up | Secretory 31, isoform A OS=Drosophila melanogaster OX=7227 GN=Sec31 PE=1 SV=1 | Sec31 |
| A0A0B4LEY5 | 73 | S | 1.575 | 0.0209426 | Up | Protein kinase N, isoform O OS=Drosophila melanogaster OX=7227 GN=Pkn PE=1 SV=1 | Pkn |
| A1Z9M4 | 96 | S | 1.657 | 0.001792187 | Up | Combgap, isoform C OS=Drosophila melanogaster OX=7227 GN=cg | cg |

| | | | | | | | |
|---|---|---|---|---|---|---|---|
| | | | | | | PE=1 SV=1 | |
| A0A0B4LF57 | 82 | S | 0.748 | 0.00928407 | Down | Calmodulin, isoform C OS=Drosophila melanogaster OX=7227 GN=Cam PE=1 SV=1 | Cam |
| A0A0B4LF97 | 1291 | S | 1.414 | 0.019872981 | Up | Bedraggled, isoform D OS=Drosophila melanogaster OX=7227 GN=bdg PE=4 SV=1 | bdg |
| A0A0B4LFB8 | 683 | S | 0.713 | 0.00564248 | Down | Optic atrophy 1, isoform D OS=Drosophila melanogaster OX=7227 GN=Opa1 PE=1 SV=1 | Opa1 |
| A0A0B4LFK0 | 11 | S | 2.398 | 2.1113E-05 | Up | Picot, isoform C OS=Drosophila melanogaster OX=7227 GN=Picot PE=4 SV=1 | Picot |
| A0A0B4LFK0 | 4 | S | 2.576 | 0.000247793 | Up | Picot, isoform C OS=Drosophila melanogaster OX=7227 GN=Picot PE=4 SV=1 | Picot |
| A0A0B4LFP7 | 548 | S | 1.346 | 0.009428116 | Up | Thin, isoform D OS=Drosophila melanogaster OX=7227 GN=tn PE=1 SV=1 | tn |
| A0A0B4LFS3 | 586 | S | 1.896 | 0.018343759 | Up | Uncharacterized protein, isoform B OS=Drosophila melanogaster OX=7227 GN=Naus PE=4 SV=1 | Naus |
| A0A0B4LFS3 | 580 | S | 1.625 | 0.003464142 | Up | Uncharacterized protein, isoform B OS=Drosophila melanogaster OX=7227 GN=Naus PE=4 SV=1 | Naus |
| A0A0B4LFS3 | 582 | S | 1.611 | 0.007014189 | Up | Uncharacterized protein, isoform B OS=Drosophila melanogaster OX=7227 GN=Naus PE=4 SV=1 | Naus |
| A0A0B4LFV0 | 759 | S | 1.332 | 0.000549449 | Up | Non-specific serine/threonine protein kinase OS=Drosophila melanogaster OX=7227 GN=hppy PE=1 SV=1 | hppy |
| A0A0B4LFV0 | 760 | S | 1.315 | 0.000475735 | Up | Non-specific serine/threonine protein kinase OS=Drosophila melanogaster OX=7227 GN=hppy PE=1 SV=1 | hppy |
| A0A0B4LG23 | 488 | S | 1.99 | 0.005187264 | Up | Coracle, isoform G OS=Drosophila melanogaster OX=7227 GN=cora PE=1 SV=1 | cora |
| A0A0B4LG23 | 329 | S | 2.567 | 0.002191996 | Up | Coracle, isoform G OS=Drosophila melanogaster OX=7227 GN=cora PE=1 SV=1 | cora |
| A0A0B4LG23 | 566 | S | 1.362 | 0.002168962 | Up | Coracle, isoform G OS=Drosophila melanogaster OX=7227 GN=cora PE=1 SV=1 | cora |
| A0A0B4LG23 | 366 | S | 1.885 | 0.001311961 | Up | Coracle, isoform G OS=Drosophila melanogaster OX=7227 GN=cora PE=1 SV=1 | cora |
| A0A0B4LG36 | 102 | S | 0.726 | 0.032262606 | Down | HMG protein Z, isoform C OS=Drosophila melanogaster OX=7227 GN=HmgZ PE=4 SV=1 | HmgZ |
| A0A0B4LGB0 | 925 | S | 1.34 | 0.004898407 | Up | Gp150, isoform E OS=Drosophila melanogaster OX=7227 GN=Gp150 PE=1 SV=1 | Gp150 |
| A0A0B4LGB7 | 346 | S | 1.674 | 0.005336477 | Up | Calcium-transporting ATPase OS=Drosophila melanogaster OX=7227 GN=SERCA PE=1 SV=1 | SERCA |
| A0A0B4LGG4 | 155 | S | 1.344 | 0.006308999 | Up | Sestrin, isoform D OS=Drosophila melanogaster OX=7227 GN=Sesn PE=4 SV=1 | Sesn |
| A0A0B4LGI9 | 258 | S | 1.447 | 0.015660027 | Up | Uncharacterized protein, isoform D OS=Drosophila melanogaster OX=7227 GN=anon-EST:fe2C9 PE=1 SV=1 | anon-EST:fe 2C9 |
| A0A0B4LGW2 | 827 | S | 1.321 | 0.032840055 | Up | Eag-like K[+] channel, isoform B OS=Drosophila melanogaster OX=7227 GN=Elk PE=4 SV=1 | Elk |
| Q9VH91 | 371 | S | 1.789 | 0.010998557 | Up | DUNC-115l OS=Drosophila melanogaster OX=7227 GN=Unc-115a PE=2 SV=4 | Unc-115a |
| Q9VH91 | 501 | S | 1.366 | 0.009331209 | Up | DUNC-115l OS=Drosophila melanogaster OX=7227 GN=Unc-115a PE=2 SV=4 | Unc-115a |
| Q9VH91 | 735 | S | 1.432 | 0.017145199 | Up | DUNC-115l OS=Drosophila melanogaster OX=7227 GN=Unc-115a PE=2 SV=4 | Unc-115a |
| Q9VH91 | 737 | S | 1.586 | 0.005993411 | Up | DUNC-115l OS=Drosophila melanogaster OX=7227 GN=Unc-115a PE=2 SV=4 | Unc-115a |
| Q9VH91 | 625 | S | 1.43 | 0.018048137 | Up | DUNC-115l OS=Drosophila melanogaster OX=7227 GN=Unc-115a PE=2 SV=4 | Unc-115a |
| Q9VH91 | 341 | S | 1.565 | 0.000566905 | Up | DUNC-115l OS=Drosophila melanogaster OX=7227 GN=Unc-115a PE=2 SV=4 | Unc-115a |
| A0A0B4LH71 | 219 | S | 1.379 | 0.00443457 | Up | Synapse-associated protein 47kD, isoform K OS=Drosophila melanogaster OX=7227 GN=Sap47 PE=1 SV=1 | Sap47 |
| A0A0B4LH99 | 615 | S | 1.513 | 0.000839303 | Up | Uncharacterized protein, isoform B OS=Drosophila melanogaster OX=7227 GN=Dmel\CG7218 PE=4 SV=1 | Dmel\CG721 8 |
| Q8IN24 | 740 | S | 1.363 | 0.000826195 | Up | Metabotropic GABA-B receptor subtype 2, isoform A OS=Drosophila melanogaster OX=7227 GN=GABA-B-R2 PE=3 SV=1 | GABA-B-R2 |
| Q8IN24 | 1206 | S | 1.439 | 0.01024477 | Up | Metabotropic GABA-B receptor subtype 2, isoform A OS=Drosophila melanogaster OX=7227 GN=GABA-B-R2 PE=3 SV=1 | GABA-B-R2 |
| A0A0B4LIP5 | 137 | S | 1.433 | 0.010852363 | Up | Lnk, isoform F OS=Drosophila melanogaster OX=7227 GN=Lnk PE=4 SV=1 | Lnk |
| A0A0B4LJ12 | 45 | S | 0.581 | 0.000314937 | Down | Nucleoside diphosphate kinase OS=Drosophila melanogaster OX=7227 GN=awd PE=1 SV=1 | awd |
| Q4QQA3 | 169 | S | 1.638 | 0.011159317 | Up | Protein kinase C OS=Drosophila melanogaster OX=7227 GN=Pkc98E PE=2 SV=1 | Pkc98E |
| Q9VE88 | 531 | S | 1.432 | 0.004728975 | Up | Uncharacterized protein, isoform B OS=Drosophila melanogaster OX=7227 GN=lincRNA.804 PE=4 SV=3 | lincRNA.804 |
| Q9VB13 | 184 | S | 1.424 | 0.00543994 | Up | Microtubule-associated protein OS=Drosophila melanogaster OX=7227 GN=tau PE=1 SV=2 | tau |
| Q9VB13 | 113 | S | 0.742 | 0.021581965 | Down | Microtubule-associated protein OS=Drosophila melanogaster OX=7227 GN=tau PE=1 SV=2 | tau |
| Q9VB13 | 343 | S | 1.635 | 0.000796627 | Up | Microtubule-associated protein OS=Drosophila melanogaster OX=7227 GN=tau PE=1 SV=2 | tau |
| Q9VB13 | 135 | S | 2.279 | 0.004140988 | Up | Microtubule-associated protein OS=Drosophila melanogaster OX=7227 GN=tau PE=1 SV=2 | tau |
| A0A0B7P9G0 | 712 | S | 1.523 | 0.018045778 | Up | Unextended, isoform E OS=Drosophila melanogaster OX=7227 GN=uex PE=1 SV=1 | uex |
| A0A0B7P9G0 | 753 | S | 1.433 | 0.035809595 | Up | Unextended, isoform E OS=Drosophila melanogaster OX=7227 GN=uex PE=1 SV=1 | uex |
| A0A0C4DHA8 | 259 | S | 1.5 | 0.033258663 | Up | Uncharacterized protein, isoform I OS=Drosophila melanogaster OX=7227 GN=Dmel\CG17816 PE=1 SV=1 | Dmel\CG178 16 |
| Q0KHZ4 | 608 | S | 1.577 | 0.027006731 | Up | Uncharacterized protein, isoform B OS=Drosophila melanogaster OX=7227 GN=Dmel\CG34133 PE=1 SV=1 | Dmel\CG341 33 |
| Q0KHZ4 | 151 | S | 1.75 | 0.004895028 | Up | Uncharacterized protein, isoform B OS=Drosophila melanogaster OX=7227 GN=Dmel\CG34133 PE=1 SV=1 | Dmel\CG341 33 |
| Q0KHZ4 | 154 | S | 1.75 | 0.004895028 | Up | Uncharacterized protein, isoform B OS=Drosophila melanogaster OX=7227 GN=Dmel\CG34133 PE=1 SV=1 | Dmel\CG341 33 |
| Q0KHZ4 | 404 | S | 1.381 | 0.024269602 | Up | Uncharacterized protein, isoform B OS=Drosophila melanogaster OX=7227 GN=Dmel\CG34133 PE=1 SV=1 | Dmel\CG341 33 |
| A1ZA89 | 828 | S | 1.486 | 0.00015755 | Up | Dystroglycan, isoform B OS=Drosophila melanogaster OX=7227 GN=Dg PE=1 SV=1 | Dg |
| Q9VJX4 | 32 | S | 1.402 | 0.002838757 | Up | RH51758p OS=Drosophila melanogaster OX=7227 GN=CG16851 PE=2 SV=3 | CG16851 |
| A0A0S0WGX2 | 320 | S | 0.647 | 0.000468995 | Down | Uncharacterized protein OS=Drosophila melanogaster OX=7227 GN=lovit PE=4 SV=1 | lovit |
| A0A0S0WGX2 | 323 | S | 0.67 | 0.004454633 | Down | Uncharacterized protein OS=Drosophila melanogaster OX=7227 GN=lovit PE=4 SV=1 | lovit |
| Q8IQ62 | 1393 | S | 0.762 | 0.017272945 | Down | Still life, isoform F OS=Drosophila melanogaster OX=7227 GN=sif PE=1 SV=2 | sif |
| A0A0S0WIA2 | 891 | S | 1.816 | 0.012309761 | Up | Multidrug-Resistance like protein 1, isoform R OS=Drosophila melanogaster OX=7227 GN=MRP PE=1 SV=1 | MRP |
| A0A0S0WIA2 | 892 | S | 1.625 | 0.003851884 | Up | Multidrug-Resistance like protein 1, isoform R OS=Drosophila melanogaster OX=7227 GN=MRP PE=1 SV=1 | MRP |
| A0A0S0X7U5 | 1383 | S | 1.982 | 0.00036618 | Up | Rab3 GDP-GTP exchange factor, isoform I OS=Drosophila melanogaster OX=7227 GN=Rab3-GEF PE=4 SV=1 | Rab3-GEF |
| A0A0S0X7U5 | 1384 | S | 1.381 | 0.000568166 | Up | Rab3 GDP-GTP exchange factor, isoform I OS=Drosophila melanogaster OX=7227 GN=Rab3-GEF PE=4 SV=1 | Rab3-GEF |
| A0A0S0X7U5 | 1387 | S | 1.363 | 0.001165558 | Up | Rab3 GDP-GTP exchange factor, isoform I OS=Drosophila melanogaster OX=7227 GN=Rab3-GEF PE=4 SV=1 | Rab3-GEF |
| A0A0S0X7U5 | 899 | S | 1.693 | 0.001365181 | Up | Rab3 GDP-GTP exchange factor, isoform I OS=Drosophila melanogaster OX=7227 GN=Rab3-GEF PE=4 SV=1 | Rab3-GEF |
| A0A0S0X7U5 | 1414 | S | 1.434 | 0.004466481 | Up | Rab3 GDP-GTP exchange factor, isoform I OS=Drosophila melanogaster | Rab3-GEF |

| | | | | | | | |
|---|---|---|---|---|---|---|---|
| | | | | | | OX=7227 GN=Rab3-GEF PE=4 SV=1 | |
| A0A0S0X7U5 | 172 | S | 1.476 | 0.025362613 | Up | Rab3 GDP-GTP exchange factor, isoform I OS=Drosophila melanogaster OX=7227 GN=Rab3-GEF PE=4 SV=1 | Rab3-GEF |
| A0A0S0X7U5 | 1932 | S | 1.405 | 1.41983E-05 | Up | Rab3 GDP-GTP exchange factor, isoform I OS=Drosophila melanogaster OX=7227 GN=Rab3-GEF PE=4 SV=1 | Rab3-GEF |
| A0A0S0X7W8 | 749 | S | 0.768 | 0.010858216 | Down | Protein tweety homolog OS=Drosophila melanogaster OX=7227 GN=Dmel\CG3638 PE=3 SV=1 | Dmel\CG3638 |
| A0A126GUN6 | 1364 | S | 1.372 | 0.030875506 | Up | Z band alternatively spliced PDZ-motif protein 52, isoform X OS=Drosophila melanogaster OX=7227 GN=Zasp52 PE=1 SV=1 | Zasp52 |
| A0A140SRF8 | 235 | S | 1.653 | 0.000246905 | Up | Uncharacterized protein, isoform C OS=Drosophila melanogaster OX=7227 GN=Dmel\CG11857 PE=1 SV=1 | Dmel\CG11857 |
| M9PCF6 | 362 | S | 1.398 | 0.008415668 | Up | Lilliputian, isoform H OS=Drosophila melanogaster OX=7227 GN=lilli PE=4 SV=1 | lilli |
| M9PH69 | 1915 | S | 1.352 | 0.023791257 | Up | Cyclic nucleotide-gated ion channel-like, isoform I OS=Drosophila melanogaster OX=7227 GN=Cngl PE=4 SV=2 | Cngl |
| A4V1P2 | 273 | S | 1.567 | 0.013449116 | Up | Orb2, isoform B OS=Drosophila melanogaster OX=7227 GN=orb2 PE=4 SV=1 | orb2 |
| Q9BKJ2 | 26 | S | 1.306 | 0.016975084 | Up | Four wheel drive, isoform C OS=Drosophila melanogaster OX=7227 GN=fwd PE=2 SV=2 | fwd |
| Q8IPA2 | 223 | S | 0.602 | 0.000131746 | Down | FI17312p1 OS=Drosophila melanogaster OX=7227 GN=Mal-B2 PE=1 SV=1 | Mal-B2 |
| A1A708 | 307 | S | 1.413 | 0.024607214 | Up | Uncharacterized protein CG4951 OS=Drosophila melanogaster OX=7227 GN=CG4951 PE=1 SV=1 | CG4951 |
| A1A708 | 251 | S | 1.34 | 0.043287752 | Up | Uncharacterized protein CG4951 OS=Drosophila melanogaster OX=7227 GN=CG4951 PE=1 SV=1 | CG4951 |
| A1Z6N4 | 525 | S | 1.968 | 0.000727965 | Up | MIP05841p OS=Drosophila melanogaster OX=7227 GN=Tdc2 PE=2 SV=1 | Tdc2 |
| A1Z6N4 | 530 | S | 1.903 | 0.005626296 | Up | MIP05841p OS=Drosophila melanogaster OX=7227 GN=Tdc2 PE=2 SV=1 | Tdc2 |
| A1Z6R7 | 218 | S | 1.657 | 0.00156938 | Up | FI21445p1 OS=Drosophila melanogaster OX=7227 GN=Dmel\CG30158 PE=2 SV=1 | Dmel\CG30158 |
| E1JGZ4 | 128 | S | 1.345 | 0.000723885 | Up | Blown fuse, isoform B OS=Drosophila melanogaster OX=7227 GN=blow PE=4 SV=1 | blow |
| E1JGZ4 | 131 | S | 1.345 | 0.000723885 | Up | Blown fuse, isoform B OS=Drosophila melanogaster OX=7227 GN=blow PE=4 SV=1 | blow |
| A1Z784 | 227 | S | 0.478 | 0.000145054 | Down | Acetyl-CoA carboxylase, isoform A OS=Drosophila melanogaster OX=7227 GN=ACC PE=1 SV=1 | ACC |
| A1Z784 | 1398 | S | 0.715 | 0.032743347 | Down | Acetyl-CoA carboxylase, isoform A OS=Drosophila melanogaster OX=7227 GN=ACC PE=1 SV=1 | ACC |
| A1Z784 | 1400 | S | 1.329 | 0.011214567 | Up | Acetyl-CoA carboxylase, isoform A OS=Drosophila melanogaster OX=7227 GN=ACC PE=1 SV=1 | ACC |
| E2QCN3 | 418 | S | 1.647 | 0.000132546 | Up | Alpha/beta hydrolase 1, isoform B OS=Drosophila melanogaster OX=7227 GN=Hydr1 PE=4 SV=1 | Hydr1 |
| Q95SG9 | 114 | S | 1.339 | 0.008538994 | Up | Alicorn, isoform B OS=Drosophila melanogaster OX=7227 GN=alc PE=2 SV=1 | alc |
| A1Z7S3 | 38 | S | 0.762 | 0.001048159 | Down | Rab32, isoform B OS=Drosophila melanogaster OX=7227 GN=Rab32 PE=1 SV=1 | Rab32 |
| A1Z7S3 | 191 | S | 1.325 | 0.030699715 | Up | Rab32, isoform B OS=Drosophila melanogaster OX=7227 GN=Rab32 PE=1 SV=1 | Rab32 |
| A1Z7Y7 | 530 | S | 1.844 | 0.00165432 | Up | Spaghetti-squash activator, isoform C OS=Drosophila melanogaster OX=7227 GN=sqa PE=4 SV=2 | sqa |
| A1Z8N1 | 118 | S | 1.33 | 0.000171829 | Up | Facilitated trehalose transporter Tret1-1 OS=Drosophila melanogaster OX=7227 GN=Tret1-1 PE=1 SV=1 | Tret1-1 |
| A1Z8N1 | 320 | S | 1.327 | 0.020267619 | Up | Facilitated trehalose transporter Tret1-1 OS=Drosophila melanogaster OX=7227 GN=Tret1-1 PE=1 SV=1 | Tret1-1 |
| A1Z8N1 | 322 | S | 1.327 | 0.020267619 | Up | Facilitated trehalose transporter Tret1-1 OS=Drosophila melanogaster OX=7227 GN=Tret1-1 PE=1 SV=1 | Tret1-1 |
| A1Z8N1 | 248 | S | 1.311 | 0.000427778 | Up | Facilitated trehalose transporter Tret1-1 OS=Drosophila melanogaster OX=7227 GN=Tret1-1 PE=1 SV=1 | Tret1-1 |
| A1Z8N1 | 249 | S | 1.369 | 0.000219667 | Up | Facilitated trehalose transporter Tret1-1 OS=Drosophila melanogaster OX=7227 GN=Tret1-1 PE=1 SV=1 | Tret1-1 |
| A1Z8N1 | 250 | S | 1.416 | 5.93683E-05 | Up | Facilitated trehalose transporter Tret1-1 OS=Drosophila melanogaster OX=7227 GN=Tret1-1 PE=1 SV=1 | Tret1-1 |
| A1Z8N1 | 268 | S | 1.607 | 0.022192504 | Up | Facilitated trehalose transporter Tret1-1 OS=Drosophila melanogaster OX=7227 GN=Tret1-1 PE=1 SV=1 | Tret1-1 |
| A1Z8N1 | 232 | S | 2.827 | 0.001063766 | Up | Facilitated trehalose transporter Tret1-1 OS=Drosophila melanogaster OX=7227 GN=Tret1-1 PE=1 SV=1 | Tret1-1 |
| A1Z8N1 | 845 | S | 1.822 | 0.003302056 | Up | Facilitated trehalose transporter Tret1-1 OS=Drosophila melanogaster OX=7227 GN=Tret1-1 PE=1 SV=1 | Tret1-1 |
| A1Z8N1 | 846 | S | 1.822 | 0.003302056 | Up | Facilitated trehalose transporter Tret1-1 OS=Drosophila melanogaster OX=7227 GN=Tret1-1 PE=1 SV=1 | Tret1-1 |
| A1Z8U2 | 86 | S | 1.517 | 0.011716046 | Up | Ornithine decarboxylase antizyme, isoform B OS=Drosophila melanogaster OX=7227 GN=Oda PE=4 SV=1 | Oda |
| A1Z8Y5 | 5 | S | 1.41 | 0.002384806 | Up | FI07730p OS=Drosophila melanogaster OX=7227 GN=Dmel\CG8501 PE=2 SV=1 | Dmel\CG8501 |
| A1Z968 | 389 | S | 1.896 | 0.007510641 | Up | NAT1, isoform D OS=Drosophila melanogaster OX=7227 GN=NAT1 PE=1 SV=1 | NAT1 |
| A1Z971 | 310 | S | 2.093 | 0.001400702 | Up | Uncharacterized protein, isoform A OS=Drosophila melanogaster OX=7227 GN=dRNF34 PE=1 SV=1 | dRNF34 |
| A1Z971 | 313 | S | 2.039 | 0.001145661 | Up | Uncharacterized protein, isoform A OS=Drosophila melanogaster OX=7227 GN=dRNF34 PE=1 SV=1 | dRNF34 |
| A1Z9E9 | 45 | S | 1.372 | 0.035897599 | Up | Uncharacterized protein OS=Drosophila melanogaster OX=7227 GN=Dmel\CG13332 PE=4 SV=1 | Dmel\CG13332 |
| A1Z9I0 | 420 | S | 5.376 | 1.25547E-05 | Up | Uncharacterized protein OS=Drosophila melanogaster OX=7227 GN=Dmel\CG6357 PE=1 SV=1 | Dmel\CG6357 |
| A1Z9I0 | 149 | S | 3.502 | 1.71781E-05 | Up | Uncharacterized protein OS=Drosophila melanogaster OX=7227 GN=Dmel\CG6357 PE=1 SV=1 | Dmel\CG6357 |
| A1Z9J3 | 3343 | S | 2.043 | 0.000196969 | Up | Short stop, isoform H OS=Drosophila melanogaster OX=7227 GN=shot PE=1 SV=1 | shot |
| A1Z9J3 | 3344 | S | 2.807 | 4.92965E-05 | Up | Short stop, isoform H OS=Drosophila melanogaster OX=7227 GN=shot PE=1 SV=1 | shot |
| A1Z9J3 | 1656 | S | 1.422 | 0.024160167 | Up | Short stop, isoform H OS=Drosophila melanogaster OX=7227 GN=shot PE=1 SV=1 | shot |
| A1Z9J3 | 3573 | S | 1.48 | 0.001786989 | Up | Short stop, isoform H OS=Drosophila melanogaster OX=7227 GN=shot PE=1 SV=1 | shot |
| A1Z9N8 | 1260 | S | 1.36 | 0.006732481 | Up | I[[h]] channel, isoform B OS=Drosophila melanogaster OX=7227 GN=Ih PE=4 SV=1 | Ih |
| A1Z9N8 | 1261 | S | 1.36 | 0.006732481 | Up | I[[h]] channel, isoform B OS=Drosophila melanogaster OX=7227 GN=Ih PE=4 SV=1 | Ih |
| A1Z9V4 | 110 | S | 1.335 | 0.01490508 | Up | Uncharacterized protein, isoform A OS=Drosophila melanogaster OX=7227 GN=CT28709 PE=4 SV=1 | CT28709 |
| A1ZA22 | 253 | S | 1.411 | 0.000860153 | Up | Eukaryotic translation initiation factor 2B subunit gamma OS=Drosophila melanogaster OX=7227 GN=eIF2Bgamma PE=1 SV=1 | eIF2Bgamma |
| A1ZA23 | 85 | S | 1.391 | 0.034244086 | Up | FI18007p1 OS=Drosophila melanogaster OX=7227 GN=Dmel\CG8192 PE=2 SV=1 | Dmel\CG8192 |
| A1ZA45 | 1087 | S | 1.466 | 0.003570379 | Up | Uncharacterized protein, isoform A OS=Drosophila melanogaster OX=7227 GN=CG12967 PE=4 SV=1 | CG12967 |
| A1ZA87 | 98 | S | 2.069 | 0.000997295 | Up | SP2353, isoform A OS=Drosophila melanogaster OX=7227 GN=SP2353 PE=4 SV=1 | SP2353 |
| A1ZAG3 | 181 | S | 2.306 | 6.95091E-07 | Up | Uncharacterized protein OS=Drosophila melanogaster OX=7227 GN=Dmel\CG4409 PE=4 SV=2 | Dmel\CG4409 |
| A1ZAG3 | 182 | S | 1.856 | 0.00024738 | Up | Uncharacterized protein OS=Drosophila melanogaster OX=7227 GN=Dmel\CG4409 PE=4 SV=2 | Dmel\CG4409 |
| Q8IFW6 | 460 | S | 0.571 | 0.0078939 | Down | DEK protein OS=Drosophila melanogaster OX=7227 GN=Dek PE=1 SV=1 | Dek |
| Q8IFW6 | 462 | S | 0.571 | 0.0078939 | Down | DEK protein OS=Drosophila melanogaster OX=7227 GN=Dek PE=1 | Dek |

| | | | | | | | |
|---|---|---|---|---|---|---|---|
| | | | | | | SV=1 | |
| Q8IFW6 | 468 | S | 0.571 | 0.0078939 | Down | DEK protein OS=Drosophila melanogaster OX=7227 GN=Dek PE=1 SV=1 | Dek |
| B7YZI7 | 345 | S | 1.422 | 0.028089124 | Up | Uncharacterized protein, isoform G OS=Drosophila melanogaster OX=7227 GN=anon-WO0140519.237 PE=4 SV=1 | anon-WO0140519.237 |
| A1ZAW5 | 202 | S | 1.318 | 0.017382437 | Up | Methylosome subunit pICln OS=Drosophila melanogaster OX=7227 GN=icln PE=1 SV=1 | icln |
| Q0E930 | 1573 | S | 1.565 | 0.005242037 | Up | Uncharacterized protein, isoform B OS=Drosophila melanogaster OX=7227 GN=BcDNA:GH04922 PE=4 SV=1 | BcDNA:GH04922 |
| Q0E930 | 1604 | S | 2.03 | 0.000371495 | Up | Uncharacterized protein, isoform B OS=Drosophila melanogaster OX=7227 GN=BcDNA:GH04922 PE=4 SV=1 | BcDNA:GH04922 |
| Q0E930 | 1623 | S | 1.918 | 0.020650155 | Up | Uncharacterized protein, isoform B OS=Drosophila melanogaster OX=7227 GN=BcDNA:GH04922 PE=4 SV=1 | BcDNA:GH04922 |
| Q0E930 | 1637 | S | 1.918 | 0.020650155 | Up | Uncharacterized protein, isoform B OS=Drosophila melanogaster OX=7227 GN=BcDNA:GH04922 PE=4 SV=1 | BcDNA:GH04922 |
| B3DN64 | 938 | S | 1.321 | 0.001945908 | Up | Guanine nucleotide exchange factor in mesoderm, isoform G OS=Drosophila melanogaster OX=7227 GN=GEFmeso PE=2 SV=1 | GEFmeso |
| A8E6M1 | 63 | S | 1.894 | 0.004981446 | Up | GH01011p OS=Drosophila melanogaster OX=7227 GN=CG5477 PE=1 SV=1 | CG5477 |
| A8E6M1 | 65 | S | 1.894 | 0.004981446 | Up | GH01011p OS=Drosophila melanogaster OX=7227 GN=CG5477 PE=1 SV=1 | CG5477 |
| A1ZBL7 | 13 | S | 1.314 | 0.029786776 | Up | Par-1, isoform H OS=Drosophila melanogaster OX=7227 GN=par-1 PE=1 SV=1 | par-1 |
| A1ZBL7 | 14 | S | 1.314 | 0.029786776 | Up | Par-1, isoform H OS=Drosophila melanogaster OX=7227 GN=par-1 PE=1 SV=1 | par-1 |
| E1JGN0 | 853 | S | 2.534 | 0.000956734 | Up | Non-specific serine/threonine protein kinase OS=Drosophila melanogaster OX=7227 GN=par-1 PE=1 SV=1 | par-1 |
| A1ZBW6 | 611 | S | 1.559 | 0.002808497 | Up | Uncharacterized protein, isoform A OS=Drosophila melanogaster OX=7227 GN=Dmel\CG11180 PE=1 SV=1 | Dmel\CG11180 |
| A1ZBW7 | 1282 | S | 1.71 | 8.89299E-05 | Up | WASH complex subunit 2 OS=Drosophila melanogaster OX=7227 GN=CG16742 PE=1 SV=1 | CG16742 |
| A1ZBW7 | 1245 | S | 1.453 | 0.001665292 | Up | WASH complex subunit 2 OS=Drosophila melanogaster OX=7227 GN=CG16742 PE=1 SV=1 | CG16742 |
| A2RVF0 | 514 | S | 1.577 | 0.000467271 | Up | ABC transporter expressed in trachea, isoform C OS=Drosophila melanogaster OX=7227 GN=Atet PE=2 SV=1 | Atet |
| A2RVF0 | 526 | S | 1.43 | 0.001963491 | Up | ABC transporter expressed in trachea, isoform C OS=Drosophila melanogaster OX=7227 GN=Atet PE=2 SV=1 | Atet |
| A2RVF0 | 529 | S | 1.43 | 0.001963491 | Up | ABC transporter expressed in trachea, isoform C OS=Drosophila melanogaster OX=7227 GN=Atet PE=2 SV=1 | Atet |
| A4UZZ4 | 249 | S | 0.543 | 0.002436373 | Down | Alpha-1,4 glucan phosphorylase OS=Drosophila melanogaster OX=7227 GN=GlyP PE=1 SV=1 | GlyP |
| A4V0B5 | 35 | S | 2.041 | 0.000614073 | Up | Nervana 2, isoform F OS=Drosophila melanogaster OX=7227 GN=nrv2 PE=1 SV=1 | nrv2 |
| A4V134 | 356 | S | 1.416 | 0.013181105 | Up | Calcium/calmodulin-dependent protein kinase II, isoform E OS=Drosophila melanogaster OX=7227 GN=CaMKII PE=1 SV=1 | CaMKII |
| A4V1B2 | 443 | S | 1.415 | 0.03122058 | Up | Patj, isoform B OS=Drosophila melanogaster OX=7227 GN=Patj PE=1 SV=1 | Patj |
| A4V1N9 | 606 | S | 1.48 | 0.002839077 | Up | Paramyosin, isoform D OS=Drosophila melanogaster OX=7227 GN=Prm PE=1 SV=1 | Prm |
| A4V1N9 | 607 | S | 1.48 | 0.002839077 | Up | Paramyosin, isoform D OS=Drosophila melanogaster OX=7227 GN=Prm PE=1 SV=1 | Prm |
| A4V4W0 | 700 | S | 1.582 | 0.003346499 | Up | RH38069p1 OS=Drosophila melanogaster OX=7227 GN=stnA PE=1 SV=1 | stnA |
| A4V4W0 | 702 | S | 1.513 | 0.003121293 | Up | RH38069p1 OS=Drosophila melanogaster OX=7227 GN=stnA PE=1 SV=1 | stnA |
| A4V4W0 | 611 | S | 2.175 | 0.00197278 | Up | RH38069p1 OS=Drosophila melanogaster OX=7227 GN=stnA PE=1 SV=1 | stnA |
| A8DY43 | 1326 | S | 1.792 | 0.000380953 | Up | G_PROTEIN_RECEP_F3_4 domain-containing protein OS=Drosophila melanogaster OX=7227 GN=Dmel\CG43795 PE=4 SV=2 | Dmel\CG43795 |
| A8DY95 | 742 | S | 2.052 | 0.00780461 | Up | Uncharacterized protein, isoform C OS=Drosophila melanogaster OX=7227 GN=Dmel\CG11883 PE=1 SV=1 | Dmel\CG11883 |
| A8DY95 | 699 | S | 1.843 | 0.007713915 | Up | Uncharacterized protein, isoform C OS=Drosophila melanogaster OX=7227 GN=Dmel\CG11883 PE=1 SV=1 | Dmel\CG11883 |
| A8DY95 | 726 | S | 2.542 | 0.005766027 | Up | Uncharacterized protein, isoform C OS=Drosophila melanogaster OX=7227 GN=Dmel\CG11883 PE=1 SV=1 | Dmel\CG11883 |
| A8DY95 | 194 | S | 1.505 | 0.016234771 | Up | Uncharacterized protein, isoform C OS=Drosophila melanogaster OX=7227 GN=Dmel\CG11883 PE=1 SV=1 | Dmel\CG11883 |
| A8DYB0 | 1724 | S | 7.066 | 0.000318327 | Up | Midasin OS=Drosophila melanogaster OX=7227 GN=Dmel\CG13185 PE=1 SV=2 | Dmel\CG13185 |
| A8DYI9 | 613 | S | 1.842 | 0.001357454 | Up | Uncharacterized protein, isoform E OS=Drosophila melanogaster OX=7227 GN=anon-EST:fe1H12 PE=1 SV=1 | anon-EST:fe1H12 |
| A8DYW2 | 202 | S | 1.341 | 0.030591968 | Up | Uncharacterized protein, isoform B OS=Drosophila melanogaster OX=7227 GN=CG13768 PE=4 SV=1 | CG13768 |
| A8DYW5 | 638 | S | 1.399 | 0.00371373 | Up | Sodium/hydrogen exchanger OS=Drosophila melanogaster OX=7227 GN=Nhe3 PE=3 SV=3 | Nhe3 |
| A8DYW5 | 640 | S | 1.409 | 0.003090064 | Up | Sodium/hydrogen exchanger OS=Drosophila melanogaster OX=7227 GN=Nhe3 PE=3 SV=3 | Nhe3 |
| Q59DZ3 | 949 | S | 1.489 | 0.004811128 | Up | FI21209p1 OS=Drosophila melanogaster OX=7227 GN=kuz PE=1 SV=1 | kuz |
| M9PBE8 | 585 | S | 1.345 | 0.026377101 | Up | Rho-type guanine nucleotide exchange factor, isoform H OS=Drosophila melanogaster OX=7227 GN=RtGEF PE=1 SV=1 | RtGEF |
| O61380 | 1197 | S | 1.357 | 0.016617304 | Up | Eukaryotic translation initiation factor 4G1, isoform A OS=Drosophila melanogaster OX=7227 GN=eIF4G1 PE=1 SV=3 | eIF4G1 |
| O61380 | 327 | S | 1.359 | 0.011857096 | Up | Eukaryotic translation initiation factor 4G1, isoform A OS=Drosophila melanogaster OX=7227 GN=eIF4G1 PE=1 SV=3 | eIF4G1 |
| M9MRV3 | 390 | S | 1.898 | 0.003541021 | Up | Uncharacterized protein, isoform G OS=Drosophila melanogaster OX=7227 GN=CG10860 PE=4 SV=1 | CG10860 |
| M9PE05 | 1584 | S | 1.795 | 0.033185789 | Up | Encore, isoform H OS=Drosophila melanogaster OX=7227 GN=enc PE=1 SV=1 | enc |
| A8JNM0 | 442 | S | 1.355 | 0.029204908 | Up | Target of wingless, isoform C OS=Drosophila melanogaster OX=7227 GN=tow PE=4 SV=1 | tow |
| A8JNN5 | 148 | S | 1.494 | 3.50213E-05 | Up | GTPase activating protein and centrosome-associated, isoform B OS=Drosophila melanogaster OX=7227 GN=GAPcenA PE=1 SV=1 | GAPcenA |
| A8JQT5 | 4072 | S | 1.462 | 0.026650268 | Up | Chondrocyte-derived ezrin-like domain containing protein, isoform G OS=Drosophila melanogaster OX=7227 GN=Cdep PE=1 SV=2 | Cdep |
| A8JQT5 | 2720 | S | 0.72 | 0.010494497 | Down | Chondrocyte-derived ezrin-like domain containing protein, isoform G OS=Drosophila melanogaster OX=7227 GN=Cdep PE=1 SV=2 | Cdep |
| A8JQT5 | 2604 | S | 0.722 | 0.022081234 | Down | Chondrocyte-derived ezrin-like domain containing protein, isoform G OS=Drosophila melanogaster OX=7227 GN=Cdep PE=1 SV=2 | Cdep |
| A8JQV2 | 770 | S | 0.7 | 0.010964752 | Down | Uncharacterized protein, isoform E OS=Drosophila melanogaster OX=7227 GN=Dmel\CG17816 PE=1 SV=1 | Dmel\CG17816 |
| A8JR00 | 352 | S | 1.454 | 0.011570574 | Up | Uncharacterized protein, isoform E OS=Drosophila melanogaster OX=7227 GN=kmr PE=4 SV=1 | kmr |
| A8JR00 | 355 | S | 1.52 | 0.010850549 | Up | Uncharacterized protein, isoform E OS=Drosophila melanogaster OX=7227 GN=kmr PE=4 SV=1 | kmr |
| A8JR05 | 734 | S | 1.686 | 0.000208891 | Up | Crossveinless c, isoform C OS=Drosophila melanogaster OX=7227 GN=cv-c PE=4 SV=1 | cv-c |
| Q9VDH1 | 35 | S | 1.89 | 0.006431912 | Up | Uncharacterized protein, isoform A OS=Drosophila melanogaster OX=7227 GN=CG17276 PE=1 SV=2 | CG17276 |
| A8JTM7 | 4738 | S | 3.172 | 2.4463E-05 | Up | Megalin, isoform A OS=Drosophila melanogaster OX=7227 GN=mgl PE=1 SV=2 | mgl |
| A8JTM7 | 4767 | S | 1.995 | 0.003895929 | Up | Megalin, isoform A OS=Drosophila melanogaster OX=7227 GN=mgl PE=1 SV=2 | mgl |
| A8JTM7 | 4628 | S | 1.738 | 0.000210255 | Up | Megalin, isoform A OS=Drosophila melanogaster OX=7227 GN=mgl PE=1 SV=2 | mgl |
| A8JUP3 | 1302 | S | 1.521 | 0.040954974 | Up | Chitinase 6, isoform H OS=Drosophila melanogaster OX=7227 GN=Cht6 PE=1 SV=2 | Cht6 |

| | | | | | | | |
|---|---|---|---|---|---|---|---|
| A8JUZ7 | 301 | S | 1.367 | 0.007949107 | Up | Uncharacterized protein, isoform B OS=Drosophila melanogaster OX=7227 GN=Dmel\CG15894 PE=1 SV=1 | Dmel\CG15894 |
| A8JUZ7 | 1409 | S | 1.428 | 0.044080634 | Up | Uncharacterized protein, isoform B OS=Drosophila melanogaster OX=7227 GN=Dmel\CG15894 PE=1 SV=1 | Dmel\CG15894 |
| A8JUZ7 | 572 | S | 1.649 | 0.014117714 | Up | Uncharacterized protein, isoform B OS=Drosophila melanogaster OX=7227 GN=Dmel\CG15894 PE=1 SV=1 | Dmel\CG15894 |
| A8JUZ7 | 573 | S | 1.741 | 0.017115021 | Up | Uncharacterized protein, isoform B OS=Drosophila melanogaster OX=7227 GN=Dmel\CG15894 PE=1 SV=1 | Dmel\CG15894 |
| A8JUZ7 | 1329 | S | 1.551 | 0.017399848 | Up | Uncharacterized protein, isoform B OS=Drosophila melanogaster OX=7227 GN=Dmel\CG15894 PE=1 SV=1 | Dmel\CG15894 |
| A8JUZ7 | 1037 | S | 1.347 | 0.029762368 | Up | Uncharacterized protein, isoform B OS=Drosophila melanogaster OX=7227 GN=Dmel\CG15894 PE=1 SV=1 | Dmel\CG15894 |
| Q9W3W1 | 575 | S | 3.27 | 0.001036458 | Up | Coronin OS=Drosophila melanogaster OX=7227 GN=pod1 PE=1 SV=1 | pod1 |
| Q9W3W1 | 534 | S | 2.817 | 0.001730158 | Up | Coronin OS=Drosophila melanogaster OX=7227 GN=pod1 PE=1 SV=1 | pod1 |
| Q9W3W1 | 542 | S | 1.644 | 0.000685799 | Up | Coronin OS=Drosophila melanogaster OX=7227 GN=pod1 PE=1 SV=1 | pod1 |
| Q9W3W1 | 431 | S | 1.674 | 0.001425597 | Up | Coronin OS=Drosophila melanogaster OX=7227 GN=pod1 PE=1 SV=1 | pod1 |
| A8JV25 | 23 | S | 2.295 | 9.30947E-06 | Up | Uncharacterized protein, isoform D OS=Drosophila melanogaster OX=7227 GN=Dmel\CG4829 PE=1 SV=1 | Dmel\CG4829 |
| B5RIV2 | 16 | S | 1.563 | 0.0118952 | Up | FI05478p OS=Drosophila melanogaster OX=7227 GN=CG9801-RA PE=2 SV=1 | CG9801-RA |
| B5RIV2 | 158 | S | 1.329 | 0.010834204 | Up | FI05478p OS=Drosophila melanogaster OX=7227 GN=CG9801-RA PE=2 SV=1 | CG9801-RA |
| B5RIV2 | 29 | S | 1.416 | 0.014852244 | Up | FI05478p OS=Drosophila melanogaster OX=7227 GN=CG9801-RA PE=2 SV=1 | CG9801-RA |
| B5RIZ2 | 593 | S | 1.592 | 0.003820898 | Up | FI06314p OS=Drosophila melanogaster OX=7227 GN=RhoGAP92B PE=1 SV=1 | RhoGAP92B |
| B5RJS0 | 526 | S | 0.727 | 0.004263203 | Down | IP20241p OS=Drosophila melanogaster OX=7227 GN=Nost PE=1 SV=1 | Nost |
| B5X0J4 | 32 | S | 2.817 | 0.000352209 | Up | FI04429p OS=Drosophila melanogaster OX=7227 GN=ple PE=1 SV=1 | ple |
| B7YZL6 | 99 | S | 1.417 | 0.010728155 | Up | Uncharacterized protein, isoform R OS=Drosophila melanogaster OX=7227 GN=Dmel\CG10737 PE=1 SV=1 | Dmel\CG10737 |
| B7YZQ1 | 482 | S | 1.523 | 0.013672711 | Up | Sterile20-like kinase, isoform F OS=Drosophila melanogaster OX=7227 GN=Slik PE=1 SV=1 | Slik |
| B7YZQ1 | 1376 | S | 1.537 | 0.01728036 | Up | Sterile20-like kinase, isoform F OS=Drosophila melanogaster OX=7227 GN=Slik PE=1 SV=1 | Slik |
| B7YZQ3 | 1067 | S | 0.648 | 0.012002889 | Down | Prominin, isoform D OS=Drosophila melanogaster OX=7227 GN=prom PE=1 SV=1 | prom |
| B7YZQ3 | 1068 | S | 0.566 | 0.003441366 | Down | Prominin, isoform D OS=Drosophila melanogaster OX=7227 GN=prom PE=1 SV=1 | prom |
| B7YZQ3 | 1070 | S | 0.564 | 0.000915643 | Down | Prominin, isoform D OS=Drosophila melanogaster OX=7227 GN=prom PE=1 SV=1 | prom |
| B7YZQ3 | 969 | S | 1.617 | 0.031473451 | Up | Prominin, isoform D OS=Drosophila melanogaster OX=7227 GN=prom PE=1 SV=1 | prom |
| Q9VJJ8 | 1239 | S | 1.537 | 0.00718789 | Up | LD30602p OS=Drosophila melanogaster OX=7227 GN=mtgo PE=1 SV=2 | mtgo |
| B7YZY0 | 892 | S | 1.309 | 0.000152424 | Up | Uncharacterized protein, isoform C OS=Drosophila melanogaster OX=7227 GN=CG17465 PE=1 SV=1 | CG17465 |
| B7YZY0 | 1103 | S | 1.575 | 0.023320944 | Up | Uncharacterized protein, isoform C OS=Drosophila melanogaster OX=7227 GN=CG17465 PE=1 SV=1 | CG17465 |
| Q9VQL7 | 2141 | S | 0.706 | 0.000717686 | Down | Fatty acid synthase 1, isoform A OS=Drosophila melanogaster OX=7227 GN=FASN1 PE=1 SV=1 | FASN1 |
| Q9VQL7 | 2144 | S | 0.64 | 0.000184223 | Down | Fatty acid synthase 1, isoform A OS=Drosophila melanogaster OX=7227 GN=FASN1 PE=1 SV=1 | FASN1 |
| Q9VQL7 | 2145 | S | 0.645 | 0.000258169 | Down | Fatty acid synthase 1, isoform A OS=Drosophila melanogaster OX=7227 GN=FASN1 PE=1 SV=1 | FASN1 |
| Q9VQL7 | 1676 | S | 0.473 | 0.000460421 | Down | Fatty acid synthase 1, isoform A OS=Drosophila melanogaster OX=7227 GN=FASN1 PE=1 SV=1 | FASN1 |
| Q9VQL7 | 2081 | S | 0.697 | 0.000379235 | Down | Fatty acid synthase 1, isoform A OS=Drosophila melanogaster OX=7227 GN=FASN1 PE=1 SV=1 | FASN1 |
| Q9VQL7 | 2431 | S | 1.313 | 0.000715068 | Up | Fatty acid synthase 1, isoform A OS=Drosophila melanogaster OX=7227 GN=FASN1 PE=1 SV=1 | FASN1 |
| M9NEL3 | 1545 | S | 1.685 | 0.014115559 | Up | Kismet, isoform F OS=Drosophila melanogaster OX=7227 GN=kis PE=1 SV=1 | kis |
| M9NEL3 | 1601 | S | 1.812 | 0.005733811 | Up | Kismet, isoform F OS=Drosophila melanogaster OX=7227 GN=kis PE=1 SV=1 | kis |
| B7Z010 | 67 | S | 1.956 | 0.005505756 | Up | Cappuccino, isoform E OS=Drosophila melanogaster OX=7227 GN=capu PE=4 SV=1 | capu |
| B7Z010 | 381 | S | 1.418 | 0.002628878 | Up | Cappuccino, isoform E OS=Drosophila melanogaster OX=7227 GN=capu PE=4 SV=1 | capu |
| B7Z061 | 216 | S | 0.701 | 0.00353176 | Down | Photoreceptor dehydrogenase, isoform D OS=Drosophila melanogaster OX=7227 GN=Pdh PE=1 SV=1 | Pdh |
| B7Z0A9 | 179 | S | 1.536 | 0.001121516 | Up | Sosondowah, isoform H OS=Drosophila melanogaster OX=7227 GN=sowah PE=1 SV=2 | sowah |
| B7Z0A9 | 183 | S | 1.548 | 0.001612891 | Up | Sosondowah, isoform H OS=Drosophila melanogaster OX=7227 GN=sowah PE=1 SV=2 | sowah |
| Q9VBK2 | 153 | S | 1.447 | 0.002749416 | Up | RE27406p OS=Drosophila melanogaster OX=7227 GN=CG8957 PE=2 SV=2 | CG8957 |
| B7Z0T3 | 500 | S | 0.724 | 0.009144274 | Down | Mustard, isoform S OS=Drosophila melanogaster OX=7227 GN=mtd PE=1 SV=1 | mtd |
| B7Z0T3 | 485 | S | 2.282 | 0.004818737 | Up | Mustard, isoform S OS=Drosophila melanogaster OX=7227 GN=mtd PE=1 SV=1 | mtd |
| Q9V3J6 | 610 | S | 2.014 | 0.000200322 | Up | Bestrophin homolog OS=Drosophila melanogaster OX=7227 GN=Best1 PE=2 SV=1 | Best1 |
| Q9V3J6 | 616 | S | 2.014 | 0.000200322 | Up | Bestrophin homolog OS=Drosophila melanogaster OX=7227 GN=Best1 PE=2 SV=1 | Best1 |
| B7Z0W8 | 1363 | S | 0.685 | 0.011880433 | Down | Rugose, isoform M OS=Drosophila melanogaster OX=7227 GN=rg PE=1 SV=2 | rg |
| B7Z0W9 | 1560 | S | 1.367 | 0.020533711 | Up | Proton channel OtopLc OS=Drosophila melanogaster OX=7227 GN=OtopLc PE=3 SV=2 | OtopLc |
| B7Z0W9 | 522 | S | 0.751 | 0.007233426 | Down | Proton channel OtopLc OS=Drosophila melanogaster OX=7227 GN=OtopLc PE=3 SV=2 | OtopLc |
| X2JFE8 | 1173 | S | 0.693 | 0.018854314 | Down | Sodium channel protein OS=Drosophila melanogaster OX=7227 GN=para PE=3 SV=1 | para |
| B7Z122 | 12 | S | 1.666 | 0.004375505 | Up | Transporter OS=Drosophila melanogaster OX=7227 GN=Dmel\CG10804 PE=3 SV=1 | Dmel\CG10804 |
| B7Z122 | 13 | S | 1.652 | 0.002239093 | Up | Transporter OS=Drosophila melanogaster OX=7227 GN=Dmel\CG10804 PE=3 SV=1 | Dmel\CG10804 |
| B7Z122 | 23 | S | 1.501 | 0.0172405 | Up | Transporter OS=Drosophila melanogaster OX=7227 GN=Dmel\CG10804 PE=3 SV=1 | Dmel\CG10804 |
| X2JIR4 | 112 | S | 1.727 | 0.007021209 | Up | C3G guanyl-nucleotide exchange factor, isoform L OS=Drosophila melanogaster OX=7227 GN=C3G PE=4 SV=1 | C3G |
| X2JIR4 | 333 | S | 1.323 | 0.000774143 | Up | C3G guanyl-nucleotide exchange factor, isoform L OS=Drosophila melanogaster OX=7227 GN=C3G PE=4 SV=1 | C3G |
| B7Z145 | 54 | S | 1.393 | 0.044116855 | Up | Tenascin accessory, isoform E OS=Drosophila melanogaster OX=7227 GN=Ten-a PE=4 SV=1 | Ten-a |
| C0HK95 | 44 | S | 1.461 | 0.044289836 | Up | Protein anoxia up-regulated OS=Drosophila melanogaster OX=7227 GN=fau PE=2 SV=1 | fau |
| C0HK95 | 94 | S | 1.45 | 0.040260443 | Up | Protein anoxia up-regulated OS=Drosophila melanogaster OX=7227 GN=fau PE=2 SV=1 | fau |
| C0MJA0 | 103 | S | 0.636 | 0.004356148 | Down | CG17950-PA OS=Drosophila melanogaster OX=7227 GN=HmgD PE=1 SV=1 | HmgD |
| C7LA93 | 870 | S | 1.789 | 0.000300247 | Up | Dorsal, isoform F OS=Drosophila melanogaster OX=7227 GN=dl PE=1 SV=1 | dl |
| C7LA93 | 873 | S | 1.514 | 0.00207693 | Up | Dorsal, isoform F OS=Drosophila melanogaster OX=7227 GN=dl PE=1 SV=1 | dl |
| C7LA93 | 770 | S | 1.621 | 0.004499842 | Up | Dorsal, isoform F OS=Drosophila melanogaster OX=7227 GN=dl PE=1 SV=1 | dl |

| | | | | | | | |
|---|---|---|---|---|---|---|---|
| C8VV14 | 36 | S | 0.634 | 0.001099458 | Down | Fructose-bisphosphate aldolase OS=Drosophila melanogaster OX=7227 GN=Ald1 PE=1 SV=1 | Ald1 |
| L0MN91 | 1001 | S | 1.818 | 0.001818625 | Up | Bent, isoform I OS=Drosophila melanogaster OX=7227 GN=bt PE=1 SV=1 | bt |
| L0MN91 | 1360 | S | 1.322 | 0.003654085 | Up | Bent, isoform I OS=Drosophila melanogaster OX=7227 GN=bt PE=1 SV=1 | bt |
| L0MN91 | 7066 | S | 0.593 | 0.000100694 | Down | Bent, isoform I OS=Drosophila melanogaster OX=7227 GN=bt PE=1 SV=1 | bt |
| L0MN91 | 7069 | S | 0.593 | 0.000100694 | Down | Bent, isoform I OS=Drosophila melanogaster OX=7227 GN=bt PE=1 SV=1 | bt |
| L0MN91 | 7070 | S | 0.593 | 0.000100694 | Down | Bent, isoform I OS=Drosophila melanogaster OX=7227 GN=bt PE=1 SV=1 | bt |
| L0MN91 | 1364 | S | 1.852 | 0.016092332 | Up | Bent, isoform I OS=Drosophila melanogaster OX=7227 GN=bt PE=1 SV=1 | bt |
| L0MN91 | 4737 | S | 0.721 | 0.019475162 | Down | Bent, isoform I OS=Drosophila melanogaster OX=7227 GN=bt PE=1 SV=1 | bt |
| L0MN91 | 6588 | S | 0.699 | 0.017865576 | Down | Bent, isoform I OS=Drosophila melanogaster OX=7227 GN=bt PE=1 SV=1 | bt |
| D1YSG7 | 356 | S | 1.416 | 0.013181105 | Up | Calcium/calmodulin-dependent protein kinase II, isoform I OS=Drosophila melanogaster OX=7227 GN=CaMKII PE=1 SV=1 | CaMKII |
| E1JGN0 | 235 | S | 1.446 | 0.002128494 | Up | Non-specific serine/threonine protein kinase OS=Drosophila melanogaster OX=7227 GN=par-1 PE=1 SV=1 | par-1 |
| Q6NNF2 | 28 | S | 0.768 | 0.00222178 | Down | High affinity cAMP-specific and IBMX-insensitive 3',5'-cyclic phosphodiesterase 8 OS=Drosophila melanogaster OX=7227 GN=Pde8 PE=2 SV=1 | Pde8 |
| Q6NNF2 | 138 | S | 1.324 | 0.000838687 | Up | High affinity cAMP-specific and IBMX-insensitive 3',5'-cyclic phosphodiesterase 8 OS=Drosophila melanogaster OX=7227 GN=Pde8 PE=2 SV=1 | Pde8 |
| Q6NNF2 | 139 | S | 1.324 | 0.000838687 | Up | High affinity cAMP-specific and IBMX-insensitive 3',5'-cyclic phosphodiesterase 8 OS=Drosophila melanogaster OX=7227 GN=Pde8 PE=2 SV=1 | Pde8 |
| Q6NNF2 | 548 | S | 1.608 | 0.014126591 | Up | High affinity cAMP-specific and IBMX-insensitive 3',5'-cyclic phosphodiesterase 8 OS=Drosophila melanogaster OX=7227 GN=Pde8 PE=2 SV=1 | Pde8 |
| Q9VM55 | 3523 | S | 1.661 | 0.018465955 | Up | Uninflatable, isoform B OS=Drosophila melanogaster OX=7227 GN=uif PE=1 SV=3 | uif |
| Q9VM55 | 3526 | S | 1.661 | 0.018465955 | Up | Uninflatable, isoform B OS=Drosophila melanogaster OX=7227 GN=uif PE=1 SV=3 | uif |
| M9PCG4 | 81 | S | 1.415 | 0.001171395 | Up | Uncharacterized protein, isoform H OS=Drosophila melanogaster OX=7227 GN=CG9585 PE=4 SV=1 | CG9585 |
| E1JHJ2 | 205 | S | 1.783 | 0.006333174 | Up | Uncharacterized protein, isoform C OS=Drosophila melanogaster OX=7227 GN=Dmel\CG5953 PE=1 SV=1 | Dmel\CG5953 |
| M9NF46 | 742 | S | 1.984 | 0.007575897 | Up | Myosin heavy chain, isoform R OS=Drosophila melanogaster OX=7227 GN=Mhc PE=1 SV=1 | Mhc |
| M9NF46 | 1280 | S | 0.696 | 0.016550438 | Down | Myosin heavy chain, isoform R OS=Drosophila melanogaster OX=7227 GN=Mhc PE=1 SV=1 | Mhc |
| E1JHM0 | 1215 | S | 1.609 | 0.005476436 | Up | Cd GTPase activating protein-related, isoform C OS=Drosophila melanogaster OX=7227 GN=CdGAPr PE=1 SV=1 | CdGAPr |
| E1JHM0 | 1750 | S | 1.502 | 0.002600045 | Up | Cd GTPase activating protein-related, isoform C OS=Drosophila melanogaster OX=7227 GN=CdGAPr PE=1 SV=1 | CdGAPr |
| E1JHM0 | 1570 | S | 3.606 | 0.000351336 | Up | Cd GTPase activating protein-related, isoform C OS=Drosophila melanogaster OX=7227 GN=CdGAPr PE=1 SV=1 | CdGAPr |
| E1JHM0 | 407 | S | 1.482 | 0.00121565 | Up | Cd GTPase activating protein-related, isoform C OS=Drosophila melanogaster OX=7227 GN=CdGAPr PE=1 SV=1 | CdGAPr |
| E1JHM0 | 813 | S | 1.822 | 0.007247896 | Up | Cd GTPase activating protein-related, isoform C OS=Drosophila melanogaster OX=7227 GN=CdGAPr PE=1 SV=1 | CdGAPr |
| E1JHM0 | 837 | S | 1.432 | 0.001350489 | Up | Cd GTPase activating protein-related, isoform C OS=Drosophila melanogaster OX=7227 GN=CdGAPr PE=1 SV=1 | CdGAPr |
| E1JHM0 | 840 | S | 1.432 | 0.001350489 | Up | Cd GTPase activating protein-related, isoform C OS=Drosophila melanogaster OX=7227 GN=CdGAPr PE=1 SV=1 | CdGAPr |
| Q9V9Q9 | 175 | S | 1.328 | 0.013144124 | Up | GH19218p OS=Drosophila melanogaster OX=7227 GN=nolo PE=1 SV=2 | nolo |
| Q9VMV9 | 263 | S | 1.599 | 0.001335769 | Up | Reticulon-like protein OS=Drosophila melanogaster OX=7227 GN=Rtnl1 PE=1 SV=2 | Rtnl1 |
| Q9VMV9 | 269 | S | 1.7 | 0.002296965 | Up | Reticulon-like protein OS=Drosophila melanogaster OX=7227 GN=Rtnl1 PE=1 SV=2 | Rtnl1 |
| M9PE12 | 2336 | S | 1.465 | 0.002465988 | Up | Enhancer of bithorax, isoform I OS=Drosophila melanogaster OX=7227 GN=E(bx) PE=1 SV=1 | E(bx) |
| M9NDT7 | 1159 | S | 1.316 | 0.009800195 | Up | Disabled, isoform C OS=Drosophila melanogaster OX=7227 GN=Dab PE=4 SV=1 | Dab |
| M9NDT7 | 1949 | S | 1.415 | 0.001421256 | Up | Disabled, isoform C OS=Drosophila melanogaster OX=7227 GN=Dab PE=4 SV=1 | Dab |
| M9NDT7 | 1595 | S | 1.347 | 0.000721614 | Up | Disabled, isoform C OS=Drosophila melanogaster OX=7227 GN=Dab PE=4 SV=1 | Dab |
| M9NDT7 | 727 | S | 1.328 | 0.028335918 | Up | Disabled, isoform C OS=Drosophila melanogaster OX=7227 GN=Dab PE=4 SV=1 | Dab |
| M9NDT7 | 1658 | S | 1.372 | 0.008962915 | Up | Disabled, isoform C OS=Drosophila melanogaster OX=7227 GN=Dab PE=4 SV=1 | Dab |
| M9NDT7 | 1539 | S | 1.334 | 0.007184161 | Up | Disabled, isoform C OS=Drosophila melanogaster OX=7227 GN=Dab PE=4 SV=1 | Dab |
| E1JI22 | 203 | S | 1.364 | 0.046890629 | Up | Clathrin light chain OS=Drosophila melanogaster OX=7227 GN=Clc PE=1 SV=1 | Clc |
| M9PIA6 | 275 | S | 0.568 | 0.027279898 | Down | Mi-2, isoform D OS=Drosophila melanogaster OX=7227 GN=Mi-2 PE=1 SV=1 | Mi-2 |
| M9PIA6 | 276 | S | 0.568 | 0.027279898 | Down | Mi-2, isoform D OS=Drosophila melanogaster OX=7227 GN=Mi-2 PE=1 SV=1 | Mi-2 |
| M9PIA6 | 201 | S | 2.03 | 0.001090031 | Up | Mi-2, isoform D OS=Drosophila melanogaster OX=7227 GN=Mi-2 PE=1 SV=1 | Mi-2 |
| M9PIA6 | 212 | S | 1.319 | 0.006761732 | Up | Mi-2, isoform D OS=Drosophila melanogaster OX=7227 GN=Mi-2 PE=1 SV=1 | Mi-2 |
| E1JIT7 | 1152 | S | 1.85 | 0.001066046 | Up | PH and SEC7 domain-containing protein OS=Drosophila melanogaster OX=7227 GN=Efa6 PE=1 SV=2 | Efa6 |
| E1JIT7 | 686 | S | 1.437 | 0.023928584 | Up | PH and SEC7 domain-containing protein OS=Drosophila melanogaster OX=7227 GN=Efa6 PE=1 SV=2 | Efa6 |
| Q59E07 | 16 | S | 1.35 | 0.000159911 | Up | Uncharacterized protein, isoform C OS=Drosophila melanogaster OX=7227 GN=BcDNA:GH02439 PE=1 SV=1 | BcDNA:GH02439 |
| Q59E07 | 320 | S | 1.64 | 0.000391315 | Up | Uncharacterized protein, isoform C OS=Drosophila melanogaster OX=7227 GN=BcDNA:GH02439 PE=1 SV=1 | BcDNA:GH02439 |
| Q59E07 | 321 | S | 1.678 | 0.000483249 | Up | Uncharacterized protein, isoform C OS=Drosophila melanogaster OX=7227 GN=BcDNA:GH02439 PE=1 SV=1 | BcDNA:GH02439 |
| Q59E07 | 323 | S | 1.374 | 0.001288844 | Up | Uncharacterized protein, isoform C OS=Drosophila melanogaster OX=7227 GN=BcDNA:GH02439 PE=1 SV=1 | BcDNA:GH02439 |
| Q59E07 | 330 | S | 1.364 | 0.009601851 | Up | Uncharacterized protein, isoform C OS=Drosophila melanogaster OX=7227 GN=BcDNA:GH02439 PE=1 SV=1 | BcDNA:GH02439 |
| Q59E07 | 338 | S | 1.515 | 0.000548137 | Up | Uncharacterized protein, isoform C OS=Drosophila melanogaster OX=7227 GN=BcDNA:GH02439 PE=1 SV=1 | BcDNA:GH02439 |
| Q59E07 | 351 | S | 2.594 | 0.002608993 | Up | Uncharacterized protein, isoform C OS=Drosophila melanogaster OX=7227 GN=BcDNA:GH02439 PE=1 SV=1 | BcDNA:GH02439 |
| E1JJ98 | 736 | S | 1.37 | 0.011589333 | Up | FI20257p1 OS=Drosophila melanogaster OX=7227 GN=sol PE=2 SV=1 | sol |
| E1JJA5 | 788 | S | 1.407 | 0.049829665 | Up | Shibire, isoform M OS=Drosophila melanogaster OX=7227 GN=shi PE=1 SV=1 | shi |
| Q9W572 | 568 | S | 1.489 | 0.032398892 | Up | Broad, isoform C OS=Drosophila melanogaster OX=7227 GN=br PE=4 SV=3 | br |
| M9PGC1 | 1177 | S | 1.627 | 0.020898948 | Up | Protostome-specific GEF, isoform G OS=Drosophila melanogaster OX=7227 GN=PsGEF PE=4 SV=1 | PsGEF |
| M9PGC1 | 1180 | S | 1.627 | 0.020898948 | Up | Protostome-specific GEF, isoform G OS=Drosophila melanogaster OX=7227 GN=PsGEF PE=4 SV=1 | PsGEF |

| | | | | | | | |
|---|---|---|---|---|---|---|---|
| Q9W4A6 | 340 | S | 0.63 | 0.000582988 | Down | Otopetrin-like a, isoform A OS=Drosophila melanogaster OX=7227 GN=OtopLa PE=1 SV=1 | OtopLa |
| E1JJK8 | 375 | S | 1.362 | 0.010046179 | Up | Retinal degeneration B, isoform G OS=Drosophila melanogaster OX=7227 GN=rdgB PE=1 SV=1 | rdgB |
| E1JJK8 | 379 | S | 1.362 | 0.010046179 | Up | Retinal degeneration B, isoform G OS=Drosophila melanogaster OX=7227 GN=rdgB PE=1 SV=1 | rdgB |
| E1JJK8 | 382 | S | 1.362 | 0.010046179 | Up | Retinal degeneration B, isoform G OS=Drosophila melanogaster OX=7227 GN=rdgB PE=1 SV=1 | rdgB |
| E1JJK8 | 838 | S | 1.848 | 0.003891808 | Up | Retinal degeneration B, isoform G OS=Drosophila melanogaster OX=7227 GN=rdgB PE=1 SV=1 | rdgB |
| M9PJM8 | 358 | S | 1.707 | 0.042887852 | Up | Retinal degeneration B, isoform I OS=Drosophila melanogaster OX=7227 GN=rdgB PE=1 SV=1 | rdgB |
| Q9VX48 | 1260 | S | 1.458 | 0.002210303 | Up | SD02996p OS=Drosophila melanogaster OX=7227 GN=CG12432 PE=1 SV=2 | CG12432 |
| X2JFY0 | 9 | S | 1.353 | 0.007266646 | Up | N-acetylglucosamine-6-phosphate deacetylase OS=Drosophila melanogaster OX=7227 GN=Dmel\CG17065 PE=3 SV=1 | Dmel\CG17065 |
| E1JJS2 | 744 | S | 1.415 | 0.015670714 | Up | Rho GTPase activating protein at 19D, isoform B OS=Drosophila melanogaster OX=7227 GN=RhoGAP19D PE=1 SV=1 | RhoGAP19D |
| E2QCY4 | 1889 | S | 0.716 | 0.020130042 | Down | Furry, isoform C OS=Drosophila melanogaster OX=7227 GN=fry PE=1 SV=1 | fry |
| E2QCY9 | 952 | S | 1.685 | 0.000996841 | Up | Synapsin, isoform D OS=Drosophila melanogaster OX=7227 GN=Syn PE=1 SV=1 | Syn |
| E2QCY9 | 954 | S | 1.939 | 0.001728733 | Up | Synapsin, isoform D OS=Drosophila melanogaster OX=7227 GN=Syn PE=1 SV=1 | Syn |
| E2QCY9 | 574 | S | 2.674 | 0.006272268 | Up | Synapsin, isoform D OS=Drosophila melanogaster OX=7227 GN=Syn PE=1 SV=1 | Syn |
| E2QCY9 | 549 | S | 1.664 | 0.024596649 | Up | Synapsin, isoform D OS=Drosophila melanogaster OX=7227 GN=Syn PE=1 SV=1 | Syn |
| E2QCY9 | 22 | S | 1.593 | 0.00155476 | Up | Synapsin, isoform D OS=Drosophila melanogaster OX=7227 GN=Syn PE=1 SV=1 | Syn |
| E2QCY9 | 99 | S | 2.436 | 0.014143545 | Up | Synapsin, isoform D OS=Drosophila melanogaster OX=7227 GN=Syn PE=1 SV=1 | Syn |
| E2QCY9 | 925 | S | 1.382 | 0.048971324 | Up | Synapsin, isoform D OS=Drosophila melanogaster OX=7227 GN=Syn PE=1 SV=1 | Syn |
| E2QCY9 | 475 | S | 1.354 | 0.012450894 | Up | Synapsin, isoform D OS=Drosophila melanogaster OX=7227 GN=Syn PE=1 SV=1 | Syn |
| E2QCY9 | 476 | S | 1.522 | 0.005302582 | Up | Synapsin, isoform D OS=Drosophila melanogaster OX=7227 GN=Syn PE=1 SV=1 | Syn |
| E2QCY9 | 522 | S | 1.618 | 0.002608601 | Up | Synapsin, isoform D OS=Drosophila melanogaster OX=7227 GN=Syn PE=1 SV=1 | Syn |
| E2QCY9 | 523 | S | 1.667 | 0.001745282 | Up | Synapsin, isoform D OS=Drosophila melanogaster OX=7227 GN=Syn PE=1 SV=1 | Syn |
| E2QCY9 | 526 | S | 1.618 | 0.002608601 | Up | Synapsin, isoform D OS=Drosophila melanogaster OX=7227 GN=Syn PE=1 SV=1 | Syn |
| E2QCY9 | 783 | S | 1.68 | 0.002235301 | Up | Synapsin, isoform D OS=Drosophila melanogaster OX=7227 GN=Syn PE=1 SV=1 | Syn |
| E2QCY9 | 789 | S | 1.397 | 0.021359425 | Up | Synapsin, isoform D OS=Drosophila melanogaster OX=7227 GN=Syn PE=1 SV=1 | Syn |
| E2QCY9 | 641 | S | 3.743 | 0.018755498 | Up | Synapsin, isoform D OS=Drosophila melanogaster OX=7227 GN=Syn PE=1 SV=1 | Syn |
| E2QCY9 | 919 | S | 1.778 | 0.007912687 | Up | Synapsin, isoform D OS=Drosophila melanogaster OX=7227 GN=Syn PE=1 SV=1 | Syn |
| E2QCY9 | 922 | S | 1.515 | 0.016034702 | Up | Synapsin, isoform D OS=Drosophila melanogaster OX=7227 GN=Syn PE=1 SV=1 | Syn |
| E2QCY9 | 979 | S | 1.972 | 0.002942412 | Up | Synapsin, isoform D OS=Drosophila melanogaster OX=7227 GN=Syn PE=1 SV=1 | Syn |
| E2QCZ8 | 681 | S | 1.85 | 0.034196502 | Up | Smallish, isoform F OS=Drosophila melanogaster OX=7227 GN=smash PE=1 SV=1 | smash |
| F0JAH7 | 384 | S | 1.597 | 0.037731841 | Up | Hoepel1, isoform H OS=Drosophila melanogaster OX=7227 GN=hoe1 PE=1 SV=1 | hoe1 |
| F0JAN1 | 111 | S | 0.576 | 0.00026097 | Down | PAICS bifunctional enzyme, isoform B OS=Drosophila melanogaster OX=7227 GN=Paics PE=1 SV=1 | Paics |
| F6J1D0 | 22 | S | 2.276 | 0.004499857 | Up | CG3595 OS=Drosophila melanogaster OX=7227 GN=sqh PE=1 SV=1 | sqh |
| G3M3A2 | 241 | S | 1.314 | 0.022605346 | Up | Ribosomal protein S3, isoform B OS=Drosophila melanogaster OX=7227 GN=RpS3 PE=1 SV=1 | RpS3 |
| H1UUB1 | 68 | S | 1.758 | 0.000107004 | Up | GEO12465p1 OS=Drosophila melanogaster OX=7227 GN=Syb PE=1 SV=1 | Syb |
| H1UUB1 | 54 | S | 1.72 | 0.000204282 | Up | GEO12465p1 OS=Drosophila melanogaster OX=7227 GN=Syb PE=1 SV=1 | Syb |
| M9NDX8 | 669 | S | 1.317 | 0.001441983 | Up | Spinophilin, isoform I OS=Drosophila melanogaster OX=7227 GN=Spn PE=1 SV=1 | Spn |
| M9NDX8 | 673 | S | 1.317 | 0.001441983 | Up | Spinophilin, isoform I OS=Drosophila melanogaster OX=7227 GN=Spn PE=1 SV=1 | Spn |
| M9NDX8 | 678 | S | 1.317 | 0.001441983 | Up | Spinophilin, isoform I OS=Drosophila melanogaster OX=7227 GN=Spn PE=1 SV=1 | Spn |
| H9XVM5 | 338 | S | 1.675 | 0.004314716 | Up | Uncharacterized protein, isoform B OS=Drosophila melanogaster OX=7227 GN=CG11572 PE=4 SV=1 | CG11572 |
| H9XVP2 | 1445 | S | 1.36 | 0.032331187 | Up | Unc-13, isoform E OS=Drosophila melanogaster OX=7227 GN=unc-13 PE=4 SV=1 | unc-13 |
| H9XVP2 | 1227 | S | 1.378 | 0.047147067 | Up | Unc-13, isoform E OS=Drosophila melanogaster OX=7227 GN=unc-13 PE=4 SV=1 | unc-13 |
| H9XVP2 | 1398 | S | 1.356 | 0.03949778 | Up | Unc-13, isoform E OS=Drosophila melanogaster OX=7227 GN=unc-13 PE=4 SV=1 | unc-13 |
| L0MLJ3 | 308 | S | 1.429 | 0.025393104 | Up | Uncharacterized protein, isoform M OS=Drosophila melanogaster OX=7227 GN=Dmel\CG1674 PE=1 SV=1 | Dmel\CG1674 |
| L0MLJ3 | 577 | S | 1.693 | 0.001287401 | Up | Uncharacterized protein, isoform M OS=Drosophila melanogaster OX=7227 GN=Dmel\CG1674 PE=1 SV=1 | Dmel\CG1674 |
| L0MLM5 | 14 | S | 2.384 | 5.23044E-05 | Up | Uncharacterized protein, isoform B OS=Drosophila melanogaster OX=7227 GN=BEST:GH26713 PE=4 SV=1 | BEST:GH26713 |
| L0MPN7 | 370 | S | 1.927 | 0.014923627 | Up | Asator, isoform H OS=Drosophila melanogaster OX=7227 GN=Asator PE=1 SV=1 | Asator |
| M9PC84 | 9413 | S | 1.533 | 0.02504952 | Up | Muscle-specific protein 300 kDa, isoform L OS=Drosophila melanogaster OX=7227 GN=Msp300 PE=1 SV=1 | Msp300 |
| M9PC84 | 757 | S | 1.368 | 0.002666036 | Up | Muscle-specific protein 300 kDa, isoform L OS=Drosophila melanogaster OX=7227 GN=Msp300 PE=1 SV=1 | Msp300 |
| M9PC84 | 439 | S | 1.34 | 0.000868611 | Up | Muscle-specific protein 300 kDa, isoform L OS=Drosophila melanogaster OX=7227 GN=Msp300 PE=1 SV=1 | Msp300 |
| M9MRG0 | 646 | S | 1.832 | 0.003152055 | Up | Nckx30C, isoform E OS=Drosophila melanogaster OX=7227 GN=Nckx30C PE=3 SV=1 | Nckx30C |
| X2JDP6 | 8 | S | 1.595 | 0.006487999 | Up | Nucleosome-destabilizing factor, isoform C OS=Drosophila melanogaster OX=7227 GN=Ndf PE=4 SV=1 | Ndf |
| X2JDP6 | 10 | S | 1.345 | 0.048252815 | Up | Nucleosome-destabilizing factor, isoform C OS=Drosophila melanogaster OX=7227 GN=Ndf PE=4 SV=1 | Ndf |
| M9MRJ4 | 2444 | S | 1.533 | 0.002316475 | Up | Muscle-specific protein 300 kDa, isoform G OS=Drosophila melanogaster OX=7227 GN=Msp300 PE=1 SV=1 | Msp300 |
| M9MSI8 | 3958 | S | 1.395 | 0.043652818 | Up | Tweek, isoform I OS=Drosophila melanogaster OX=7227 GN=tweek PE=4 SV=1 | tweek |
| M9MRX4 | 11739 | S | 0.67 | 0.004557465 | Down | Ankyrin 2, isoform U OS=Drosophila melanogaster OX=7227 GN=Ank2 PE=1 SV=1 | Ank2 |
| M9MRX4 | 1600 | S | 2.38 | 0.003508208 | Up | Ankyrin 2, isoform U OS=Drosophila melanogaster OX=7227 GN=Ank2 PE=1 SV=1 | Ank2 |
| M9MRX4 | 12636 | S | 1.572 | 0.028269857 | Up | Ankyrin 2, isoform U OS=Drosophila melanogaster OX=7227 GN=Ank2 PE=1 SV=1 | Ank2 |
| Q9VVB6 | 618 | S | 1.314 | 0.038703494 | Up | GH17145p OS=Drosophila melanogaster OX=7227 GN=Lmpt PE=1 SV=2 | Lmpt |
| Q9VVB6 | 535 | S | 1.407 | 0.008090952 | Up | GH17145p OS=Drosophila melanogaster OX=7227 GN=Lmpt PE=1 SV=2 | Lmpt |
| Q9VVB6 | 365 | S | 1.389 | 0.006865539 | Up | GH17145p OS=Drosophila melanogaster OX=7227 GN=Lmpt PE=1 SV=2 | Lmpt |

| | | | | | | | |
|---|---|---|---|---|---|---|---|
| Q9VVB6 | 431 | S | 1.721 | 0.01140899 | Up | GH17145p OS=Drosophila melanogaster OX=7227 GN=Lmpt PE=1 SV=2 | Lmpt |
| M9MRX4 | 5063 | S | 0.685 | 0.014021135 | Down | Ankyrin 2, isoform U OS=Drosophila melanogaster OX=7227 GN=Ank2 PE=1 SV=1 | Ank2 |
| M9MRX4 | 4987 | S | 0.753 | 0.035871077 | Down | Ankyrin 2, isoform U OS=Drosophila melanogaster OX=7227 GN=Ank2 PE=1 SV=1 | Ank2 |
| M9MRX4 | 3199 | S | 1.429 | 0.005352431 | Up | Ankyrin 2, isoform U OS=Drosophila melanogaster OX=7227 GN=Ank2 PE=1 SV=1 | Ank2 |
| M9MRX4 | 5443 | S | 0.587 | 0.000868214 | Down | Ankyrin 2, isoform U OS=Drosophila melanogaster OX=7227 GN=Ank2 PE=1 SV=1 | Ank2 |
| M9MRX4 | 5438 | S | 0.581 | 0.001260957 | Down | Ankyrin 2, isoform U OS=Drosophila melanogaster OX=7227 GN=Ank2 PE=1 SV=1 | Ank2 |
| M9MRX4 | 3194 | S | 1.466 | 0.014302742 | Up | Ankyrin 2, isoform U OS=Drosophila melanogaster OX=7227 GN=Ank2 PE=1 SV=1 | Ank2 |
| M9MRX4 | 10552 | S | 0.673 | 0.01349332 | Down | Ankyrin 2, isoform U OS=Drosophila melanogaster OX=7227 GN=Ank2 PE=1 SV=1 | Ank2 |
| M9MRX4 | 3442 | S | 1.877 | 8.12769E-07 | Up | Ankyrin 2, isoform U OS=Drosophila melanogaster OX=7227 GN=Ank2 PE=1 SV=1 | Ank2 |
| M9MS14 | 242 | S | 1.833 | 0.000267599 | Up | Moody, isoform C OS=Drosophila melanogaster OX=7227 GN=moody PE=3 SV=1 | moody |
| M9MS14 | 424 | S | 1.979 | 0.000265142 | Up | Moody, isoform C OS=Drosophila melanogaster OX=7227 GN=moody PE=3 SV=1 | moody |
| M9MS14 | 396 | S | 1.866 | 0.000237679 | Up | Moody, isoform C OS=Drosophila melanogaster OX=7227 GN=moody PE=3 SV=1 | moody |
| M9MS14 | 397 | S | 1.866 | 0.000237679 | Up | Moody, isoform C OS=Drosophila melanogaster OX=7227 GN=moody PE=3 SV=1 | moody |
| M9MS15 | 256 | S | 1.534 | 0.00170168 | Up | Neurotactin, isoform C OS=Drosophila melanogaster OX=7227 GN=Nrt PE=4 SV=1 | Nrt |
| M9MS26 | 512 | S | 1.36 | 0.038038906 | Up | Slipper, isoform C OS=Drosophila melanogaster OX=7227 GN=slpr PE=4 SV=1 | slpr |
| M9MS26 | 1141 | S | 1.446 | 0.004244895 | Up | Slipper, isoform C OS=Drosophila melanogaster OX=7227 GN=slpr PE=4 SV=1 | slpr |
| M9MS31 | 1291 | S | 0.754 | 0.000190042 | Down | Diacylglycerol kinase OS=Drosophila melanogaster OX=7227 GN=rdgA PE=3 SV=1 | rdgA |
| Q9VPA7 | 93 | S | 2.375 | 0.004644609 | Up | Protein kinase, cAMP-dependent, regulatory subunit type 1, isoform O OS=Drosophila melanogaster OX=7227 GN=Pka-R1 PE=1 SV=2 | Pka-R1 |
| Q9VPA7 | 72 | S | 1.563 | 0.03805615 | Up | Protein kinase, cAMP-dependent, regulatory subunit type 1, isoform O OS=Drosophila melanogaster OX=7227 GN=Pka-R1 PE=1 SV=2 | Pka-R1 |
| M9PGT0 | 1076 | S | 1.523 | 0.000227492 | Up | Uncharacterized protein, isoform R OS=Drosophila melanogaster OX=7227 GN=CG3960 PE=1 SV=1 | CG3960 |
| M9PGT0 | 2888 | S | 1.314 | 0.000363746 | Up | Uncharacterized protein, isoform R OS=Drosophila melanogaster OX=7227 GN=CG3960 PE=1 SV=1 | CG3960 |
| M9PGT0 | 2731 | S | 1.338 | 0.00695759 | Up | Uncharacterized protein, isoform R OS=Drosophila melanogaster OX=7227 GN=CG3960 PE=1 SV=1 | CG3960 |
| M9PGT0 | 1190 | S | 2.194 | 9.68267E-06 | Up | Uncharacterized protein, isoform R OS=Drosophila melanogaster OX=7227 GN=CG3960 PE=1 SV=1 | CG3960 |
| M9PGT0 | 1118 | S | 1.378 | 0.00919118 | Up | Uncharacterized protein, isoform R OS=Drosophila melanogaster OX=7227 GN=CG3960 PE=1 SV=1 | CG3960 |
| M9PHM3 | 802 | S | 1.49 | 0.003088238 | Up | Ether a go-go, isoform G OS=Drosophila melanogaster OX=7227 GN=eag PE=4 SV=1 | eag |
| Q7YZA4 | 387 | S | 1.317 | 0.003784535 | Up | Dishevelled associated activator of morphogenesis, isoform A OS=Drosophila melanogaster OX=7227 GN=DAAM PE=4 SV=1 | DAAM |
| M9MSQ1 | 741 | S | 1.379 | 0.019211216 | Up | Uncharacterized protein, isoform B OS=Drosophila melanogaster OX=7227 GN=anon-15Db PE=4 SV=1 | anon-15Db |
| M9NCQ1 | 726 | S | 1.485 | 0.005079895 | Up | Uncharacterized protein, isoform B OS=Drosophila melanogaster OX=7227 GN=Dmel\CG31663 PE=1 SV=1 | Dmel\CG31663 |
| M9NCQ1 | 890 | S | 2.49 | 0.000461489 | Up | Uncharacterized protein, isoform B OS=Drosophila melanogaster OX=7227 GN=Dmel\CG31663 PE=1 SV=1 | Dmel\CG31663 |
| M9NCS8 | 927 | S | 1.339 | 0.039321206 | Up | Glutactin, isoform D OS=Drosophila melanogaster OX=7227 GN=Glt PE=4 SV=1 | Glt |
| M9PBJ2 | 679 | S | 1.308 | 0.003431937 | Up | Lethal (2) giant larvae, isoform K OS=Drosophila melanogaster OX=7227 GN=l(2)gl PE=4 SV=1 | l(2)gl |
| M9PBJ2 | 984 | S | 1.373 | 0.027739649 | Up | Lethal (2) giant larvae, isoform K OS=Drosophila melanogaster OX=7227 GN=l(2)gl PE=4 SV=1 | l(2)gl |
| Q7KV91 | 69 | S | 0.685 | 0.020187939 | Down | Purine nucleoside phosphorylase OS=Drosophila melanogaster OX=7227 GN=Dmel\CG16758 PE=1 SV=1 | Dmel\CG16758 |
| Q8MV37 | 399 | S | 1.605 | 0.003134879 | Up | Myosin binding subunit of myosin phosphatase OS=Drosophila melanogaster OX=7227 GN=Mbs PE=1 SV=1 | Mbs |
| Q8MV37 | 370 | S | 1.362 | 0.019784179 | Up | Myosin binding subunit of myosin phosphatase OS=Drosophila melanogaster OX=7227 GN=Mbs PE=1 SV=1 | Mbs |
| M9NDD2 | 624 | S | 1.516 | 0.031569732 | Up | Adenylyl cyclase 76E, isoform B OS=Drosophila melanogaster OX=7227 GN=Ac76E PE=3 SV=1 | Ac76E |
| M9NDD2 | 627 | S | 1.516 | 0.031569732 | Up | Adenylyl cyclase 76E, isoform B OS=Drosophila melanogaster OX=7227 GN=Ac76E PE=3 SV=1 | Ac76E |
| M9NDE3 | 2800 | S | 1.311 | 4.91375E-05 | Up | Protein bark beetle OS=Drosophila melanogaster OX=7227 GN=bark PE=1 SV=1 | bark |
| M9NDE3 | 2803 | S | 1.311 | 4.91375E-05 | Up | Protein bark beetle OS=Drosophila melanogaster OX=7227 GN=bark PE=1 SV=1 | bark |
| M9NDE3 | 2755 | S | 1.755 | 0.002192141 | Up | Protein bark beetle OS=Drosophila melanogaster OX=7227 GN=bark PE=1 SV=1 | bark |
| M9NDE3 | 2759 | S | 1.755 | 0.002192141 | Up | Protein bark beetle OS=Drosophila melanogaster OX=7227 GN=bark PE=1 SV=1 | bark |
| M9NDE3 | 2975 | S | 1.392 | 0.003399953 | Up | Protein bark beetle OS=Drosophila melanogaster OX=7227 GN=bark PE=1 SV=1 | bark |
| M9NDE3 | 2978 | S | 1.689 | 0.001256687 | Up | Protein bark beetle OS=Drosophila melanogaster OX=7227 GN=bark PE=1 SV=1 | bark |
| M9NDE4 | 578 | S | 1.613 | 0.014346133 | Up | Uncharacterized protein, isoform C OS=Drosophila melanogaster OX=7227 GN=Dmel\CG12502 PE=4 SV=1 | Dmel\CG12502 |
| M9NDE4 | 16 | S | 1.366 | 0.039000628 | Up | Uncharacterized protein, isoform C OS=Drosophila melanogaster OX=7227 GN=Dmel\CG12502 PE=4 SV=1 | Dmel\CG12502 |
| M9NDL7 | 637 | S | 1.463 | 0.024972276 | Up | RALBP1 associated Eps domain containing, isoform B OS=Drosophila melanogaster OX=7227 GN=Reps PE=1 SV=1 | Reps |
| M9NDL7 | 452 | S | 1.432 | 0.023790809 | Up | RALBP1 associated Eps domain containing, isoform B OS=Drosophila melanogaster OX=7227 GN=Reps PE=1 SV=1 | Reps |
| X2JFA0 | 122 | S | 1.543 | 0.000412856 | Up | AMP deaminase OS=Drosophila melanogaster OX=7227 GN=AMPdeam PE=1 SV=1 | AMPdeam |
| X2JFA0 | 139 | S | 0.755 | 0.00175879 | Down | AMP deaminase OS=Drosophila melanogaster OX=7227 GN=AMPdeam PE=1 SV=1 | AMPdeam |
| M9PHR5 | 1561 | S | 1.475 | 0.010097954 | Up | Ectoderm-expressed 4, isoform K OS=Drosophila melanogaster OX=7227 GN=Sarm PE=1 SV=1 | Sarm |
| M9NF47 | 1286 | S | 1.582 | 0.000619899 | Up | Uncharacterized protein, isoform J OS=Drosophila melanogaster OX=7227 GN=CG32355 PE=1 SV=1 | CG32355 |
| M9NF47 | 1559 | S | 3.347 | 0.006197769 | Up | Uncharacterized protein, isoform J OS=Drosophila melanogaster OX=7227 GN=CG32355 PE=1 SV=1 | CG32355 |
| M9NF47 | 1308 | S | 1.936 | 0.049659158 | Up | Uncharacterized protein, isoform J OS=Drosophila melanogaster OX=7227 GN=CG32355 PE=1 SV=1 | CG32355 |
| M9NF47 | 1185 | S | 1.786 | 0.002925284 | Up | Uncharacterized protein, isoform J OS=Drosophila melanogaster OX=7227 GN=CG32355 PE=1 SV=1 | CG32355 |
| M9NE59 | 369 | S | 1.377 | 0.017181696 | Up | Uncharacterized protein, isoform G OS=Drosophila melanogaster OX=7227 GN=CG6239 PE=1 SV=1 | CG6239 |
| M9NE59 | 7 | S | 1.88 | 0.029982572 | Up | Uncharacterized protein, isoform G OS=Drosophila melanogaster OX=7227 GN=CG6239 PE=1 SV=1 | CG6239 |
| M9NE73 | 769 | S | 1.318 | 0.002513767 | Up | Calpain-B, isoform B OS=Drosophila melanogaster OX=7227 GN=CalpB PE=3 SV=1 | CalpB |
| M9NEA5 | 975 | S | 1.573 | 0.026779366 | Up | Homeobox protein cut-like OS=Drosophila melanogaster OX=7227 GN=ct PE=1 SV=1 | ct |
| Q8IR69 | 278 | S | 1.313 | 0.012373651 | Up | Histone deacetylase OS=Drosophila melanogaster OX=7227 GN=HDAC4 | HDAC4 |

| | | | | | | | |
|---|---|---|---|---|---|---|---|
| | | | | | | PE=1 SV=1 | |
| M9NEH6 | 926 | S | 1.404 | 0.003521756 | Up | Histone deacetylase 6, isoform F OS=Drosophila melanogaster OX=7227 GN=HDAC6 PE=1 SV=1 | HDAC6 |
| M9NFE1 | 1157 | S | 1.462 | 0.006051258 | Up | Uncharacterized protein, isoform C OS=Drosophila melanogaster OX=7227 GN=Dmel\CG14200 PE=1 SV=1 | Dmel\CG14200 |
| X2J8V5 | 147 | S | 0.422 | 0.00175161 | Down | Arrestin 1, isoform C OS=Drosophila melanogaster OX=7227 GN=Arr1 PE=4 SV=1 | Arr1 |
| M9NEW2 | 368 | S | 5.954 | 1.36849E-06 | Up | Histone acetyltransferase OS=Drosophila melanogaster OX=7227 GN=chm PE=3 SV=1 | chm |
| M9NF18 | 764 | S | 2.978 | 0.000808791 | Up | Uncharacterized protein, isoform D OS=Drosophila melanogaster OX=7227 GN=Dmel\CG2247 PE=1 SV=1 | Dmel\CG2247 |
| M9NF47 | 775 | S | 1.302 | 0.016525894 | Up | Uncharacterized protein, isoform J OS=Drosophila melanogaster OX=7227 GN=CG32355 PE=1 SV=1 | CG32355 |
| M9NF47 | 683 | S | 1.764 | 0.00020721 | Up | Uncharacterized protein, isoform J OS=Drosophila melanogaster OX=7227 GN=CG32355 PE=1 SV=1 | CG32355 |
| M9NF51 | 1968 | S | 1.79 | 8.55123E-06 | Up | Rutabaga, isoform B OS=Drosophila melanogaster OX=7227 GN=rut PE=3 SV=1 | rut |
| M9NF51 | 1971 | S | 1.79 | 8.55123E-06 | Up | Rutabaga, isoform B OS=Drosophila melanogaster OX=7227 GN=rut PE=3 SV=1 | rut |
| M9NFC0 | 85 | S | 0.711 | 0.021361036 | Down | Wings up A, isoform I OS=Drosophila melanogaster OX=7227 GN=wupA PE=1 SV=1 | wupA |
| Q9VVA4 | 241 | S | 0.662 | 0.004933334 | Down | GH26789p OS=Drosophila melanogaster OX=7227 GN=GS PE=1 SV=2 | GS |
| M9PBP8 | 2067 | S | 1.87 | 0.005296711 | Up | Axotactin, isoform F OS=Drosophila melanogaster OX=7227 GN=axo PE=4 SV=1 | axo |
| M9PBP8 | 2068 | S | 1.87 | 0.005296711 | Up | Axotactin, isoform F OS=Drosophila melanogaster OX=7227 GN=axo PE=4 SV=1 | axo |
| M9PBP8 | 1869 | S | 1.479 | 0.00640866 | Up | Axotactin, isoform F OS=Drosophila melanogaster OX=7227 GN=axo PE=4 SV=1 | axo |
| M9NFQ4 | 331 | S | 1.631 | 0.01934071 | Up | Uncharacterized protein, isoform E OS=Drosophila melanogaster OX=7227 GN=CG3791 PE=4 SV=1 | CG3791 |
| M9NFZ9 | 180 | S | 1.331 | 0.006957151 | Up | Ringmaker, isoform B OS=Drosophila melanogaster OX=7227 GN=ringer PE=4 SV=1 | ringer |
| M9NGG5 | 1773 | S | 0.71 | 8.20636E-05 | Down | Futsch, isoform F OS=Drosophila melanogaster OX=7227 GN=futsch PE=1 SV=1 | futsch |
| M9NGG5 | 1581 | S | 1.667 | 0.047608451 | Up | Futsch, isoform F OS=Drosophila melanogaster OX=7227 GN=futsch PE=1 SV=1 | futsch |
| M9NGG5 | 1585 | S | 1.667 | 0.047608451 | Up | Futsch, isoform F OS=Drosophila melanogaster OX=7227 GN=futsch PE=1 SV=1 | futsch |
| M9NGG5 | 4725 | S | 1.309 | 0.021226607 | Up | Futsch, isoform F OS=Drosophila melanogaster OX=7227 GN=futsch PE=1 SV=1 | futsch |
| M9NGG5 | 4879 | S | 1.358 | 0.011307314 | Up | Futsch, isoform F OS=Drosophila melanogaster OX=7227 GN=futsch PE=1 SV=1 | futsch |
| M9NGG5 | 2537 | S | 0.765 | 0.009607982 | Down | Futsch, isoform F OS=Drosophila melanogaster OX=7227 GN=futsch PE=1 SV=1 | futsch |
| M9NGG5 | 2543 | S | 0.749 | 0.009131436 | Down | Futsch, isoform F OS=Drosophila melanogaster OX=7227 GN=futsch PE=1 SV=1 | futsch |
| M9NGG5 | 2547 | S | 0.597 | 0.002112993 | Down | Futsch, isoform F OS=Drosophila melanogaster OX=7227 GN=futsch PE=1 SV=1 | futsch |
| M9NGG5 | 5081 | S | 1.709 | 0.001638298 | Up | Futsch, isoform F OS=Drosophila melanogaster OX=7227 GN=futsch PE=1 SV=1 | futsch |
| M9NGG5 | 2914 | S | 1.578 | 0.007758441 | Up | Futsch, isoform F OS=Drosophila melanogaster OX=7227 GN=futsch PE=1 SV=1 | futsch |
| M9NGG5 | 2918 | S | 1.489 | 0.005971722 | Up | Futsch, isoform F OS=Drosophila melanogaster OX=7227 GN=futsch PE=1 SV=1 | futsch |
| M9NGG5 | 2789 | S | 1.864 | 0.00099343 | Up | Futsch, isoform F OS=Drosophila melanogaster OX=7227 GN=futsch PE=1 SV=1 | futsch |
| M9NGG5 | 2427 | S | 0.59 | 0.000234359 | Down | Futsch, isoform F OS=Drosophila melanogaster OX=7227 GN=futsch PE=1 SV=1 | futsch |
| M9NGG5 | 4394 | S | 0.765 | 0.002075862 | Down | Futsch, isoform F OS=Drosophila melanogaster OX=7227 GN=futsch PE=1 SV=1 | futsch |
| M9NGG5 | 2025 | S | 0.76 | 0.031075783 | Down | Futsch, isoform F OS=Drosophila melanogaster OX=7227 GN=futsch PE=1 SV=1 | futsch |
| M9NGG5 | 2029 | S | 0.713 | 0.013707606 | Down | Futsch, isoform F OS=Drosophila melanogaster OX=7227 GN=futsch PE=1 SV=1 | futsch |
| M9NGG5 | 2033 | S | 1.807 | 0.000213105 | Up | Futsch, isoform F OS=Drosophila melanogaster OX=7227 GN=futsch PE=1 SV=1 | futsch |
| M9NGG5 | 2551 | S | 0.735 | 0.009198884 | Down | Futsch, isoform F OS=Drosophila melanogaster OX=7227 GN=futsch PE=1 SV=1 | futsch |
| M9NGG5 | 4124 | S | 0.749 | 0.00293085 | Down | Futsch, isoform F OS=Drosophila melanogaster OX=7227 GN=futsch PE=1 SV=1 | futsch |
| M9NGG5 | 4924 | S | 1.417 | 0.005989467 | Up | Futsch, isoform F OS=Drosophila melanogaster OX=7227 GN=futsch PE=1 SV=1 | futsch |
| M9NGG5 | 3579 | S | 0.724 | 0.006230444 | Down | Futsch, isoform F OS=Drosophila melanogaster OX=7227 GN=futsch PE=1 SV=1 | futsch |
| M9NGG5 | 3583 | S | 0.604 | 0.001744363 | Down | Futsch, isoform F OS=Drosophila melanogaster OX=7227 GN=futsch PE=1 SV=1 | futsch |
| M9NGG5 | 2972 | S | 1.307 | 0.007927903 | Up | Futsch, isoform F OS=Drosophila melanogaster OX=7227 GN=futsch PE=1 SV=1 | futsch |
| M9NGG5 | 4695 | S | 1.375 | 0.039794221 | Up | Futsch, isoform F OS=Drosophila melanogaster OX=7227 GN=futsch PE=1 SV=1 | futsch |
| M9NGG5 | 2019 | S | 0.753 | 0.043886609 | Down | Futsch, isoform F OS=Drosophila melanogaster OX=7227 GN=futsch PE=1 SV=1 | futsch |
| M9NGG5 | 3354 | S | 0.681 | 0.003858256 | Down | Futsch, isoform F OS=Drosophila melanogaster OX=7227 GN=futsch PE=1 SV=1 | futsch |
| M9NGG5 | 2070 | S | 1.361 | 0.001210834 | Up | Futsch, isoform F OS=Drosophila melanogaster OX=7227 GN=futsch PE=1 SV=1 | futsch |
| M9NGG5 | 2074 | S | 1.365 | 0.013195888 | Up | Futsch, isoform F OS=Drosophila melanogaster OX=7227 GN=futsch PE=1 SV=1 | futsch |
| M9NGG5 | 2366 | S | 1.473 | 0.000506067 | Up | Futsch, isoform F OS=Drosophila melanogaster OX=7227 GN=futsch PE=1 SV=1 | futsch |
| M9NGG5 | 2370 | S | 1.609 | 0.003578909 | Up | Futsch, isoform F OS=Drosophila melanogaster OX=7227 GN=futsch PE=1 SV=1 | futsch |
| M9NGG5 | 3735 | S | 2.444 | 0.001017692 | Up | Futsch, isoform F OS=Drosophila melanogaster OX=7227 GN=futsch PE=1 SV=1 | futsch |
| M9NGG5 | 3739 | S | 2.915 | 0.001239124 | Up | Futsch, isoform F OS=Drosophila melanogaster OX=7227 GN=futsch PE=1 SV=1 | futsch |
| M9NGG5 | 3743 | S | 2.579 | 0.001198999 | Up | Futsch, isoform F OS=Drosophila melanogaster OX=7227 GN=futsch PE=1 SV=1 | futsch |
| M9NGG5 | 3439 | S | 1.901 | 0.000596242 | Up | Futsch, isoform F OS=Drosophila melanogaster OX=7227 GN=futsch PE=1 SV=1 | futsch |
| M9NGG5 | 3443 | S | 2.648 | 0.002038938 | Up | Futsch, isoform F OS=Drosophila melanogaster OX=7227 GN=futsch PE=1 SV=1 | futsch |
| M9NGG5 | 3447 | S | 1.346 | 0.011412678 | Up | Futsch, isoform F OS=Drosophila melanogaster OX=7227 GN=futsch PE=1 SV=1 | futsch |
| M9NGG5 | 1631 | S | 0.764 | 0.003595601 | Down | Futsch, isoform F OS=Drosophila melanogaster OX=7227 GN=futsch PE=1 SV=1 | futsch |
| M9NGG5 | 1635 | S | 0.75 | 0.004251938 | Down | Futsch, isoform F OS=Drosophila melanogaster OX=7227 GN=futsch PE=1 SV=1 | futsch |
| M9NGG5 | 1679 | S | 0.661 | 0.005173348 | Down | Futsch, isoform F OS=Drosophila melanogaster OX=7227 GN=futsch PE=1 SV=1 | futsch |
| M9NGG5 | 3924 | S | 1.531 | 0.002961781 | Up | Futsch, isoform F OS=Drosophila melanogaster OX=7227 GN=futsch PE=1 SV=1 | futsch |
| M9NGG5 | 3039 | S | 0.666 | 0.005748457 | Down | Futsch, isoform F OS=Drosophila melanogaster OX=7227 GN=futsch PE=1 SV=1 | futsch |
| M9NGG5 | 2524 | S | 2.02 | 0.003687981 | Up | Futsch, isoform F OS=Drosophila melanogaster OX=7227 GN=futsch PE=1 SV=1 | futsch |

| | | | | | | | |
|---|---|---|---|---|---|---|---|
| M9NGG5 | 1039 | S | 1.456 | 0.006921377 | Up | Futsch, isoform F OS=Drosophila melanogaster OX=7227 GN=futsch PE=1 SV=1 | futsch |
| M9NGG5 | 2014 | S | 0.637 | 0.019037517 | Down | Futsch, isoform F OS=Drosophila melanogaster OX=7227 GN=futsch PE=1 SV=1 | futsch |
| M9NGG5 | 3365 | S | 0.762 | 0.002634637 | Down | Futsch, isoform F OS=Drosophila melanogaster OX=7227 GN=futsch PE=1 SV=1 | futsch |
| M9NGG5 | 2358 | S | 0.627 | 0.001297524 | Down | Futsch, isoform F OS=Drosophila melanogaster OX=7227 GN=futsch PE=1 SV=1 | futsch |
| M9NGG5 | 2362 | S | 0.462 | 0.000311323 | Down | Futsch, isoform F OS=Drosophila melanogaster OX=7227 GN=futsch PE=1 SV=1 | futsch |
| M9NGG5 | 3944 | S | 0.759 | 0.006556108 | Down | Futsch, isoform F OS=Drosophila melanogaster OX=7227 GN=futsch PE=1 SV=1 | futsch |
| M9NGG5 | 3957 | S | 2.337 | 0.001889312 | Up | Futsch, isoform F OS=Drosophila melanogaster OX=7227 GN=futsch PE=1 SV=1 | futsch |
| M9NGG5 | 3961 | S | 2.337 | 0.001889312 | Up | Futsch, isoform F OS=Drosophila melanogaster OX=7227 GN=futsch PE=1 SV=1 | futsch |
| M9NGG5 | 1695 | S | 1.626 | 0.000672628 | Up | Futsch, isoform F OS=Drosophila melanogaster OX=7227 GN=futsch PE=1 SV=1 | futsch |
| M9NGG5 | 2528 | S | 2 | 0.013172546 | Up | Futsch, isoform F OS=Drosophila melanogaster OX=7227 GN=futsch PE=1 SV=1 | futsch |
| M9NGG5 | 1648 | S | 1.502 | 0.035405422 | Up | Futsch, isoform F OS=Drosophila melanogaster OX=7227 GN=futsch PE=1 SV=1 | futsch |
| M9NGG5 | 1657 | S | 0.685 | 0.005040918 | Down | Futsch, isoform F OS=Drosophila melanogaster OX=7227 GN=futsch PE=1 SV=1 | futsch |
| M9NGG5 | 3071 | S | 0.663 | 0.005532699 | Down | Futsch, isoform F OS=Drosophila melanogaster OX=7227 GN=futsch PE=1 SV=1 | futsch |
| M9NGG5 | 1617 | S | 0.722 | 0.00343978 | Down | Futsch, isoform F OS=Drosophila melanogaster OX=7227 GN=futsch PE=1 SV=1 | futsch |
| M9NGG5 | 4035 | S | 1.613 | 0.010234378 | Up | Futsch, isoform F OS=Drosophila melanogaster OX=7227 GN=futsch PE=1 SV=1 | futsch |
| M9NGG5 | 3620 | S | 0.614 | 0.001450927 | Down | Futsch, isoform F OS=Drosophila melanogaster OX=7227 GN=futsch PE=1 SV=1 | futsch |
| M9NGG5 | 3624 | S | 0.717 | 0.012869268 | Down | Futsch, isoform F OS=Drosophila melanogaster OX=7227 GN=futsch PE=1 SV=1 | futsch |
| M9NGG5 | 4023 | S | 0.557 | 0.001030751 | Down | Futsch, isoform F OS=Drosophila melanogaster OX=7227 GN=futsch PE=1 SV=1 | futsch |
| M9NGG5 | 4027 | S | 0.578 | 0.00067124 | Down | Futsch, isoform F OS=Drosophila melanogaster OX=7227 GN=futsch PE=1 SV=1 | futsch |
| M9NGG5 | 2057 | S | 1.381 | 0.015802707 | Up | Futsch, isoform F OS=Drosophila melanogaster OX=7227 GN=futsch PE=1 SV=1 | futsch |
| M9NGG5 | 2062 | S | 0.646 | 0.000908078 | Down | Futsch, isoform F OS=Drosophila melanogaster OX=7227 GN=futsch PE=1 SV=1 | futsch |
| M9NGG5 | 2066 | S | 0.497 | 0.003011306 | Down | Futsch, isoform F OS=Drosophila melanogaster OX=7227 GN=futsch PE=1 SV=1 | futsch |
| M9NGG5 | 3839 | S | 0.754 | 0.006278528 | Down | Futsch, isoform F OS=Drosophila melanogaster OX=7227 GN=futsch PE=1 SV=1 | futsch |
| M9NGG5 | 3842 | S | 0.754 | 0.006278528 | Down | Futsch, isoform F OS=Drosophila melanogaster OX=7227 GN=futsch PE=1 SV=1 | futsch |
| M9NGG5 | 1764 | S | 1.946 | 0.0009924 | Up | Futsch, isoform F OS=Drosophila melanogaster OX=7227 GN=futsch PE=1 SV=1 | futsch |
| M9NGG5 | 3099 | S | 1.799 | 0.003452279 | Up | Futsch, isoform F OS=Drosophila melanogaster OX=7227 GN=futsch PE=1 SV=1 | futsch |
| M9NGG5 | 3246 | S | 0.636 | 0.004629283 | Down | Futsch, isoform F OS=Drosophila melanogaster OX=7227 GN=futsch PE=1 SV=1 | futsch |
| M9NGG5 | 2435 | S | 1.532 | 0.000873054 | Up | Futsch, isoform F OS=Drosophila melanogaster OX=7227 GN=futsch PE=1 SV=1 | futsch |
| M9NGG5 | 3411 | S | 2.117 | 0.001224788 | Up | Futsch, isoform F OS=Drosophila melanogaster OX=7227 GN=futsch PE=1 SV=1 | futsch |
| M9NGG5 | 3415 | S | 2.13 | 0.001510222 | Up | Futsch, isoform F OS=Drosophila melanogaster OX=7227 GN=futsch PE=1 SV=1 | futsch |
| M9NGG5 | 3419 | S | 2.124 | 0.001053497 | Up | Futsch, isoform F OS=Drosophila melanogaster OX=7227 GN=futsch PE=1 SV=1 | futsch |
| M9NGG5 | 1710 | S | 1.81 | 0.008197024 | Up | Futsch, isoform F OS=Drosophila melanogaster OX=7227 GN=futsch PE=1 SV=1 | futsch |
| M9NGG5 | 1714 | S | 2.275 | 0.002325952 | Up | Futsch, isoform F OS=Drosophila melanogaster OX=7227 GN=futsch PE=1 SV=1 | futsch |
| M9NGG5 | 4182 | S | 1.501 | 0.001746622 | Up | Futsch, isoform F OS=Drosophila melanogaster OX=7227 GN=futsch PE=1 SV=1 | futsch |
| M9NGG5 | 2835 | S | 0.672 | 0.026773253 | Down | Futsch, isoform F OS=Drosophila melanogaster OX=7227 GN=futsch PE=1 SV=1 | futsch |
| M9NGG5 | 1157 | S | 0.658 | 0.003217603 | Down | Futsch, isoform F OS=Drosophila melanogaster OX=7227 GN=futsch PE=1 SV=1 | futsch |
| M9NGG5 | 1095 | S | 1.426 | 0.035289596 | Up | Futsch, isoform F OS=Drosophila melanogaster OX=7227 GN=futsch PE=1 SV=1 | futsch |
| M9NGG5 | 2037 | S | 1.669 | 0.001223846 | Up | Futsch, isoform F OS=Drosophila melanogaster OX=7227 GN=futsch PE=1 SV=1 | futsch |
| M9NGG5 | 3603 | S | 0.511 | 0.007410229 | Down | Futsch, isoform F OS=Drosophila melanogaster OX=7227 GN=futsch PE=1 SV=1 | futsch |
| M9NGG5 | 2990 | S | 1.377 | 0.04102553 | Up | Futsch, isoform F OS=Drosophila melanogaster OX=7227 GN=futsch PE=1 SV=1 | futsch |
| M9NGG5 | 3122 | S | 0.696 | 0.002564212 | Down | Futsch, isoform F OS=Drosophila melanogaster OX=7227 GN=futsch PE=1 SV=1 | futsch |
| M9NGG5 | 3136 | S | 1.445 | 0.017327823 | Up | Futsch, isoform F OS=Drosophila melanogaster OX=7227 GN=futsch PE=1 SV=1 | futsch |
| M9NGG5 | 4059 | S | 0.645 | 0.001522991 | Down | Futsch, isoform F OS=Drosophila melanogaster OX=7227 GN=futsch PE=1 SV=1 | futsch |
| M9NGG5 | 1736 | S | 0.704 | 0.023237768 | Down | Futsch, isoform F OS=Drosophila melanogaster OX=7227 GN=futsch PE=1 SV=1 | futsch |
| M9NGG5 | 2588 | S | 1.476 | 0.023641985 | Up | Futsch, isoform F OS=Drosophila melanogaster OX=7227 GN=futsch PE=1 SV=1 | futsch |
| M9NGG5 | 2592 | S | 0.726 | 0.024810018 | Down | Futsch, isoform F OS=Drosophila melanogaster OX=7227 GN=futsch PE=1 SV=1 | futsch |
| M9NGG5 | 2390 | S | 2.234 | 0.014393714 | Up | Futsch, isoform F OS=Drosophila melanogaster OX=7227 GN=futsch PE=1 SV=1 | futsch |
| M9NGG5 | 3398 | S | 1.622 | 0.005927076 | Up | Futsch, isoform F OS=Drosophila melanogaster OX=7227 GN=futsch PE=1 SV=1 | futsch |
| M9NGG5 | 3282 | S | 3.167 | 0.004495884 | Up | Futsch, isoform F OS=Drosophila melanogaster OX=7227 GN=futsch PE=1 SV=1 | futsch |
| M9NGG5 | 1627 | S | 0.746 | 0.008078744 | Down | Futsch, isoform F OS=Drosophila melanogaster OX=7227 GN=futsch PE=1 SV=1 | futsch |
| M9NGG5 | 1667 | S | 0.755 | 0.009074208 | Down | Futsch, isoform F OS=Drosophila melanogaster OX=7227 GN=futsch PE=1 SV=1 | futsch |
| M9NGG5 | 4160 | S | 1.868 | 0.048882702 | Up | Futsch, isoform F OS=Drosophila melanogaster OX=7227 GN=futsch PE=1 SV=1 | futsch |
| M9NGG5 | 1515 | S | 0.721 | 0.043164728 | Down | Futsch, isoform F OS=Drosophila melanogaster OX=7227 GN=futsch PE=1 SV=1 | futsch |
| M9NGG5 | 4106 | S | 1.43 | 0.001389324 | Up | Futsch, isoform F OS=Drosophila melanogaster OX=7227 GN=futsch PE=1 SV=1 | futsch |
| M9NGG5 | 2875 | S | 0.76 | 0.001800112 | Down | Futsch, isoform F OS=Drosophila melanogaster OX=7227 GN=futsch PE=1 SV=1 | futsch |
| M9NGG5 | 1595 | S | 0.704 | 0.047177044 | Down | Futsch, isoform F OS=Drosophila melanogaster OX=7227 GN=futsch PE=1 SV=1 | futsch |
| M9NGG5 | 2730 | S | 1.434 | 0.04813353 | Up | Futsch, isoform F OS=Drosophila melanogaster OX=7227 GN=futsch PE=1 SV=1 | futsch |
| M9NGQ5 | 472 | S | 1.47 | 0.000711278 | Up | Uncharacterized protein, isoform B OS=Drosophila melanogaster OX=7227 GN=Dmel\CG3556 PE=4 SV=1 | Dmel\CG3556 |

| | | | | | | | |
|---|---|---|---|---|---|---|---|
| M9NGQ9 | 366 | S | 2.371 | 4.24794E-05 | Up | Trapped in endoderm 1, isoform B OS=Drosophila melanogaster OX=7227 GN=Tre1 PE=3 SV=1 | Tre1 |
| Q59E11 | 330 | S | 1.867 | 0.003929147 | Up | IA-2 protein tyrosine phosphatase, isoform D OS=Drosophila melanogaster OX=7227 GN=IA-2 PE=1 SV=2 | IA-2 |
| M9PB50 | 505 | S | 1.516 | 0.007529333 | Up | Real-time, isoform B OS=Drosophila melanogaster OX=7227 GN=retm PE=4 SV=1 | retm |
| M9PB68 | 589 | S | 1.358 | 0.049806928 | Up | Purity of essence, isoform B OS=Drosophila melanogaster OX=7227 GN=poe PE=4 SV=1 | poe |
| M9PB85 | 79 | S | 2.199 | 0.000655547 | Up | Uncharacterized protein, isoform B OS=Drosophila melanogaster OX=7227 GN=Dmel\CG31712 PE=4 SV=1 | Dmel\CG31712 |
| M9PB85 | 25 | S | 1.665 | 0.006885494 | Up | Uncharacterized protein, isoform B OS=Drosophila melanogaster OX=7227 GN=Dmel\CG31712 PE=4 SV=1 | Dmel\CG31712 |
| M9PB85 | 27 | S | 1.665 | 0.006885494 | Up | Uncharacterized protein, isoform B OS=Drosophila melanogaster OX=7227 GN=Dmel\CG31712 PE=4 SV=1 | Dmel\CG31712 |
| M9PBF6 | 111 | S | 1.476 | 0.007548982 | Up | Dynamin associated protein 160, isoform G OS=Drosophila melanogaster OX=7227 GN=Dap160 PE=1 SV=1 | Dap160 |
| M9PBF6 | 305 | S | 1.547 | 0.00779512 | Up | Dynamin associated protein 160, isoform G OS=Drosophila melanogaster OX=7227 GN=Dap160 PE=1 SV=1 | Dap160 |
| M9PBF6 | 308 | S | 1.507 | 0.0085089 | Up | Dynamin associated protein 160, isoform G OS=Drosophila melanogaster OX=7227 GN=Dap160 PE=1 SV=1 | Dap160 |
| M9PBF6 | 328 | S | 1.384 | 0.030815851 | Up | Dynamin associated protein 160, isoform G OS=Drosophila melanogaster OX=7227 GN=Dap160 PE=1 SV=1 | Dap160 |
| M9PBF6 | 293 | S | 1.672 | 0.003076795 | Up | Dynamin associated protein 160, isoform G OS=Drosophila melanogaster OX=7227 GN=Dap160 PE=1 SV=1 | Dap160 |
| M9PBF6 | 925 | S | 2.182 | 0.004914042 | Up | Dynamin associated protein 160, isoform G OS=Drosophila melanogaster OX=7227 GN=Dap160 PE=1 SV=1 | Dap160 |
| M9PBF6 | 929 | S | 1.874 | 0.002931153 | Up | Dynamin associated protein 160, isoform G OS=Drosophila melanogaster OX=7227 GN=Dap160 PE=1 SV=1 | Dap160 |
| M9PBH6 | 143 | S | 1.498 | 0.000482688 | Up | Bric a brac 2, isoform B OS=Drosophila melanogaster OX=7227 GN=bab2 PE=4 SV=1 | bab2 |
| M9PBH6 | 147 | S | 1.498 | 0.000482688 | Up | Bric a brac 2, isoform B OS=Drosophila melanogaster OX=7227 GN=bab2 PE=4 SV=1 | bab2 |
| M9PDS3 | 6392 | S | 1.788 | 0.002222491 | Up | Sallimus, isoform T OS=Drosophila melanogaster OX=7227 GN=sls PE=4 SV=1 | sls |
| M9PDS3 | 16510 | S | 1.321 | 0.024506314 | Up | Sallimus, isoform T OS=Drosophila melanogaster OX=7227 GN=sls PE=4 SV=1 | sls |
| M9PDS3 | 16511 | S | 1.526 | 0.014211566 | Up | Sallimus, isoform T OS=Drosophila melanogaster OX=7227 GN=sls PE=4 SV=1 | sls |
| M9PDS3 | 5025 | S | 1.777 | 0.001995544 | Up | Sallimus, isoform T OS=Drosophila melanogaster OX=7227 GN=sls PE=4 SV=1 | sls |
| M9PDS3 | 5028 | S | 1.745 | 0.002136453 | Up | Sallimus, isoform T OS=Drosophila melanogaster OX=7227 GN=sls PE=4 SV=1 | sls |
| M9PDS3 | 15005 | S | 5.289 | 9.76861E-05 | Up | Sallimus, isoform T OS=Drosophila melanogaster OX=7227 GN=sls PE=4 SV=1 | sls |
| M9PDS3 | 5202 | S | 1.527 | 0.020693012 | Up | Sallimus, isoform T OS=Drosophila melanogaster OX=7227 GN=sls PE=4 SV=1 | sls |
| M9PDS3 | 5182 | S | 1.629 | 0.01290426 | Up | Sallimus, isoform T OS=Drosophila melanogaster OX=7227 GN=sls PE=4 SV=1 | sls |
| M9PDS3 | 16272 | S | 1.309 | 0.01712471 | Up | Sallimus, isoform T OS=Drosophila melanogaster OX=7227 GN=sls PE=4 SV=1 | sls |
| M9PDS3 | 16505 | S | 1.321 | 0.024506314 | Up | Sallimus, isoform T OS=Drosophila melanogaster OX=7227 GN=sls PE=4 SV=1 | sls |
| M9PDS3 | 16507 | S | 1.321 | 0.024506314 | Up | Sallimus, isoform T OS=Drosophila melanogaster OX=7227 GN=sls PE=4 SV=1 | sls |
| M9PDS3 | 4807 | S | 1.431 | 0.022760111 | Up | Sallimus, isoform T OS=Drosophila melanogaster OX=7227 GN=sls PE=4 SV=1 | sls |
| M9PBS9 | 318 | S | 1.426 | 3.07236E-05 | Up | Uncharacterized protein, isoform F OS=Drosophila melanogaster OX=7227 GN=Ist1 PE=1 SV=1 | Ist1 |
| M9PBV6 | 218 | S | 0.722 | 0.001216012 | Down | Nuclear localized protein 1, isoform B OS=Drosophila melanogaster OX=7227 GN=Nulp1 PE=4 SV=1 | Nulp1 |
| Q9VQF1 | 326 | S | 1.301 | 0.00768007 | Up | FI05237p OS=Drosophila melanogaster OX=7227 GN=Dmel\CG31689 PE=1 SV=1 | Dmel\CG31689 |
| M9PI01 | 26 | S | 1.515 | 1.71003E-05 | Up | Uncharacterized protein, isoform F OS=Drosophila melanogaster OX=7227 GN=CG34239 PE=4 SV=1 | CG34239 |
| M9PC73 | 632 | S | 1.378 | 0.023195686 | Up | Liprin-alpha, isoform C OS=Drosophila melanogaster OX=7227 GN=Liprin-alpha PE=1 SV=1 | Liprin-alpha |
| M9PI37 | 126 | S | 1.651 | 0.003980813 | Up | Sodium chloride cotransporter 69, isoform E OS=Drosophila melanogaster OX=7227 GN=Ncc69 PE=1 SV=1 | Ncc69 |
| M9PI37 | 129 | S | 2.701 | 0.00614983 | Up | Sodium chloride cotransporter 69, isoform E OS=Drosophila melanogaster OX=7227 GN=Ncc69 PE=1 SV=1 | Ncc69 |
| M9PI37 | 132 | S | 2.251 | 0.005797363 | Up | Sodium chloride cotransporter 69, isoform E OS=Drosophila melanogaster OX=7227 GN=Ncc69 PE=1 SV=1 | Ncc69 |
| M9PI37 | 892 | S | 1.486 | 0.000956852 | Up | Sodium chloride cotransporter 69, isoform E OS=Drosophila melanogaster OX=7227 GN=Ncc69 PE=1 SV=1 | Ncc69 |
| M9PI37 | 895 | S | 1.564 | 0.00216835 | Up | Sodium chloride cotransporter 69, isoform E OS=Drosophila melanogaster OX=7227 GN=Ncc69 PE=1 SV=1 | Ncc69 |
| M9PI37 | 913 | S | 2.229 | 2.26261E-05 | Up | Sodium chloride cotransporter 69, isoform E OS=Drosophila melanogaster OX=7227 GN=Ncc69 PE=1 SV=1 | Ncc69 |
| M9PI37 | 917 | S | 2.084 | 4.05668E-05 | Up | Sodium chloride cotransporter 69, isoform E OS=Drosophila melanogaster OX=7227 GN=Ncc69 PE=1 SV=1 | Ncc69 |
| M9PI37 | 194 | S | 5.817 | 0.003818892 | Up | Sodium chloride cotransporter 69, isoform E OS=Drosophila melanogaster OX=7227 GN=Ncc69 PE=1 SV=1 | Ncc69 |
| M9PI37 | 974 | S | 3.005 | 9.85193E-05 | Up | Sodium chloride cotransporter 69, isoform E OS=Drosophila melanogaster OX=7227 GN=Ncc69 PE=1 SV=1 | Ncc69 |
| M9PCA7 | 925 | S | 2.198 | 1.87393E-05 | Up | Sodium chloride cotransporter 69, isoform C OS=Drosophila melanogaster OX=7227 GN=Ncc69 PE=1 SV=1 | Ncc69 |
| M9PI37 | 874 | S | 3.393 | 0.000186464 | Up | Sodium chloride cotransporter 69, isoform E OS=Drosophila melanogaster OX=7227 GN=Ncc69 PE=1 SV=1 | Ncc69 |
| M9PI37 | 855 | S | 1.722 | 0.000155591 | Up | Sodium chloride cotransporter 69, isoform E OS=Drosophila melanogaster OX=7227 GN=Ncc69 PE=1 SV=1 | Ncc69 |
| M9PCK5 | 390 | S | 1.498 | 0.000572673 | Up | Mextli, isoform F OS=Drosophila melanogaster OX=7227 GN=mxt PE=4 SV=1 | mxt |
| Q7KUM8 | 182 | S | 0.66 | 0.017215267 | Down | Ecdysone-induced protein 28/29kD, isoform E OS=Drosophila melanogaster OX=7227 GN=Eip71CD PE=1 SV=2 | Eip71CD |
| M9PCQ6 | 991 | S | 1.68 | 0.00187422 | Up | Discs large 5, isoform C OS=Drosophila melanogaster OX=7227 GN=Dlg5 PE=1 SV=1 | Dlg5 |
| M9PCQ8 | 555 | S | 1.526 | 0.002989499 | Up | Chico, isoform B OS=Drosophila melanogaster OX=7227 GN=chico PE=4 SV=1 | chico |
| M9PD06 | 1356 | S | 2.047 | 0.001379501 | Up | Ubiquitin carboxyl-terminal hydrolase 32 OS=Drosophila melanogaster OX=7227 GN=Usp32 PE=3 SV=1 | Usp32 |
| Q9VK62 | 375 | S | 1.8 | 0.001375271 | Up | GM13388p OS=Drosophila melanogaster OX=7227 GN=spict PE=2 SV=1 | spict |
| M9PD16 | 784 | S | 1.389 | 0.00838747 | Up | Chiffon, isoform D OS=Drosophila melanogaster OX=7227 GN=chif PE=4 SV=1 | chif |
| M9PDG9 | 487 | S | 1.688 | 0.001230963 | Up | Uncharacterized protein, isoform B OS=Drosophila melanogaster OX=7227 GN=Dmel\CG12773 PE=1 SV=1 | Dmel\CG12773 |
| M9PDH8 | 157 | S | 1.474 | 0.03639844 | Up | A6, isoform B OS=Drosophila melanogaster OX=7227 GN=a6 PE=4 SV=1 | a6 |
| Q9W4X4 | 1456 | S | 1.43 | 0.0008553 | Up | Phosphatidylinositol 4-kinase III alpha, isoform C OS=Drosophila melanogaster OX=7227 GN=Pi4KIIIalpha PE=1 SV=3 | Pi4KIIIalpha |
| M9PDV6 | 468 | S | 1.498 | 0.008599289 | Up | Fife, isoform B OS=Drosophila melanogaster OX=7227 GN=Fife PE=1 SV=1 | Fife |
| M9PDV6 | 242 | S | 1.663 | 0.014426839 | Up | Fife, isoform B OS=Drosophila melanogaster OX=7227 GN=Fife PE=1 SV=1 | Fife |
| M9PDV6 | 936 | S | 1.313 | 0.03214178 | Up | Fife, isoform B OS=Drosophila melanogaster OX=7227 GN=Fife PE=1 SV=1 | Fife |
| M9PDV6 | 193 | S | 1.749 | 0.006446686 | Up | Fife, isoform B OS=Drosophila melanogaster OX=7227 GN=Fife PE=1 SV=1 | Fife |

| | | | | | | | |
|---|---|---|---|---|---|---|---|
| M9PDV6 | 391 | S | 1.471 | 0.014558547 | Up | Fife, isoform B OS=Drosophila melanogaster OX=7227 GN=Fife PE=1 SV=1 | Fife |
| M9PDV6 | 399 | S | 1.609 | 0.003587811 | Up | Fife, isoform B OS=Drosophila melanogaster OX=7227 GN=Fife PE=1 SV=1 | Fife |
| M9PDV6 | 402 | S | 1.526 | 0.014188088 | Up | Fife, isoform B OS=Drosophila melanogaster OX=7227 GN=Fife PE=1 SV=1 | Fife |
| M9PDX6 | 922 | S | 2.133 | 0.000647298 | Up | Prominin-like, isoform D OS=Drosophila melanogaster OX=7227 GN=prominin-like PE=1 SV=1 | prominin-like |
| M9PE01 | 530 | S | 1.443 | 0.016854098 | Up | Ecdysone-induced protein 63E, isoform N OS=Drosophila melanogaster OX=7227 GN=Eip63E PE=4 SV=1 | Eip63E |
| M9PE30 | 121 | S | 1.73 | 1.08135E-05 | Up | Transgelin OS=Drosophila melanogaster OX=7227 GN=Chd64 PE=1 SV=1 | Chd64 |
| M9PE30 | 141 | S | 1.731 | 0.000714567 | Up | Transgelin OS=Drosophila melanogaster OX=7227 GN=Chd64 PE=1 SV=1 | Chd64 |
| M9PE74 | 1001 | S | 1.595 | 0.001828062 | Up | No circadian temperature entrainment, isoform D OS=Drosophila melanogaster OX=7227 GN=nocte PE=1 SV=1 | nocte |
| M9PEB5 | 562 | S | 1.625 | 0.005507957 | Up | Uncharacterized protein, isoform B OS=Drosophila melanogaster OX=7227 GN=AMT PE=4 SV=1 | AMT |
| M9PEB5 | 575 | S | 2.371 | 0.000109477 | Up | Uncharacterized protein, isoform B OS=Drosophila melanogaster OX=7227 GN=AMT PE=4 SV=1 | AMT |
| M9PEB5 | 578 | S | 1.415 | 0.000257935 | Up | Uncharacterized protein, isoform B OS=Drosophila melanogaster OX=7227 GN=AMT PE=4 SV=1 | AMT |
| Q7KU82 | 506 | S | 1.75 | 0.001192523 | Up | MIP20544p OS=Drosophila melanogaster OX=7227 GN=smid PE=1 SV=2 | smid |
| M9PEG1 | 685 | S | 0.751 | 0.00346656 | Down | Uncharacterized protein OS=Drosophila melanogaster OX=7227 GN=Dmel\CG44195 PE=1 SV=1 | Dmel\CG44195 |
| M9PEG8 | 204 | S | 1.336 | 0.015697553 | Up | Tantalus, isoform B OS=Drosophila melanogaster OX=7227 GN=tant PE=4 SV=1 | tant |
| M9PEK9 | 859 | S | 2.161 | 0.002266998 | Up | Smog, isoform F OS=Drosophila melanogaster OX=7227 GN=smog PE=1 SV=1 | smog |
| Q9VXU0 | 599 | S | 1.452 | 0.02975942 | Up | Adenylyl cyclase 13E, isoform A OS=Drosophila melanogaster OX=7227 GN=Ac13E PE=3 SV=1 | Ac13E |
| Q8IQD4 | 132 | S | 3.19 | 0.002158929 | Up | Anion exchange protein OS=Drosophila melanogaster OX=7227 GN=Ae2 PE=1 SV=1 | Ae2 |
| Q8IQD4 | 136 | S | 2.341 | 0.001252798 | Up | Anion exchange protein OS=Drosophila melanogaster OX=7227 GN=Ae2 PE=1 SV=1 | Ae2 |
| Q8IQD4 | 367 | S | 1.779 | 0.001901881 | Up | Anion exchange protein OS=Drosophila melanogaster OX=7227 GN=Ae2 PE=1 SV=1 | Ae2 |
| Q8IQD4 | 540 | S | 1.748 | 0.002943769 | Up | Anion exchange protein OS=Drosophila melanogaster OX=7227 GN=Ae2 PE=1 SV=1 | Ae2 |
| Q8IQD4 | 341 | S | 1.709 | 0.000415586 | Up | Anion exchange protein OS=Drosophila melanogaster OX=7227 GN=Ae2 PE=1 SV=1 | Ae2 |
| M9PFF9 | 218 | S | 1.302 | 0.04024853 | Up | Trailer hitch, isoform D OS=Drosophila melanogaster OX=7227 GN=tral PE=1 SV=1 | tral |
| Q00963 | 2268 | S | 1.661 | 0.038040949 | Up | Spectrin beta chain OS=Drosophila melanogaster OX=7227 GN=beta-Spec PE=1 SV=2 | beta-Spec |
| M9PFG7 | 145 | S | 2.036 | 0.010034284 | Up | Kinesin light chain, isoform C OS=Drosophila melanogaster OX=7227 GN=Klc PE=4 SV=1 | Klc |
| M9PFG7 | 485 | S | 1.476 | 0.00132915 | Up | Kinesin light chain, isoform C OS=Drosophila melanogaster OX=7227 GN=Klc PE=4 SV=1 | Klc |
| M9PF68 | 258 | S | 1.586 | 0.036688747 | Up | Kinesin-like protein OS=Drosophila melanogaster OX=7227 GN=Klp68D PE=3 SV=1 | Klp68D |
| M9PF94 | 126 | S | 1.432 | 0.049994457 | Up | Doublecortin-domain-containing echinoderm-microtubule-associated protein, isoform G OS=Drosophila melanogaster OX=7227 GN=DCX-EMAP PE=4 SV=1 | DCX-EMAP |
| M9PF94 | 134 | S | 1.432 | 0.049994457 | Up | Doublecortin-domain-containing echinoderm-microtubule-associated protein, isoform G OS=Drosophila melanogaster OX=7227 GN=DCX-EMAP PE=4 SV=1 | DCX-EMAP |
| M9PF94 | 137 | S | 1.432 | 0.049994457 | Up | Doublecortin-domain-containing echinoderm-microtubule-associated protein, isoform G OS=Drosophila melanogaster OX=7227 GN=DCX-EMAP PE=4 SV=1 | DCX-EMAP |
| M9PFS3 | 612 | S | 1.356 | 0.000999953 | Up | Sugar-free frosting, isoform B OS=Drosophila melanogaster OX=7227 GN=sff PE=4 SV=1 | sff |
| M9PFS3 | 613 | S | 1.374 | 0.002851826 | Up | Sugar-free frosting, isoform B OS=Drosophila melanogaster OX=7227 GN=sff PE=4 SV=1 | sff |
| M9PFS3 | 504 | S | 1.444 | 0.022869314 | Up | Sugar-free frosting, isoform B OS=Drosophila melanogaster OX=7227 GN=sff PE=4 SV=1 | sff |
| Q8SXD5 | 95 | S | 1.455 | 0.002865166 | Up | GH02216p OS=Drosophila melanogaster OX=7227 GN=vir-1 PE=1 SV=1 | vir-1 |
| Q8SXD5 | 106 | S | 1.546 | 0.001898658 | Up | GH02216p OS=Drosophila melanogaster OX=7227 GN=vir-1 PE=1 SV=1 | vir-1 |
| Q8SXD5 | 109 | S | 1.525 | 0.003836165 | Up | GH02216p OS=Drosophila melanogaster OX=7227 GN=vir-1 PE=1 SV=1 | vir-1 |
| Q8SXD5 | 113 | S | 1.549 | 0.011212408 | Up | GH02216p OS=Drosophila melanogaster OX=7227 GN=vir-1 PE=1 SV=1 | vir-1 |
| Q9Y102 | 244 | S | 2.388 | 9.85458E-06 | Up | BcDNA.GH11973 OS=Drosophila melanogaster OX=7227 GN=rgn PE=1 SV=1 | rgn |
| Q9Y102 | 402 | S | 2.434 | 0.00050345 | Up | BcDNA.GH11973 OS=Drosophila melanogaster OX=7227 GN=rgn PE=1 SV=1 | rgn |
| M9PFY1 | 168 | S | 1.43 | 0.013206942 | Up | Weckle, isoform B OS=Drosophila melanogaster OX=7227 GN=wek PE=4 SV=1 | wek |
| M9PID3 | 612 | S | 1.351 | 0.019333004 | Up | HP1 and insulator partner protein 1, isoform D OS=Drosophila melanogaster OX=7227 GN=HIPP1 PE=1 SV=1 | HIPP1 |
| M9PG50 | 471 | S | 1.664 | 0.016428033 | Up | Cyclin-dependent kinase 12, isoform C OS=Drosophila melanogaster OX=7227 GN=Cdk12 PE=4 SV=1 | Cdk12 |
| M9PG50 | 473 | S | 1.664 | 0.016428033 | Up | Cyclin-dependent kinase 12, isoform C OS=Drosophila melanogaster OX=7227 GN=Cdk12 PE=4 SV=1 | Cdk12 |
| Q7KI49 | 51 | S | 2.599 | 0.000329872 | Up | LD13050p OS=Drosophila melanogaster OX=7227 GN=M6 PE=2 SV=1 | M6 |
| Q7KI49 | 54 | S | 2.599 | 0.000329872 | Up | LD13050p OS=Drosophila melanogaster OX=7227 GN=M6 PE=2 SV=1 | M6 |
| Q7KI49 | 57 | S | 2.599 | 0.000329872 | Up | LD13050p OS=Drosophila melanogaster OX=7227 GN=M6 PE=2 SV=1 | M6 |
| M9PGC5 | 1960 | S | 0.689 | 0.001882811 | Down | Wnk kinase, isoform M OS=Drosophila melanogaster OX=7227 GN=Wnk PE=1 SV=2 | Wnk |
| M9PGC5 | 403 | S | 1.327 | 0.00337994 | Up | Wnk kinase, isoform M OS=Drosophila melanogaster OX=7227 GN=Wnk PE=1 SV=2 | Wnk |
| M9PGE1 | 240 | S | 1.33 | 0.033406093 | Up | Dpr-interacting protein alpha, isoform B OS=Drosophila melanogaster OX=7227 GN=DIP-alpha PE=4 SV=1 | DIP-alpha |
| M9PGF7 | 177 | S | 1.477 | 0.002837331 | Up | Tenascin major, isoform E OS=Drosophila melanogaster OX=7227 GN=Ten-m PE=1 SV=1 | Ten-m |
| M9PGI7 | 272 | S | 1.335 | 0.00912381 | Up | Uncharacterized protein, isoform B OS=Drosophila melanogaster OX=7227 GN=PRP4 PE=1 SV=1 | PRP4 |
| M9PHA0 | 870 | S | 1.663 | 0.020104474 | Up | Bifocal, isoform F OS=Drosophila melanogaster OX=7227 GN=bif PE=1 SV=1 | bif |
| M9PHG0 | 988 | S | 1.312 | 0.03895011 | Up | Rho GTPase activating protein p190, isoform D OS=Drosophila melanogaster OX=7227 GN=RhoGAPp190 PE=1 SV=1 | RhoGAPp190 |
| M9PHI0 | 718 | S | 1.845 | 0.005481834 | Up | Hyperkinetic, isoform M OS=Drosophila melanogaster OX=7227 GN=Hk PE=4 SV=1 | Hk |
| M9PHI0 | 763 | S | 1.431 | 0.005692585 | Up | Hyperkinetic, isoform M OS=Drosophila melanogaster OX=7227 GN=Hk PE=4 SV=1 | Hk |
| M9PHJ0 | 2 | S | 0.54 | 0.044527788 | Down | Upheld, isoform P OS=Drosophila melanogaster OX=7227 GN=up PE=1 SV=1 | up |
| M9PHJ0 | 9 | S | 0.548 | 0.044076149 | Down | Upheld, isoform P OS=Drosophila melanogaster OX=7227 GN=up PE=1 SV=1 | up |
| M9PHL0 | 1916 | S | 2.067 | 0.0027888 | Up | Enhancer of yellow 3, isoform B OS=Drosophila melanogaster OX=7227 GN=e(y)3 PE=4 SV=1 | e(y)3 |
| M9PHL0 | 419 | S | 1.382 | 0.018735457 | Up | Enhancer of yellow 3, isoform B OS=Drosophila melanogaster OX=7227 GN=e(y)3 PE=4 SV=1 | e(y)3 |
| M9PHM0 | 493 | S | 0.711 | 0.0213889 | Down | Uncharacterized protein, isoform B OS=Drosophila melanogaster OX=7227 GN=Dmel\CG12531 PE=4 SV=1 | Dmel\CG12531 |
| M9PI37 | 934 | S | 1.569 | 0.000696177 | Up | Sodium chloride cotransporter 69, isoform E OS=Drosophila melanogaster OX=7227 GN=Ncc69 PE=1 SV=1 | Ncc69 |

| | | | | | | | |
|---|---|---|---|---|---|---|---|
| Q9VU80 | 352 | S | 1.592 | 0.016751026 | Up | FI14633p OS=Drosophila melanogaster OX=7227 GN=Dmel\CG10089 PE=2 SV=2 | Dmel\CG10089 |
| M9PJC7 | 1971 | S | 1.501 | 0.01800164 | Up | Female sterile (1) homeotic, isoform G OS=Drosophila melanogaster OX=7227 GN=fs(1)h PE=1 SV=1 | fs(1)h |
| M9PJE4 | 63 | S | 0.709 | 0.008491951 | Down | LIM homeobox 1, isoform B OS=Drosophila melanogaster OX=7227 GN=Lim1 PE=4 SV=1 | Lim1 |
| M9PJM8 | 379 | S | 1.337 | 0.009732439 | Up | Retinal degeneration B, isoform I OS=Drosophila melanogaster OX=7227 GN=rdgB PE=1 SV=1 | rdgB |
| N0D8I3 | 156 | S | 1.393 | 0.049124911 | Up | Transcription elongation factor spt6 OS=Drosophila melanogaster OX=7227 GN=Spt6 PE=1 SV=1 | Spt6 |
| O02193 | 154 | S | 1.301 | 0.002287398 | Up | Males-absent on the first protein OS=Drosophila melanogaster OX=7227 GN=mof PE=1 SV=1 | mof |
| O18332 | 187 | S | 1.32 | 0.010436589 | Up | FI01544p OS=Drosophila melanogaster OX=7227 GN=Rab1 PE=1 SV=1 | Rab1 |
| O44424 | 409 | S | 1.331 | 0.011722574 | Up | Splicing factor ESS-2 homolog OS=Drosophila melanogaster OX=7227 GN=Es2 PE=1 SV=2 | Es2 |
| O44424 | 411 | S | 1.331 | 0.011722574 | Up | Splicing factor ESS-2 homolog OS=Drosophila melanogaster OX=7227 GN=Es2 PE=1 SV=2 | Es2 |
| O61604 | 12 | S | 1.43 | 0.016179051 | Up | Fimbrin OS=Drosophila melanogaster OX=7227 GN=Fim PE=1 SV=1 | Fim |
| O62530 | 231 | S | 1.943 | 0.004149857 | Up | Adaptor protein complex 2, mu subunit, isoform A OS=Drosophila melanogaster OX=7227 GN=AP-2mu PE=1 SV=1 | AP-2mu |
| O76742 | 34 | S | 1.39 | 0.000274425 | Up | CG5915 protein OS=Drosophila melanogaster OX=7227 GN=Rab7 PE=1 SV=1 | Rab7 |
| O76874 | 264 | S | 2.099 | 0.000337712 | Up | SD17974p OS=Drosophila melanogaster OX=7227 GN=EG:125H10.1 PE=1 SV=2 | EG:125H10.1 |
| Q2PE37 | 15 | S | 1.42 | 0.015326569 | Up | Pickel, isoform D OS=Drosophila melanogaster OX=7227 GN=pck PE=4 SV=1 | pck |
| O77062 | 456 | S | 1.885 | 0.000322366 | Up | Amino acid transporter OS=Drosophila melanogaster OX=7227 GN=Eaat1 PE=1 SV=1 | Eaat1 |
| O97125 | 509 | S | 4.748 | 8.17858E-06 | Up | Heat shock protein 68 OS=Drosophila melanogaster OX=7227 GN=Hsp68 PE=1 SV=1 | Hsp68 |
| O97182 | 311 | S | 1.582 | 0.002949611 | Up | Actin-related protein 2/3 complex subunit OS=Drosophila melanogaster OX=7227 GN=Arpc1 PE=1 SV=1 | Arpc1 |
| P00334 | 165 | S | 0.706 | 0.000950023 | Down | Alcohol dehydrogenase OS=Drosophila melanogaster OX=7227 GN=Adh PE=1 SV=2 | Adh |
| X2JD55 | 185 | S | 7.982 | 0.001300289 | Up | Yolk protein 1, isoform B OS=Drosophila melanogaster OX=7227 GN=Yp1 PE=1 SV=1 | Yp1 |
| X2JD55 | 186 | S | 5.6 | 0.001290446 | Up | Yolk protein 1, isoform B OS=Drosophila melanogaster OX=7227 GN=Yp1 PE=1 SV=1 | Yp1 |
| X2JD55 | 191 | S | 5.6 | 0.001290446 | Up | Yolk protein 1, isoform B OS=Drosophila melanogaster OX=7227 GN=Yp1 PE=1 SV=1 | Yp1 |
| X2JD55 | 175 | S | 6.133 | 0.005898108 | Up | Yolk protein 1, isoform B OS=Drosophila melanogaster OX=7227 GN=Yp1 PE=1 SV=1 | Yp1 |
| X2JB25 | 33 | S | 3.839 | 0.0223691 | Up | Yolk protein 2, isoform B OS=Drosophila melanogaster OX=7227 GN=Yp2 PE=1 SV=1 | Yp2 |
| P06002 | 357 | S | 0.544 | 0.040221537 | Down | Opsin Rh1 OS=Drosophila melanogaster OX=7227 GN=ninaE PE=1 SV=1 | ninaE |
| P06002 | 358 | S | 0.507 | 0.017510701 | Down | Opsin Rh1 OS=Drosophila melanogaster OX=7227 GN=ninaE PE=1 SV=1 | ninaE |
| P07668 | 74 | S | 2.911 | 0.015213109 | Up | Choline O-acetyltransferase OS=Drosophila melanogaster OX=7227 GN=ChAT PE=1 SV=3 | ChAT |
| P07668 | 28 | S | 1.456 | 0.031676413 | Up | Choline O-acetyltransferase OS=Drosophila melanogaster OX=7227 GN=ChAT PE=1 SV=3 | ChAT |
| P08255 | 363 | S | 1.515 | 0.04039281 | Up | Opsin Rh4 OS=Drosophila melanogaster OX=7227 GN=Rh4 PE=1 SV=2 | Rh4 |
| P10676 | 1410 | S | 1.515 | 0.017658966 | Up | Neither inactivation nor afterpotential protein C OS=Drosophila melanogaster OX=7227 GN=ninaC PE=1 SV=2 | ninaC |
| P10676 | 1397 | S | 1.513 | 0.001635815 | Up | Neither inactivation nor afterpotential protein C OS=Drosophila melanogaster OX=7227 GN=ninaC PE=1 SV=2 | ninaC |
| X2J8L3 | 4 | S | 0.647 | 0.003513435 | Down | Adenine phosphoribosyltransferase, isoform C OS=Drosophila melanogaster OX=7227 GN=Aprt PE=3 SV=1 | Aprt |
| P13496 | 375 | S | 0.35 | 0.000135105 | Down | Dynactin subunit 1 OS=Drosophila melanogaster OX=7227 GN=DCTN1-p150 PE=1 SV=2 | DCTN1-p150 |
| P13496 | 568 | S | 1.568 | 0.000217956 | Up | Dynactin subunit 1 OS=Drosophila melanogaster OX=7227 GN=DCTN1-p150 PE=1 SV=2 | DCTN1-p150 |
| P13496 | 573 | S | 1.303 | 0.00107502 | Up | Dynactin subunit 1 OS=Drosophila melanogaster OX=7227 GN=DCTN1-p150 PE=1 SV=2 | DCTN1-p150 |
| P13496 | 85 | S | 1.338 | 0.01807608 | Up | Dynactin subunit 1 OS=Drosophila melanogaster OX=7227 GN=DCTN1-p150 PE=1 SV=2 | DCTN1-p150 |
| P15348 | 1385 | S | 0.747 | 0.00242086 | Down | DNA topoisomerase 2 OS=Drosophila melanogaster OX=7227 GN=Top2 PE=1 SV=1 | Top2 |
| P15348 | 1392 | S | 0.747 | 0.00242086 | Down | DNA topoisomerase 2 OS=Drosophila melanogaster OX=7227 GN=Top2 PE=1 SV=1 | Top2 |
| P19107 | 186 | S | 0.718 | 0.008671611 | Down | Phosrestin-1 OS=Drosophila melanogaster OX=7227 GN=Arr2 PE=1 SV=2 | Arr2 |
| P19334 | 867 | S | 1.349 | 0.002692878 | Up | Transient receptor potential protein OS=Drosophila melanogaster OX=7227 GN=trp PE=1 SV=3 | trp |
| P20240 | 50 | S | 1.321 | 0.01823105 | Up | Otefin OS=Drosophila melanogaster OX=7227 GN=Ote PE=1 SV=2 | Ote |
| P20240 | 54 | S | 1.459 | 0.040010997 | Up | Otefin OS=Drosophila melanogaster OX=7227 GN=Ote PE=1 SV=2 | Ote |
| X2JJG8 | 59 | S | 2.987 | 1.68646E-05 | Up | Glutamine synthetase OS=Drosophila melanogaster OX=7227 GN=Gs2 PE=1 SV=1 | Gs2 |
| X2JJG8 | 349 | S | 1.367 | 0.000745904 | Up | Glutamine synthetase OS=Drosophila melanogaster OX=7227 GN=Gs2 PE=1 SV=1 | Gs2 |
| X2J979 | 21 | S | 0.504 | 0.000433429 | Down | Synaptotagmin 1, isoform H OS=Drosophila melanogaster OX=7227 GN=Syt1 PE=4 SV=1 | Syt1 |
| P22815 | 883 | S | 1.329 | 0.015095202 | Up | Protein bride of sevenless OS=Drosophila melanogaster OX=7227 GN=boss PE=2 SV=3 | boss |
| P23128 | 458 | S | 2.102 | 0.002164889 | Up | ATP-dependent RNA helicase me31b OS=Drosophila melanogaster OX=7227 GN=me31B PE=1 SV=3 | me31B |
| Q8IMF5 | 712 | S | 1.413 | 0.000326375 | Up | Microtubule-associated protein 205, isoform B OS=Drosophila melanogaster OX=7227 GN=Map205 PE=1 SV=1 | Map205 |
| P29746 | 97 | S | 1.497 | 0.013380509 | Up | Protein bangles and beads OS=Drosophila melanogaster OX=7227 GN=bnb PE=1 SV=1 | bnb |
| P29746 | 192 | S | 1.968 | 0.014437196 | Up | Protein bangles and beads OS=Drosophila melanogaster OX=7227 GN=bnb PE=1 SV=1 | bnb |
| P29746 | 302 | S | 1.664 | 0.002413832 | Up | Protein bangles and beads OS=Drosophila melanogaster OX=7227 GN=bnb PE=1 SV=1 | bnb |
| P34082 | 814 | S | 1.404 | 0.029732276 | Up | Fasciclin-2 OS=Drosophila melanogaster OX=7227 GN=Fas2 PE=1 SV=1 | Fas2 |
| P35381 | 47 | S | 0.684 | 0.001933306 | Down | ATP synthase subunit alpha, mitochondrial OS=Drosophila melanogaster OX=7227 GN=blw PE=1 SV=2 | blw |
| P35992 | 1240 | S | 1.688 | 0.000485563 | Up | Tyrosine-protein phosphatase 10D OS=Drosophila melanogaster OX=7227 GN=Ptp10D PE=1 SV=4 | Ptp10D |
| P35992 | 1244 | S | 1.688 | 0.000485563 | Up | Tyrosine-protein phosphatase 10D OS=Drosophila melanogaster OX=7227 GN=Ptp10D PE=1 SV=4 | Ptp10D |
| P45594 | 8 | S | 2.069 | 0.040107062 | Up | Cofilin/actin-depolymerizing factor homolog OS=Drosophila melanogaster OX=7227 GN=tsr PE=1 SV=1 | tsr |
| P48596 | 43 | S | 0.713 | 0.02362953 | Down | GTP cyclohydrolase 1 OS=Drosophila melanogaster OX=7227 GN=Pu PE=1 SV=3 | Pu |
| P61851 | 101 | S | 1.312 | 0.010635515 | Up | Superoxide dismutase [Cu-Zn] OS=Drosophila melanogaster OX=7227 GN=Sod1 PE=1 SV=2 | Sod1 |
| P91926 | 634 | S | 1.356 | 0.000960797 | Up | AP-2 complex subunit alpha OS=Drosophila melanogaster OX=7227 GN=AP-2alpha PE=1 SV=1 | AP-2alpha |
| Q00748 | 685 | S | 1.886 | 0.001360091 | Up | Multidrug resistance protein homolog 65 OS=Drosophila melanogaster OX=7227 GN=Mdr65 PE=1 SV=2 | Mdr65 |
| Q00963 | 2134 | S | 1.517 | 0.006823476 | Up | Spectrin beta chain OS=Drosophila melanogaster OX=7227 GN=beta-Spec PE=1 SV=2 | beta-Spec |
| Q00963 | 2138 | S | 1.476 | 0.001985882 | Up | Spectrin beta chain OS=Drosophila melanogaster OX=7227 GN=beta-Spec PE=1 SV=2 | beta-Spec |
| Q9VWL9 | 211 | S | 2.023 | 0.000398097 | Up | LD25711p OS=Drosophila melanogaster OX=7227 GN=RhoGAP18B | RhoGAP18B |

| | | | | | | | |
|---|---|---|---|---|---|---|---|
| | | | | | | PE=2 SV=2 | |
| Q03427 | 441 | S | 1.406 | 0.001717801 | Up | Lamin-C OS=Drosophila melanogaster OX=7227 GN=LamC PE=1 SV=2 | LamC |
| Q07407 | 46 | S | 2.177 | 0.000498936 | Up | Fibroblast growth factor receptor homolog 1 OS=Drosophila melanogaster OX=7227 GN=htl PE=1 SV=3 | htl |
| Q0E8P5 | 145 | S | 1.647 | 0.000554278 | Up | FI05614p OS=Drosophila melanogaster OX=7227 GN=Dmel\CG5758 PE=2 SV=1 | Dmel\CG5758 |
| Q0E8U7 | 1168 | S | 1.308 | 0.036979749 | Up | WD repeat domain 62, isoform C OS=Drosophila melanogaster OX=7227 GN=Wdr62 PE=4 SV=1 | Wdr62 |
| Q0E8U7 | 1380 | S | 1.402 | 0.004343768 | Up | WD repeat domain 62, isoform C OS=Drosophila melanogaster OX=7227 GN=Wdr62 PE=4 SV=1 | Wdr62 |
| Q7KN97 | 495 | S | 0.702 | 0.002925625 | Down | Pyruvate carboxylase OS=Drosophila melanogaster OX=7227 GN=PCB PE=1 SV=1 | PCB |
| Q0KHR5 | 32 | S | 1.313 | 4.83743E-05 | Up | Rho GTPase activating protein at 15B, isoform B OS=Drosophila melanogaster OX=7227 GN=RhoGAP15B PE=1 SV=1 | RhoGAP15B |
| Q0KHZ9 | 569 | S | 1.628 | 0.000317618 | Up | Calnexin 99A, isoform E OS=Drosophila melanogaster OX=7227 GN=Cnx99A PE=1 SV=2 | Cnx99A |
| Q0KID3 | 420 | S | 1.331 | 0.007643624 | Up | Uncharacterized protein, isoform E OS=Drosophila melanogaster OX=7227 GN=CG9818 PE=3 SV=2 | CG9818 |
| Q0KID3 | 429 | S | 1.331 | 0.007643624 | Up | Uncharacterized protein, isoform E OS=Drosophila melanogaster OX=7227 GN=CG9818 PE=3 SV=2 | CG9818 |
| Q24050 | 192 | S | 1.358 | 0.000989108 | Up | Anon-i1 protein OS=Drosophila melanogaster OX=7227 GN=Dmel\CG2034 PE=1 SV=1 | Dmel\CG2034 |
| Q24050 | 197 | S | 1.358 | 0.000989108 | Up | Anon-i1 protein OS=Drosophila melanogaster OX=7227 GN=Dmel\CG2034 PE=1 SV=1 | Dmel\CG2034 |
| Q24238 | 34 | S | 2.067 | 0.000696489 | Up | Alkaline phosphatase 4 OS=Drosophila melanogaster OX=7227 GN=Alp4 PE=2 SV=3 | Alp4 |
| Q24400 | 88 | S | 1.588 | 0.001493631 | Up | Muscle LIM protein Mlp84B OS=Drosophila melanogaster OX=7227 GN=Mlp84B PE=1 SV=1 | Mlp84B |
| Q24418 | 954 | S | 1.432 | 0.021880065 | Up | Glutamate [NMDA] receptor subunit 1 OS=Drosophila melanogaster OX=7227 GN=Nmdar1 PE=1 SV=1 | Nmdar1 |
| Q24472 | 791 | S | 1.426 | 0.003234657 | Up | Retinoblastoma family protein OS=Drosophila melanogaster OX=7227 GN=Rbf PE=1 SV=2 | Rbf |
| Q24478 | 233 | S | 2.04 | 0.00142509 | Up | Centrosome-associated zinc finger protein CP190 OS=Drosophila melanogaster OX=7227 GN=Cp190 PE=1 SV=2 | Cp190 |
| Q24478 | 197 | S | 1.332 | 0.038067825 | Up | Centrosome-associated zinc finger protein CP190 OS=Drosophila melanogaster OX=7227 GN=Cp190 PE=1 SV=2 | Cp190 |
| X2J847 | 289 | S | 1.413 | 0.02222688 | Up | Bunched, isoform P OS=Drosophila melanogaster OX=7227 GN=bun PE=4 SV=1 | bun |
| Q24546 | 6 | S | 1.72 | 0.004662714 | Up | Synapsin OS=Drosophila melanogaster OX=7227 GN=Syn PE=1 SV=2 | Syn |
| Q24546 | 7 | S | 2.976 | 0.002316774 | Up | Synapsin OS=Drosophila melanogaster OX=7227 GN=Syn PE=1 SV=2 | Syn |
| Q24546 | 11 | S | 1.414 | 0.004596771 | Up | Synapsin OS=Drosophila melanogaster OX=7227 GN=Syn PE=1 SV=2 | Syn |
| Q24595 | 859 | S | 1.346 | 0.000236908 | Up | DNA repair protein complementing XP-C cells homolog OS=Drosophila melanogaster OX=7227 GN=Xpc PE=1 SV=2 | Xpc |
| Q24595 | 533 | S | 1.5 | 9.9631E-05 | Up | DNA repair protein complementing XP-C cells homolog OS=Drosophila melanogaster OX=7227 GN=Xpc PE=1 SV=2 | Xpc |
| Q24595 | 537 | S | 1.5 | 9.9631E-05 | Up | DNA repair protein complementing XP-C cells homolog OS=Drosophila melanogaster OX=7227 GN=Xpc PE=1 SV=2 | Xpc |
| X2JCI8 | 114 | S | 1.427 | 0.030502708 | Up | Msr-110, isoform D OS=Drosophila melanogaster OX=7227 GN=Msr-110 PE=1 SV=1 | Msr-110 |
| X2JCI8 | 118 | S | 1.498 | 0.033466961 | Up | Msr-110, isoform D OS=Drosophila melanogaster OX=7227 GN=Msr-110 PE=1 SV=1 | Msr-110 |
| X2JCI8 | 141 | S | 4.941 | 5.52591E-06 | Up | Msr-110, isoform D OS=Drosophila melanogaster OX=7227 GN=Msr-110 PE=1 SV=1 | Msr-110 |
| X2JCI8 | 145 | S | 4.46 | 7.01513E-06 | Up | Msr-110, isoform D OS=Drosophila melanogaster OX=7227 GN=Msr-110 PE=1 SV=1 | Msr-110 |
| X2JCI8 | 337 | S | 2.67 | 5.56496E-06 | Up | Msr-110, isoform D OS=Drosophila melanogaster OX=7227 GN=Msr-110 PE=1 SV=1 | Msr-110 |
| X2JCI8 | 338 | S | 3.258 | 7.23708E-06 | Up | Msr-110, isoform D OS=Drosophila melanogaster OX=7227 GN=Msr-110 PE=1 SV=1 | Msr-110 |
| X2JCI8 | 250 | S | 2.363 | 0.002571274 | Up | Msr-110, isoform D OS=Drosophila melanogaster OX=7227 GN=Msr-110 PE=1 SV=1 | Msr-110 |
| Q29QE1 | 412 | S | 1.444 | 0.026916034 | Up | GH15984p OS=Drosophila melanogaster OX=7227 GN=RhoGAP1A PE=1 SV=1 | RhoGAP1A |
| Q29R16 | 49 | S | 2.115 | 0.000398597 | Up | LP13067p OS=Drosophila melanogaster OX=7227 GN=ome PE=1 SV=1 | ome |
| Q2PDQ9 | 78 | S | 1.482 | 0.000437854 | Up | FI23737p1 OS=Drosophila melanogaster OX=7227 GN=Dmel\CG8927 PE=1 SV=1 | Dmel\CG8927 |
| Q494G8 | 223 | S | 1.506 | 0.002469861 | Up | F-box and leucine-rich repeat protein 6, isoform A OS=Drosophila melanogaster OX=7227 GN=Fbl6 PE=1 SV=1 | Fbl6 |
| Q56JH9 | 8 | S | 1.809 | 0.000729831 | Up | Hyperpolarization-activated ion channel variant DMIH-A2B1C1 OS=Drosophila melanogaster OX=7227 GN=Ih PE=2 SV=1 | Ih |
| Q58CJ5 | 299 | S | 1.41 | 0.039721523 | Up | GH01093p OS=Drosophila melanogaster OX=7227 GN=sals PE=1 SV=1 | sals |
| Q59DP9 | 1129 | S | 1.44 | 0.029253666 | Up | Calcium-transporting ATPase OS=Drosophila melanogaster OX=7227 GN=PMCA PE=1 SV=3 | PMCA |
| Q59DP9 | 1132 | S | 1.44 | 0.029253666 | Up | Calcium-transporting ATPase OS=Drosophila melanogaster OX=7227 GN=PMCA PE=1 SV=3 | PMCA |
| Q59DP9 | 1135 | S | 1.44 | 0.029253666 | Up | Calcium-transporting ATPase OS=Drosophila melanogaster OX=7227 GN=PMCA PE=1 SV=3 | PMCA |
| Q5LJZ2 | 589 | S | 1.357 | 0.014861953 | Up | Histone-lysine N-methyltransferase SETD1 OS=Drosophila melanogaster OX=7227 GN=Set1 PE=1 SV=1 | Set1 |
| Q7JV09 | 972 | S | 1.766 | 0.033207187 | Up | GH28348p OS=Drosophila melanogaster OX=7227 GN=jp PE=2 SV=1 | jp |
| Q9VZU9 | 107 | S | 1.481 | 0.009257619 | Up | Uncharacterized protein, isoform A OS=Drosophila melanogaster OX=7227 GN=Dmel\CG11537 PE=4 SV=1 | Dmel\CG11537 |
| Q9VZU9 | 676 | S | 1.545 | 0.00320395 | Up | Uncharacterized protein, isoform A OS=Drosophila melanogaster OX=7227 GN=Dmel\CG11537 PE=4 SV=1 | Dmel\CG11537 |
| Q6NP69 | 697 | S | 2.109 | 0.001926322 | Up | GST-containing FLYWCH zinc-finger protein OS=Drosophila melanogaster OX=7227 GN=gfzf PE=1 SV=1 | gfzf |
| Q6NP91 | 704 | S | 1.929 | 0.000978955 | Up | RE44586p OS=Drosophila melanogaster OX=7227 GN=Tmem63 PE=1 SV=1 | Tmem63 |
| Q7JQX9 | 419 | S | 1.906 | 0.035911143 | Up | FI14001p1 OS=Drosophila melanogaster OX=7227 GN=SNF4Agamma PE=2 SV=1 | SNF4Agamma |
| Q7JR73 | 289 | S | 1.613 | 0.007526548 | Up | LD20239p OS=Drosophila melanogaster OX=7227 GN=Rcd6 PE=2 SV=1 | Rcd6 |
| Q86P32 | 106 | S | 1.312 | 0.010438583 | Up | RE35194p OS=Drosophila melanogaster OX=7227 GN=Syx6 PE=2 SV=1 | Syx6 |
| Q86P32 | 120 | S | 1.312 | 0.010438583 | Up | RE35194p OS=Drosophila melanogaster OX=7227 GN=Syx6 PE=2 SV=1 | Syx6 |
| Q7JV09 | 806 | S | 1.575 | 0.039304093 | Up | GH28348p OS=Drosophila melanogaster OX=7227 GN=jp PE=2 SV=1 | jp |
| Q7JV09 | 930 | S | 1.791 | 0.033035548 | Up | GH28348p OS=Drosophila melanogaster OX=7227 GN=jp PE=2 SV=1 | jp |
| Q7JVF1 | 38 | S | 2.1 | 0.041938802 | Up | LP02169p OS=Drosophila melanogaster OX=7227 GN=Socs44A PE=2 SV=1 | Socs44A |
| Q7JVP4 | 1018 | S | 1.393 | 0.011145136 | Up | Bromodomain-containing protein, 140kD, isoform A OS=Drosophila melanogaster OX=7227 GN=Br140 PE=1 SV=1 | Br140 |
| Q7JZ25 | 14 | S | 1.656 | 5.84256E-05 | Up | Chloride channel protein OS=Drosophila melanogaster OX=7227 GN=ClC-b PE=1 SV=1 | ClC-b |
| Q7JZ25 | 15 | S | 1.558 | 3.4217E-05 | Up | Chloride channel protein OS=Drosophila melanogaster OX=7227 GN=ClC-b PE=1 SV=1 | ClC-b |
| Q7K0L8 | 358 | S | 1.731 | 0.00037481 | Up | FLICE-associated huge protein, isoform A OS=Drosophila melanogaster OX=7227 GN=FLASH PE=1 SV=1 | FLASH |
| Q7K0S5 | 301 | S | 0.76 | 0.000467157 | Down | GDI interacting protein 3, isoform B OS=Drosophila melanogaster OX=7227 GN=Gint3 PE=1 SV=1 | Gint3 |
| Q7K180 | 241 | S | 1.372 | 0.031661911 | Up | LD02709p OS=Drosophila melanogaster OX=7227 GN=Map60 PE=1 SV=1 | Map60 |
| Q7K1D7 | 14 | S | 1.479 | 0.00077952 | Up | GH15861p OS=Drosophila melanogaster OX=7227 GN=Dmel\CG1358 PE=1 SV=1 | Dmel\CG1358 |
| Q7K1D7 | 23 | S | 1.732 | 0.000310913 | Up | GH15861p OS=Drosophila melanogaster OX=7227 GN=Dmel\CG1358 PE=1 SV=1 | Dmel\CG1358 |
| Q7K1L4 | 422 | S | 1.691 | 0.006687921 | Up | SD10469p OS=Drosophila melanogaster OX=7227 GN=Dmel\CG8468 PE=1 SV=1 | Dmel\CG8468 |

| | | | | | | | |
|---|---|---|---|---|---|---|---|
| Q7K204 | 544 | S | 1.427 | 0.001228012 | Up | Barricade, isoform A OS=Drosophila melanogaster OX=7227 GN=barc PE=1 SV=1 | barc |
| Q7K3H0 | 112 | S | 3.079 | 3.89172E-05 | Up | LD28067p OS=Drosophila melanogaster OX=7227 GN=Pi3K68D PE=2 SV=1 | Pi3K68D |
| Q7K3H0 | 102 | S | 1.475 | 0.004016703 | Up | LD28067p OS=Drosophila melanogaster OX=7227 GN=Pi3K68D PE=2 SV=1 | Pi3K68D |
| Q7K3Y9 | 566 | S | 2.319 | 2.52311E-06 | Up | GH02025p OS=Drosophila melanogaster OX=7227 GN=Dmel\CG17739 PE=2 SV=1 | Dmel\CG17739 |
| Q7K3Z3 | 117 | S | 0.747 | 0.024196316 | Down | GH01724p OS=Drosophila melanogaster OX=7227 GN=p47 PE=1 SV=1 | p47 |
| Q7K3Z3 | 119 | S | 0.756 | 0.019378681 | Down | GH01724p OS=Drosophila melanogaster OX=7227 GN=p47 PE=1 SV=1 | p47 |
| Q7K3Z3 | 114 | S | 1.75 | 0.004339736 | Up | GH01724p OS=Drosophila melanogaster OX=7227 GN=p47 PE=1 SV=1 | p47 |
| Q7K4R2 | 177 | S | 1.322 | 0.00226224 | Up | LD26477p OS=Drosophila melanogaster OX=7227 GN=Dmel\CG11504 PE=2 SV=1 | Dmel\CG11504 |
| Q7K7G0 | 241 | S | 1.606 | 0.000150132 | Up | Coatomer subunit delta OS=Drosophila melanogaster OX=7227 GN=deltaCOP PE=1 SV=1 | deltaCOP |
| Q7K9H6 | 636 | S | 1.772 | 0.000771501 | Up | Zinc finger FYVE domain-containing protein OS=Drosophila melanogaster OX=7227 GN=Sara PE=1 SV=1 | Sara |
| Q7KK54 | 747 | S | 1.774 | 0.000314901 | Up | MIP07328p OS=Drosophila melanogaster OX=7227 GN=Sema1b PE=1 SV=1 | Sema1b |
| Q7KLE5 | 340 | S | 2.06 | 0.000333445 | Up | Amphiphysin OS=Drosophila melanogaster OX=7227 GN=Amph PE=1 SV=1 | Amph |
| Q7KLI1 | 1409 | S | 1.489 | 0.006492511 | Up | Apaf-1 related killer DARK OS=Drosophila melanogaster OX=7227 GN=Dark PE=1 SV=1 | Dark |
| Q7KLI1 | 1411 | S | 1.489 | 0.006492511 | Up | Apaf-1 related killer DARK OS=Drosophila melanogaster OX=7227 GN=Dark PE=1 SV=1 | Dark |
| Q7KRT4 | 378 | S | 0.7 | 0.000366164 | Down | GH07253p OS=Drosophila melanogaster OX=7227 GN=stops PE=2 SV=1 | stops |
| Q7KRT4 | 382 | S | 0.666 | 0.001571025 | Down | GH07253p OS=Drosophila melanogaster OX=7227 GN=stops PE=2 SV=1 | stops |
| Q7KRW8 | 252 | S | 1.495 | 0.001281337 | Up | Pre-mRNA-processing factor 39 OS=Drosophila melanogaster OX=7227 GN=CG1646 PE=1 SV=1 | CG1646 |
| Q7KRW8 | 151 | S | 1.477 | 0.001395484 | Up | Pre-mRNA-processing factor 39 OS=Drosophila melanogaster OX=7227 GN=CG1646 PE=1 SV=1 | CG1646 |
| Q7KRW8 | 25 | S | 1.415 | 0.008230522 | Up | Pre-mRNA-processing factor 39 OS=Drosophila melanogaster OX=7227 GN=CG1646 PE=1 SV=1 | CG1646 |
| Q9VBI4 | 664 | S | 1.802 | 0.041408442 | Up | Exocyst 84, isoform A OS=Drosophila melanogaster OX=7227 GN=Exo84 PE=1 SV=1 | Exo84 |
| Q7KSE4 | 81 | S | 1.482 | 0.001216076 | Up | GH05443p OS=Drosophila melanogaster OX=7227 GN=repo PE=2 SV=1 | repo |
| Q7KSP6 | 1736 | S | 1.454 | 0.000574542 | Up | SET domain binding factor, isoform B OS=Drosophila melanogaster OX=7227 GN=Sbf PE=1 SV=1 | Sbf |
| X2JE45 | 486 | S | 1.309 | 0.006178748 | Up | Uncharacterized protein, isoform C OS=Drosophila melanogaster OX=7227 GN=Dmel\CG33090 PE=4 SV=1 | Dmel\CG33090 |
| Q7KTG4 | 383 | S | 1.343 | 0.011572065 | Up | Organic anion transporting polypeptide 30B, isoform D OS=Drosophila melanogaster OX=7227 GN=Oatp30B PE=4 SV=1 | Oatp30B |
| Q7KTG4 | 1034 | S | 1.391 | 0.001142162 | Up | Organic anion transporting polypeptide 30B, isoform D OS=Drosophila melanogaster OX=7227 GN=Oatp30B PE=4 SV=1 | Oatp30B |
| Q7KTG4 | 1038 | S | 1.391 | 0.001142162 | Up | Organic anion transporting polypeptide 30B, isoform D OS=Drosophila melanogaster OX=7227 GN=Oatp30B PE=4 SV=1 | Oatp30B |
| Q7KU45 | 977 | S | 1.353 | 0.029226859 | Up | Lethal (2) k05819, isoform C OS=Drosophila melanogaster OX=7227 GN=l(2)k05819 PE=4 SV=1 | l(2)k05819 |
| Q7KU45 | 980 | S | 1.353 | 0.029226859 | Up | Lethal (2) k05819, isoform C OS=Drosophila melanogaster OX=7227 GN=l(2)k05819 PE=4 SV=1 | l(2)k05819 |
| Q7KUW6 | 1020 | S | 1.973 | 0.003354064 | Up | Chascon, isoform B OS=Drosophila melanogaster OX=7227 GN=chas PE=4 SV=1 | chas |
| Q9VXF8 | 20 | S | 1.633 | 0.010970772 | Up | Eukaryotic translation initiation factor 4H1, isoform A OS=Drosophila melanogaster OX=7227 GN=eIF4H1 PE=1 SV=1 | eIF4H1 |
| Q9VXF8 | 186 | S | 1.413 | 0.011097208 | Up | Eukaryotic translation initiation factor 4H1, isoform A OS=Drosophila melanogaster OX=7227 GN=eIF4H1 PE=1 SV=1 | eIF4H1 |
| Q7KV34 | 208 | S | 0.642 | 0.003053782 | Down | CHK domain-containing protein OS=Drosophila melanogaster OX=7227 GN=Dmel\CG1561 PE=1 SV=1 | Dmel\CG1561 |
| Q7KV69 | 3961 | S | 1.517 | 0.000693669 | Up | Karst, isoform B OS=Drosophila melanogaster OX=7227 GN=kst PE=1 SV=1 | kst |
| Q7KVX5 | 4 | S | 1.4 | 0.013894922 | Up | VAMP-associated protein 33kDa, isoform A OS=Drosophila melanogaster OX=7227 GN=Vap33 PE=1 SV=1 | Vap33 |
| Q7PLL3 | 262 | S | 1.769 | 0.000696484 | Up | Eukaryotic initiation factor 4B OS=Drosophila melanogaster OX=7227 GN=eIF4B PE=1 SV=1 | eIF4B |
| Q7PLL3 | 233 | S | 1.614 | 0.005786517 | Up | Eukaryotic initiation factor 4B OS=Drosophila melanogaster OX=7227 GN=eIF4B PE=1 SV=1 | eIF4B |
| Q86B74 | 272 | S | 1.538 | 0.012453949 | Up | WASp, isoform C OS=Drosophila melanogaster OX=7227 GN=WASp PE=4 SV=1 | WASp |
| Q86B74 | 275 | S | 1.538 | 0.012453949 | Up | WASp, isoform C OS=Drosophila melanogaster OX=7227 GN=WASp PE=4 SV=1 | WASp |
| Q9VLL3 | 135 | S | 1.521 | 0.009716793 | Up | A-kinase anchor protein 200 OS=Drosophila melanogaster OX=7227 GN=Akap200 PE=1 SV=3 | Akap200 |
| Q9VLL3 | 137 | S | 1.644 | 0.006468912 | Up | A-kinase anchor protein 200 OS=Drosophila melanogaster OX=7227 GN=Akap200 PE=1 SV=3 | Akap200 |
| Q9VLL3 | 468 | S | 1.365 | 0.005330627 | Up | A-kinase anchor protein 200 OS=Drosophila melanogaster OX=7227 GN=Akap200 PE=1 SV=3 | Akap200 |
| Q86BS3 | 380 | S | 1.351 | 0.016835924 | Up | Chromator, isoform A OS=Drosophila melanogaster OX=7227 GN=Chro PE=1 SV=1 | Chro |
| Q86BS3 | 542 | S | 1.351 | 0.004445894 | Up | Chromator, isoform A OS=Drosophila melanogaster OX=7227 GN=Chro PE=1 SV=1 | Chro |
| Q8IMM7 | 80 | S | 0.741 | 0.005180953 | Down | Minotaur, isoform B OS=Drosophila melanogaster OX=7227 GN=mino PE=1 SV=1 | mino |
| Q8IMT3 | 47 | S | 3.426 | 4.81688E-05 | Up | IP12392p OS=Drosophila melanogaster OX=7227 GN=Dmel\CG31436 PE=2 SV=1 | Dmel\CG31436 |
| Q8IMV1 | 30 | S | 1.76 | 0.000512842 | Up | Elastin-like, isoform B OS=Drosophila melanogaster OX=7227 GN=Elal PE=4 SV=1 | Elal |
| Q8IMV6 | 476 | S | 1.57 | 0.006376606 | Up | Scaffold attachment factor B, isoform B OS=Drosophila melanogaster OX=7227 GN=Saf-B PE=1 SV=2 | Saf-B |
| Q8IMV6 | 447 | S | 2.466 | 0.001998166 | Up | Scaffold attachment factor B, isoform B OS=Drosophila melanogaster OX=7227 GN=Saf-B PE=1 SV=2 | Saf-B |
| Q8IMY7 | 75 | S | 2.297 | 0.001591082 | Up | Inwardly rectifying potassium channel 2, isoform B OS=Drosophila melanogaster OX=7227 GN=Irk2 PE=3 SV=1 | Irk2 |
| Q9VD13 | 251 | S | 1.452 | 0.000682809 | Up | GH02671p OS=Drosophila melanogaster OX=7227 GN=lqfR PE=1 SV=4 | lqfR |
| Q9VD13 | 334 | S | 1.332 | 0.029390185 | Up | GH02671p OS=Drosophila melanogaster OX=7227 GN=lqfR PE=1 SV=4 | lqfR |
| Q9VD13 | 362 | S | 1.88 | 0.002799554 | Up | GH02671p OS=Drosophila melanogaster OX=7227 GN=lqfR PE=1 SV=4 | lqfR |
| Q8INF8 | 591 | S | 1.504 | 0.020753064 | Up | FI20128p1 OS=Drosophila melanogaster OX=7227 GN=BcDNA:RH31685 PE=2 SV=1 | BcDNA:RH31685 |
| Q8ING0 | 184 | S | 0.641 | 0.001208369 | Down | Peptidase S1 domain-containing protein OS=Drosophila melanogaster OX=7227 GN=CG9645 PE=3 SV=1 | CG9645 |
| Q8INX7 | 458 | S | 1.392 | 0.000163729 | Up | Fasciclin 3, isoform C OS=Drosophila melanogaster OX=7227 GN=Fas3 PE=4 SV=2 | Fas3 |
| Q9V475 | 742 | S | 1.368 | 0.018427096 | Up | Cullin 3, isoform C OS=Drosophila melanogaster OX=7227 GN=Cul3 PE=1 SV=1 | Cul3 |
| Q8IPG9 | 258 | S | 1.316 | 0.018222443 | Up | Basigin, isoform A OS=Drosophila melanogaster OX=7227 GN=Bsg PE=1 SV=1 | Bsg |
| Q9I7P8 | 219 | S | 0.758 | 0.000893594 | Down | Tenzing norgay, isoform B OS=Drosophila melanogaster OX=7227 GN=tzn PE=1 SV=1 | tzn |
| Q8IQ30 | 894 | S | 1.413 | 0.001215293 | Up | Uncharacterized protein, isoform C OS=Drosophila melanogaster OX=7227 GN=tbc PE=4 SV=3 | tbc |
| Q8IQ30 | 257 | S | 1.663 | 0.002896733 | Up | Uncharacterized protein, isoform C OS=Drosophila melanogaster OX=7227 GN=tbc PE=4 SV=3 | tbc |
| Q8IQC8 | 928 | S | 1.599 | 0.022643124 | Up | FI05293p OS=Drosophila melanogaster OX=7227 GN=CG6740 PE=2 SV=4 | CG6740 |
| Q8IQE6 | 485 | S | 0.183 | 0.000106978 | Down | Insulin receptor substrate 53 kDa, isoform A OS=Drosophila melanogaster OX=7227 GN=IRSp53 PE=4 SV=2 | IRSp53 |

| | | | | | | | |
|---|---|---|---|---|---|---|---|
| Q8IQQ7 | 754 | S | 2.151 | 0.005419217 | Up | Bloated tubules, isoform B OS=Drosophila melanogaster OX=7227 GN=blot PE=2 SV=1 | blot |
| Q8IQW5 | 92 | S | 1.301 | 0.040441968 | Up | RE23625p OS=Drosophila melanogaster OX=7227 GN=HspB8 PE=1 SV=1 | HspB8 |
| Q8IQX3 | 155 | S | 1.611 | 0.004969168 | Up | Uncharacterized protein, isoform A OS=Drosophila melanogaster OX=7227 GN=Dmel\CG32544 PE=1 SV=1 | Dmel\CG32544 |
| Q8IQX3 | 116 | S | 2.185 | 0.00648617 | Up | Uncharacterized protein, isoform A OS=Drosophila melanogaster OX=7227 GN=Dmel\CG32544 PE=1 SV=1 | Dmel\CG32544 |
| Q8IQX3 | 119 | S | 2.185 | 0.00648617 | Up | Uncharacterized protein, isoform A OS=Drosophila melanogaster OX=7227 GN=Dmel\CG32544 PE=1 SV=1 | Dmel\CG32544 |
| Q8IR72 | 134 | S | 1.927 | 0.000747048 | Up | FI19011p1 OS=Drosophila melanogaster OX=7227 GN=Dmel\CG32638 PE=1 SV=2 | Dmel\CG32638 |
| Q8IRE3 | 1182 | S | 1.331 | 0.001590591 | Up | Gryzun OS=Drosophila melanogaster OX=7227 GN=gry PE=1 SV=2 | gry |
| Q8IRI5 | 140 | S | 1.998 | 0.003140096 | Up | Trio, isoform D OS=Drosophila melanogaster OX=7227 GN=trio PE=1 SV=1 | trio |
| Q8MLR7 | 26 | S | 1.353 | 0.001745526 | Up | GH09546p OS=Drosophila melanogaster OX=7227 GN=BcDNA PE=1 SV=1 | BcDNA |
| Q8MLV1 | 211 | S | 1.43 | 0.009338284 | Up | Lamin-B receptor OS=Drosophila melanogaster OX=7227 GN=LBR PE=1 SV=1 | LBR |
| Q9W2I6 | 270 | S | 1.478 | 0.001169736 | Up | Major facilitator superfamily transporter 16, isoform A OS=Drosophila melanogaster OX=7227 GN=MFS16 PE=2 SV=3 | MFS16 |
| Q8MMD2 | 777 | S | 1.632 | 0.00242684 | Up | Epidermal growth factor receptor pathway substrate clone 15, isoform B OS=Drosophila melanogaster OX=7227 GN=Eps-15 PE=1 SV=1 | Eps-15 |
| Q8MQW8 | 1704 | S | 1.342 | 0.018588615 | Up | Protein sprint OS=Drosophila melanogaster OX=7227 GN=spri PE=2 SV=3 | spri |
| Q9VX44 | 510 | S | 1.423 | 0.045269169 | Up | Ankyrin repeat and LEM domain containing 2, isoform F OS=Drosophila melanogaster OX=7227 GN=Ankle2 PE=1 SV=2 | Ankle2 |
| Q8MRM6 | 230 | S | 2.35 | 9.29218E-05 | Up | GH15213p OS=Drosophila melanogaster OX=7227 GN=Dmel\CG12065 PE=1 SV=1 | Dmel\CG12065 |
| Q8MRM6 | 234 | S | 1.951 | 0.000112127 | Up | GH15213p OS=Drosophila melanogaster OX=7227 GN=Dmel\CG12065 PE=1 SV=1 | Dmel\CG12065 |
| Q8MRM6 | 238 | S | 1.71 | 8.45662E-05 | Up | GH15213p OS=Drosophila melanogaster OX=7227 GN=Dmel\CG12065 PE=1 SV=1 | Dmel\CG12065 |
| Q8MRQ1 | 221 | S | 2.419 | 0.006149341 | Up | GH06222p OS=Drosophila melanogaster OX=7227 GN=Dmel\CG13124 PE=2 SV=1 | Dmel\CG13124 |
| Q8MSQ4 | 1176 | S | 1.867 | 0.013869413 | Up | Mekk1, isoform B OS=Drosophila melanogaster OX=7227 GN=Mekk1 PE=2 SV=1 | Mekk1 |
| Q8MUJ1 | 235 | S | 2.8 | 1.027E-05 | Up | Protein eiger OS=Drosophila melanogaster OX=7227 GN=egr PE=1 SV=1 | egr |
| Q8MUJ1 | 240 | S | 2.8 | 1.027E-05 | Up | Protein eiger OS=Drosophila melanogaster OX=7227 GN=egr PE=1 SV=1 | egr |
| Q8SWR8 | 25 | S | 1.74 | 0.03512938 | Up | Ataxin-2 homolog OS=Drosophila melanogaster OX=7227 GN=Atx2 PE=1 SV=1 | Atx2 |
| Q8SX76 | 417 | S | 1.845 | 0.013061827 | Up | LD24646p OS=Drosophila melanogaster OX=7227 GN=pch2 PE=1 SV=1 | pch2 |
| Q8SXX1 | 310 | S | 0.58 | 0.005236156 | Down | Phosphatidylinositol 5-phosphate 4-kinase, isoform A OS=Drosophila melanogaster OX=7227 GN=PIP4K PE=1 SV=1 | PIP4K |
| Q8T043 | 591 | S | 2.27 | 0.02857849 | Up | Chaski, isoform A OS=Drosophila melanogaster OX=7227 GN=chk PE=2 SV=1 | chk |
| Q8T051 | 260 | S | 1.727 | 0.005044903 | Up | LD28458p OS=Drosophila melanogaster OX=7227 GN=CG4639 PE=2 SV=1 | CG4639 |
| Q8T0Q2 | 44 | S | 2.041 | 0.01170165 | Up | GH14260p OS=Drosophila melanogaster OX=7227 GN=Dmel\CG9775 PE=2 SV=1 | Dmel\CG9775 |
| Q8T0V2 | 123 | S | 1.798 | 0.001217064 | Up | GH02495p OS=Drosophila melanogaster OX=7227 GN=MESK2 PE=1 SV=1 | MESK2 |
| Q8T0V2 | 432 | S | 1.63 | 0.040359929 | Up | GH02495p OS=Drosophila melanogaster OX=7227 GN=MESK2 PE=1 SV=1 | MESK2 |
| Q94522 | 145 | S | 0.745 | 0.003354226 | Down | Succinate--CoA ligase [ADP/GDP-forming] subunit alpha, mitochondrial OS=Drosophila melanogaster OX=7227 GN=Scsalpha1 PE=2 SV=3 | Scsalpha1 |
| X2JD82 | 373 | S | 0.654 | 3.34867E-05 | Down | Open rectifier K[+] channel 1, isoform D OS=Drosophila melanogaster OX=7227 GN=Ork1 PE=3 SV=1 | Ork1 |
| X2JD82 | 914 | S | 1.321 | 0.043393203 | Up | Open rectifier K[+] channel 1, isoform D OS=Drosophila melanogaster OX=7227 GN=Ork1 PE=3 SV=1 | Ork1 |
| X2JD82 | 332 | S | 0.722 | 0.00219257 | Down | Open rectifier K[+] channel 1, isoform D OS=Drosophila melanogaster OX=7227 GN=Ork1 PE=3 SV=1 | Ork1 |
| Q94527 | 45 | S | 1.356 | 0.01782631 | Up | Nuclear factor NF-kappa-B p110 subunit OS=Drosophila melanogaster OX=7227 GN=Rel PE=1 SV=1 | Rel |
| Q94915 | 254 | S | 0.595 | 0.000370416 | Down | Rhythmically expressed gene 2 protein OS=Drosophila melanogaster OX=7227 GN=Reg-2 PE=2 SV=1 | Reg-2 |
| Q95RB1 | 219 | S | 1.876 | 0.00052547 | Up | LD46870p OS=Drosophila melanogaster OX=7227 GN=Dmel\CG14641 PE=1 SV=1 | Dmel\CG14641 |
| Q95SH0 | 449 | S | 1.492 | 0.000954881 | Up | GH26463p OS=Drosophila melanogaster OX=7227 GN=Pask PE=2 SV=1 | Pask |
| Q95SH7 | 283 | S | 1.505 | 0.002027153 | Up | GH26007p OS=Drosophila melanogaster OX=7227 GN=Dmel\CG14969 PE=1 SV=1 | Dmel\CG14969 |
| Q95SH7 | 284 | S | 1.505 | 0.002027153 | Up | GH26007p OS=Drosophila melanogaster OX=7227 GN=Dmel\CG14969 PE=1 SV=1 | Dmel\CG14969 |
| Q95SH7 | 300 | S | 1.702 | 2.3903E-05 | Up | GH26007p OS=Drosophila melanogaster OX=7227 GN=Dmel\CG14969 PE=1 SV=1 | Dmel\CG14969 |
| Q95SH7 | 303 | S | 1.52 | 0.000452879 | Up | GH26007p OS=Drosophila melanogaster OX=7227 GN=Dmel\CG14969 PE=1 SV=1 | Dmel\CG14969 |
| Q95SH7 | 304 | S | 1.539 | 0.000386393 | Up | GH26007p OS=Drosophila melanogaster OX=7227 GN=Dmel\CG14969 PE=1 SV=1 | Dmel\CG14969 |
| Q9VEH0 | 129 | S | 1.372 | 0.03619336 | Up | Aluminum tubes, isoform A OS=Drosophila melanogaster OX=7227 GN=alt PE=1 SV=1 | alt |
| Q961D1 | 251 | S | 1.714 | 0.000364302 | Up | Cyclin K, isoform A OS=Drosophila melanogaster OX=7227 GN=CycK PE=1 SV=1 | CycK |
| Q961J5 | 585 | S | 1.521 | 0.001649882 | Up | Beta-alanine transporter OS=Drosophila melanogaster OX=7227 GN=Balat PE=2 SV=1 | Balat |
| Q9I7S6 | 1160 | S | 1.415 | 0.000527715 | Up | High affinity cGMP-specific 3',5'-cyclic phosphodiesterase 9A OS=Drosophila melanogaster OX=7227 GN=Pde9 PE=2 SV=2 | Pde9 |
| X2JF19 | 2619 | S | 1.348 | 0.002573066 | Up | Highwire, isoform B OS=Drosophila melanogaster OX=7227 GN=hiw PE=4 SV=1 | hiw |
| Q9NHE5 | 142 | S | 1.345 | 0.00557374 | Up | Calcium-dependent secretion activator OS=Drosophila melanogaster OX=7227 GN=Cadps PE=1 SV=3 | Cadps |
| Q9TVP3 | 164 | S | 1.846 | 0.005540737 | Up | J domain-containing protein OS=Drosophila melanogaster OX=7227 GN=jdp PE=2 SV=2 | jdp |
| Q9V396 | 29 | S | 1.439 | 0.002012492 | Up | Carbonic anhydrase 1 OS=Drosophila melanogaster OX=7227 GN=CAH1 PE=1 SV=1 | CAH1 |
| Q9V3C8 | 256 | S | 1.449 | 0.001245647 | Up | DShc protein OS=Drosophila melanogaster OX=7227 GN=Shc PE=1 SV=1 | Shc |
| Q9V3H9 | 233 | S | 3.728 | 0.000134385 | Up | BcDNA.LD27873 OS=Drosophila melanogaster OX=7227 GN=Nab2 PE=1 SV=1 | Nab2 |
| Q9V3Q9 | 751 | S | 0.715 | 0.026419236 | Down | IP03705p OS=Drosophila melanogaster OX=7227 GN=GABA-B-R1 PE=2 SV=3 | GABA-B-R1 |
| Q9V3V3 | 805 | S | 1.365 | 0.006062113 | Up | Pez, isoform A OS=Drosophila melanogaster OX=7227 GN=Pez PE=2 SV=1 | Pez |
| Q9V3X4 | 334 | S | 1.414 | 0.003355234 | Up | Seipin OS=Drosophila melanogaster OX=7227 GN=Seipin PE=1 SV=1 | Seipin |
| Q9V477 | 1275 | S | 2.197 | 0.003266923 | Up | Toll-like receptor Tollo OS=Drosophila melanogaster OX=7227 GN=Tollo PE=1 SV=1 | Tollo |
| Q9V483 | 867 | S | 0.366 | 3.10775E-05 | Down | Unc-13, isoform C OS=Drosophila melanogaster OX=7227 GN=unc-13 PE=4 SV=4 | unc-13 |
| Q9V483 | 873 | S | 1.342 | 0.013554303 | Up | Unc-13, isoform C OS=Drosophila melanogaster OX=7227 GN=unc-13 PE=4 SV=4 | unc-13 |
| Q9V4D4 | 1204 | S | 1.431 | 0.009552231 | Up | Bip2 OS=Drosophila melanogaster OX=7227 GN=bip2 PE=1 SV=1 | bip2 |
| Q9V4E7 | 635 | S | 3.27 | 1.22824E-05 | Up | Transporter OS=Drosophila melanogaster OX=7227 GN=Gat PE=1 SV=4 | Gat |
| Q9V4E7 | 32 | S | 2.322 | 0.022339886 | Up | Transporter OS=Drosophila melanogaster OX=7227 GN=Gat PE=1 SV=4 | Gat |
| Q9V4E7 | 37 | S | 1.947 | 0.005088745 | Up | Transporter OS=Drosophila melanogaster OX=7227 GN=Gat PE=1 SV=4 | Gat |
| Q9V4E7 | 20 | S | 1.934 | 1.28348E-05 | Up | Transporter OS=Drosophila melanogaster OX=7227 GN=Gat PE=1 SV=4 | Gat |

| | | | | | | | |
|---|---|---|---|---|---|---|---|
| Q9V4E7 | 5 | S | 2.548 | 0.002066728 | Up | Transporter OS=Drosophila melanogaster OX=7227 GN=Gat PE=1 SV=4 | Gat |
| Q9V4E7 | 7 | S | 2.629 | 0.002909583 | Up | Transporter OS=Drosophila melanogaster OX=7227 GN=Gat PE=1 SV=4 | Gat |
| Q9V5M6 | 374 | S | 1.306 | 0.000280615 | Up | Longitudinals lacking protein, isoforms J/P/Q/S/Z OS=Drosophila melanogaster OX=7227 GN=lola PE=1 SV=4 | lola |
| Q9V5M6 | 378 | S | 1.513 | 0.002706616 | Up | Longitudinals lacking protein, isoforms J/P/Q/S/Z OS=Drosophila melanogaster OX=7227 GN=lola PE=1 SV=4 | lola |
| Q9V7P1 | 164 | S | 1.304 | 0.001795931 | Up | U3 small nucleolar RNA-associated protein 18 homolog OS=Drosophila melanogaster OX=7227 GN=wcd PE=1 SV=1 | wcd |
| Q9V7P1 | 165 | S | 1.304 | 0.001795931 | Up | U3 small nucleolar RNA-associated protein 18 homolog OS=Drosophila melanogaster OX=7227 GN=wcd PE=1 SV=1 | wcd |
| Q9VA38 | 624 | S | 1.673 | 0.040364773 | Up | Serine/threonine-protein kinase Warts OS=Drosophila melanogaster OX=7227 GN=wts PE=1 SV=1 | wts |
| Q9VAP3 | 4 | S | 1.606 | 0.001730408 | Up | CTL-like protein 2 OS=Drosophila melanogaster OX=7227 GN=CG11880 PE=3 SV=1 | CG11880 |
| Q9VAU5 | 496 | S | 6.067 | 4.13475E-05 | Up | ANK_REP_REGION domain-containing protein OS=Drosophila melanogaster OX=7227 GN=Dmel\CG10011 PE=4 SV=1 | Dmel\CG10011 |
| Q9VB20 | 1301 | S | 1.338 | 0.011160524 | Up | Distracted, isoform B OS=Drosophila melanogaster OX=7227 GN=dsd PE=4 SV=2 | dsd |
| Q9VB22 | 436 | S | 1.478 | 0.000234552 | Up | LD33695p OS=Drosophila melanogaster OX=7227 GN=pins PE=1 SV=1 | pins |
| Q9VBH8 | 12 | S | 1.521 | 0.040817605 | Up | FI02850p OS=Drosophila melanogaster OX=7227 GN=RpL34a PE=1 SV=2 | RpL34a |
| Q9VBJ2 | 2526 | S | 1.617 | 0.003158082 | Up | Neurofibromin 1, isoform B OS=Drosophila melanogaster OX=7227 GN=Nf1 PE=4 SV=3 | Nf1 |
| Q9VBJ2 | 2558 | S | 1.645 | 0.011508207 | Up | Neurofibromin 1, isoform B OS=Drosophila melanogaster OX=7227 GN=Nf1 PE=4 SV=3 | Nf1 |
| Q9VBJ3 | 942 | S | 1.724 | 0.00812143 | Up | Fire dancer OS=Drosophila melanogaster OX=7227 GN=fid PE=4 SV=3 | fid |
| Q9VBX1 | 738 | S | 0.57 | 0.000426171 | Down | Nuclear export mediator factor NEMF homolog OS=Drosophila melanogaster OX=7227 GN=Clbn PE=1 SV=2 | Clbn |
| Q9VC02 | 20 | S | 2.082 | 0.002923217 | Up | Mahogany OS=Drosophila melanogaster OX=7227 GN=mah PE=4 SV=1 | mah |
| Q9VC54 | 299 | S | 1.315 | 0.005690579 | Up | SD06668p OS=Drosophila melanogaster OX=7227 GN=Dmel\CG6695 PE=2 SV=3 | Dmel\CG6695 |
| Q9VC54 | 824 | S | 1.563 | 0.000873126 | Up | SD06668p OS=Drosophila melanogaster OX=7227 GN=Dmel\CG6695 PE=2 SV=3 | Dmel\CG6695 |
| Q9VC60 | 283 | S | 1.586 | 0.019862131 | Up | GH07383p OS=Drosophila melanogaster OX=7227 GN=anon-WO0118547.376 PE=2 SV=1 | anon-WO0118547.376 |
| Q9VC60 | 286 | S | 1.586 | 0.019862131 | Up | GH07383p OS=Drosophila melanogaster OX=7227 GN=anon-WO0118547.376 PE=2 SV=1 | anon-WO0118547.376 |
| Q9VC94 | 1112 | S | 1.361 | 0.022325399 | Up | LD41803p OS=Drosophila melanogaster OX=7227 GN=Dmel\CG5728 PE=1 SV=2 | Dmel\CG5728 |
| Q9VC94 | 1113 | S | 1.361 | 0.022325399 | Up | LD41803p OS=Drosophila melanogaster OX=7227 GN=Dmel\CG5728 PE=1 SV=2 | Dmel\CG5728 |
| Q9VC94 | 1108 | S | 1.361 | 0.022325399 | Up | LD41803p OS=Drosophila melanogaster OX=7227 GN=Dmel\CG5728 PE=1 SV=2 | Dmel\CG5728 |
| Q9VCD4 | 38 | S | 2.427 | 6.2211E-05 | Up | Uncharacterized protein, isoform A OS=Drosophila melanogaster OX=7227 GN=Dmel\CG18428 PE=4 SV=2 | Dmel\CG18428 |
| Q9VCE6 | 169 | S | 1.317 | 0.015990194 | Up | N6-adenosine-methyltransferase MT-A70-like protein OS=Drosophila melanogaster OX=7227 GN=Mettl3 PE=1 SV=1 | Mettl3 |
| Q9VCE6 | 171 | S | 1.317 | 0.015990194 | Up | N6-adenosine-methyltransferase MT-A70-like protein OS=Drosophila melanogaster OX=7227 GN=Mettl3 PE=1 SV=1 | Mettl3 |
| Q9VCF1 | 226 | S | 1.33 | 0.003098567 | Up | GH22341p OS=Drosophila melanogaster OX=7227 GN=Dmel\CG13603 PE=1 SV=1 | Dmel\CG13603 |
| Q9VCF1 | 227 | S | 1.334 | 0.002659501 | Up | GH22341p OS=Drosophila melanogaster OX=7227 GN=Dmel\CG13603 PE=1 SV=1 | Dmel\CG13603 |
| Q9VCI3 | 8 | S | 0.706 | 0.026767908 | Down | Lipid storage droplets surface-binding protein 1 OS=Drosophila melanogaster OX=7227 GN=Lsd-1 PE=2 SV=2 | Lsd-1 |
| Q9VCI3 | 20 | S | 0.629 | 0.004133443 | Down | Lipid storage droplets surface-binding protein 1 OS=Drosophila melanogaster OX=7227 GN=Lsd-1 PE=2 SV=2 | Lsd-1 |
| Q9VCU6 | 35 | S | 1.423 | 0.018954854 | Up | Heterochromatin protein 1c OS=Drosophila melanogaster OX=7227 GN=HP1c PE=1 SV=1 | HP1c |
| Q9VDD2 | 343 | S | 1.608 | 0.030845876 | Up | LD22662p OS=Drosophila melanogaster OX=7227 GN=SNF4Agamma PE=1 SV=2 | SNF4Agamma |
| Q9VDD8 | 486 | S | 1.498 | 0.024925078 | Up | Ubiquitin carboxyl-terminal hydrolase OS=Drosophila melanogaster OX=7227 GN=Usp8 PE=1 SV=2 | Usp8 |
| Q9VDE9 | 506 | S | 1.366 | 0.000740188 | Up | FI04035p OS=Drosophila melanogaster OX=7227 GN=RhoGAP93B PE=2 SV=2 | RhoGAP93B |
| Q9VDE9 | 410 | S | 1.61 | 0.00099252 | Up | FI04035p OS=Drosophila melanogaster OX=7227 GN=RhoGAP93B PE=2 SV=2 | RhoGAP93B |
| Q9VDE9 | 413 | S | 1.61 | 0.00099252 | Up | FI04035p OS=Drosophila melanogaster OX=7227 GN=RhoGAP93B PE=2 SV=2 | RhoGAP93B |
| Q9VDH5 | 848 | S | 1.482 | 0.016736642 | Up | Kainate-type ionotropic glutamate receptor subunit 1D OS=Drosophila melanogaster OX=7227 GN=KaiR1D PE=1 SV=1 | KaiR1D |
| Q9VDK9 | 509 | S | 1.991 | 0.000379174 | Up | GH12359p OS=Drosophila melanogaster OX=7227 GN=Dmel\CG16953 PE=1 SV=3 | Dmel\CG16953 |
| Q9VDK9 | 538 | S | 1.424 | 1.18879E-05 | Up | GH12359p OS=Drosophila melanogaster OX=7227 GN=Dmel\CG16953 PE=1 SV=3 | Dmel\CG16953 |
| Q9VDK9 | 541 | S | 1.477 | 0.00035755 | Up | GH12359p OS=Drosophila melanogaster OX=7227 GN=Dmel\CG16953 PE=1 SV=3 | Dmel\CG16953 |
| Q9VDK9 | 553 | S | 1.39 | 0.028933824 | Up | GH12359p OS=Drosophila melanogaster OX=7227 GN=Dmel\CG16953 PE=1 SV=3 | Dmel\CG16953 |
| Q9VDV5 | 537 | S | 1.311 | 0.004501717 | Up | FI18650p1 OS=Drosophila melanogaster OX=7227 GN=Dmel\CG6231 PE=2 SV=2 | Dmel\CG6231 |
| Q9VDV5 | 549 | S | 2.186 | 0.001342513 | Up | FI18650p1 OS=Drosophila melanogaster OX=7227 GN=Dmel\CG6231 PE=2 SV=2 | Dmel\CG6231 |
| Q9VDV5 | 569 | S | 2.17 | 0.00034528 | Up | FI18650p1 OS=Drosophila melanogaster OX=7227 GN=Dmel\CG6231 PE=2 SV=2 | Dmel\CG6231 |
| Q9VE02 | 1537 | S | 1.906 | 0.006184802 | Up | Uncharacterized protein OS=Drosophila melanogaster OX=7227 GN=Dmel\CG6040 PE=4 SV=2 | Dmel\CG6040 |
| Q9VE73 | 13 | S | 1.537 | 0.012798177 | Up | GH17724p OS=Drosophila melanogaster OX=7227 GN=Wdr37 PE=1 SV=1 | Wdr37 |
| Q9VEF0 | 94 | S | 1.328 | 0.003310295 | Up | LD33778p OS=Drosophila melanogaster OX=7227 GN=Odj PE=1 SV=2 | Odj |
| Q9VEF0 | 98 | S | 1.53 | 0.00063096 | Up | LD33778p OS=Drosophila melanogaster OX=7227 GN=Odj PE=1 SV=2 | Odj |
| Q9VEF9 | 127 | S | 0.707 | 0.001998581 | Down | Uncharacterized protein, isoform E OS=Drosophila melanogaster OX=7227 GN=CG7397 PE=4 SV=1 | CG7397 |
| Q9VEN3 | 455 | S | 2.002 | 0.001382441 | Up | FI20177p1 OS=Drosophila melanogaster OX=7227 GN=GckIII PE=1 SV=1 | GckIII |
| Q9VEP8 | 489 | S | 1.859 | 0.000965612 | Up | GH11385p OS=Drosophila melanogaster OX=7227 GN=Irc PE=1 SV=2 | Irc |
| Q9VF06 | 158 | S | 1.426 | 0.029798266 | Up | GH08991p OS=Drosophila melanogaster OX=7227 GN=Dmel\CG4287 PE=2 SV=1 | Dmel\CG4287 |
| Q9VF45 | 26 | S | 1.42 | 0.001038907 | Up | GH25012p OS=Drosophila melanogaster OX=7227 GN=Dmel\CG5404 PE=2 SV=2 | Dmel\CG5404 |
| Q9VF45 | 617 | S | 2.324 | 0.001866906 | Up | GH25012p OS=Drosophila melanogaster OX=7227 GN=Dmel\CG5404 PE=2 SV=2 | Dmel\CG5404 |
| Q9VFD9 | 245 | S | 1.672 | 0.011321061 | Up | Defective proboscis extension response 9, isoform A OS=Drosophila melanogaster OX=7227 GN=dpr9 PE=1 SV=3 | dpr9 |
| Q9VFS2 | 411 | S | 1.452 | 0.001657173 | Up | Carotenoid isomerooxygenase OS=Drosophila melanogaster OX=7227 GN=ninaB PE=1 SV=1 | ninaB |
| Q9VFS4 | 162 | S | 1.751 | 0.000144196 | Up | ARL2_Bind_BART domain-containing protein OS=Drosophila melanogaster OX=7227 GN=Dmel\CG14367 PE=4 SV=1 | Dmel\CG14367 |
| Q9VG39 | 551 | S | 1.966 | 0.000724107 | Up | Karmoisin, isoform A OS=Drosophila melanogaster OX=7227 GN=kar PE=2 SV=2 | kar |
| Q9VG45 | 415 | S | 2.155 | 0.000100487 | Up | Dpp target protein OS=Drosophila melanogaster OX=7227 GN=Dtg PE=2 SV=1 | Dtg |
| Q9VG69 | 312 | S | 1.569 | 0.027753922 | Up | LP03547p OS=Drosophila melanogaster OX=7227 GN=Srlp PE=1 SV=1 | Srlp |
| Q9VG84 | 656 | S | 1.342 | 0.002535526 | Up | DNA ligase OS=Drosophila melanogaster OX=7227 GN=DNAlig3 PE=1 SV=3 | DNAlig3 |

| | | | | | | | |
|---|---|---|---|---|---|---|---|
| Q9VGA4 | 756 | S | 1.376 | 0.006073383 | Up | MBD-R2 OS=Drosophila melanogaster OX=7227 GN=MBD-R2 PE=1 SV=2 | MBD-R2 |
| Q9VGE4 | 980 | S | 1.405 | 0.002830699 | Up | FI04457p OS=Drosophila melanogaster OX=7227 GN=GCC185 PE=1 SV=3 | GCC185 |
| Q9VGG5 | 1909 | S | 1.509 | 0.004127702 | Up | Cadherin-87A OS=Drosophila melanogaster OX=7227 GN=Cad87A PE=1 SV=4 | Cad87A |
| Q9VGH7 | 805 | S | 1.892 | 0.00036868 | Up | Chloride channel protein 2 OS=Drosophila melanogaster OX=7227 GN=ClC-a PE=2 SV=3 | ClC-a |
| Q9VGH7 | 860 | S | 1.352 | 0.017734665 | Up | Chloride channel protein 2 OS=Drosophila melanogaster OX=7227 GN=ClC-a PE=2 SV=3 | ClC-a |
| Q9VGH7 | 862 | S | 1.352 | 0.017734665 | Up | Chloride channel protein 2 OS=Drosophila melanogaster OX=7227 GN=ClC-a PE=2 SV=3 | ClC-a |
| Q9VGH7 | 866 | S | 1.352 | 0.017734665 | Up | Chloride channel protein 2 OS=Drosophila melanogaster OX=7227 GN=ClC-a PE=2 SV=3 | ClC-a |
| Q9VGP1 | 582 | S | 1.788 | 0.01837801 | Up | Zinc finger FYVE domain-containing protein 26 homolog OS=Drosophila melanogaster OX=7227 GN=CG5270 PE=1 SV=3 | CG5270 |
| Q9VGP3 | 563 | S | 0.581 | 0.0002199 | Down | FI02016p OS=Drosophila melanogaster OX=7227 GN=Dmel\CG6723 PE=2 SV=1 | Dmel\CG6723 |
| Q9VGP9 | 532 | S | 35.145 | 0.000472778 | Up | KP78a OS=Drosophila melanogaster OX=7227 GN=KP78a PE=2 SV=1 | KP78a |
| Q9VH63 | 538 | S | 1.909 | 0.000117862 | Up | Uncharacterized protein OS=Drosophila melanogaster OX=7227 GN=Dmel\CG8516 PE=4 SV=1 | Dmel\CG8516 |
| Q9VH63 | 503 | S | 1.513 | 0.006985582 | Up | Uncharacterized protein OS=Drosophila melanogaster OX=7227 GN=Dmel\CG8516 PE=4 SV=1 | Dmel\CG8516 |
| Q9VH93 | 50 | S | 1.994 | 0.00021248 | Up | Uncharacterized protein, isoform A OS=Drosophila melanogaster OX=7227 GN=Dmel\CG9444 PE=4 SV=1 | Dmel\CG9444 |
| Q9VHC0 | 6 | S | 1.665 | 0.011555576 | Up | LD23870p OS=Drosophila melanogaster OX=7227 GN=RnpS1 PE=1 SV=2 | RnpS1 |
| Q9VHC3 | 362 | S | 1.366 | 0.002565616 | Up | Blistery, isoform A OS=Drosophila melanogaster OX=7227 GN=by PE=2 SV=1 | by |
| Q9VHC3 | 133 | S | 1.372 | 0.003047248 | Up | Blistery, isoform A OS=Drosophila melanogaster OX=7227 GN=by PE=2 SV=1 | by |
| Q9VHF5 | 10 | S | 2.165 | 0.033993592 | Up | LD42595p OS=Drosophila melanogaster OX=7227 GN=pasi2 PE=2 SV=1 | pasi2 |
| Q9VHI7 | 340 | S | 1.342 | 0.005877725 | Up | E3 ubiquitin-protein ligase Iruka OS=Drosophila melanogaster OX=7227 GN=Iru PE=1 SV=1 | Iru |
| Q9VHK6 | 42 | S | 0.714 | 0.016207337 | Down | GEO04668p1 OS=Drosophila melanogaster OX=7227 GN=IscU PE=1 SV=1 | IscU |
| Q9VHM0 | 275 | S | 1.379 | 0.039912162 | Up | Uncharacterized protein, isoform A OS=Drosophila melanogaster OX=7227 GN=Dmel\CG11768 PE=4 SV=1 | Dmel\CG11768 |
| Q9VHR5 | 127 | S | 1.39 | 0.002367763 | Up | Uncharacterized protein OS=Drosophila melanogaster OX=7227 GN=Dmel\CG9684 PE=1 SV=2 | Dmel\CG9684 |
| Q9VHR5 | 101 | S | 1.342 | 0.021034916 | Up | Uncharacterized protein OS=Drosophila melanogaster OX=7227 GN=Dmel\CG9684 PE=1 SV=2 | Dmel\CG9684 |
| Q9VHR5 | 6 | S | 1.82 | 0.017651337 | Up | Uncharacterized protein OS=Drosophila melanogaster OX=7227 GN=Dmel\CG9684 PE=1 SV=2 | Dmel\CG9684 |
| Q9VHW1 | 908 | S | 2.141 | 0.002187704 | Up | Muscarinic acetylcholine Receptor, B-type, isoform A OS=Drosophila melanogaster OX=7227 GN=mAChR-B PE=2 SV=2 | mAChR-B |
| Q9VHY7 | 21 | S | 4.386 | 0.004753726 | Up | HL04706p OS=Drosophila melanogaster OX=7227 GN=wtrw PE=2 SV=2 | wtrw |
| Q9VID9 | 1920 | S | 1.623 | 0.000327453 | Up | Uncharacterized protein, isoform B OS=Drosophila melanogaster OX=7227 GN=BEST:LD19244 PE=1 SV=3 | BEST:LD19244 |
| Q9VID9 | 946 | S | 2.461 | 0.00084161 | Up | Uncharacterized protein, isoform B OS=Drosophila melanogaster OX=7227 GN=BEST:LD19244 PE=1 SV=3 | BEST:LD19244 |
| Q9VIF2 | 12 | S | 1.499 | 0.002521689 | Up | GM01519p OS=Drosophila melanogaster OX=7227 GN=Pld3 PE=1 SV=1 | Pld3 |
| Q9VIK2 | 647 | S | 1.402 | 0.033471188 | Up | Carcinine transporter OS=Drosophila melanogaster OX=7227 GN=CarT PE=2 SV=1 | CarT |
| Q9VIK2 | 656 | S | 1.656 | 0.00667521 | Up | Carcinine transporter OS=Drosophila melanogaster OX=7227 GN=CarT PE=2 SV=1 | CarT |
| Q9VIL0 | 317 | S | 1.321 | 0.005349954 | Up | Protein brunelleschi OS=Drosophila melanogaster OX=7227 GN=brun PE=1 SV=2 | brun |
| Q9VIQ5 | 220 | S | 2.012 | 0.005611229 | Up | RH02620p OS=Drosophila melanogaster OX=7227 GN=Dmel\CG10680 PE=2 SV=1 | Dmel\CG10680 |
| Q9VIT5 | 15 | S | 0.652 | 0.005281793 | Down | RH01238p OS=Drosophila melanogaster OX=7227 GN=Dmel\CG13077 PE=2 SV=1 | Dmel\CG13077 |
| Q9VIU5 | 798 | S | 1.536 | 0.020626372 | Up | GH24474p OS=Drosophila melanogaster OX=7227 GN=Dmel\CG10137 PE=2 SV=2 | Dmel\CG10137 |
| Q9VIW9 | 18 | S | 1.999 | 2.18886E-07 | Up | Alkaline phosphatase 12, isoform A OS=Drosophila melanogaster OX=7227 GN=Alp12 PE=1 SV=1 | Alp12 |
| Q9VIX7 | 36 | S | 1.523 | 0.042672419 | Up | Fondue, isoform B OS=Drosophila melanogaster OX=7227 GN=fon PE=1 SV=1 | fon |
| Q9VJ12 | 315 | S | 1.433 | 0.022959711 | Up | Acinus, isoform A OS=Drosophila melanogaster OX=7227 GN=Acn PE=1 SV=1 | Acn |
| Q9VJ12 | 326 | S | 1.302 | 0.032236071 | Up | Acinus, isoform A OS=Drosophila melanogaster OX=7227 GN=Acn PE=1 SV=1 | Acn |
| Q9VJ30 | 1403 | S | 1.582 | 0.000857427 | Up | Numb-associated kinase, isoform A OS=Drosophila melanogaster OX=7227 GN=Nak PE=1 SV=2 | Nak |
| Q9VJ59 | 223 | S | 1.366 | 0.003704629 | Up | PRA1 family protein OS=Drosophila melanogaster OX=7227 GN=Jwa PE=1 SV=5 | Jwa |
| Q9VJI7 | 20 | S | 0.753 | 0.007290936 | Down | GEO08441p1 OS=Drosophila melanogaster OX=7227 GN=LSm7 PE=1 SV=1 | LSm7 |
| Q9VJZ7 | 632 | S | 5.336 | 0.000449051 | Up | Ribosomal RNA processing protein 1 homolog OS=Drosophila melanogaster OX=7227 GN=Nnp-1 PE=1 SV=1 | Nnp-1 |
| Q9VK59 | 471 | S | 0.606 | 0.021715577 | Down | LD23647p OS=Drosophila melanogaster OX=7227 GN=Dmel\CG5787 PE=1 SV=2 | Dmel\CG5787 |
| Q9VKC1 | 58 | S | 1.699 | 0.022084932 | Up | IP16805p OS=Drosophila melanogaster OX=7227 GN=RpL7-like PE=1 SV=2 | RpL7-like |
| Q9VKC3 | 118 | S | 2.118 | 0.000169086 | Up | Uncharacterized protein, isoform A OS=Drosophila melanogaster OX=7227 GN=PLCXD PE=4 SV=1 | PLCXD |
| Q9VKD0 | 1162 | S | 1.485 | 0.005064648 | Up | GH12955p OS=Drosophila melanogaster OX=7227 GN=Wdr81 PE=2 SV=1 | Wdr81 |
| Q9VKF0 | 124 | S | 1.493 | 0.001645955 | Up | Cyclin Y, isoform A OS=Drosophila melanogaster OX=7227 GN=CycY PE=1 SV=1 | CycY |
| Q9VKF0 | 166 | S | 1.331 | 0.001702512 | Up | Cyclin Y, isoform A OS=Drosophila melanogaster OX=7227 GN=CycY PE=1 SV=1 | CycY |
| Q9VKI8 | 54 | S | 0.505 | 0.005240089 | Down | GH03305p OS=Drosophila melanogaster OX=7227 GN=GH26 PE=1 SV=1 | GH26 |
| Q9VKK1 | 759 | S | 1.315 | 0.000573237 | Up | Enhancer of mRNA-decapping protein 4 homolog OS=Drosophila melanogaster OX=7227 GN=Ge-1 PE=1 SV=2 | Ge-1 |
| Q9VKP5 | 496 | S | 1.476 | 0.001774814 | Up | Uncharacterized protein, isoform A OS=Drosophila melanogaster OX=7227 GN=Dmel\CG6700 PE=1 SV=1 | Dmel\CG6700 |
| Q9VKQ9 | 10 | S | 1.582 | 0.000694337 | Up | Protein dpy-30 homolog OS=Drosophila melanogaster OX=7227 GN=Dpy-30L1 PE=1 SV=1 | Dpy-30L1 |
| Q9VKV6 | 12 | S | 1.555 | 0.030291359 | Up | FI04439p OS=Drosophila melanogaster OX=7227 GN=TBC1D16 PE=2 SV=2 | TBC1D16 |
| Q9VKV6 | 159 | S | 1.45 | 0.00334389 | Up | FI04439p OS=Drosophila melanogaster OX=7227 GN=TBC1D16 PE=2 SV=2 | TBC1D16 |
| Q9VKW5 | 660 | S | 1.357 | 0.00082773 | Up | LP07359p OS=Drosophila melanogaster OX=7227 GN=Dmel\CG5355 PE=1 SV=2 | Dmel\CG5355 |
| Q9VKX2 | 241 | S | 0.623 | 0.000565249 | Down | Malate dehydrogenase OS=Drosophila melanogaster OX=7227 GN=Mdh1 PE=1 SV=2 | Mdh1 |
| Q9VKX2 | 242 | S | 0.635 | 0.000168574 | Down | Malate dehydrogenase OS=Drosophila melanogaster OX=7227 GN=Mdh1 PE=1 SV=2 | Mdh1 |
| Q9VL50 | 153 | S | 1.515 | 0.007606383 | Up | RE23984p OS=Drosophila melanogaster OX=7227 GN=Trp1 PE=1 SV=1 | Trp1 |
| Q9VL50 | 313 | S | 1.332 | 0.000347099 | Up | RE23984p OS=Drosophila melanogaster OX=7227 GN=Trp1 PE=1 SV=1 | Trp1 |
| Q9VL70 | 24 | S | 0.502 | 0.004658394 | Down | HL08109p OS=Drosophila melanogaster OX=7227 GN=yip2 PE=1 SV=1 | yip2 |
| Q9VLA2 | 256 | S | 1.427 | 0.009407478 | Up | Transcription factor glial cells missing 2 OS=Drosophila melanogaster OX=7227 GN=gcm2 PE=2 SV=4 | gcm2 |

| | | | | | | | |
|---|---|---|---|---|---|---|---|
| Q9VLA2 | 259 | S | 1.427 | 0.009407478 | Up | Transcription factor glial cells missing 2 OS=Drosophila melanogaster OX=7227 GN=gcm2 PE=2 SV=4 | gcm2 |
| Q9VLC5 | 265 | S | 0.721 | 0.013130003 | Down | Aldehyde dehydrogenase OS=Drosophila melanogaster OX=7227 GN=Aldh PE=1 SV=1 | Aldh |
| Q9VLI5 | 149 | S | 1.433 | 0.000483476 | Up | FI05272p OS=Drosophila melanogaster OX=7227 GN=raw PE=2 SV=2 | raw |
| Q9VM10 | 526 | S | 1.528 | 0.000216084 | Up | Scavenger receptor acting in neural tissue and majority of rhodopsin is absent, isoform A OS=Drosophila melanogaster OX=7227 GN=santa-maria PE=1 SV=2 | santa-maria |
| Q9VM10 | 529 | S | 1.482 | 0.000163031 | Up | Scavenger receptor acting in neural tissue and majority of rhodopsin is absent, isoform A OS=Drosophila melanogaster OX=7227 GN=santa-maria PE=1 SV=2 | santa-maria |
| Q9VM10 | 530 | S | 1.456 | 0.000170631 | Up | Scavenger receptor acting in neural tissue and majority of rhodopsin is absent, isoform A OS=Drosophila melanogaster OX=7227 GN=santa-maria PE=1 SV=2 | santa-maria |
| Q9VM10 | 502 | S | 1.974 | 3.58866E-05 | Up | Scavenger receptor acting in neural tissue and majority of rhodopsin is absent, isoform A OS=Drosophila melanogaster OX=7227 GN=santa-maria PE=1 SV=2 | santa-maria |
| Q9VM10 | 505 | S | 2.148 | 1.65458E-05 | Up | Scavenger receptor acting in neural tissue and majority of rhodopsin is absent, isoform A OS=Drosophila melanogaster OX=7227 GN=santa-maria PE=1 SV=2 | santa-maria |
| Q9VM10 | 506 | S | 1.571 | 5.5262E-05 | Up | Scavenger receptor acting in neural tissue and majority of rhodopsin is absent, isoform A OS=Drosophila melanogaster OX=7227 GN=santa-maria PE=1 SV=2 | santa-maria |
| Q9VM10 | 500 | S | 1.609 | 0.000167728 | Up | Scavenger receptor acting in neural tissue and majority of rhodopsin is absent, isoform A OS=Drosophila melanogaster OX=7227 GN=santa-maria PE=1 SV=2 | santa-maria |
| Q9VM49 | 33 | S | 1.405 | 0.000313654 | Up | Caper, isoform A OS=Drosophila melanogaster OX=7227 GN=Caper PE=1 SV=1 | Caper |
| Q9VM49 | 35 | S | 1.405 | 0.000313654 | Up | Caper, isoform A OS=Drosophila melanogaster OX=7227 GN=Caper PE=1 SV=1 | Caper |
| Q9VM87 | 184 | S | 1.441 | 0.012176196 | Up | RH29724p OS=Drosophila melanogaster OX=7227 GN=Dmel\CG17378 PE=2 SV=2 | Dmel\CG17378 |
| Q9VM87 | 187 | S | 1.441 | 0.012176196 | Up | RH29724p OS=Drosophila melanogaster OX=7227 GN=Dmel\CG17378 PE=2 SV=2 | Dmel\CG17378 |
| Q9VM87 | 190 | S | 1.441 | 0.012176196 | Up | RH29724p OS=Drosophila melanogaster OX=7227 GN=Dmel\CG17378 PE=2 SV=2 | Dmel\CG17378 |
| Q9VM95 | 26 | S | 1.389 | 0.000371255 | Up | Protein Aatf OS=Drosophila melanogaster OX=7227 GN=Aatf PE=1 SV=1 | Aatf |
| Q9VM95 | 28 | S | 1.389 | 0.000371255 | Up | Protein Aatf OS=Drosophila melanogaster OX=7227 GN=Aatf PE=1 SV=1 | Aatf |
| Q9VM95 | 32 | S | 1.389 | 0.000371255 | Up | Protein Aatf OS=Drosophila melanogaster OX=7227 GN=Aatf PE=1 SV=1 | Aatf |
| X2J786 | 1374 | S | 1.716 | 0.008285594 | Up | Transport and golgi organization 1, isoform D OS=Drosophila melanogaster OX=7227 GN=Tango1 PE=4 SV=1 | Tango1 |
| X2J786 | 1376 | S | 1.519 | 0.030367397 | Up | Transport and golgi organization 1, isoform D OS=Drosophila melanogaster OX=7227 GN=Tango1 PE=4 SV=1 | Tango1 |
| Q9VMC7 | 226 | S | 1.333 | 0.001062427 | Up | LP11564p OS=Drosophila melanogaster OX=7227 GN=CG9545 PE=1 SV=3 | CG9545 |
| X2JDD7 | 874 | S | 3.478 | 2.74346E-05 | Up | Tiggrin, isoform B OS=Drosophila melanogaster OX=7227 GN=Tig PE=1 SV=1 | Tig |
| Q9VMF0 | 417 | S | 0.532 | 0.000612505 | Down | FI04444p OS=Drosophila melanogaster OX=7227 GN=Dmel\CG9498 PE=1 SV=1 | Dmel\CG9498 |
| Q9VMI2 | 31 | S | 1.341 | 0.013693551 | Up | GEO08832p1 OS=Drosophila melanogaster OX=7227 GN=Dmel\CG13994 PE=2 SV=1 | Dmel\CG13994 |
| Q9VML2 | 1576 | S | 1.635 | 0.012632282 | Up | Blue cheese OS=Drosophila melanogaster OX=7227 GN=bchs PE=4 SV=4 | bchs |
| Q9VML2 | 2619 | S | 1.301 | 0.003199616 | Up | Blue cheese OS=Drosophila melanogaster OX=7227 GN=bchs PE=4 SV=4 | bchs |
| Q9VMR0 | 138 | S | 0.575 | 0.003008737 | Down | GM05057p OS=Drosophila melanogaster OX=7227 GN=Dmel\CG7382 PE=1 SV=1 | Dmel\CG7382 |
| Q9VMY3 | 580 | S | 1.754 | 0.00306294 | Up | FI19613p1 OS=Drosophila melanogaster OX=7227 GN=Dmel\CG12581 PE=2 SV=2 | Dmel\CG12581 |
| Q9VMY3 | 583 | S | 1.754 | 0.00306294 | Up | FI19613p1 OS=Drosophila melanogaster OX=7227 GN=Dmel\CG12581 PE=2 SV=2 | Dmel\CG12581 |
| Q9VMY9 | 416 | S | 0.55 | 0.000121984 | Down | Guanine deaminase OS=Drosophila melanogaster OX=7227 GN=DhpD PE=1 SV=1 | DhpD |
| Q9VN21 | 63 | S | 1.423 | 0.002980095 | Up | LD30155p OS=Drosophila melanogaster OX=7227 GN=lost PE=1 SV=1 | lost |
| Q9VN55 | 413 | S | 1.412 | 0.000317063 | Up | Antimeros, isoform A OS=Drosophila melanogaster OX=7227 GN=atms PE=1 SV=1 | atms |
| Q9VN93 | 496 | S | 1.325 | 0.023343728 | Up | Putative cysteine proteinase CG12163 OS=Drosophila melanogaster OX=7227 GN=CG12163 PE=2 SV=2 | CG12163 |
| Q9VNC7 | 119 | S | 1.373 | 0.014668174 | Up | GH09711p OS=Drosophila melanogaster OX=7227 GN=NKCC PE=2 SV=2 | NKCC |
| Q9VND3 | 237 | S | 1.342 | 0.027335189 | Up | LD24833p OS=Drosophila melanogaster OX=7227 GN=Pi4KIIalpha PE=1 SV=3 | Pi4KIIalpha |
| Q9VND3 | 240 | S | 1.342 | 0.027335189 | Up | LD24833p OS=Drosophila melanogaster OX=7227 GN=Pi4KIIalpha PE=1 SV=3 | Pi4KIIalpha |
| Q9VND3 | 243 | S | 1.342 | 0.027335189 | Up | LD24833p OS=Drosophila melanogaster OX=7227 GN=Pi4KIIalpha PE=1 SV=3 | Pi4KIIalpha |
| Q9VNH6 | 456 | S | 1.405 | 0.000713939 | Up | Exocyst complex component 4 OS=Drosophila melanogaster OX=7227 GN=Sec8 PE=1 SV=3 | Sec8 |
| Q9VNH6 | 471 | S | 1.366 | 0.010859975 | Up | Exocyst complex component 4 OS=Drosophila melanogaster OX=7227 GN=Sec8 PE=1 SV=3 | Sec8 |
| Q9VNI5 | 890 | S | 1.319 | 0.034231181 | Up | LD10772p OS=Drosophila melanogaster OX=7227 GN=Dmel\CG10979 PE=1 SV=1 | Dmel\CG10979 |
| Q9VNI5 | 893 | S | 1.319 | 0.034231181 | Up | LD10772p OS=Drosophila melanogaster OX=7227 GN=Dmel\CG10979 PE=1 SV=1 | Dmel\CG10979 |
| Q9VNR6 | 406 | S | 1.388 | 0.030252204 | Up | Lethal (3) 04053 OS=Drosophila melanogaster OX=7227 GN=l(3)04053 PE=1 SV=1 | l(3)04053 |
| Q9VP46 | 1177 | S | 1.332 | 0.000504396 | Up | GH16847p OS=Drosophila melanogaster OX=7227 GN=Dmel\CG7324 PE=1 SV=1 | Dmel\CG7324 |
| Q9VP46 | 1180 | S | 1.383 | 0.043271115 | Up | GH16847p OS=Drosophila melanogaster OX=7227 GN=Dmel\CG7324 PE=1 SV=1 | Dmel\CG7324 |
| Q9VPE3 | 83 | S | 1.647 | 0.002774551 | Up | FI23527p1 OS=Drosophila melanogaster OX=7227 GN=ZnT77C PE=2 SV=1 | ZnT77C |
| Q9VPE3 | 395 | S | 1.892 | 0.000100063 | Up | FI23527p1 OS=Drosophila melanogaster OX=7227 GN=ZnT77C PE=2 SV=1 | ZnT77C |
| Q9VPW9 | 544 | S | 1.381 | 0.005564841 | Up | Uncharacterized protein OS=Drosophila melanogaster OX=7227 GN=TBC1D23 PE=1 SV=2 | TBC1D23 |
| Q9VPY4 | 154 | S | 1.521 | 0.000808329 | Up | FI21118p1 OS=Drosophila melanogaster OX=7227 GN=Dmel\CG4887 PE=1 SV=1 | Dmel\CG4887 |
| Q9VQD8 | 31 | S | 1.526 | 0.00379293 | Up | Actin-related protein 2/3 complex subunit 5 OS=Drosophila melanogaster OX=7227 GN=Arpc5 PE=1 SV=1 | Arpc5 |
| Q9VQF7 | 55 | S | 1.573 | 0.009051778 | Up | Bacchus OS=Drosophila melanogaster OX=7227 GN=Bacc PE=2 SV=1 | Bacc |
| Q9VQI6 | 129 | S | 1.693 | 0.003273586 | Up | LD25952p OS=Drosophila melanogaster OX=7227 GN=DmCG8814 PE=2 SV=1 | DmCG8814 |
| Q9VQI6 | 137 | S | 1.39 | 0.009170381 | Up | LD25952p OS=Drosophila melanogaster OX=7227 GN=DmCG8814 PE=2 SV=1 | DmCG8814 |
| Q9VQK5 | 660 | S | 1.346 | 0.010392937 | Up | LD24714p OS=Drosophila melanogaster OX=7227 GN=Dmel\CG3542 PE=1 SV=1 | Dmel\CG3542 |
| Q9VQK5 | 662 | S | 1.346 | 0.010392937 | Up | LD24714p OS=Drosophila melanogaster OX=7227 GN=Dmel\CG3542 PE=1 SV=1 | Dmel\CG3542 |
| Q9VQK5 | 664 | S | 1.346 | 0.010392937 | Up | LD24714p OS=Drosophila melanogaster OX=7227 GN=Dmel\CG3542 PE=1 SV=1 | Dmel\CG3542 |
| Q9VQM0 | 2015 | S | 1.857 | 0.020560876 | Up | Toucan, isoform A OS=Drosophila melanogaster OX=7227 GN=toc PE=4 SV=2 | toc |
| Q9VQV7 | 444 | S | 1.567 | 0.00034588 | Up | Centrosomal protein 97kDa, isoform B OS=Drosophila melanogaster OX=7227 GN=Cep97 PE=4 SV=3 | Cep97 |
| Q9VQZ0 | 14 | S | 1.45 | 0.000498815 | Up | Protein YIPF OS=Drosophila melanogaster OX=7227 GN=Dmel\CG3652 PE=1 SV=2 | Dmel\CG3652 |
| Q9VQZ0 | 16 | S | 1.45 | 0.000498815 | Up | Protein YIPF OS=Drosophila melanogaster OX=7227 GN=Dmel\CG3652 | Dmel\CG365 |

| | | | | | | | |
|---|---|---|---|---|---|---|---|
| | | | | | | PE=1 SV=2 | 2 |
| Q9VQZ3 | 750 | S | 1.458 | 0.036081287 | Up | Uncharacterized protein, isoform A OS=Drosophila melanogaster OX=7227 GN=Dmel\CG15431 PE=4 SV=1 | Dmel\CG15431 |
| Q9VR30 | 327 | S | 1.621 | 0.007784012 | Up | RE58324p OS=Drosophila melanogaster OX=7227 GN=anon-WO02059370.23 PE=2 SV=3 | anon-WO02059370.23 |
| Q9VR44 | 479 | S | 1.49 | 0.000114342 | Up | GH05102p OS=Drosophila melanogaster OX=7227 GN=CT10168 PE=1 SV=1 | CT10168 |
| Q9VRM3 | 238 | S | 1.348 | 0.002107471 | Up | Sinuous OS=Drosophila melanogaster OX=7227 GN=sinu PE=1 SV=3 | sinu |
| Q9VRP5 | 672 | S | 1.313 | 0.019857107 | Up | Ubiquitin carboxyl-terminal hydrolase 36 OS=Drosophila melanogaster OX=7227 GN=scny PE=1 SV=3 | scny |
| Q9VRP5 | 674 | S | 1.313 | 0.019857107 | Up | Ubiquitin carboxyl-terminal hydrolase 36 OS=Drosophila melanogaster OX=7227 GN=scny PE=1 SV=3 | scny |
| Q9VRV5 | 164 | S | 1.518 | 0.015255452 | Up | D19A OS=Drosophila melanogaster OX=7227 GN=D19A PE=1 SV=2 | D19A |
| Q9VS39 | 154 | S | 1.635 | 0.002205731 | Up | FI19525p1 OS=Drosophila melanogaster OX=7227 GN=Dmel\CG14830 PE=2 SV=1 | Dmel\CG14830 |
| Q9VS41 | 437 | S | 1.419 | 0.003686906 | Up | Unc-13-4A, isoform B OS=Drosophila melanogaster OX=7227 GN=unc-13-4A PE=4 SV=4 | unc-13-4A |
| Q9VSA3 | 347 | S | 0.576 | 0.020308338 | Down | Probable medium-chain specific acyl-CoA dehydrogenase, mitochondrial OS=Drosophila melanogaster OX=7227 GN=CG12262 PE=2 SV=1 | CG12262 |
| Q9VSC3 | 61 | S | 1.531 | 0.001428774 | Up | Ribonuclease X25 OS=Drosophila melanogaster OX=7227 GN=RNaseX25 PE=1 SV=1 | RNaseX25 |
| Q9VSC3 | 66 | S | 2.202 | 0.002939911 | Up | Ribonuclease X25 OS=Drosophila melanogaster OX=7227 GN=RNaseX25 PE=1 SV=1 | RNaseX25 |
| Q9VSF2 | 850 | S | 1.604 | 0.024077319 | Up | Mediator of RNA polymerase II transcription subunit 24 OS=Drosophila melanogaster OX=7227 GN=MED24 PE=1 SV=2 | MED24 |
| Q9VSK4 | 168 | S | 1.828 | 0.015684625 | Up | GM13032p OS=Drosophila melanogaster OX=7227 GN=Dmel\CG6983 PE=1 SV=1 | Dmel\CG6983 |
| Q9VSK4 | 171 | S | 1.828 | 0.015684625 | Up | GM13032p OS=Drosophila melanogaster OX=7227 GN=Dmel\CG6983 PE=1 SV=1 | Dmel\CG6983 |
| Q9VT27 | 295 | S | 1.938 | 0.013313011 | Up | CNMamide receptor OS=Drosophila melanogaster OX=7227 GN=CNMaR PE=2 SV=3 | CNMaR |
| Q9VT41 | 132 | S | 1.686 | 0.014723892 | Up | Integrator complex subunit 7 OS=Drosophila melanogaster OX=7227 GN=defl PE=1 SV=1 | defl |
| Q9VT61 | 84 | S | 1.838 | 0.004934475 | Up | LD27033p OS=Drosophila melanogaster OX=7227 GN=BcDNA:GH07921 PE=1 SV=1 | BcDNA:GH07921 |
| Q9VTB3 | 170 | S | 1.563 | 0.001972444 | Up | GH06691p OS=Drosophila melanogaster OX=7227 GN=Dmel\CG11811 PE=1 SV=1 | Dmel\CG11811 |
| Q9VTE6 | 1077 | S | 1.327 | 0.014721128 | Up | IP14658p OS=Drosophila melanogaster OX=7227 GN=Dmel\CG7839 PE=1 SV=3 | Dmel\CG7839 |
| Q9VTG0 | 383 | S | 1.575 | 0.042258595 | Up | Phosphate transporter OS=Drosophila melanogaster OX=7227 GN=NaPi-III PE=2 SV=2 | NaPi-III |
| Q9VTG0 | 440 | S | 1.362 | 0.001845323 | Up | Phosphate transporter OS=Drosophila melanogaster OX=7227 GN=NaPi-III PE=2 SV=2 | NaPi-III |
| Q9VTG8 | 54 | S | 4.666 | 1.26191E-05 | Up | Ig-like domain-containing protein OS=Drosophila melanogaster OX=7227 GN=CT23219 PE=4 SV=1 | CT23219 |
| Q9VTW8 | 904 | S | 2.224 | 4.86835E-05 | Up | GH27027p OS=Drosophila melanogaster OX=7227 GN=Ncc69 PE=1 SV=1 | Ncc69 |
| Q9VTW8 | 905 | S | 2.147 | 4.2312E-05 | Up | GH27027p OS=Drosophila melanogaster OX=7227 GN=Ncc69 PE=1 SV=1 | Ncc69 |
| Q9VTW8 | 907 | S | 2.16 | 0.000136769 | Up | GH27027p OS=Drosophila melanogaster OX=7227 GN=Ncc69 PE=1 SV=1 | Ncc69 |
| Q9VTW8 | 910 | S | 2.421 | 4.52828E-05 | Up | GH27027p OS=Drosophila melanogaster OX=7227 GN=Ncc69 PE=1 SV=1 | Ncc69 |
| Q9VU43 | 174 | S | 1.352 | 0.004971795 | Up | FI04407p OS=Drosophila melanogaster OX=7227 GN=Srrm1 PE=1 SV=1 | Srrm1 |
| Q9VU88 | 228 | S | 2.075 | 0.010753385 | Up | AT31531p OS=Drosophila melanogaster OX=7227 GN=Liprin-beta PE=2 SV=1 | Liprin-beta |
| Q9VUB5 | 63 | S | 1.556 | 0.010760979 | Up | UpSET, isoform A OS=Drosophila melanogaster OX=7227 GN=upSET PE=1 SV=3 | upSET |
| Q9VUB5 | 2214 | S | 1.347 | 0.015306674 | Up | UpSET, isoform A OS=Drosophila melanogaster OX=7227 GN=upSET PE=1 SV=3 | upSET |
| Q9VUC6 | 1175 | S | 1.765 | 0.000409494 | Up | Formin-like protein OS=Drosophila melanogaster OX=7227 GN=Frl PE=1 SV=3 | Frl |
| Q9VUC6 | 225 | S | 1.965 | 0.000558728 | Up | Formin-like protein OS=Drosophila melanogaster OX=7227 GN=Frl PE=1 SV=3 | Frl |
| Q9VUN9 | 19 | S | 1.316 | 0.015567915 | Up | FI09629p OS=Drosophila melanogaster OX=7227 GN=Dmel\CG7857 PE=1 SV=1 | Dmel\CG7857 |
| Q9VUQ5 | 93 | S | 2.683 | 9.65911E-05 | Up | Protein argonaute-2 OS=Drosophila melanogaster OX=7227 GN=AGO2 PE=1 SV=3 | AGO2 |
| Q9VUY9 | 116 | S | 0.587 | 3.1173E-05 | Down | Phosphoglucomutase OS=Drosophila melanogaster OX=7227 GN=Pgm1 PE=1 SV=1 | Pgm1 |
| Q9VV74 | 64 | S | 1.425 | 0.017121507 | Up | Survival motor neuron protein OS=Drosophila melanogaster OX=7227 GN=Smn PE=1 SV=1 | Smn |
| Q9VVL8 | 436 | S | 1.759 | 0.014842271 | Up | FI03887p OS=Drosophila melanogaster OX=7227 GN=TrpRS-m PE=1 SV=1 | TrpRS-m |
| Q9VVN1 | 265 | S | 1.483 | 0.004408334 | Up | GH19567p OS=Drosophila melanogaster OX=7227 GN=Dmel\CG13698 PE=2 SV=1 | Dmel\CG13698 |
| Q9VVN1 | 270 | S | 1.483 | 0.004408334 | Up | GH19567p OS=Drosophila melanogaster OX=7227 GN=Dmel\CG13698 PE=2 SV=1 | Dmel\CG13698 |
| Q9VVU4 | 323 | S | 1.766 | 0.042298099 | Up | Cir_N domain-containing protein OS=Drosophila melanogaster OX=7227 GN=Dmel\CG6843 PE=4 SV=2 | Dmel\CG6843 |
| X2JGV8 | 47 | S | 0.562 | 0.003354946 | Down | Uncharacterized protein, isoform B OS=Drosophila melanogaster OX=7227 GN=Dmel\CG9231 PE=4 SV=1 | Dmel\CG9231 |
| Q9VW17 | 75 | S | 2.375 | 9.42048E-05 | Up | RE58036p OS=Drosophila melanogaster OX=7227 GN=BcDNA:RH51268 PE=2 SV=1 | BcDNA:RH51268 |
| Q9VW17 | 144 | S | 1.494 | 2.0128E-05 | Up | RE58036p OS=Drosophila melanogaster OX=7227 GN=BcDNA:RH51268 PE=2 SV=1 | BcDNA:RH51268 |
| Q9VW22 | 1259 | S | 1.782 | 4.22766E-05 | Up | FI18195p1 OS=Drosophila melanogaster OX=7227 GN=l(3)76BDm PE=1 SV=2 | l(3)76BDm |
| Q9VW22 | 1260 | S | 1.764 | 0.000646761 | Up | FI18195p1 OS=Drosophila melanogaster OX=7227 GN=l(3)76BDm PE=1 SV=2 | l(3)76BDm |
| Q9VW54 | 358 | S | 1.346 | 0.004746898 | Up | 26S proteasome non-ATPase regulatory subunit 2 OS=Drosophila melanogaster OX=7227 GN=Rpn1 PE=1 SV=2 | Rpn1 |
| Q9VW73 | 57 | S | 1.542 | 0.009227186 | Up | FI16105p1 OS=Drosophila melanogaster OX=7227 GN=p18 PE=1 SV=1 | p18 |
| Q9VWA2 | 87 | S | 0.697 | 0.001027202 | Down | Probable deoxycytidylate deaminase OS=Drosophila melanogaster OX=7227 GN=CG6951 PE=2 SV=1 | CG6951 |
| Q9VWD4 | 231 | S | 1.328 | 0.013695452 | Up | Probable RNA-binding protein CG14230 OS=Drosophila melanogaster OX=7227 GN=CG14230 PE=1 SV=2 | CG14230 |
| X2JG50 | 1840 | S | 1.625 | 0.007837641 | Up | Elys, isoform B OS=Drosophila melanogaster OX=7227 GN=Elys PE=4 SV=1 | Elys |
| Q9VWI6 | 497 | S | 1.679 | 0.004796436 | Up | Kekkon 5, isoform A OS=Drosophila melanogaster OX=7227 GN=kek5 PE=1 SV=2 | kek5 |
| Q9VWJ0 | 316 | S | 1.952 | 0.002566498 | Up | SD10554p OS=Drosophila melanogaster OX=7227 GN=anon-WO03002137.1 PE=2 SV=2 | anon-WO03002137.1 |
| Q9VWJ0 | 319 | S | 1.952 | 0.002566498 | Up | SD10554p OS=Drosophila melanogaster OX=7227 GN=anon-WO03002137.1 PE=2 SV=2 | anon-WO03002137.1 |
| X2JL73 | 1904 | S | 1.458 | 0.003713325 | Up | Rapamycin-insensitive companion of Tor, isoform B OS=Drosophila melanogaster OX=7227 GN=rictor PE=1 SV=1 | rictor |
| Q9VWK5 | 1115 | S | 1.745 | 0.005893726 | Up | GH15225p OS=Drosophila melanogaster OX=7227 GN=Ulp1 PE=1 SV=1 | Ulp1 |
| Q9VWN4 | 368 | S | 1.406 | 0.00155741 | Up | Fl(2)d-associated complex component OS=Drosophila melanogaster OX=7227 GN=Flacc PE=1 SV=1 | Flacc |
| Q9VWN4 | 881 | S | 1.431 | 0.026051841 | Up | Fl(2)d-associated complex component OS=Drosophila melanogaster OX=7227 GN=Flacc PE=1 SV=1 | Flacc |
| Q9VWP5 | 1797 | S | 1.492 | 0.023516778 | Up | Hat-trick, isoform D OS=Drosophila melanogaster OX=7227 GN=htk PE=1 SV=4 | htk |
| Q9VWP8 | 914 | S | 0.767 | 0.022459732 | Down | Uncharacterized protein, isoform A OS=Drosophila melanogaster | CG32543 |

| | | | | | | | |
|---|---|---|---|---|---|---|---|
| | | | | | | OX=7227 GN=CG32543 PE=1 SV=2 | |
| Q9VWP8 | 916 | S | 0.767 | 0.022459732 | Down | Uncharacterized protein, isoform A OS=Drosophila melanogaster OX=7227 GN=CG32543 PE=1 SV=2 | CG32543 |
| Q9VWP8 | 1194 | S | 0.714 | 0.000767032 | Down | Uncharacterized protein, isoform A OS=Drosophila melanogaster OX=7227 GN=CG32543 PE=1 SV=2 | CG32543 |
| X2JFP6 | 416 | S | 1.793 | 0.005654879 | Up | Rab11 interacting protein, isoform E OS=Drosophila melanogaster OX=7227 GN=Rip11 PE=1 SV=1 | Rip11 |
| X2JFP6 | 137 | S | 1.301 | 0.049845369 | Up | Rab11 interacting protein, isoform E OS=Drosophila melanogaster OX=7227 GN=Rip11 PE=1 SV=1 | Rip11 |
| X2JFP6 | 635 | S | 1.372 | 0.00159761 | Up | Rab11 interacting protein, isoform E OS=Drosophila melanogaster OX=7227 GN=Rip11 PE=1 SV=1 | Rip11 |
| Q9VWV6 | 509 | S | 3.164 | 0.000216644 | Up | Transferrin OS=Drosophila melanogaster OX=7227 GN=Tsf1 PE=1 SV=1 | Tsf1 |
| X2JFW6 | 296 | S | 1.391 | 0.043603902 | Up | Uncharacterized protein, isoform B OS=Drosophila melanogaster OX=7227 GN=Dmel\CG8289 PE=1 SV=1 | Dmel\CG8289 |
| Q9VXA3 | 638 | S | 1.54 | 0.010976509 | Up | LP21163p OS=Drosophila melanogaster OX=7227 GN=RSG7 PE=2 SV=2 | RSG7 |
| Q9VXH3 | 491 | S | 1.553 | 0.002415355 | Up | 1-phosphatidylinositol 4,5-bisphosphate phosphodiesterase gamma OS=Drosophila melanogaster OX=7227 GN=sl PE=1 SV=3 | sl |
| Q9VXV4 | 445 | S | 1.336 | 0.017983841 | Up | RH04535p OS=Drosophila melanogaster OX=7227 GN=Dmel\CG11655 PE=1 SV=1 | Dmel\CG11655 |
| Q9VXW2 | 239 | S | 1.305 | 0.001217672 | Up | LD41277p OS=Drosophila melanogaster OX=7227 GN=Dmel\CG6227 PE=1 SV=1 | Dmel\CG6227 |
| Q9VYK6 | 924 | S | 2.715 | 0.004262524 | Up | Tomosyn, isoform C OS=Drosophila melanogaster OX=7227 GN=Tomosyn PE=1 SV=3 | Tomosyn |
| Q9VYM7 | 324 | S | 1.47 | 0.000752068 | Up | Uncharacterized protein, isoform C OS=Drosophila melanogaster OX=7227 GN=Pripl4 PE=1 SV=2 | Pripl4 |
| Q9VYM7 | 176 | S | 1.773 | 0.004951406 | Up | Uncharacterized protein, isoform C OS=Drosophila melanogaster OX=7227 GN=Pripl4 PE=1 SV=2 | Pripl4 |
| X2JEX7 | 1112 | S | 1.381 | 0.000462387 | Up | Ubiquitin-specific protease 7, isoform C OS=Drosophila melanogaster OX=7227 GN=Usp7 PE=3 SV=1 | Usp7 |
| Q9VYS2 | 307 | S | 1.726 | 0.034762723 | Up | GH09980p OS=Drosophila melanogaster OX=7227 GN=pot PE=2 SV=1 | pot |
| Q9VYU0 | 638 | S | 1.749 | 0.005375282 | Up | Rudhira, isoform C OS=Drosophila melanogaster OX=7227 GN=rudhira PE=4 SV=3 | rudhira |
| Q9VYX8 | 2 | S | 1.829 | 0.000342552 | Up | LD04844p OS=Drosophila melanogaster OX=7227 GN=Dmel\CG1572 PE=1 SV=1 | Dmel\CG1572 |
| Q9VYX8 | 4 | S | 1.919 | 0.019517846 | Up | LD04844p OS=Drosophila melanogaster OX=7227 GN=Dmel\CG1572 PE=1 SV=1 | Dmel\CG1572 |
| Q9VYY7 | 523 | S | 1.414 | 0.001961066 | Up | Heterochromatin protein 5, isoform B OS=Drosophila melanogaster OX=7227 GN=HP5 PE=1 SV=1 | HP5 |
| Q9VYY7 | 418 | S | 1.581 | 0.026595137 | Up | Heterochromatin protein 5, isoform B OS=Drosophila melanogaster OX=7227 GN=HP5 PE=1 SV=1 | HP5 |
| X2JEU3 | 63 | S | 3.651 | 0.012651814 | Up | Uncharacterized protein, isoform C OS=Drosophila melanogaster OX=7227 GN=Dmel\CG1737 PE=1 SV=1 | Dmel\CG1737 |
| Q9VZ08 | 1088 | S | 2.547 | 0.000515078 | Up | Receptor-mediated endocytosis protein 6 homolog OS=Drosophila melanogaster OX=7227 GN=CG1657 PE=1 SV=2 | CG1657 |
| Q9VZ08 | 1190 | S | 1.339 | 0.031315702 | Up | Receptor-mediated endocytosis protein 6 homolog OS=Drosophila melanogaster OX=7227 GN=CG1657 PE=1 SV=2 | CG1657 |
| X2JES6 | 41 | S | 1.651 | 0.000244208 | Up | Uncharacterized protein, isoform C OS=Drosophila melanogaster OX=7227 GN=anon-EST:Posey230 PE=4 SV=1 | anon-EST:Posey230 |
| Q9VZ67 | 16 | S | 1.41 | 0.004506799 | Up | Uncharacterized protein OS=Drosophila melanogaster OX=7227 GN=Dmel\CG15210 PE=4 SV=1 | Dmel\CG15210 |
| Q9VZK7 | 377 | S | 1.352 | 0.013128888 | Up | Uncharacterized protein, isoform B OS=Drosophila melanogaster OX=7227 GN=Dmel\CG14982 PE=4 SV=2 | Dmel\CG14982 |
| Q9VZL1 | 18 | S | 1.344 | 0.020623668 | Up | LP07226p OS=Drosophila melanogaster OX=7227 GN=mge PE=1 SV=1 | mge |
| Q9VZL5 | 266 | S | 1.314 | 0.001402301 | Up | Caffeine, calcium, zinc sensitivity 1 OS=Drosophila melanogaster OX=7227 GN=Ccz1 PE=2 SV=3 | Ccz1 |
| Q9VZS7 | 208 | S | 1.47 | 0.00044778 | Up | Hobbit, isoform B OS=Drosophila melanogaster OX=7227 GN=hob PE=1 SV=2 | hob |
| Q9VZT8 | 654 | S | 1.408 | 0.014715234 | Up | Uncharacterized protein OS=Drosophila melanogaster OX=7227 GN=Dmel\CG14964 PE=4 SV=3 | Dmel\CG14964 |
| Q9VZZ9 | 513 | S | 1.328 | 0.000324396 | Up | Protein daughter of sevenless OS=Drosophila melanogaster OX=7227 GN=dos PE=1 SV=1 | dos |
| Q9VZZ9 | 519 | S | 1.366 | 0.00496285 | Up | Protein daughter of sevenless OS=Drosophila melanogaster OX=7227 GN=dos PE=1 SV=1 | dos |
| Q9VZZ9 | 401 | S | 1.326 | 0.014472432 | Up | Protein daughter of sevenless OS=Drosophila melanogaster OX=7227 GN=dos PE=1 SV=1 | dos |
| Q9W002 | 1153 | S | 1.457 | 0.021121548 | Up | Misshapen, isoform A OS=Drosophila melanogaster OX=7227 GN=msn PE=1 SV=3 | msn |
| Q9W019 | 119 | S | 1.311 | 0.000392807 | Up | IP09724p OS=Drosophila melanogaster OX=7227 GN=Dmel\CG15877 PE=1 SV=1 | Dmel\CG15877 |
| X2JG46 | 153 | S | 1.334 | 0.046481533 | Up | Uncharacterized protein, isoform C OS=Drosophila melanogaster OX=7227 GN=Dmel\CG12104 PE=4 SV=1 | Dmel\CG12104 |
| Q9W0D3 | 1490 | S | 1.45 | 0.007792965 | Up | GH15728p OS=Drosophila melanogaster OX=7227 GN=Dmel\CG13917 PE=2 SV=1 | Dmel\CG13917 |
| Q9W0D9 | 380 | S | 1.917 | 3.30215E-05 | Up | GH25855p OS=Drosophila melanogaster OX=7227 GN=Pcyt2 PE=2 SV=1 | Pcyt2 |
| Q9W0H9 | 72 | S | 1.333 | 0.006842271 | Up | Rabaptin-5-associated exchange factor for Rab5 OS=Drosophila melanogaster OX=7227 GN=Rabex-5 PE=1 SV=1 | Rabex-5 |
| Q9W0L6 | 30 | S | 1.354 | 0.028498712 | Up | GH22266p OS=Drosophila melanogaster OX=7227 GN=Dmel\CG13907 PE=1 SV=1 | Dmel\CG13907 |
| Q9W0L7 | 682 | S | 1.981 | 0.041170388 | Up | LD28815p OS=Drosophila melanogaster OX=7227 GN=Usp10 PE=1 SV=2 | Usp10 |
| Q9W1C8 | 215 | S | 1.357 | 0.004987236 | Up | LD24355p OS=Drosophila melanogaster OX=7227 GN=Dmel\CG3163 PE=1 SV=2 | Dmel\CG3163 |
| Q9W1D9 | 300 | S | 2.349 | 0.007937165 | Up | Oxysterol-binding protein OS=Drosophila melanogaster OX=7227 GN=Dmel\CG3860 PE=1 SV=1 | Dmel\CG3860 |
| Q9W1D9 | 302 | S | 2.033 | 0.002087541 | Up | Oxysterol-binding protein OS=Drosophila melanogaster OX=7227 GN=Dmel\CG3860 PE=1 SV=1 | Dmel\CG3860 |
| Q9W1G0 | 237 | S | 0.563 | 0.000464491 | Down | Probable transaldolase OS=Drosophila melanogaster OX=7227 GN=Taldo PE=2 SV=2 | Taldo |
| Q9W266 | 518 | S | 1.785 | 4.69652E-05 | Up | Protein windpipe OS=Drosophila melanogaster OX=7227 GN=wdp PE=1 SV=1 | wdp |
| Q9W266 | 623 | S | 1.383 | 0.001932414 | Up | Protein windpipe OS=Drosophila melanogaster OX=7227 GN=wdp PE=1 SV=1 | wdp |
| Q9W266 | 654 | S | 1.609 | 0.005517511 | Up | Protein windpipe OS=Drosophila melanogaster OX=7227 GN=wdp PE=1 SV=1 | wdp |
| Q9W266 | 588 | S | 1.441 | 0.000192983 | Up | Protein windpipe OS=Drosophila melanogaster OX=7227 GN=wdp PE=1 SV=1 | wdp |
| Q9W269 | 384 | S | 1.537 | 0.006255064 | Up | Solute carrier organic anion transporter family member OS=Drosophila melanogaster OX=7227 GN=Oatp58Dc PE=1 SV=1 | Oatp58Dc |
| Q9W2F3 | 509 | S | 1.318 | 0.049312265 | Up | Protein tyrosine phosphatase ERK OS=Drosophila melanogaster OX=7227 GN=PTP-ER PE=2 SV=2 | PTP-ER |
| Q9W2R3 | 117 | S | 1.509 | 0.009672317 | Up | LP03320p OS=Drosophila melanogaster OX=7227 GN=sktl PE=1 SV=2 | sktl |
| Q9W2S3 | 704 | S | 1.49 | 0.027376834 | Up | BTB/POZ domain-containing protein 9 OS=Drosophila melanogaster OX=7227 GN=BTBD9 PE=1 SV=1 | BTBD9 |
| Q9W308 | 36 | S | 6.834 | 3.28232E-05 | Up | GH05731p OS=Drosophila melanogaster OX=7227 GN=BcDNA:GH05731 PE=2 SV=1 | BcDNA:GH05731 |
| Q9W309 | 117 | S | 3.237 | 0.0001819 | Up | GEO08445p1 OS=Drosophila melanogaster OX=7227 GN=Dmel\CG9686 PE=2 SV=1 | Dmel\CG9686 |
| Q9W309 | 122 | S | 3.399 | 0.000599202 | Up | GEO08445p1 OS=Drosophila melanogaster OX=7227 GN=Dmel\CG9686 PE=2 SV=1 | Dmel\CG9686 |
| Q9W309 | 48 | S | 1.682 | 0.030027332 | Up | GEO08445p1 OS=Drosophila melanogaster OX=7227 GN=Dmel\CG9686 PE=2 SV=1 | Dmel\CG9686 |
| Q9W335 | 281 | S | 2.118 | 0.015122648 | Up | LD26546p OS=Drosophila melanogaster OX=7227 GN=l(1)G0320 PE=1 SV=1 | l(1)G0320 |
| Q9W337 | 79 | S | 1.474 | 0.044891749 | Up | GH19985p OS=Drosophila melanogaster OX=7227 GN=CG18429 PE=2 SV=2 | CG18429 |

| | | | | | | | |
|---|---|---|---|---|---|---|---|
| Q9W337 | 98 | S | 2.036 | 0.020932136 | Up | GH19985p OS=Drosophila melanogaster OX=7227 GN=CG18429 PE=2 SV=2 | CG18429 |
| Q9W337 | 151 | S | 2.388 | 0.00096073 | Up | GH19985p OS=Drosophila melanogaster OX=7227 GN=CG18429 PE=2 SV=2 | CG18429 |
| Q9W391 | 1012 | S | 0.51 | 0.000536303 | Down | Probable phosphorylase b kinase regulatory subunit alpha OS=Drosophila melanogaster OX=7227 GN=CG7766 PE=1 SV=2 | CG7766 |
| Q9W391 | 1033 | S | 1.74 | 0.006758228 | Up | Probable phosphorylase b kinase regulatory subunit alpha OS=Drosophila melanogaster OX=7227 GN=CG7766 PE=1 SV=2 | CG7766 |
| Q9W391 | 677 | S | 1.487 | 0.006739564 | Up | Probable phosphorylase b kinase regulatory subunit alpha OS=Drosophila melanogaster OX=7227 GN=CG7766 PE=1 SV=2 | CG7766 |
| Q9W391 | 1030 | S | 1.774 | 0.005980621 | Up | Probable phosphorylase b kinase regulatory subunit alpha OS=Drosophila melanogaster OX=7227 GN=CG7766 PE=1 SV=2 | CG7766 |
| Q9W392 | 254 | S | 1.374 | 0.026271512 | Up | Chaperonin containing TCP1 subunit 2 OS=Drosophila melanogaster OX=7227 GN=CCT2 PE=1 SV=2 | CCT2 |
| Q9W3E2 | 508 | S | 1.406 | 0.016714587 | Up | PIP82 OS=Drosophila melanogaster OX=7227 GN=PIP82 PE=1 SV=3 | PIP82 |
| Q9W3E2 | 837 | S | 1.86 | 0.014295621 | Up | PIP82 OS=Drosophila melanogaster OX=7227 GN=PIP82 PE=1 SV=3 | PIP82 |
| Q9W3E2 | 654 | S | 1.307 | 0.028843586 | Up | PIP82 OS=Drosophila melanogaster OX=7227 GN=PIP82 PE=1 SV=3 | PIP82 |
| Q9W3F7 | 243 | S | 1.458 | 0.049507089 | Up | Mitoguardin OS=Drosophila melanogaster OX=7227 GN=Miga PE=1 SV=2 | Miga |
| Q9W3J9 | 468 | S | 0.744 | 0.029037899 | Down | LP13385p OS=Drosophila melanogaster OX=7227 GN=Dmel\CG2116 PE=2 SV=1 | Dmel\CG2116 |
| Q9W3W9 | 123 | S | 1.601 | 0.000201653 | Up | LP17136p OS=Drosophila melanogaster OX=7227 GN=LP06294.5prime PE=1 SV=2 | LP06294.5prime |
| Q9W3Y3 | 174 | S | 1.408 | 0.000302899 | Up | LD13807p OS=Drosophila melanogaster OX=7227 GN=Dmel\CG3226 PE=1 SV=1 | Dmel\CG3226 |
| Q9W3Y4 | 479 | S | 1.564 | 0.021769566 | Up | GAS2-like protein pickled eggs OS=Drosophila melanogaster OX=7227 GN=pigs PE=2 SV=2 | pigs |
| Q9W425 | 1469 | S | 1.746 | 0.011339295 | Up | Rabconnectin-3A OS=Drosophila melanogaster OX=7227 GN=Rbcn-3A PE=1 SV=3 | Rbcn-3A |
| Q9W425 | 2798 | S | 1.541 | 0.018707916 | Up | Rabconnectin-3A OS=Drosophila melanogaster OX=7227 GN=Rbcn-3A PE=1 SV=3 | Rbcn-3A |
| Q9W425 | 2266 | S | 1.514 | 0.034310246 | Up | Rabconnectin-3A OS=Drosophila melanogaster OX=7227 GN=Rbcn-3A PE=1 SV=3 | Rbcn-3A |
| Q9W425 | 1056 | S | 1.718 | 0.003288368 | Up | Rabconnectin-3A OS=Drosophila melanogaster OX=7227 GN=Rbcn-3A PE=1 SV=3 | Rbcn-3A |
| Q9W425 | 1387 | S | 1.336 | 0.011923126 | Up | Rabconnectin-3A OS=Drosophila melanogaster OX=7227 GN=Rbcn-3A PE=1 SV=3 | Rbcn-3A |
| Q9W425 | 1362 | S | 1.332 | 0.04747651 | Up | Rabconnectin-3A OS=Drosophila melanogaster OX=7227 GN=Rbcn-3A PE=1 SV=3 | Rbcn-3A |
| Q9W425 | 967 | S | 0.585 | 0.005699251 | Down | Rabconnectin-3A OS=Drosophila melanogaster OX=7227 GN=Rbcn-3A PE=1 SV=3 | Rbcn-3A |
| Q9W425 | 976 | S | 0.585 | 0.005699251 | Down | Rabconnectin-3A OS=Drosophila melanogaster OX=7227 GN=Rbcn-3A PE=1 SV=3 | Rbcn-3A |
| Q9W425 | 977 | S | 0.585 | 0.005699251 | Down | Rabconnectin-3A OS=Drosophila melanogaster OX=7227 GN=Rbcn-3A PE=1 SV=3 | Rbcn-3A |
| Q9W425 | 516 | S | 1.343 | 0.000248689 | Up | Rabconnectin-3A OS=Drosophila melanogaster OX=7227 GN=Rbcn-3A PE=1 SV=3 | Rbcn-3A |
| Q9W425 | 518 | S | 1.369 | 0.000249036 | Up | Rabconnectin-3A OS=Drosophila melanogaster OX=7227 GN=Rbcn-3A PE=1 SV=3 | Rbcn-3A |
| Q9W425 | 1579 | S | 1.454 | 0.031092359 | Up | Rabconnectin-3A OS=Drosophila melanogaster OX=7227 GN=Rbcn-3A PE=1 SV=3 | Rbcn-3A |
| Q9W427 | 405 | S | 1.894 | 8.4445E-05 | Up | Detonator, isoform B OS=Drosophila melanogaster OX=7227 GN=dtn PE=4 SV=2 | dtn |
| Q9W441 | 267 | S | 0.446 | 4.67616E-05 | Down | Tetraspanin OS=Drosophila melanogaster OX=7227 GN=Tsp5D PE=2 SV=4 | Tsp5D |
| Q9W441 | 269 | S | 0.431 | 2.04799E-05 | Down | Tetraspanin OS=Drosophila melanogaster OX=7227 GN=Tsp5D PE=2 SV=4 | Tsp5D |
| Q9W441 | 271 | S | 1.745 | 0.00080117 | Up | Tetraspanin OS=Drosophila melanogaster OX=7227 GN=Tsp5D PE=2 SV=4 | Tsp5D |
| Q9W441 | 193 | S | 1.842 | 0.000556578 | Up | Tetraspanin OS=Drosophila melanogaster OX=7227 GN=Tsp5D PE=2 SV=4 | Tsp5D |
| Q9W490 | 187 | S | 1.332 | 0.025814188 | Up | FI04818p OS=Drosophila melanogaster OX=7227 GN=CG15774 PE=2 SV=2 | CG15774 |
| Q9W4B6 | 2 | S | 3.092 | 4.00131E-05 | Up | RE01432p OS=Drosophila melanogaster OX=7227 GN=BcDNA:RE01432 PE=2 SV=1 | BcDNA:RE01432 |
| Q9W4C1 | 283 | S | 1.451 | 0.00268053 | Up | IP13321p OS=Drosophila melanogaster OX=7227 GN=Dmel\CG15784 PE=1 SV=1 | Dmel\CG15784 |
| Q9W4C1 | 322 | S | 1.696 | 0.00456013 | Up | IP13321p OS=Drosophila melanogaster OX=7227 GN=Dmel\CG15784 PE=1 SV=1 | Dmel\CG15784 |
| Q9W4C1 | 324 | S | 1.696 | 0.00456013 | Up | IP13321p OS=Drosophila melanogaster OX=7227 GN=Dmel\CG15784 PE=1 SV=1 | Dmel\CG15784 |
| Q9W4X7 | 198 | S | 1.392 | 0.000229832 | Up | Eukaryotic translation initiation factor 3 subunit G-1 OS=Drosophila melanogaster OX=7227 GN=eIF3g1 PE=1 SV=1 | eIF3g1 |
| Q9W542 | 564 | S | 2.185 | 0.001781791 | Up | LD07342p OS=Drosophila melanogaster OX=7227 GN=mip130 PE=1 SV=3 | mip130 |
| Q9W5C2 | 361 | S | 1.372 | 7.02473E-05 | Up | Translocating chain-associated membrane protein OS=Drosophila melanogaster OX=7227 GN=TRAM PE=1 SV=1 | TRAM |
| X2JDC1 | 636 | S | 1.342 | 0.002048118 | Up | Uncharacterized protein, isoform K OS=Drosophila melanogaster OX=7227 GN=CG14632 PE=4 SV=1 | CG14632 |
| Q9W5G7 | 11 | S | 2.54 | 0.001508322 | Up | EG:BACR37P7.8 protein OS=Drosophila melanogaster OX=7227 GN=EG:BACR37P7.8 PE=4 SV=2 | EG:BACR37P7.8 |
| Q9W5G7 | 15 | S | 1.605 | 0.009299556 | Up | EG:BACR37P7.8 protein OS=Drosophila melanogaster OX=7227 GN=EG:BACR37P7.8 PE=4 SV=2 | EG:BACR37P7.8 |
| X2JFT2 | 22 | S | 2.581 | 0.002041071 | Up | Gone early, isoform B OS=Drosophila melanogaster OX=7227 GN=goe PE=1 SV=1 | goe |
| Q9Y095 | 396 | S | 1.35 | 0.000978054 | Up | FI04011p OS=Drosophila melanogaster OX=7227 GN=XRCC1 PE=1 SV=1 | XRCC1 |
| Q9Y095 | 180 | S | 1.694 | 0.002528981 | Up | FI04011p OS=Drosophila melanogaster OX=7227 GN=XRCC1 PE=1 SV=1 | XRCC1 |
| Q9Y095 | 562 | S | 1.561 | 0.018544602 | Up | FI04011p OS=Drosophila melanogaster OX=7227 GN=XRCC1 PE=1 SV=1 | XRCC1 |
| Q9Y095 | 250 | S | 1.348 | 0.000559023 | Up | FI04011p OS=Drosophila melanogaster OX=7227 GN=XRCC1 PE=1 SV=1 | XRCC1 |
| Q9Y119 | 45 | S | 0.493 | 6.79508E-05 | Down | BcDNA.GH08860 OS=Drosophila melanogaster OX=7227 GN=Tps1 PE=1 SV=1 | Tps1 |
| Q9Y119 | 345 | S | 0.607 | 0.044696844 | Down | BcDNA.GH08860 OS=Drosophila melanogaster OX=7227 GN=Tps1 PE=1 SV=1 | Tps1 |
| Q9Y128 | 143 | S | 1.482 | 0.009122636 | Up | BcDNA.GH07688 OS=Drosophila melanogaster OX=7227 GN=cert PE=1 SV=1 | cert |
| Q9Y165 | 169 | S | 1.562 | 5.38942E-05 | Up | BcDNA.GH02435 OS=Drosophila melanogaster OX=7227 GN=morgue PE=1 SV=1 | morgue |
| Q9Y1A7 | 32 | S | 1.383 | 0.003247962 | Up | LD25378p OS=Drosophila melanogaster OX=7227 GN=mnd PE=2 SV=1 | mnd |
| Q9Y1A7 | 35 | S | 1.54 | 0.000765242 | Up | LD25378p OS=Drosophila melanogaster OX=7227 GN=mnd PE=2 SV=1 | mnd |
| X2JAM4 | 911 | S | 6.009 | 1.69623E-05 | Up | Supervillin, isoform AE OS=Drosophila melanogaster OX=7227 GN=Svil PE=1 SV=1 | Svil |
| X2JAU8 | 482 | S | 0.22 | 3.16432E-06 | Down | Protein nervous wreck OS=Drosophila melanogaster OX=7227 GN=nwk PE=1 SV=1 | nwk |
| X2JAU8 | 484 | S | 0.22 | 3.16432E-06 | Down | Protein nervous wreck OS=Drosophila melanogaster OX=7227 GN=nwk PE=1 SV=1 | nwk |
| X2JAU8 | 436 | S | 1.588 | 0.008393336 | Up | Protein nervous wreck OS=Drosophila melanogaster OX=7227 GN=nwk PE=1 SV=1 | nwk |
| X2JAU8 | 622 | S | 1.812 | 0.002787292 | Up | Protein nervous wreck OS=Drosophila melanogaster OX=7227 GN=nwk PE=1 SV=1 | nwk |
| X2JAW9 | 54 | S | 2.271 | 0.000202478 | Up | Inwardly rectifying potassium channel 3, isoform C OS=Drosophila melanogaster OX=7227 GN=Irk3 PE=3 SV=1 | Irk3 |

| | | | | | | | | |
|---|---|---|---|---|---|---|---|---|
| X2JAW9 | 63 | S | 3.357 | 6.82263E-05 | Up | Inwardly rectifying potassium channel 3, isoform C OS=Drosophila melanogaster OX=7227 GN=Irk3 PE=3 SV=1 | | Irk3 |
| X2JAW9 | 23 | S | 1.826 | 0.000949609 | Up | Inwardly rectifying potassium channel 3, isoform C OS=Drosophila melanogaster OX=7227 GN=Irk3 PE=3 SV=1 | | Irk3 |
| X2JFA0 | 211 | S | 1.446 | 0.02618416 | Up | AMP deaminase OS=Drosophila melanogaster OX=7227 GN=AMPdeam PE=1 SV=1 | | AMPdeam |
| X2JKC1 | 204 | S | 0.623 | 0.00359659 | Down | Lipid storage droplet-2, isoform E OS=Drosophila melanogaster OX=7227 GN=Lsd-2 PE=1 SV=1 | | Lsd-2 |
| E5DK16 | 105 | T | 2.912 | 0.006395169 | Up | Lipin isoform J OS=Drosophila melanogaster OX=7227 GN=Lpin PE=1 SV=1 | | Lpin |
| Q9VF03 | 345 | T | 1.434 | 0.02258974 | Up | Brahma associated protein 155 kDa OS=Drosophila melanogaster OX=7227 GN=mor PE=1 SV=3 | | mor |
| A0A0B4K6V2 | 311 | T | 1.554 | 0.001178015 | Up | Like-AP180, isoform G OS=Drosophila melanogaster OX=7227 GN=lap PE=1 SV=1 | | lap |
| Q9VC62 | 607 | T | 2.248 | 0.001041027 | Up | Extended synaptotagmin-like protein 2, isoform A OS=Drosophila melanogaster OX=7227 GN=Esyt2 PE=1 SV=1 | | Esyt2 |
| A1Z9J3 | 8285 | T | 1.303 | 0.023414345 | Up | Short stop, isoform H OS=Drosophila melanogaster OX=7227 GN=shot PE=1 SV=1 | | shot |
| A0A0B4K849 | 701 | T | 2.121 | 0.005633527 | Up | Hu li tai shao, isoform O OS=Drosophila melanogaster OX=7227 GN=hts PE=1 SV=1 | | hts |
| A0A0B4K849 | 586 | T | 1.523 | 0.002118878 | Up | Hu li tai shao, isoform O OS=Drosophila melanogaster OX=7227 GN=hts PE=1 SV=1 | | hts |
| A0A0B4K849 | 596 | T | 1.424 | 0.001378392 | Up | Hu li tai shao, isoform O OS=Drosophila melanogaster OX=7227 GN=hts PE=1 SV=1 | | hts |
| Q9VCH4 | 1872 | T | 1.851 | 0.001057061 | Up | Myoblast city OS=Drosophila melanogaster OX=7227 GN=mbc PE=1 SV=2 | | mbc |
| A1ZAN7 | 1678 | T | 1.571 | 0.000271531 | Up | Rho guanine nucleotide exchange factor 2, isoform E OS=Drosophila melanogaster OX=7227 GN=RhoGEF2 PE=1 SV=1 | | RhoGEF2 |
| A0A0B4KEH0 | 235 | T | 1.387 | 0.017887471 | Up | 14-3-3zeta, isoform K OS=Drosophila melanogaster OX=7227 GN=14-3-3zeta PE=1 SV=1 | | 14-3-3zeta |
| A0A0B4KEI6 | 92 | T | 2.427 | 0.004392113 | Up | Jun-related antigen, isoform C OS=Drosophila melanogaster OX=7227 GN=Jra PE=3 SV=1 | | Jra |
| Q7JQL5 | 62 | T | 1.774 | 0.000604925 | Up | Autophagy-related protein 9 OS=Drosophila melanogaster OX=7227 GN=Atg9 PE=1 SV=1 | | Atg9 |
| Q7JQL5 | 69 | T | 1.774 | 0.000604925 | Up | Autophagy-related protein 9 OS=Drosophila melanogaster OX=7227 GN=Atg9 PE=1 SV=1 | | Atg9 |
| A0A0B4KF90 | 298 | T | 1.673 | 0.037299126 | Up | Like-AP180, isoform J OS=Drosophila melanogaster OX=7227 GN=lap PE=1 SV=1 | | lap |
| A0A0B4KGP0 | 410 | T | 1.729 | 0.000479344 | Up | Stumps, isoform F OS=Drosophila melanogaster OX=7227 GN=stumps PE=4 SV=1 | | stumps |
| A0A126GUT4 | 44 | T | 1.33 | 0.0081336 | Up | Forkhead box, sub-group O, isoform H OS=Drosophila melanogaster OX=7227 GN=foxo PE=4 SV=1 | | foxo |
| A0A0B4KGU4 | 185 | T | 1.379 | 0.018218351 | Up | Belle, isoform B OS=Drosophila melanogaster OX=7227 GN=bel PE=4 SV=1 | | bel |
| A0A0B4KHD9 | 270 | T | 1.799 | 0.013280398 | Up | Uncharacterized protein, isoform E OS=Drosophila melanogaster OX=7227 GN=Dmel\CG5746 PE=4 SV=1 | | Dmel\CG5746 |
| A0A0B4KI69 | 613 | T | 1.334 | 0.002762537 | Up | G protein-coupled receptor kinase OS=Drosophila melanogaster OX=7227 GN=Gprk2 PE=3 SV=1 | | Gprk2 |
| A0A0B4LEW1 | 842 | T | 1.893 | 0.00146471 | Up | Uncharacterized protein, isoform E OS=Drosophila melanogaster OX=7227 GN=BcDNA:GH05582 PE=4 SV=1 | | BcDNA:GH05582 |
| A0A0B4LG01 | 514 | T | 1.542 | 0.007925467 | Up | Oysgedart, isoform D OS=Drosophila melanogaster OX=7227 GN=oys PE=3 SV=1 | | oys |
| A0A0B4LGB9 | 306 | T | 1.4 | 0.02725049 | Up | Quaking related 58E-1, isoform C OS=Drosophila melanogaster OX=7227 GN=qkr58E-1 PE=1 SV=1 | | qkr58E-1 |
| Q9W1G5 | 884 | T | 3.323 | 0.000897799 | Up | Kazachoc, isoform C OS=Drosophila melanogaster OX=7227 GN=kcc PE=1 SV=2 | | kcc |
| Q9VE88 | 535 | T | 1.432 | 0.004728975 | Up | Uncharacterized protein, isoform B OS=Drosophila melanogaster OX=7227 GN=lincRNA.804 PE=4 SV=3 | | lincRNA.804 |
| A0A0B4LIJ0 | 433 | T | 0.55 | 5.04746E-05 | Down | Regulatory particle triple-A ATPase 2, isoform B OS=Drosophila melanogaster OX=7227 GN=Rpt2 PE=3 SV=1 | | Rpt2 |
| A0A126GUN6 | 239 | T | 3.155 | 0.002330303 | Up | Z band alternatively spliced PDZ-motif protein 52, isoform X OS=Drosophila melanogaster OX=7227 GN=Zasp52 PE=1 SV=1 | | Zasp52 |
| E1JGZ4 | 319 | T | 1.305 | 0.00314418 | Up | Blown fuse, isoform B OS=Drosophila melanogaster OX=7227 GN=blow PE=4 SV=1 | | blow |
| A1Z8N1 | 374 | T | 2.341 | 0.02327959 | Up | Facilitated trehalose transporter Tret1-1 OS=Drosophila melanogaster OX=7227 GN=Tret1-1 PE=1 SV=1 | | Tret1-1 |
| Q8SX22 | 514 | T | 1.457 | 0.015825271 | Up | RE68725p OS=Drosophila melanogaster OX=7227 GN=Syn2 PE=2 SV=1 | | Syn2 |
| B7YZI7 | 537 | T | 1.532 | 0.001090885 | Up | Uncharacterized protein, isoform G OS=Drosophila melanogaster OX=7227 GN=anon-WO0140519.237 PE=4 SV=1 | | anon-WO0140519.237 |
| A1ZAX6 | 620 | T | 1.309 | 0.00021692 | Up | FI02012p OS=Drosophila melanogaster OX=7227 GN=HPS4 PE=1 SV=1 | | HPS4 |
| A8E6M1 | 522 | T | 1.387 | 0.000365937 | Up | GH01011p OS=Drosophila melanogaster OX=7227 GN=CG5477 PE=1 SV=1 | | CG5477 |
| E1JGN0 | 907 | T | 1.944 | 0.007041698 | Up | Non-specific serine/threonine protein kinase OS=Drosophila melanogaster OX=7227 GN=par-1 PE=1 SV=1 | | par-1 |
| A4UZL3 | 317 | T | 1.516 | 0.008808984 | Up | Eps15 homology domain containing protein-binding protein 1, isoform F OS=Drosophila melanogaster OX=7227 GN=Ehbp1 PE=1 SV=1 | | Ehbp1 |
| D1YSG7 | 352 | T | 1.9 | 0.002240249 | Up | Calcium/calmodulin-dependent protein kinase II, isoform I OS=Drosophila melanogaster OX=7227 GN=CaMKII PE=1 SV=1 | | CaMKII |
| A8JMD5 | 659 | T | 1.463 | 0.003020097 | Up | Black match, isoform D OS=Drosophila melanogaster OX=7227 GN=bma PE=4 SV=2 | | bma |
| M9PEM7 | 381 | T | 2.125 | 0.000555893 | Up | Liquid facets, isoform G OS=Drosophila melanogaster OX=7227 GN=lqf PE=1 SV=1 | | lqf |
| A8JQT5 | 60 | T | 1.54 | 0.01234736 | Up | Chondrocyte-derived ezrin-like domain containing protein, isoform G OS=Drosophila melanogaster OX=7227 GN=Cdep PE=1 SV=2 | | Cdep |
| Q9VE16 | 514 | T | 1.365 | 0.03122871 | Up | GUK-holder, isoform B OS=Drosophila melanogaster OX=7227 GN=gukh PE=2 SV=2 | | gukh |
| M9PHE6 | 71 | T | 2.343 | 0.000408695 | Up | Uncharacterized protein, isoform I OS=Drosophila melanogaster OX=7227 GN=CG1521 PE=4 SV=1 | | CG1521 |
| B5RJS0 | 524 | T | 0.727 | 0.004263203 | Down | IP20241p OS=Drosophila melanogaster OX=7227 GN=Nost PE=1 SV=1 | | Nost |
| B7YZQ1 | 1470 | T | 1.327 | 0.004579253 | Up | Sterile20-like kinase, isoform F OS=Drosophila melanogaster OX=7227 GN=Slik PE=1 SV=1 | | Slik |
| B7YZQ1 | 1474 | T | 1.514 | 0.018283752 | Up | Sterile20-like kinase, isoform F OS=Drosophila melanogaster OX=7227 GN=Slik PE=1 SV=1 | | Slik |
| B7YZQ1 | 505 | T | 1.342 | 0.043572592 | Up | Sterile20-like kinase, isoform F OS=Drosophila melanogaster OX=7227 GN=Slik PE=1 SV=1 | | Slik |
| B7YZQ3 | 973 | T | 1.554 | 0.025374428 | Up | Prominin, isoform D OS=Drosophila melanogaster OX=7227 GN=prom PE=1 SV=1 | | prom |
| M9PDB4 | 289 | T | 1.41 | 0.036512946 | Up | Cytoplasmic linker protein 190, isoform Q OS=Drosophila melanogaster OX=7227 GN=CLIP-190 PE=1 SV=1 | | CLIP-190 |
| Q7KUB0 | 159 | T | 0.665 | 0.047111472 | Down | Isocitrate dehydrogenase [NADP] OS=Drosophila melanogaster OX=7227 GN=Idh PE=1 SV=1 | | Idh |
| Q9V3J6 | 613 | T | 2.014 | 0.000200322 | Up | Bestrophin homolog OS=Drosophila melanogaster OX=7227 GN=Best1 PE=2 SV=1 | | Best1 |
| C8VV14 | 39 | T | 0.737 | 0.007199959 | Down | Fructose-bisphosphate aldolase OS=Drosophila melanogaster OX=7227 GN=Ald1 PE=1 SV=1 | | Ald1 |
| E1JGM2 | 325 | T | 1.644 | 0.001482141 | Up | 5-hydroxytryptamine (Serotonin) receptor 1B, isoform D OS=Drosophila melanogaster OX=7227 GN=5-HT1B PE=3 SV=2 | | 5-HT1B |
| E1JHT2 | 73 | T | 0.743 | 0.00520383 | Down | Transporter OS=Drosophila melanogaster OX=7227 GN=ine PE=3 SV=1 | | ine |
| E2QCY9 | 981 | T | 1.499 | 0.005918182 | Up | Synapsin, isoform D OS=Drosophila melanogaster OX=7227 GN=Syn PE=1 SV=1 | | Syn |
| E2QCY9 | 119 | T | 2.116 | 0.008697509 | Up | Synapsin, isoform D OS=Drosophila melanogaster OX=7227 GN=Syn PE=1 SV=1 | | Syn |
| E2QD75 | 9 | T | 1.516 | 0.004821659 | Up | Cap binding protein 80, isoform B OS=Drosophila melanogaster OX=7227 GN=Cbp80 PE=1 SV=1 | | Cbp80 |
| F6J1D0 | 21 | T | 2.136 | 0.003114808 | Up | CG3595 OS=Drosophila melanogaster OX=7227 GN=sqh PE=1 SV=1 | | sqh |

| | | | | | | | |
|---|---|---|---|---|---|---|---|
| H8F4P8 | 519 | T | 2.812 | 0.028688078 | Up | Phosphodiesterase OS=Drosophila melanogaster OX=7227 GN=dnc PE=1 SV=1 | dnc |
| L0MLK7 | 1411 | T | 1.345 | 0.000411208 | Up | Zn finger homeodomain 2, isoform C OS=Drosophila melanogaster OX=7227 GN=zfh2 PE=4 SV=1 | zfh2 |
| X2JDP6 | 265 | T | 2.708 | 0.00325435 | Up | Nucleosome-destabilizing factor, isoform C OS=Drosophila melanogaster OX=7227 GN=Ndf PE=4 SV=1 | Ndf |
| M9MRJ4 | 2293 | T | 1.621 | 0.002813931 | Up | Muscle-specific protein 300 kDa, isoform G OS=Drosophila melanogaster OX=7227 GN=Msp300 PE=1 SV=1 | Msp300 |
| M9MRX4 | 2166 | T | 1.558 | 0.017373127 | Up | Ankyrin 2, isoform U OS=Drosophila melanogaster OX=7227 GN=Ank2 PE=1 SV=1 | Ank2 |
| M9MRX4 | 2213 | T | 1.509 | 0.016603176 | Up | Ankyrin 2, isoform U OS=Drosophila melanogaster OX=7227 GN=Ank2 PE=1 SV=1 | Ank2 |
| M9MRX4 | 1404 | T | 1.67 | 0.026822078 | Up | Ankyrin 2, isoform U OS=Drosophila melanogaster OX=7227 GN=Ank2 PE=1 SV=1 | Ank2 |
| M9MRX4 | 3221 | T | 1.654 | 0.000323482 | Up | Ankyrin 2, isoform U OS=Drosophila melanogaster OX=7227 GN=Ank2 PE=1 SV=1 | Ank2 |
| M9MRX4 | 3225 | T | 1.525 | 0.000258409 | Up | Ankyrin 2, isoform U OS=Drosophila melanogaster OX=7227 GN=Ank2 PE=1 SV=1 | Ank2 |
| M9MRX4 | 5067 | T | 0.741 | 0.02070789 | Down | Ankyrin 2, isoform U OS=Drosophila melanogaster OX=7227 GN=Ank2 PE=1 SV=1 | Ank2 |
| M9MRX4 | 3203 | T | 1.546 | 0.005394967 | Up | Ankyrin 2, isoform U OS=Drosophila melanogaster OX=7227 GN=Ank2 PE=1 SV=1 | Ank2 |
| M9MRX4 | 5447 | T | 0.487 | 0.000633387 | Down | Ankyrin 2, isoform U OS=Drosophila melanogaster OX=7227 GN=Ank2 PE=1 SV=1 | Ank2 |
| M9MS14 | 238 | T | 3.301 | 0.002670612 | Up | Moody, isoform C OS=Drosophila melanogaster OX=7227 GN=moody PE=3 SV=1 | moody |
| M9MS15 | 259 | T | 1.663 | 0.0014524 | Up | Neurotactin, isoform C OS=Drosophila melanogaster OX=7227 GN=Nrt PE=4 SV=1 | Nrt |
| M9MS31 | 1292 | T | 0.758 | 0.000194806 | Down | Diacylglycerol kinase OS=Drosophila melanogaster OX=7227 GN=rdgA PE=3 SV=1 | rdgA |
| M9PGT0 | 2860 | T | 1.354 | 0.000785718 | Up | Uncharacterized protein, isoform R OS=Drosophila melanogaster OX=7227 GN=CG3960 PE=1 SV=1 | CG3960 |
| M9PGT0 | 1294 | T | 1.787 | 0.003758559 | Up | Uncharacterized protein, isoform R OS=Drosophila melanogaster OX=7227 GN=CG3960 PE=1 SV=1 | CG3960 |
| Q86B48 | 137 | T | 7.584 | 5.12685E-06 | Up | Forked, isoform G OS=Drosophila melanogaster OX=7227 GN=f PE=4 SV=2 | f |
| M9NCM6 | 2274 | T | 1.332 | 0.001876988 | Up | Split ends, isoform D OS=Drosophila melanogaster OX=7227 GN=spen PE=1 SV=1 | spen |
| M9NDP0 | 1881 | T | 1.387 | 0.007378635 | Up | Uncharacterized protein, isoform C OS=Drosophila melanogaster OX=7227 GN=elF-4A PE=4 SV=1 | elF-4A |
| M9NF47 | 1183 | T | 2.005 | 0.002191009 | Up | Uncharacterized protein, isoform J OS=Drosophila melanogaster OX=7227 GN=CG32355 PE=1 SV=1 | CG32355 |
| X2J8V5 | 148 | T | 0.45 | 0.007201343 | Down | Arrestin 1, isoform C OS=Drosophila melanogaster OX=7227 GN=Arr1 PE=4 SV=1 | Arr1 |
| M9PBP8 | 1925 | T | 1.638 | 0.00037264 | Up | Axotactin, isoform F OS=Drosophila melanogaster OX=7227 GN=axo PE=4 SV=1 | axo |
| M9NGG5 | 5082 | T | 1.324 | 0.000635882 | Up | Futsch, isoform F OS=Drosophila melanogaster OX=7227 GN=futsch PE=1 SV=1 | futsch |
| M9NGG5 | 1472 | T | 1.348 | 0.002534691 | Up | Futsch, isoform F OS=Drosophila melanogaster OX=7227 GN=futsch PE=1 SV=1 | futsch |
| M9NGG5 | 1637 | T | 0.659 | 0.01794307 | Down | Futsch, isoform F OS=Drosophila melanogaster OX=7227 GN=futsch PE=1 SV=1 | futsch |
| M9NGG5 | 3553 | T | 1.782 | 0.01254404 | Up | Futsch, isoform F OS=Drosophila melanogaster OX=7227 GN=futsch PE=1 SV=1 | futsch |
| M9NGG5 | 4939 | T | 0.767 | 0.017719641 | Down | Futsch, isoform F OS=Drosophila melanogaster OX=7227 GN=futsch PE=1 SV=1 | futsch |
| M9NGG5 | 4180 | T | 1.759 | 0.004452645 | Up | Futsch, isoform F OS=Drosophila melanogaster OX=7227 GN=futsch PE=1 SV=1 | futsch |
| M9NGG5 | 3181 | T | 1.825 | 0.001097553 | Up | Futsch, isoform F OS=Drosophila melanogaster OX=7227 GN=futsch PE=1 SV=1 | futsch |
| M9NGG5 | 1156 | T | 0.658 | 0.003217603 | Down | Futsch, isoform F OS=Drosophila melanogaster OX=7227 GN=futsch PE=1 SV=1 | futsch |
| M9NGG5 | 2797 | T | 1.305 | 0.001038099 | Up | Futsch, isoform F OS=Drosophila melanogaster OX=7227 GN=futsch PE=1 SV=1 | futsch |
| M9PAY9 | 586 | T | 1.311 | 0.000857413 | Up | Uncharacterized protein, isoform B OS=Drosophila melanogaster OX=7227 GN=Dmel\CG4133 PE=4 SV=1 | Dmel\CG4133 |
| M9PDS3 | 5730 | T | 5.961 | 5.37929E-06 | Up | Sallimus, isoform T OS=Drosophila melanogaster OX=7227 GN=sls PE=4 SV=1 | sls |
| M9PBL3 | 332 | T | 1.605 | 0.000136525 | Up | Heat shock protein 83, isoform B OS=Drosophila melanogaster OX=7227 GN=Hsp83 PE=3 SV=1 | Hsp83 |
| M9PBW9 | 150 | T | 1.955 | 0.023991858 | Up | Rhea, isoform G OS=Drosophila melanogaster OX=7227 GN=rhea PE=1 SV=1 | rhea |
| M9PBW9 | 152 | T | 1.955 | 0.023991858 | Up | Rhea, isoform G OS=Drosophila melanogaster OX=7227 GN=rhea PE=1 SV=1 | rhea |
| M9PI37 | 1121 | T | 5.264 | 0.001083515 | Up | Sodium chloride cotransporter 69, isoform E OS=Drosophila melanogaster OX=7227 GN=Ncc69 PE=1 SV=1 | Ncc69 |
| M9PEG1 | 678 | T | 0.763 | 0.03438059 | Down | Uncharacterized protein OS=Drosophila melanogaster OX=7227 GN=Dmel\CG44195 PE=1 SV=1 | Dmel\CG44195 |
| Q960V1 | 227 | T | 1.312 | 0.049692198 | Up | LD33361p OS=Drosophila melanogaster OX=7227 GN=simj PE=1 SV=1 | simj |
| M9PFF9 | 222 | T | 1.315 | 0.003769548 | Up | Trailer hitch, isoform D OS=Drosophila melanogaster OX=7227 GN=tral PE=1 SV=1 | tral |
| Q00963 | 1842 | T | 1.48 | 0.002004103 | Up | Spectrin beta chain OS=Drosophila melanogaster OX=7227 GN=beta-Spec PE=1 SV=2 | beta-Spec |
| M9PF68 | 260 | T | 1.586 | 0.036688747 | Up | Kinesin-like protein OS=Drosophila melanogaster OX=7227 GN=Klp68D PE=3 SV=1 | Klp68D |
| M9PFS3 | 609 | T | 1.356 | 0.000999953 | Up | Sugar-free frosting, isoform B OS=Drosophila melanogaster OX=7227 GN=sff PE=4 SV=1 | sff |
| M9PFU0 | 527 | T | 1.315 | 0.003734712 | Up | Uncharacterized protein, isoform A OS=Drosophila melanogaster OX=7227 GN=CG15639 PE=4 SV=1 | CG15639 |
| M9PG50 | 184 | T | 1.776 | 1.23651E-05 | Up | Cyclin-dependent kinase 12, isoform C OS=Drosophila melanogaster OX=7227 GN=Cdk12 PE=4 SV=1 | Cdk12 |
| M9PGC5 | 250 | T | 1.367 | 0.002403932 | Up | Wnk kinase, isoform M OS=Drosophila melanogaster OX=7227 GN=Wnk PE=1 SV=2 | Wnk |
| M9PGF7 | 178 | T | 1.477 | 0.002837331 | Up | Tenascin major, isoform E OS=Drosophila melanogaster OX=7227 GN=Ten-m PE=1 SV=1 | Ten-m |
| M9PHJ0 | 8 | T | 0.563 | 0.03627413 | Down | Upheld, isoform P OS=Drosophila melanogaster OX=7227 GN=up PE=1 SV=1 | up |
| M9PJM6 | 8 | T | 0.517 | 0.015333973 | Down | Upheld, isoform Q OS=Drosophila melanogaster OX=7227 GN=up PE=1 SV=1 | up |
| O76928 | 282 | T | 0.739 | 0.000142375 | Down | Carmine, isoform A OS=Drosophila melanogaster OX=7227 GN=cm PE=1 SV=1 | cm |
| X2JD55 | 184 | T | 4.708 | 0.002301687 | Up | Yolk protein 1, isoform B OS=Drosophila melanogaster OX=7227 GN=Yp1 PE=1 SV=1 | Yp1 |
| P07668 | 75 | T | 2.772 | 0.016850493 | Up | Choline O-acetyltransferase OS=Drosophila melanogaster OX=7227 GN=ChAT PE=1 SV=3 | ChAT |
| P13677 | 673 | T | 0.757 | 0.003769511 | Down | Protein kinase C, eye isozyme OS=Drosophila melanogaster OX=7227 GN=inaC PE=1 SV=1 | inaC |
| P14199 | 364 | T | 2.195 | 0.000476824 | Up | Protein ref(2)P OS=Drosophila melanogaster OX=7227 GN=ref(2)P PE=1 SV=2 | ref(2)P |
| P14199 | 305 | T | 2.625 | 0.000338287 | Up | Protein ref(2)P OS=Drosophila melanogaster OX=7227 GN=ref(2)P PE=1 SV=2 | ref(2)P |
| P18432 | 22 | T | 1.413 | 0.021136468 | Up | Myosin regulatory light chain 2 OS=Drosophila melanogaster OX=7227 GN=Mlc2 PE=1 SV=2 | Mlc2 |
| P19334 | 864 | T | 2.313 | 0.000165774 | Up | Transient receptor potential protein OS=Drosophila melanogaster OX=7227 GN=trp PE=1 SV=3 | trp |
| Q540V7 | 82 | T | 3.242 | 0.001265969 | Up | LD06131p OS=Drosophila melanogaster OX=7227 GN=rho PE=2 SV=1 | rho |

| | | | | | | | |
|---|---|---|---|---|---|---|---|
| X2JJG8 | 52 | T | 1.991 | 4.03077E-05 | Up | Glutamine synthetase OS=Drosophila melanogaster OX=7227 GN=Gs2 PE=1 SV=1 | Gs2 |
| X2JJG8 | 113 | T | 2.244 | 0.00028685 | Up | Glutamine synthetase OS=Drosophila melanogaster OX=7227 GN=Gs2 PE=1 SV=1 | Gs2 |
| X2JJG8 | 117 | T | 2.106 | 0.000358394 | Up | Glutamine synthetase OS=Drosophila melanogaster OX=7227 GN=Gs2 PE=1 SV=1 | Gs2 |
| P34082 | 841 | T | 1.421 | 0.00276204 | Up | Fasciclin-2 OS=Drosophila melanogaster OX=7227 GN=Fas2 PE=1 SV=1 | Fas2 |
| P40417 | 198 | T | 1.464 | 0.023290618 | Up | Mitogen-activated protein kinase ERK-A OS=Drosophila melanogaster OX=7227 GN=rl PE=1 SV=3 | rl |
| P48596 | 67 | T | 1.508 | 0.000201321 | Up | GTP cyclohydrolase 1 OS=Drosophila melanogaster OX=7227 GN=Pu PE=1 SV=3 | Pu |
| X2JC55 | 65 | T | 1.457 | 0.003315425 | Up | Phosphatidate cytidylyltransferase OS=Drosophila melanogaster OX=7227 GN=Cds PE=1 SV=1 | Cds |
| P83097 | 448 | T | 1.587 | 0.003175028 | Up | Putative tyrosine-protein kinase Wsck OS=Drosophila melanogaster OX=7227 GN=Wsck PE=2 SV=2 | Wsck |
| Q03427 | 457 | T | 2.774 | 0.008529346 | Up | Lamin-C OS=Drosophila melanogaster OX=7227 GN=LamC PE=1 SV=2 | LamC |
| Q03427 | 443 | T | 1.717 | 0.002535603 | Up | Lamin-C OS=Drosophila melanogaster OX=7227 GN=LamC PE=1 SV=2 | LamC |
| Q03427 | 7 | T | 2.312 | 0.004522404 | Up | Lamin-C OS=Drosophila melanogaster OX=7227 GN=LamC PE=1 SV=2 | LamC |
| Q03427 | 597 | T | 1.563 | 4.38459E-05 | Up | Lamin-C OS=Drosophila melanogaster OX=7227 GN=LamC PE=1 SV=2 | LamC |
| Q0KID3 | 427 | T | 1.331 | 0.007643624 | Up | Uncharacterized protein, isoform E OS=Drosophila melanogaster OX=7227 GN=CG9818 PE=3 SV=2 | CG9818 |
| X2JCI8 | 549 | T | 1.399 | 0.02897833 | Up | Msr-110, isoform D OS=Drosophila melanogaster OX=7227 GN=Msr-110 PE=1 SV=1 | Msr-110 |
| Q58CJ5 | 257 | T | 1.308 | 0.010159493 | Up | GH01093p OS=Drosophila melanogaster OX=7227 GN=sals PE=1 SV=1 | sals |
| Q7K1C5 | 388 | T | 2.203 | 0.009262173 | Up | GH21176p OS=Drosophila melanogaster OX=7227 GN=Hyccin PE=1 SV=1 | Hyccin |
| Q7K1L4 | 333 | T | 1.561 | 0.003974861 | Up | SD10469p OS=Drosophila melanogaster OX=7227 GN=Dmel\CG8468 PE=1 SV=1 | Dmel\CG8468 |
| Q7K204 | 555 | T | 1.427 | 0.001228012 | Up | Barricade, isoform A OS=Drosophila melanogaster OX=7227 GN=barc PE=1 SV=1 | barc |
| Q7KTJ7 | 635 | T | 1.403 | 0.004706048 | Up | Basigin, isoform G OS=Drosophila melanogaster OX=7227 GN=Bsg PE=1 SV=1 | Bsg |
| Q7KU01 | 648 | T | 1.309 | 0.000840389 | Up | PNUTS, isoform D OS=Drosophila melanogaster OX=7227 GN=PNUTS PE=1 SV=1 | PNUTS |
| Q7KV34 | 190 | T | 0.693 | 0.004181924 | Down | CHK domain-containing protein OS=Drosophila melanogaster OX=7227 GN=Dmel\CG1561 PE=1 SV=1 | Dmel\CG1561 |
| Q8IRI5 | 329 | T | 1.648 | 0.025533783 | Up | Trio, isoform D OS=Drosophila melanogaster OX=7227 GN=trio PE=1 SV=1 | trio |
| Q8IMF3 | 222 | T | 0.76 | 0.018211051 | Down | Uncharacterized protein, isoform C OS=Drosophila melanogaster OX=7227 GN=Dmel\CG2201 PE=4 SV=1 | Dmel\CG2201 |
| Q8IMG1 | 274 | T | 1.498 | 0.018301404 | Up | Uncharacterized protein, isoform C OS=Drosophila melanogaster OX=7227 GN=anon-WO0118547.745 PE=1 SV=1 | anon-WO0118547.745 |
| Q8IMY7 | 418 | T | 2.055 | 0.000779119 | Up | Inwardly rectifying potassium channel 2, isoform B OS=Drosophila melanogaster OX=7227 GN=Irk2 PE=3 SV=1 | Irk2 |
| Q8IPG9 | 260 | T | 1.348 | 0.008413333 | Up | Basigin, isoform A OS=Drosophila melanogaster OX=7227 GN=Bsg PE=1 SV=1 | Bsg |
| Q8IQQ7 | 748 | T | 1.74 | 0.033139331 | Up | Bloated tubules, isoform B OS=Drosophila melanogaster OX=7227 GN=blot PE=2 SV=1 | blot |
| Q8MLQ7 | 284 | T | 1.608 | 0.001355231 | Up | RH09188p OS=Drosophila melanogaster OX=7227 GN=Dmel\CG4797 PE=2 SV=2 | Dmel\CG4797 |
| Q8MLS1 | 8 | T | 0.452 | 0.008353796 | Down | 5-formyltetrahydrofolate cyclo-ligase OS=Drosophila melanogaster OX=7227 GN=Mthfs PE=3 SV=1 | Mthfs |
| Q8MMD2 | 915 | T | 1.914 | 0.04295554 | Up | Epidermal growth factor receptor pathway substrate clone 15, isoform B OS=Drosophila melanogaster OX=7227 GN=Eps-15 PE=1 SV=1 | Eps-15 |
| Q8SWV5 | 580 | T | 2.407 | 3.35876E-05 | Up | LD47995p OS=Drosophila melanogaster OX=7227 GN=dSLC5A3 PE=1 SV=1 | dSLC5A3 |
| Q95RI2 | 36 | T | 1.431 | 0.004382737 | Up | LD28549p OS=Drosophila melanogaster OX=7227 GN=Dmel\CG10973 PE=1 SV=1 | Dmel\CG10973 |
| Q961J5 | 582 | T | 1.731 | 0.000403142 | Up | Beta-alanine transporter OS=Drosophila melanogaster OX=7227 GN=Balat PE=2 SV=1 | Balat |
| Q9TVP3 | 107 | T | 1.632 | 0.018884941 | Up | J domain-containing protein OS=Drosophila melanogaster OX=7227 GN=jdp PE=2 SV=2 | jdp |
| Q9V4W1 | 47 | T | 3.454 | 0.000348504 | Up | Nucleoporin Gle1 OS=Drosophila melanogaster OX=7227 GN=Gle1 PE=2 SV=1 | Gle1 |
| Q9VDD2 | 338 | T | 1.608 | 0.030845876 | Up | LD22662p OS=Drosophila melanogaster OX=7227 GN=SNF4Agamma PE=1 SV=2 | SNF4Agamma |
| Q9VG84 | 654 | T | 1.318 | 0.004523233 | Up | DNA ligase OS=Drosophila melanogaster OX=7227 GN=DNAlig3 PE=1 SV=3 | DNAlig3 |
| Q9VJ12 | 319 | T | 1.322 | 0.011577268 | Up | Acinus, isoform A OS=Drosophila melanogaster OX=7227 GN=Acn PE=1 SV=1 | Acn |
| X2J9M9 | 408 | T | 1.721 | 0.001933819 | Up | Phospholipid-transporting ATPase OS=Drosophila melanogaster OX=7227 GN=P4-ATPase PE=1 SV=1 | P4-ATPase |
| Q9VMI2 | 35 | T | 1.341 | 0.013693551 | Up | GEO08832p1 OS=Drosophila melanogaster OX=7227 GN=Dmel\CG13994 PE=2 SV=1 | Dmel\CG13994 |
| Q9VML2 | 3303 | T | 1.459 | 0.048476378 | Up | Blue cheese OS=Drosophila melanogaster OX=7227 GN=bchs PE=4 SV=4 | bchs |
| Q9VN21 | 479 | T | 1.508 | 0.045067511 | Up | LD30155p OS=Drosophila melanogaster OX=7227 GN=lost PE=1 SV=1 | lost |
| Q9VN21 | 370 | T | 1.401 | 0.000338847 | Up | LD30155p OS=Drosophila melanogaster OX=7227 GN=lost PE=1 SV=1 | lost |
| Q9VN77 | 89 | T | 1.311 | 6.16677E-05 | Up | LD31216p OS=Drosophila melanogaster OX=7227 GN=RabGGTalpha PE=1 SV=1 | RabGGTalpha |
| Q9VPX6 | 326 | T | 1.503 | 0.000442246 | Up | Adenylyl cyclase-associated protein OS=Drosophila melanogaster OX=7227 GN=capt PE=1 SV=2 | capt |
| Q9VQF7 | 54 | T | 1.388 | 0.018420077 | Up | Bacchus OS=Drosophila melanogaster OX=7227 GN=Bacc PE=2 SV=1 | Bacc |
| Q9VQV9 | 168 | T | 1.572 | 0.003243824 | Up | Cappuccino, isoform D OS=Drosophila melanogaster OX=7227 GN=capu PE=4 SV=2 | capu |
| Q9VQZ0 | 13 | T | 1.45 | 0.000498815 | Up | Protein YIPF OS=Drosophila melanogaster OX=7227 GN=Dmel\CG3652 PE=1 SV=2 | Dmel\CG3652 |
| Q9VQZ3 | 748 | T | 1.325 | 0.00355695 | Up | Uncharacterized protein, isoform A OS=Drosophila melanogaster OX=7227 GN=Dmel\CG15431 PE=4 SV=1 | Dmel\CG15431 |
| Q9VR44 | 482 | T | 1.481 | 0.000331607 | Up | GH05102p OS=Drosophila melanogaster OX=7227 GN=CT10168 PE=1 SV=1 | CT10168 |
| Q9VS39 | 36 | T | 1.759 | 0.043808009 | Up | FI19525p1 OS=Drosophila melanogaster OX=7227 GN=Dmel\CG14830 PE=2 SV=1 | Dmel\CG14830 |
| Q9VSK4 | 149 | T | 1.303 | 0.040738507 | Up | GM13032p OS=Drosophila melanogaster OX=7227 GN=Dmel\CG6983 PE=1 SV=1 | Dmel\CG6983 |
| Q9VSW5 | 744 | T | 1.358 | 0.029160276 | Up | Kinesin-like protein at 67A, isoform A OS=Drosophila melanogaster OX=7227 GN=Klp67A PE=1 SV=1 | Klp67A |
| Q9VTC3 | 25 | T | 0.7 | 0.003132041 | Down | GH07049p OS=Drosophila melanogaster OX=7227 GN=Dmel\CG6409 PE=1 SV=2 | Dmel\CG6409 |
| Q9VTW8 | 906 | T | 2.08 | 5.68553E-05 | Up | GH27027p OS=Drosophila melanogaster OX=7227 GN=Ncc69 PE=1 SV=1 | Ncc69 |
| Q9VW22 | 1263 | T | 1.66 | 0.000589837 | Up | FI18195p1 OS=Drosophila melanogaster OX=7227 GN=l(3)76BDm PE=1 SV=2 | l(3)76BDm |
| Q9VWJ0 | 303 | T | 1.652 | 0.000913397 | Up | SD10554p OS=Drosophila melanogaster OX=7227 GN=anon-WO03002137.1 PE=2 SV=2 | anon-WO03002137.1 |
| Q9VWK0 | 299 | T | 2.029 | 0.001699036 | Up | IP13170p OS=Drosophila melanogaster OX=7227 GN=Dmel\CG14196 PE=2 SV=1 | Dmel\CG14196 |
| Q9VWK2 | 116 | T | 2.427 | 0.001233728 | Up | Mucin 18B, isoform A OS=Drosophila melanogaster OX=7227 GN=Muc18B PE=2 SV=1 | Muc18B |
| Q9VWK2 | 123 | T | 2.427 | 0.001233728 | Up | Mucin 18B, isoform A OS=Drosophila melanogaster OX=7227 GN=Muc18B PE=2 SV=1 | Muc18B |
| Q9VWN4 | 371 | T | 1.406 | 0.00155741 | Up | Fl(2)d-associated complex component OS=Drosophila melanogaster OX=7227 GN=Flacc PE=1 SV=1 | Flacc |
| Q9VWP5 | 914 | T | 1.965 | 0.003984786 | Up | Hat-trick, isoform D OS=Drosophila melanogaster OX=7227 GN=htk PE=1 SV=4 | htk |
| Q9VWP8 | 840 | T | 1.948 | 0.017314984 | Up | Uncharacterized protein, isoform A OS=Drosophila melanogaster | CG32543 |

| | | | | | | | |
|---|---|---|---|---|---|---|---|
| | | | | | | OX=7227 GN=CG32543 PE=1 SV=2 | |
| Q9VX34 | 608 | T | 1.334 | 0.029902423 | Up | RNA helicase OS=Drosophila melanogaster OX=7227 GN=cg5800 PE=1 SV=2 | cg5800 |
| Q9VXV4 | 443 | T | 1.336 | 0.017983841 | Up | RH04535p OS=Drosophila melanogaster OX=7227 GN=Dmel\CG11655 PE=1 SV=1 | Dmel\CG11655 |
| Q9VYM7 | 322 | T | 1.591 | 0.001341345 | Up | Uncharacterized protein, isoform C OS=Drosophila melanogaster OX=7227 GN=Pripl4 PE=1 SV=2 | Pripl4 |
| Q9VYV4 | 281 | T | 1.327 | 0.025792997 | Up | Amun, isoform A OS=Drosophila melanogaster OX=7227 GN=Amun PE=1 SV=1 | Amun |
| Q9VYV4 | 250 | T | 1.408 | 0.006943477 | Up | Amun, isoform A OS=Drosophila melanogaster OX=7227 GN=Amun PE=1 SV=1 | Amun |
| Q9VYW6 | 52 | T | 1.399 | 0.015105841 | Up | Rhomboid-4 OS=Drosophila melanogaster OX=7227 GN=rho-4 PE=1 SV=2 | rho-4 |
| Q9W002 | 1009 | T | 1.442 | 0.001746061 | Up | Misshapen, isoform A OS=Drosophila melanogaster OX=7227 GN=msn PE=1 SV=3 | msn |
| Q9W019 | 114 | T | 1.882 | 0.00056097 | Up | IP09724p OS=Drosophila melanogaster OX=7227 GN=Dmel\CG15877 PE=1 SV=1 | Dmel\CG15877 |
| Q9W0L6 | 28 | T | 1.851 | 0.001616876 | Up | GH22266p OS=Drosophila melanogaster OX=7227 GN=Dmel\CG13907 PE=1 SV=1 | Dmel\CG13907 |
| Q9W1H5 | 223 | T | 1.411 | 0.005442928 | Up | Decapping protein 1, isoform A OS=Drosophila melanogaster OX=7227 GN=DCP1 PE=1 SV=1 | DCP1 |
| Q9W1K4 | 947 | T | 1.627 | 0.001541973 | Up | Egalitarian, isoform B OS=Drosophila melanogaster OX=7227 GN=egl PE=1 SV=4 | egl |
| Q9W266 | 645 | T | 2.524 | 0.001066536 | Up | Protein windpipe OS=Drosophila melanogaster OX=7227 GN=wdp PE=1 SV=1 | wdp |
| Q9W266 | 585 | T | 1.645 | 0.004164749 | Up | Protein windpipe OS=Drosophila melanogaster OX=7227 GN=wdp PE=1 SV=1 | wdp |
| Q9W2H0 | 466 | T | 2.04 | 2.05423E-05 | Up | Eukaryotic translation elongation factor, selenocysteine-specific OS=Drosophila melanogaster OX=7227 GN=eEFSec PE=2 SV=1 | eEFSec |
| Q9W2R3 | 591 | T | 2.006 | 0.010207804 | Up | LP03320p OS=Drosophila melanogaster OX=7227 GN=sktl PE=1 SV=2 | sktl |
| Q9W335 | 272 | T | 2.096 | 0.018377784 | Up | LD26546p OS=Drosophila melanogaster OX=7227 GN=l(1)G0320 PE=1 SV=1 | l(1)G0320 |
| Q9W3D3 | 1139 | T | 2.096 | 0.005595397 | Up | DENN domain-containing protein Crag OS=Drosophila melanogaster OX=7227 GN=Crag PE=1 SV=3 | Crag |
| Q9W3E2 | 288 | T | 1.587 | 0.042413852 | Up | PIP82 OS=Drosophila melanogaster OX=7227 GN=PIP82 PE=1 SV=3 | PIP82 |
| Q9W3N6 | 729 | T | 1.324 | 5.57812E-05 | Up | General vesicular transport factor p115 OS=Drosophila melanogaster OX=7227 GN=p115 PE=1 SV=2 | p115 |
| Q9W425 | 1365 | T | 1.441 | 0.043826709 | Up | Rabconnectin-3A OS=Drosophila melanogaster OX=7227 GN=Rbcn-3A PE=1 SV=3 | Rbcn-3A |
| Q9W425 | 2340 | T | 1.323 | 0.026326986 | Up | Rabconnectin-3A OS=Drosophila melanogaster OX=7227 GN=Rbcn-3A PE=1 SV=3 | Rbcn-3A |
| Q9W441 | 276 | T | 1.766 | 0.003248106 | Up | Tetraspanin OS=Drosophila melanogaster OX=7227 GN=Tsp5D PE=2 SV=4 | Tsp5D |
| X2JAA8 | 336 | T | 1.31 | 0.000438633 | Up | Wapl-RA OS=Drosophila melanogaster OX=7227 GN=wapl PE=1 SV=1 | wapl |
| Q9Y095 | 399 | T | 1.35 | 0.000978054 | Up | FI04011p OS=Drosophila melanogaster OX=7227 GN=XRCC1 PE=1 SV=1 | XRCC1 |
| Q9Y095 | 566 | T | 1.561 | 0.018544602 | Up | FI04011p OS=Drosophila melanogaster OX=7227 GN=XRCC1 PE=1 SV=1 | XRCC1 |
| X2JAM4 | 913 | T | 6.009 | 1.69623E-05 | Up | Supervillin, isoform AE OS=Drosophila melanogaster OX=7227 GN=Svil PE=1 SV=1 | Svil |
| X2JAM4 | 1299 | T | 1.321 | 0.000838616 | Up | Supervillin, isoform AE OS=Drosophila melanogaster OX=7227 GN=Svil PE=1 SV=1 | Svil |
| X2JAU8 | 480 | T | 0.243 | 1.85133E-06 | Down | Protein nervous wreck OS=Drosophila melanogaster OX=7227 GN=nwk PE=1 SV=1 | nwk |
| X2JF15 | 120 | T | 1.587 | 0.007465326 | Up | Radish, isoform L OS=Drosophila melanogaster OX=7227 GN=rad PE=4 SV=1 | rad |
| A0A0B4JDF4 | 538 | Y | 3.052 | 5.74315E-06 | Up | Protein tyrosine phosphatase 99A, isoform G OS=Drosophila melanogaster OX=7227 GN=Ptp99A PE=4 SV=1 | Ptp99A |
| A0A0B4KGI7 | 704 | Y | 2.041 | 0.040917308 | Up | Signal transducer and activator of transcription OS=Drosophila melanogaster OX=7227 GN=Stat92E PE=3 SV=1 | Stat92E |
| A0A0B4LGS4 | 112 | Y | 1.484 | 0.00096826 | Up | Vacuolar H[+]-ATPase 26kD subunit, isoform C OS=Drosophila melanogaster OX=7227 GN=Vha26 PE=3 SV=1 | Vha26 |
| M9PCU1 | 1381 | Y | 2.404 | 0.014218475 | Up | Abl tyrosine kinase, isoform H OS=Drosophila melanogaster OX=7227 GN=Abl PE=1 SV=1 | Abl |
| A8JQW5 | 36 | Y | 8.494 | 0.0001242 | Up | Uncharacterized protein OS=Drosophila melanogaster OX=7227 GN=Dmel\CG12947 PE=4 SV=1 | Dmel\CG12947 |
| A8JUQ9 | 695 | Y | 2.451 | 0.000220513 | Up | Vav guanine nucleotide exchange factor, isoform C OS=Drosophila melanogaster OX=7227 GN=Vav PE=1 SV=1 | Vav |
| A8JV07 | 664 | Y | 3.889 | 0.000535869 | Up | Uncharacterized protein, isoform B OS=Drosophila melanogaster OX=7227 GN=Dmel\CG9915 PE=1 SV=1 | Dmel\CG9915 |
| E1JHB6 | 1380 | Y | 2.138 | 0.001456469 | Up | Receptor protein-tyrosine kinase OS=Drosophila melanogaster OX=7227 GN=Pvr PE=1 SV=1 | Pvr |
| E1JHD6 | 183 | Y | 1.817 | 0.008108348 | Up | Mitogen-activated protein kinase OS=Drosophila melanogaster OX=7227 GN=bsk PE=1 SV=1 | bsk |
| E8NH67 | 56 | Y | 1.411 | 0.008677021 | Up | GH07746p OS=Drosophila melanogaster OX=7227 GN=CG9572-RB PE=2 SV=1 | CG9572-RB |
| Q95RW2 | 185 | Y | 1.584 | 0.001596248 | Up | Crk oncogene, isoform B OS=Drosophila melanogaster OX=7227 GN=Crk PE=1 SV=1 | Crk |
| Q9VNP2 | 1025 | Y | 1.309 | 0.006774414 | Up | Neuromusculin, isoform A OS=Drosophila melanogaster OX=7227 GN=nrm PE=1 SV=4 | nrm |
| O61443 | 185 | Y | 1.425 | 0.006931912 | Up | Mitogen-activated protein kinase p38b OS=Drosophila melanogaster OX=7227 GN=p38b PE=1 SV=1 | p38b |
| P40417 | 200 | Y | 1.665 | 0.028273004 | Up | Mitogen-activated protein kinase ERK-A OS=Drosophila melanogaster OX=7227 GN=rl PE=1 SV=3 | rl |
| Q95SH7 | 302 | Y | 1.632 | 0.001634181 | Up | GH26007p OS=Drosophila melanogaster OX=7227 GN=Dmel\CG14969 PE=1 SV=1 | Dmel\CG14969 |
